# Prevalence and polymorphism of a mussel transmissible cancer in Europe

**DOI:** 10.1101/2021.03.31.436483

**Authors:** Maurine Hammel, Alexis Simon, Christine Arbiol, Antonio Villalba, Erika AV Burioli, Jean-François Pépin, Jean-Baptiste Lamy, Abdellah Benabdelmouna, Ismael Bernard, Maryline Houssin, Guillaume M Charrière, Delphine Destoumieux-Garzon, John Welch, Michael J Metzger, Nicolas Bierne

## Abstract

Transmissible cancers are parasitic malignant cell lineages that acquired the ability to infect new hosts from the same species, or sometimes related species. First described in dogs and Tasmanian devils, transmissible cancers were later discovered in some marine bivalves affected by a leukemia-like disease. In *Mytilus* mussels, two lineages of Bivalve Transmissible Neoplasia (BTN), both emerged in a *M. trossulus* founder individual, have been described to date (MtrBTN1 and MtrBTN2). Here, we performed an extensive screening of genetic chimerism, a hallmark of transmissible cancer, by genotyping hundred SNPs of thousands of European *Mytilus* mussels. The genetic analysis allowed us to simultaneously obtain the genotype of hosts *-M. edulis*, *M. galloprovincialis* or hybrids- and the genotype of tumors of heavily infected individuals. In addition, a subset of individuals were systematically genotyped and analysed by histology in order to screen for possible non-transmissible cancers. We detected MtrBTN2 at low prevalence in *M. edulis*, and also in *M. galloprovincialis* and hybrids although at a much lower prevalence. No MtrBTN1 or new BTN were found but a few individuals with non-transmissible neoplasia were observed at a single polluted site on the same sampling date. We observed a diversity of MtrBTN2 genotypes that appeared more introgressed or more ancestral than MtrBTN1 and reference healthy *M. trossulus* individuals. The observed polymorphism is most likely due to somatic null alleles caused by structural variations or point mutations in primer-binding sites leading to enhanced detection of the host alleles. Despite low prevalence, two divergent sublineages, confirmed by mtCOI sequences, are co-spreading in the same geographic area, suggesting a complex diversification of MtrBTN2 since its emergence and host species shift.

## 1. Introduction

Transmissible cancers are malignant cell lines, derived from a founder host, which can spread and colonize other individuals by transmission of living cancer cells. Several lineages of transmissible cancer have been described: one in dogs (Canine Transmissible Venereal Tumor, CTVT, Murgia et al. 2006, Rebbeck et al. 2009), two in Tasmanian devils (Devil Facial Tumor Disease, DFT1 and 2, Pearse et al. 2006, Pye et al. 2016) and, more recently, six in different bivalve species (Bivalve Transmissible Neoplasia, BTN, Metzger et al. 2015, 2016, Yonemitsu et al. 2019). The three lineages found in vertebrates are the best studied to date. In dogs, CTVT arose about 4000-8000 ybp and spread worldwide (Leathlobhair et al. 2018, Baez-Ortega et al. 2019). In Tasmanian devils two independent transmissible cancers lineages have emerged in the last 30 years. In bother cases, although at different spatio-temporal scales, genome studies have shown early diversification into sublineages and secondary co-existence of some divergent sublineages (Baez-Ortega et al. 2019, Kwon et al. 2020; Patton et al. 2020). Studies in marine bivalves also showed multiple emergences (two in *Cerastoderma edule*, Metzger et al. 2016 and two in *Mytilus trossulus*, Yonemitsu et al. 2019) and for the first time in transmissible cancer research, transmissions crossing the species barrier (BTN lineage in *Polititapes* aureus has a *Venerupis corrugata* origin, Metzger et al. 2016, and MtrBTN2 in *M. edulis* and *M. chilensis* has a *M. trossulus* origin, Yonemitsu et al. 2019).

The *Mytilus edulis* complex of species is composed of three species in the northern hemisphere: *M. edulis*, *M. galloprovincialis* and *M. trossulus*, and three species in the southern hemisphere: *M. chilensis*, *M. platensis* and *M. planulatus*. They are incompletely reproductively isolated and can hybridize when they come into contact either naturally or via human-induced introductions (Fraisse et al. 2016, Simon et al. 2020, Popovic et al. 2020). Disseminated neoplasia (DN) has been previously observed in four of this species around the world: *M. trossulus* (Rasmussen 1986, Sunila 1987, Moore et al. 1991, Elston et al. 1992, Usheva and Frolova 2000, Vassilenko and Baldwin 2014), *M. chilensis* (Campalans et al. 1998, Cremonte et al. 2015, Lohrmann et al. 2019), *M. edulis* (Farley 1969, Lowe and Moore 1978, Green and Alderman 1983, Elston et al. 1988, Galimany and Sunila 2008) and *M. galloprovincialis* (Figueras et al. 1991, Villalba et al. 1997, Carrasco et al. 2008, Gombac et al. 2013, Carella et al. 2013, Matozzo et al. 2018). Cancer cells are characterised by rounded, basophilic and polyploid cells with high nucleus to cell ratio (for review see Carballal et al. 2015) and their presence is determined by histological observation, and cytology or flow cytometry of hemolymph (Benabdelmouna and Ledu 2016, Benabdelmouna et al. 2018, Burioli et al. 2019). However, in order to demonstrate that a cancer is transmissible, a genetic study is required in order to identify (i) genetic differences between tumor and host cells and (ii) genetic similarity between tumors of different individuals (hallmark of transmissible cancer). Thus, two independent lineages of BTN from two *M. trossulus* founder mussels have been described in three mussel species: one lineage (MtrBTN1) found in East Pacific *M. trossulus* populations, and the other lineage (MtrBTN2) with a worldwide distribution in West Pacific *M. trossulus* populations, European *M. edulis* and South American *M. chilensis* hosts (Metzger et al. 2016, Yonemitsu et al. 2019, Skazina et al. 2020). MtrBTN1 was discovered in *M. trossulus* populations in Western Canada by genetic similarity of neoplastic hemocytes on a mitochondrial and a nuclear marker (Metzger et al. 2016). Although disseminated neoplasia was first described 50 years ago in *Mytilus sp*. (Farley 1969) we do not know if all reports of disseminated neoplasia correspond to a transmissible cancer. A population genetic study of French mussels detected chimeric individuals with an abnormal presence of *M. trossulus* alleles in *M. edulis* populations devoid of *M. trossulus* individuals, leading the authors to hypothesize the presence of a *M. trossulus* transmissible cancer (Riquet et al. 2017). The presence of *M. trossulus* transmissible cancer in *M. edulis* was later confirmed by the detection of a *M. trossulus* genetic signal in the hemolymph of cancerous *M. edulis* grown in Brittany (Burioli et al. 2019). By comparing two nuclear and two mitochondrial markers from MtrBTN1 tumors, and tumors found in South-American *M. chilensis* and European *M. edulis*, Yonetmistu et *al*. (2019) showed the latter are an independent emergence, called MtrBTN2. One mitochondrial marker indicated that MtrBTN2 from two *M. chilensis* mussels have the *M. chilensis*-derived haplotype indicating that recombination between host and cancer mitochondria might occur in this cancer lineage (Yonetmistu et al. 2019). Very recently, MtrBTN2 has been observed in *M. trossulus* hosts in the Sea of Japan (Skazina et al. 2020) with divergent mtDNA sequences and a lower ploïdy (~4N) than the hyperploids MtrBTN2 found in *M. edulis* (>8N, Burioli et al. 2019).

Here, we performed a large genetic screening of European mussels (*M. edulis*, *M. galloprovincialis* and their hybrids) in order to detect genetic evidence of transmissible cancer (chimerism) of any origin and compare the prevalence between species. For this study we performed a dedicated sampling and also reanalysed data from previous population genetics studies. We finally inferred the genotypes of tumors of the most cancerous individuals and studied polymorphism of these tumors. Most candidate chimeric individuals were subsequently observed in histology, cytology or flow cytometry in order to obtain a direct validation that the chimerism was the consequence of a transmissible cancer. Because disseminated neoplasia could also be non-transmissible, a subsample of 222 mussels was systematically inspected by histology for the presence of neoplasia before genetic screening. MtrBTN2 was found at a low prevalence in Europe, mostly in *M. edulis* and also in *M. galloprovincialis* and their hybrids. Some neoplastic mussels did not show evidence of genetic chimerism suggesting that conventional non-transmissible cancers were also present. Finally, the study of MtrBTN2 genotypes from the 6 most infected *M. edulis* hosts revealed polymorphism and allow to describe two divergent sub-lineages that are co-spreading in French populations.

## 2. Material and methods

### a. Sampling and genotyping

We analysed a total of 5907 European mussels (*Mytilus edulis*, *M. galloprovincialis*, and some hybrids, subsequently classified with SNPs, see below), collected between 2005 and 2019 (Figure 1, Table S1). For the purpose of this study n = 1749 mussels were sampled on two tissues: hemolymph (higher amount of cancer cells, increasing with cancer severity) and mantle (higher amount of host cells, decreasing with cancer severity). We also analysed n = 4158 mussels sampled for population genetics studies at only one tissue (gill or hemolymph). For comparison, we also analysed two known MtrBTN1 samples and their *M. trossulus* hosts (described in Metzger et al. 2016), and 32 *M. trossulus* (10 from British Columbia and 10 from Saint-Laurent, Canada and 8 from the Baltic Sea, Gdansk, Poland) (TableS1).

**Fig. 1:**
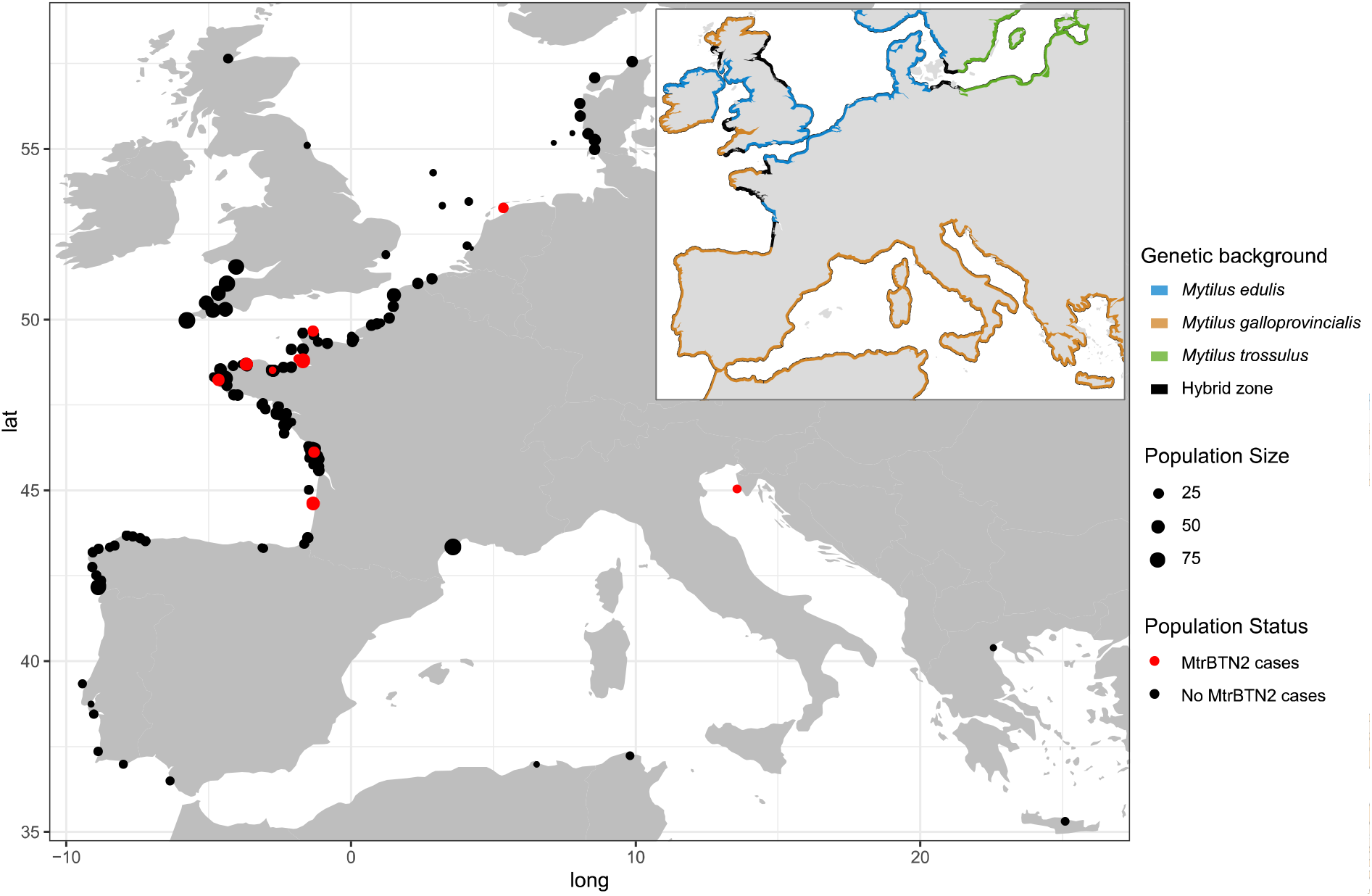
Locations of sampling sites and distribution of genetic lineages of *Mytilus* mussels in Europe. Point size indicates sampling size and color indicates the absence (black) or presence (red) of MtrBTN2 cancer. Coastline colors represent the different genetic lineages: edulis (blue), *M. galloprovincialis* (orange), *M. trossulus* (green) and hybrid zone (black). All the information about sites is reported in Table S1.

Solid tissues (mantle and gill) were either fixed in 96% ethanol or directly used for DNA extraction. For hemolymph samples, collected from adductor muscle (1ml syringe, 26G needle), cells were centrifuged (10 min at 2500g) and the supernatant removed and replaced by 96% ethanol. DNA extraction was done with the Nucleomag^TM^ 96 Tissue kit (Macherey-Nagel) in combination with a Kingfisher Flex^TM^ extraction robot (serial number 711-920, ThermoFisher Scientific). We followed the kit protocol with modified volumes of some reagents: beads 2x diluted in water, 200μL of MB3 and MB4, 300μL of MB5 and 100μL of MB6.

A total of 106 (103 nuclear and 3 mitochondrial) biallelic single nucleotide polymorphisms (SNPs) described in Simon et al. 2020 were analysed. However 29 nuclear and 1 mitochondrial SNPs produced too much missing data and unsatisfactorily fluorescennce leve and were removed for subsequent analyses. The list of 76 SNPs used and their sequences are provided in Table S2. They were identified to be strongly differentiated between species and lineages of the *Mytilus edulis* complex of species in the study of Fraisse et al. (2016), in order to develop an ancestry-informative panel of SNPs, and its efficacy was confirmed by the analysis of large datasets (Simon et al. 2020, 2021). Genotyping was subcontracted to LGC genomics (Hoddensdon, UK) and performed with the KASP^TM^ method (Cuppen, 2007, Semagn et *al*. 2014). This method puts two “allele specific” primers with two different fluorochromes in competition during a PCR amplification and gives two values of fluorescence level, one for allele X and the other for allele Y. Homozygous genotypes have a high fluorescence for allele X or allele Y only, while heterozygous genotypes have an intermediate fluorescence for each allele (Figure 2). The genotype classification at each SNP of each sample was determined by two LGC genomics experts. Assays with too low fluorescence were classified as badly amplified (“Bad”) and those with an unbalanced intermediate fluorescence that was neither compatible with a homozygous nor with a heterozygous genotype of a standard diploid genotype were classified as “uncalled” (Figure 2). In order to assign mussels to species (*M. edulis*, *M. galloprovincialis* or hybrid), we performed a Principal Component Analysis (PCA, dudi.pca(), package adegenet in R, Jombart et al. 2011) based on the 74 nuclear SNPs and used the first axis coordinate to infer a hybrid index (Väinölä and Strelkov 2011; Brisbin et al. 2012).

**Fig. 2:**
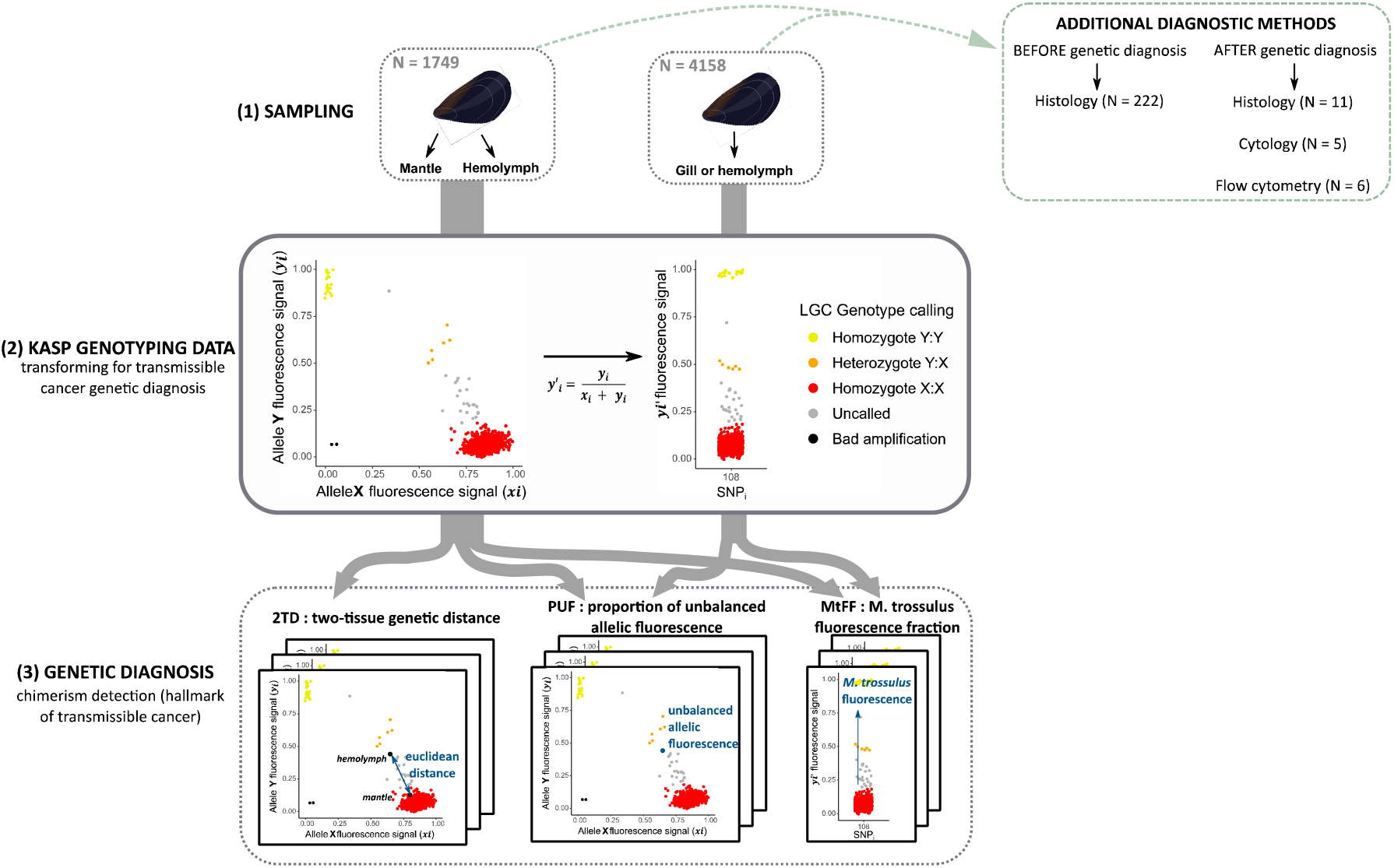
Experimental design diagram, from sampling to genetic diagnosis. (1) Mussel sampling. 1749 individuals double sampled on hemolymph and mantle and 4158 individuals only sampled on gill or hemolymph. The additional diagnostic methods used are reported in the green box (see Table S3). (2) Transformation of KASP genotyping data for transmissible cancer detection. Example of KASP fluorescence data plot (left) and the transformed fluorescence value y’_i_ for the bi-allelic SNP “108” (right). This SNP is diagnostic and allele Y (y_i_) corresponds to the most frequent allele in *M. trossulus* reference samples (Table S2). Thus, homozygotes X:X (red) are *M. edulis*-*M. galloprovincialis* individuals while homozygotes Y:Y (yellow) are *M. trossulus* individuals. Heterozygotes Y:X (orange) are introgressed genotypes, healthy individuals with a *trossulus*-state allele at a locus due to shared polymorphism. Uncalled (grey) are *M. edulis*-*M. galloprovincialis* hosts with an excess of *M. trossulus* flurorescence likely due to *M. trossulus* transmissible cancer. Poor amplification (black) were removed from the analysis. (3) Genetic diagnosis, 3 indexes to detect genetic chimerism. Two-tissue genetic distance (2TD) calculated for double sampled mussels (n=1749). Proportion of unbalanced allelic fluorescence (PUF). *M. trossulus* fluorescence fraction (MtFF) computed from 10 diagnostic SNPs.

### b. Transforming KASP genotyping data for transmissible cancer genetic diagnostics

In contrast to healthy mussels, mussels affected by transmissible cancer are chimeric, with their infected tissues composed of a mix of neoplastic and host cells, usually with a proportion deviating from 50:50 (a minority of cancerous cells mainly in the hemolymph at the earlier stages of the disease, and a majority of cancerous cells in every tissue at the later stages but usually with a greater proportion of cancer cells in the hemolymph). As a consequence, the KASP^TM^ amplification is likely to result in an unbalanced amplification of the two alleles at SNPs for which the genotype is different between cancerous and host cells. Such deviations from standard diploid genotype callings are likely to result in the genotype to be classified as “uncalled” (Figure 2). Consequently, we chose to use allele fluorescence values directly rather than genotype calling. To be able to compare samples from independent genotyping experiments, we first had to standardize the X and Y fluorescence values to have all of them in a range from 0 to 1 by using the following formulas within each experimental plate: ***x_i,j_ = (X_i,j_ - min(X_i_))/(max(X_i_)-min(X_i_)) and y_i,j_ = (Y_i,j_ - min(Y_i_))/(max(Y_i_)-min(Y_i_))***, where *X_i,j_* and *Y_i,j_* are the allele X and allele Y fluorescence values respectively for the *ith* SNP of *jth* sample. Then, following Cuenca et al. (2013) method for genotyping polyploid samples, we transformed the data to have a measure of the proportion of allele Y fluorescence at each SNP, with a value going from 0 when allele X (*x_i,j_*) fluorescence dominates to 1 when allele Y (*y_i,j_*) fluorescence dominates (Figure 2). Furthermore, SNP alleles were oriented in order for allele Y to be the most frequent in *M. trossulus* reference samples (see Table S2). To do this, we used the following formula ***y’_i,j_ = *Y_i,j_* /(*X_i,j_* + *Y_i,j_*)*** where *y_i_* and *x_i_* are the fluorescence values of allele X and Y respectively for the *ith* SNP of *jth* sample (Figure 2). We called *y’_i,j_* the “Variant Allele Fluorescence Fraction (VAFF)” (it is similar to the Variant Allele Fraction statistics used with NGS dataset where the fluorescence signal is used instead of the read coverage).

To filter the data out from badly amplified low fluorescence level SNPs and samples, we calculated an index of amplification following this formula ***a_i,j_ = X_i,j_ + Y_i,j_***, where *X_i,j_* and *Y_i,j_* are the allele X and allele Y original fluorescence values. Fluorescence levels were removed from the dataset when ***_i,j_*** was below 0.3. Samples with an average index of amplification below 0.8 were also removed.

### c. Genetic diagnosis: 3 indexes to detect chimerism

We developed two indexes to identify chimeric mussels (a hallmark of transmissible cancer), and a third one designed to specifically identify chimera with an excess of *M. trossulus* allele fluorescence (Figure2).

When two tissues of the same individual were sampled we were able to investigate genetic differences between the two tissue samples within individuals. We define the two-tissue genetic distance (2TD) as the euclidean distance between the hemolymph and the mantle based on allele fluorescences ***x_i_*** and ***y_i_*** (*dist()*, package *stats* in R, R Core Team 2020) (Figure 2). Chimerism was considered detected when 2TD was above 1.54 (highest percentile, see results). We expected a higher proportion of cancer cells in the hemolymph than in hard tissues, at least in the early stage of the disease. However, we noticed that for more infected individuals, chimerism was observed in both tissues with sometimes little differences in fluorescence rates between hemolymph and mantle. We therefore developed a second index to identify fluorescence rates that deviate from a balanced amplification of the two alleles. The proportion of unbalanced allelic fluorescence (PUF) was computed as the proportion of SNPs classified as “uncalled” in an individual tissue (classification performed by two LGC experts) (Figure2). For individuals genotyped at two tissues we obtained two values of PUFs, one for each tissue. Chimerism was considered detected when PUF was above 0.20 (highest percentile, see results). Because only MtrBTN2 has been found in Europe to date (Riquet et al. 2017, Yonemitsu et al. 2019, Burioli et al. 2019) and as MtrBTN2 originated in a *M. trossulus* host, we computed a “*M. trossulus* fluorescence fraction” index (MtFF) to specifically detect *M. trossulus* chimerism (Figure2). We selected 10 SNPs of the array that are diagnostic between *M. trossulus* and both *M. edulis* and *M. galloprovincialis* (allele Y fixed or nearly fixed in *M. trossulus* populations and absent or very rare in *M. edulis* and *M. galloprovincialis* populations, see Table S2). We computed the average VAFF *(y’_i,j_)* at these 10 SNPs in order to detect samples with abnormal excess of *M. trossulus* fluorescence. Remember that *M. trossulus* is absent from the areas investigated in this study. As for PUFs, we obtained two values of MtFF for individuals genotyped at two tissues. We considered the *M. trossulus* fluorescence signal as positive when MtFF was above 0.22 (highest percentile, see results).

Because apparent chimerism can result from experimental artifacts rather than to transmissible cancer, we inferred a clustering tree to determine if positive individuals grouped together, as expected for a clonally propagating transmissible cancer, but not for independent contaminations. We used the Neighbor-Joining method (*nj()*, package *ape* in R, Paradis and Schliep 2018) based on Euclidean distance of VAFF value (*y’_i,j_*).

### d. Method evaluation: analysis of in silico chimera

We evaluated our ability to detect non-*trossulus* chimerism, and conversely our ability to conclude that neoplasia are non-transmissible, using the 2TD and PUF statistics by computing the expected distribution of in-silico generated chimera with an increasing fraction of cancer cells in the sample. For the 2TD statistic we first had to estimate the allele fluorescence of chimera (chim). We combined the fluorescence values of two randomly selected hemolymph samples (“host” and “cancer”) following this formulas : ***x_i,chim_* = (1-F)\**x_i,host_* + F\**x_i,cancer_*** and ***y_i,chim_* = (1-F)\**y_i,host_* + F\**y_i,cancer_***, where *x_i_* and *y_i_* are the allele X and allele Y fluorescence values respectively for the *ith* SNP of the “host” and “cancer” samples and F the fraction of cancer cells. Then, we calculated the 2TD as the euclidean distance between the computed “chimera” hemolymph sample and the “host” mantle sample (as described previously). For the PUF statistic we classified as “uncalled” each SNPs for which the “host” and “cancer” hemolymph samples had different genotypes calling and calculated the proportion of “uncalled” SNPs for each chimera (as described previously). Because transmissible cancer can occur at intraspecific and interspecific levels, we generated different categories of in silico chimera by selecting “host” and “cancer” samples from the same or different species (*M. edulis* - *M. edulis*, *M. edulis* - *M. galloprovincialis*, *M. edulis* - *M. trossulus*, *M. galloprovincialis* - *M. galloprovincialis* and *M. galloprovincialis* - *M. trossulus*).

### e. Additional diagnosis methods: histology, cytology and flow cytometry

A total of n = 222 mussels have been screened for neoplasia using histology (Figure2, Table S3) before genetic diagnosis. Shells were opened by cutting the adductor muscle and the soft tissues were removed. An approximately 5 mm thick transverse section of mussel tissue containing mantle lobes, visceral mass (gut, digestive gland) and gills was excised, fixed in Davidson’s fixative for 48h and then preserved in 70 % ethanol. Fixed tissues were dehydrated through an ascending ethanol series and embedded in paraffin wax. Five μm thick sections were produced with a rotary microtome and stained with Harris’ hematoxylin and eosin (HHE) (Howard et al. 2004). Histological sections were examined with a light microscope Olympus BX50.

In addition to genetic diagnosis, n = 17 mussels have been investigated by histology (n = 11), cytology (n = 5) and/or flow cytometry (n = 6) after the genetic screening. The methods used are referenced in Table S3.

### f. Inference of cancer genotypes and their genetic relationship

Determining the genotype of a cancer lineage is complicated by the presence of host cells. However, 6 samples exhibited a MtFF close to our *M. trossulus* reference samples consistent with them being composed of a vast majority of cancerous cells (up to 90%). We used these 6 samples to infer the genotypes of tumors. In order to infer genotypes, we used the x/y fluorescence plots as well as the correlation between fluorescence rate at individual SNP and MtFF (Figure S1). As we were working with *M. edulis* hosts only (see results) we used a new *M. trossulus* fluorescence fraction (MtFF2) with an extended number of 17 *M. edulis* - *M. trossulus* diagnostic SNPs (Table S2). MtFF2 was used as a proxy of the proportion of cancerous cells in the sample (see Figure S1). Genotype inference was performed independently by two users and cross-checked.

To have a global comparison of cancerous genotypes with *Mytilus* references, we performed a maximum parsimony (MP) tree using MEGA v7.0.26 (Kumar, Stecher and Tumura 2016) with all sites and Subtree-Pruning-Regrafting method. SNP genotypes were coded with two nucleotides per position (a single SNP was represented by two positions in a “pseudo-fasta” file) to be able to represent heterozygous and homozygous genotypes and then we combined all SNPs in a single “pseudo-sequence” for each sample before converting to fasta format (*as.fasta()* package *bio3d* v2.4-1 in R, Grant et al. 2006).

### g. mtCOI sequences

Individuals STBRI_69, L19_8, L9_3, L11_4, CamaretB_1, Chausey5, Hol8 and Barf30 affected by MtrBTN2, were sequenced for a parallel research project and short reads produced by Illumina sequencing were available (Novaseq, paired-end 150 bp). We skimmed mitochondrial reads by mapping them to a set of eight *Mytilus* mitogenomes, using bwa-mem2 (v2.0, Md et al. 2019) and SAMtools (v1.10, Li et al. 2009) with a minimal mapping quality of 5. The mitochondrion genomes from female and male *M. galloprovincialis* (FJ890849 and FJ890850, respectively), *M. edulis* (DQ198231 and AY823623, respectively), *M. trossulus* (AY823625 and HM462081, respectively), and M. californianus (JX486124 and JX486123, respectively) were retrieved from GenBank and used as references. Note that the *M. edulis* female mitochondrion is attributed to a *M. trossulus* individual in GenBank but corresponds to an introgression of the *M. edulis* mitochondrion in the Baltic sea. For each cancerous individual, mitochondrial reads were then assembled with MEGAHIT (v1.2.9, Li et al. 2016). Contigs produced with megahit were finally processed with MitoFinder (Allio et al. 2020) to automatically retrieve contigs corresponding to mitochondria and annotate them (using the invertebrate mitochondrial code as organism type). Then, we extracted COI sequences from all mitochondrial produced contigs for each individual, and aligned them with sequences of Yonemitsu et al. (2019), Skazina et al. (2020) as well as *M. trossulus* mtCOI sequences gathered from Genbank (Riquet et al. 2017). We also extracted COI sequences from the mitogenome of Mtx10A, a 100,000 year old *M. trossulus* mussel (Der Sarkissian et al. 2020, European Nucleotides Archive SAMEA6367443). Finally, we performed a maximum parsimony (MP) tree using MEGA v7.0.26 (Kumar, Stecher and Tumura 2016) with all sites and Subtree-Pruning-Regrafting method.

## 3. Results

### a. Transmissible cancer genetic diagnosis of individuals sampled on two tissues

We first analysed samples of two tissues from n = 1749 mussels in order to evaluate the prevalence of genetic chimerism in this dataset and the ability of our indexes to identify mussels infected by a transmissible cancer either of *M. trossulus* origin or other *Mytilus* spp origins.

We found 11 positive MtFF outlier individuals (*M. trossulus* fluorescence fraction), 5 for both tissues and 6 for hemolymph only (Figure 3A). Ten of them were also positive at both PUF (proportion of unbalanced allelic fluorescence, 5 for both tissues and 5 for hemolymph only) and 2TD (2-tissue genetic distance), and the last one was positive at PUF but not 2TD (Figure S2). MtFF values were higher in the hemolymph samples than the mantle samples but the difference was sometimes low. Those 11 individuals are likely infected by the MtrBTN2 lineage previously described in these populations (Yonemitsu et al. 2019). Ten of these 11 mussels were also investigated by at least one of the additional diagnostic methods (histology, cytology and/or flow cytometry), which confirmed the presence of cancer cells for all of them (Figure 3A, red triangle up, Table S2). Moreover, the highest MtFF values were found in heavily infected individuals as also reported by Burioli et al. (2019) (Table S2). The individual that was negative at 2TD (L11_4) was equally infected in both tissues. Finally, we used the MtrBTN2 specific qPCR diagnosis of Yonemitsu et al. (2019) and found the 11 samples to be positive.

**Fig. 3:**
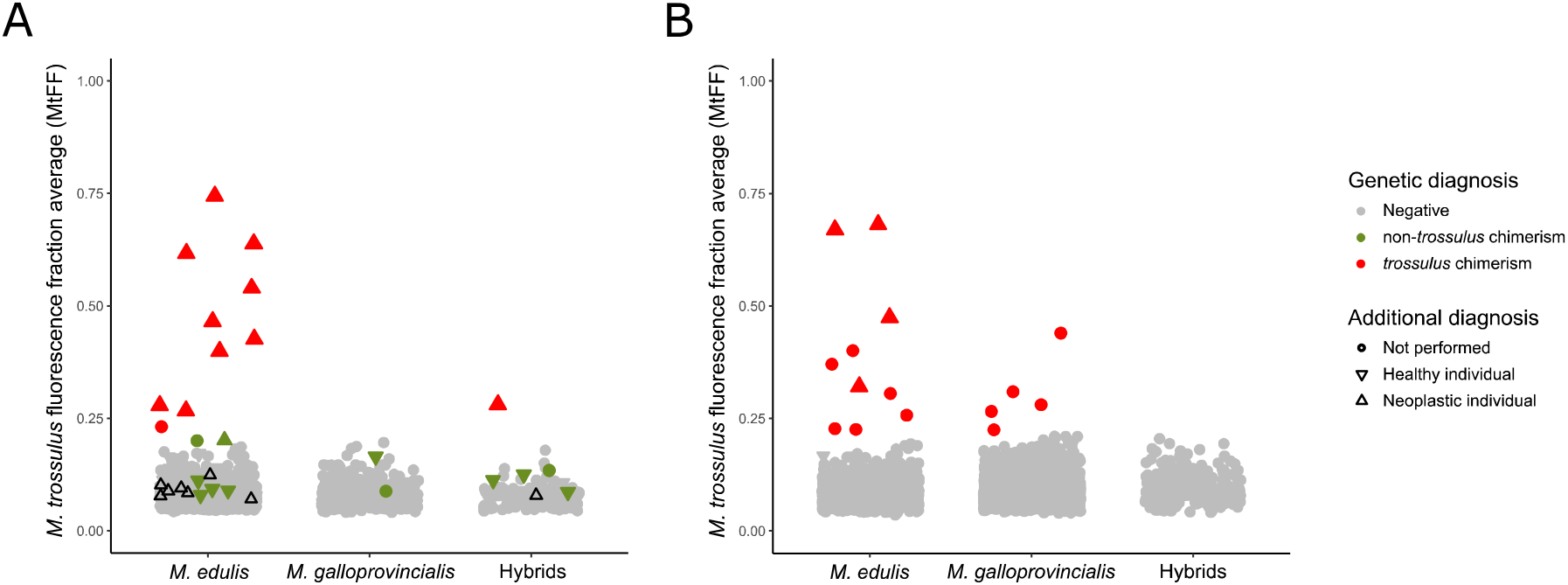
*M. trossulus* Fluorescence Fraction (MtFF) used for genetic diagnosis. (A) MtFF values for individuals sampled at two tissues. Color correspond to genetic diagnosis: red for trossulus chimeric individuals (positive for MtFF, PUF and 2TD), green for non-*trossulus* chimeric individuals (positive for PUF and 2TD indexes) and grey for non-chimeric individuals (negative to all indexes). Black triangles pointing up represent neoplastic individuals of type B which are not chimeric (negative to all indexes). (B) Single-sampled individuals for which genetic diagnosis correspond to MtFF value: red for trossulus chimerism (positive to MtFF) and grey for non-chimeric individuals (negative to MtFF). (A, B) The symbol form corresponds to additional diagnosis methods (histology, cytology or flow cytometry): triangles pointing up for neoplastic individuals, triangles pointing down for healthy individuals and dot for not performed (see TableS3).

We also found 12 individuals that were negative at MtFF but positive at both PUF and 2TD. Those individuals could be candidates for a new transmissible cancer that emerged in *M. edulis* or *M. galloprovincialis*. We obtained additional diagnostic information (histology, cytology and/or flow cytometry) for 9 of these 12 mussels and did not find any evidence of the presence of cancer cells for 8 of them, suggesting the chimerism is not related to transmissible cancer but most probably to experimental artifacts such as contamination. Only one individual (CamaretA_61) revealed the possible presence of a few cancer cells (Figure 3A, green triangle up, Table S3). Although the MtFF of this individual was not high enough to belong to outliers, the MtFF of the hemolymph sample was nonetheless high, close to the detection limit (Figure 3A, green triangle up). This suggests that it could be the *M. trossulus* cancer found in other mussels, likely MtrBTN2, at a very early stage of the disease.

To further investigate the origin of non-*trossulus* chimerism we analysed the genetic similarity of samples with a cluster tree reconstruction based on VAFF Euclidean distances between individuals. All *M. trossulus* chimeric samples grouped together (Figure S3, red points) in a cluster related to *M. trossulus* reference samples. Conversely, non-trossulus chimeric samples were sparsely distributed in the tree. Two exceptions (CameretA_61 and AUN_001_13) to this were the two samples with MtFF close to the limit and with few cancer cells for CamaretA_61, which clusters with the *M. trossulus* chimeric group. This suggests once again that they are likely early stage of the MtrBTN2 disease. Although two groups of 4 and 2 samples tended to cluster together, they were respectively sampled in the same population which was a sympatric population where *M. edulis* and *M. galloprovincialis* coexist and a contamination can easily resemble a hybrid genotype. However, the argument that non-trossulus chimeric samples do not tend to cluster together in the tree is not necessarily strong because the fluorescence signal of non-trossulus BTN cells might not be sufficient to outshine the signal of host cells. In this case PUF and 2TD statistics would not have been found with outlier values though. Our analysis of in-silico chimera with variable proportions of mixing between two individuals (FigureS4), gives an estimation of infection rate needed to detect a BTN (over 80% for a *M. edulis* BTN in *M. edulis* host, over 50% for a *M. galloprovincialis* BTN in *M. galloprovincialis* host and over 30% for an interspecific spreading). With such proportions of cancer cells chimeric samples would have tended to cluster together as illustrated by in-silico chimera with mixtures of 40, 60 and 80% added in the tree (see Figure S4, orange branches). Above all, we would have observed neoplastic cells by histological examination. We therefore conclude that non-trossulus chimerism was most likely due to technical artifacts.

### b. screening of MtrBTN2 in individuals sampled on one tissue only

As we obtained satisfactory results of MtrBTN2 diagnostic with the *M. trossulus* fluorescence fraction (MtFF), even with mantle samples, and that there is very little evidence for another transmissible cancer, we decided to investigate genotyping data previously obtained for population genetic studies from DNA extraction of gills (n = 4154) and hemolymph (n = 4). This allowed us to substantially increase our sample size from 1749 to 5907 mussels. The MtFF index allowed us to identify 15 individuals with an excess of *M. trossulus* fluorescence (Figure 3B, red point). Chimerism was confirmed by a high PUF signal for 12 of them (Figure S2). The 3 others had a very low PUF and correspond to MtFF values close to the limit. We considered as candidates only the 12 positives for both MtFF and PUF. In addition, we had additional diagnostic information for 4 of them which all proved to be infected by neoplastic cells (Figure 3B, red triangle up, Table S3).

### c. Host-specific prevalence of MtrBTN2 in Europe

We classified individual mussels as *M. edulis*, *M. galloprovincialis* or hybrid using a PCA on all SNP VAFF values (*Y_i,j_*’). The first axis of the PCA separates *M. edulis* and *M. galloprovincialis* individuals, with hybrids in-between, while the second axis separates these two species and *M. trossulus* (Figure S5). We used the first axis coordinate to infer a hybrid index (HI: Väinölä and Strelkov 2011; Brisbin et al. 2012) and arbitrarily assign mussels to *M. edulis* (HI>0.85), *M. galloprovincialis* (HI < 0.4), and hybrids (0.4 < HI < 0.85), given that asymmetric introgression generates a larger variance of *M. edulis* ancestry in *M. galloprovincialis* than the variance of *M. galloprovincialis* ancestry in *M. edulis*. The overall prevalence of MtrBTN2 was low (25/5907 = 0.42%) but was significantly higher in *M. edulis* hosts (22/2808 = 0.78%) than in hybrids and *M. galloprovincialis* hosts (1/381 = 0.26% and 2/2718 = 0.07%, respectively - Fisher’s exact test p=0.0002).

### d. Non-transmissible cancers

For 222 individuals we conducted histological inspection before genetic diagnosis. One individual L9_3 was highly infected by large neoplastic cells referred to as type A neoplasia (see micrographs A and B in Figure 4 and description below). This individual was positive at the three genetic diagnostics indexes (MtFF, PUF and 2TD) and was infected by MtrBTN2. Eight individuals from batch L6 were affected by a type of neoplasia characterised by neoplastic cells with smaller size than the type A and referred as type B neoplasia (see micrographs C and D in Figure 4). Both disseminated neoplasia type A and type B involved the proliferation of abnormally large cells through the connective tissue and hemolymph vessels of different organs. The main morphological character for discrimination between both neoplasia types when examining histological sections was the nuclear size, significantly larger in type A (mean diameter ± SE = 7.83 ± 0.136 μm; range: 6.0 – 9.9 μm; N = 31) than in type B (mean diameter ± SE = 4.56 ± 0.096 μm; range: 3.6 – 5.1 μm; N = 40); both cell types showed scant cytoplasm (high N/C ratio) and mitotic figures were more abundant in type A. Genetic analysis revealed no evidence of chimerism for these individuals with neoplasia type B (low values of PUF and 2TD, Figure S2). Note that the highest PUF and 2TD values corresponded to two different samples (L6_49 for PUF and L6_46 for 2TD). However, our analysis of in-silico chimera with variable proportions of mixing between two individuals (Figure S4) suggested that a BTN emerged in an *M. edulis* founder that infects an *M. edulis* host can be missed if the neoplastic cells do not outnumber normal cells. It would therefore be possible that the proportion of neoplastic cells could have been too low in each individual. We used our in-silico chimera to test the proportion of neoplastic cells needed to detect chimerism with 2TD when we combine the eight samples with a Fisher’s combined probability test. More than 35% neoplastic cells in each sample would be needed to obtain a significant test. Given that almost all mussels infected were in moderate or later stages of the diseases (see Table S3), the most parsimonious interpretation is that type B disseminated neoplasia are not transmissible.

**Fig. 4:**
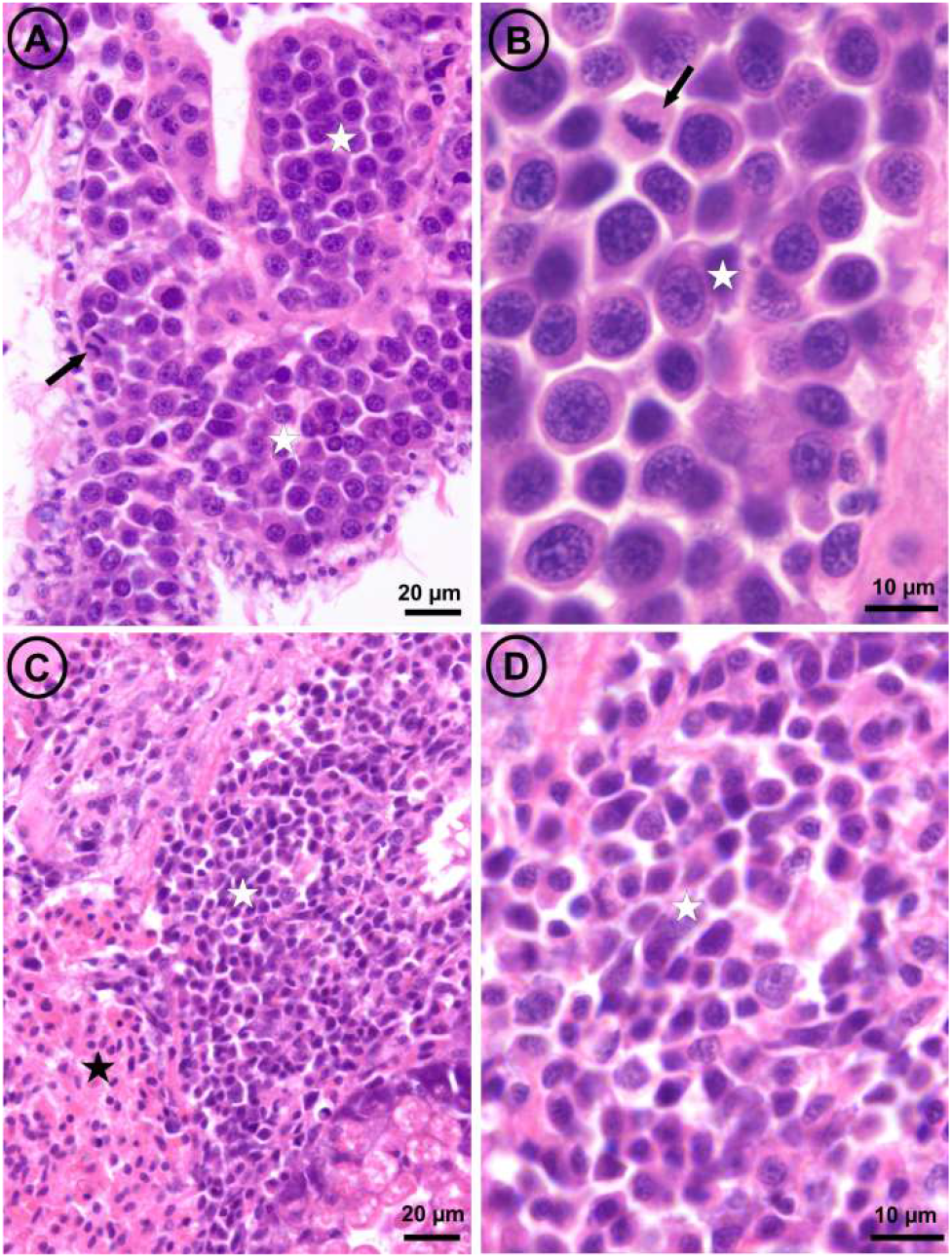
Light micrographs of histological sections through the visceral mass of two mussels, one affected by disseminated neoplasia type A (panels A and B) and the other by disseminated neoplasia type B (panels C and D). The arrows point out mitotic figures; white stars mark masses of neoplastic cells; the black star marks a mass of normal hemocytes.

### e. Genotypic differences and polymorphism of MtrBTN2 tumors

Cancerous samples were a mix of host and tumor genotypes and when the proportion of cancer cells was too low we could not infer the tumor genotype with accuracy. Hence, we inferred the tumor genotype of 6 cancerous individuals with the highest MtFF2 values (more than 0.75) and composed of a vast majority of cancer cells (up to 90%, FigureS1). Most of the SNPs were associated with *M. trossulus*-state alleles (Figure 5C, Figure S6, see “nuclear fixed”), as expected for an emergence in a *M. trossulus* host, but 5 SNPs had *M. edulis*-state alleles at the heterozygous or homozygous states, three fixed in the 6 tumors and two shared by at least two tumors (Figure 5C, red arrowhead). This shows an excess of *M. edulis* allelic states in MtrBTN2 compared to MtrBTN1 and *M. trossulus* populations used as reference, or a deficit compared to the strongly introgressed population of the Baltic Sea.

**Fig. 5:**
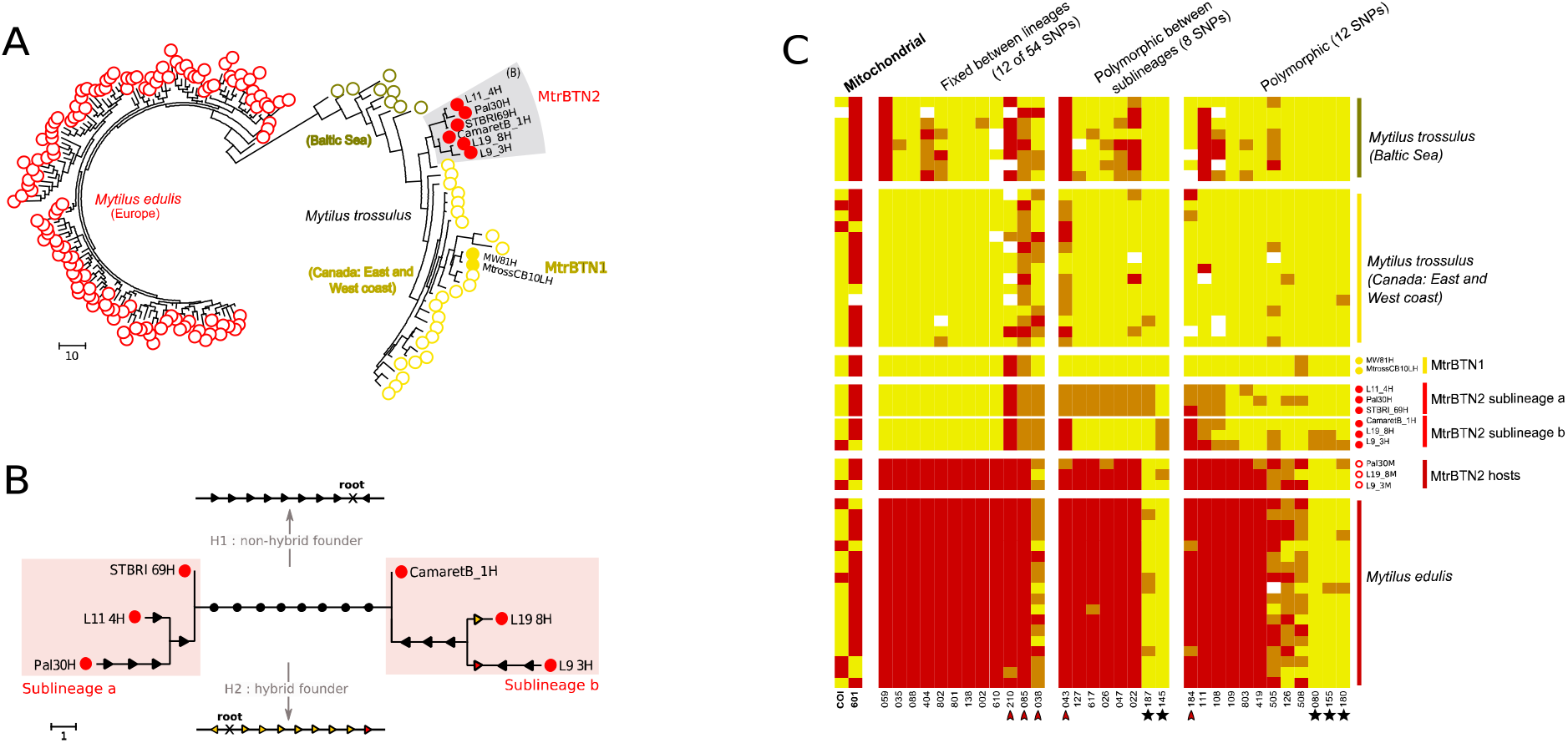
MtrBTN2 tumor genotypes. (A) Maximum parsimony tree estimated from 74 nuclear SNP genotypes of MtrBTN2 tumors (red filled circles), *M. edulis* individuals (red open circles), MtrBTN1 tumors (yellow filled circles), *M. trossulus* individuals from British Columbia (yellow open circles) and Batic Sea (dark yellow open circles). (B) Unrooted subtree of the 6 MtrBTN2 tumors. Mutations are reported and represent gain of heterozygosity (black arrow), loss of heterozygosity (colored arrow, yellow for mutation toward a *trossulus*-state allele and red toward an *edulis*-state allele). The founder (common ancestor) genotype being unknown, the 8 mutations separating the two lines (black circles) cannot be oriented without assumptions. We present two hypotheses: a non-hybrid founder (H1, all mutations are gain of heterozygosity) and a hybrid founder (H2, all mutations are loss of heterozygosity but mostly to *trossulus*-state allele). (C) Plot of genotypes for a subset of SNPs in columns and individuals in rows. Red: homozygote of *edulis*-state allele; yellow: homozygote of *trossulus*-state allele; orange: heterozygote. Individuals’ origins are reported by circle type and color (see (A)) and only 17 of the *M. edulis* individuals are represented here. The first two columns correspond to mitochondrial SNPs, the twelve next to a subset of nuclear SNPs fixed in the 6 MtrBTN2 tumors, then eight SNPs fixed between the two sublineages and then 12 SNPs revealing polymorphism within each sublineages. Red arrowhead pointed the five SNPs with *M. edulis*-state allele excess in the 6 tumors. Black star highlights the five SNPs for which *M.edulis*-state allele in tumors are not explained by the null allele hypothesis. All individuals and SNPs are detailed in Figure S5.

Using the 74 nuclear SNPs, we then reconstructed a MP-tree with *M. edulis* and *M. trossulus* healthy individuals, 2 MtrBTN1 and 6 MtrBTN2 (Figure 5A). As expected, *M. edulis* and *M. trossulus* form two genetically distinct clusters. MtrBTN2 samples group together and differ from MtrBTN1, confirming the existence of two distinct lineages (Yonemitsu et al. 2019). The two *Mytilus* BTN lineages are related to the *M. trossulus* cluster also confirming the *M. trossulus* origin of both.

Even though MtrBTN2 tumors group together, many genetic differences (polymorphisms) were observed among the 6 tumors. A zoom on the subtree with only the 6 tumors (Figure 5B) shows two groups of tumors: L11_4, PAL30 and STBRI69 (subclade a) on one side and L19_8, L9_3 and CamaretB_1 (subclade b) on the other side, separated by as many as 8 mutations (Figure 5C, see fixed between sublineages). Subclade a is homozygous for 7 of these 8 mutations (mostly *M. trossulus*-state alleles) while subclade b appears heterozygous. Mutations in each subclade can be oriented by parsimony. Surprisingly, 10 of those 12 mutations are gain of heterozygosity (GOH, Figure 5B, black arrow) while only 2 are loss of heterozygosity (LOH, Figure 5B, colored arrow). However, the 8 mutations separating the two subclades cannot be oriented without rooting the tree, and we cannot root the tree given the host is sexually reproducing. We can however work on alternative hypotheses given the SNPs are diagnostics between *M. trossulus* and *M. edulis*. Under a “non-hybrid founder” hypothesis (Figure 5B, see H1) the ancestor is assumed homozygous for *M. trossulus*-state alleles and subclade b is inferred to have an excess gain of heterozygosity (GOH). Under a “hybrid founder” hypothesis (Figure 5B, see H2) the ancestor is assumed heterozygous and we conversely infer an excess loss of heterozygosity (LOH) favoring *M. trossulus*-state alleles in subclade a. Finally, 5 of the 6 tumors share the same allele at the two mitochondrial SNPs except L9_3 which is different at these two SNPs (Figure 5C).

The mtCOI tree (Figure 6) mostly confirms the nuclear tree and sublineages with the exception of the L9_3 mtCOI sequence, which appears divergent to every other MtrBTN2 sequence sampled in *M. edulis* or *M. chilensis* while this tumor is related to L19_8 and CamaretB_1 in the nuclear tree. Together with the MtrBTN2 sequences obtained from *M. trossulus* (Skazina *et al*. 2020), the L9_3 mtCOI sequence suggests 62mc10 mitochondria came from a MtrBTN2 tumor, as extensively discussed by Skazina *et al*. (2020). Briefly, this sequence has been obtained after an extensive screening of a *M. trossulus* mitochondria in Baltic mussels known to be fixed for introgressed *M. edulis* mitochondria, and 62mc10 was found heteroplasmic for two F mitochondria (Śmietanka and Burzyński 2017). The closest two mtCOI sequences of healthy mussels, AY823625 and RET-14, have been obtained from Chester Basin (Nova Scotia, Canada) and in Retinskoye (Murmansk Oblast, Rusia) respectively. A basic molecular clock calculation using the 100Ky old Tx101A sequence for the calibration could have allowed to date the MRCA of every MtrBTN2. However, we note that AY823625 have no substitutions from the ancestral node and RET-14 only one (Figure 6B), and that the MtrBTN2 clade appears to have much longer branches, suggesting an accelerated evolutionary rate that prevent accurate datation of the MRCA.

**Fig. 6:**
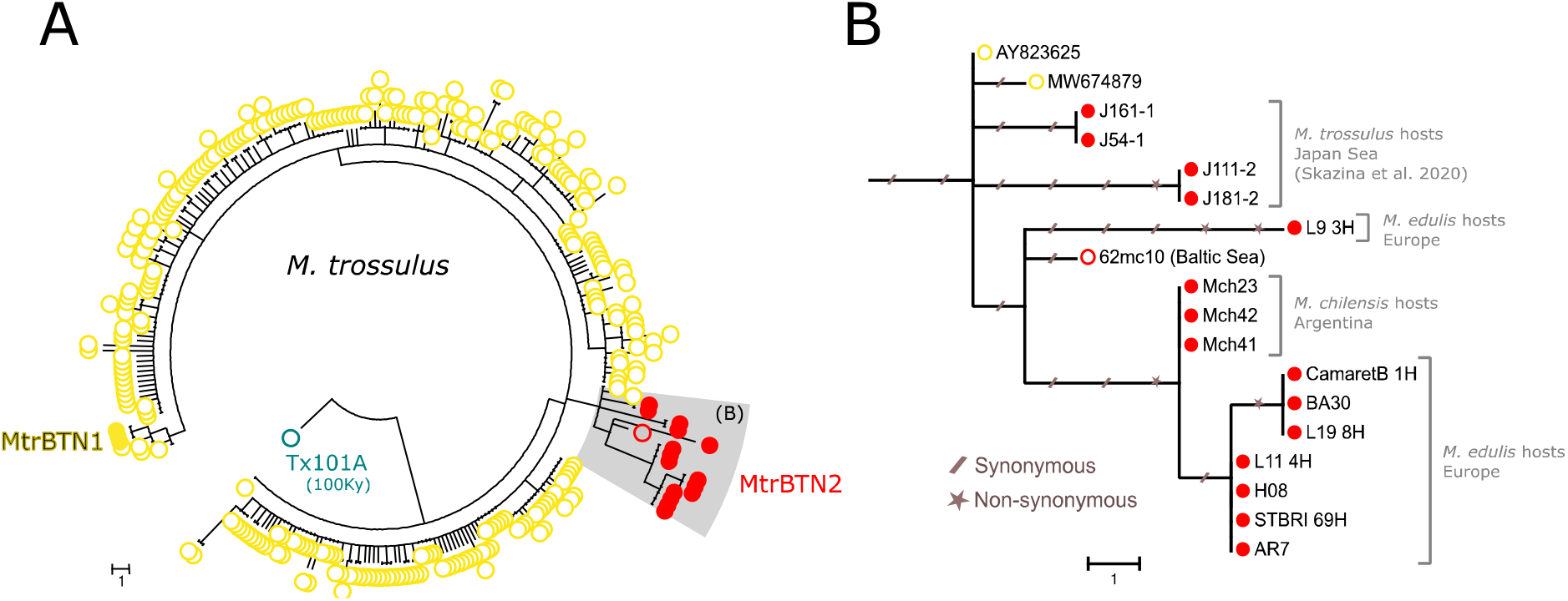
Phylogenetic analysis of mtCOI sequences. (A) Maximum parsominy tree on mtCOI 603bp aligned sequences of our 5 MtrBTN2 tumor samples (red filled circles, L9_3, CamaretB_1, L19_8, L11_4, STBRI69) and all available *M. trossulus* (GenBank) including three MtrBTN1 tumor samples (yellow filled circles), seven MtrBTN2 tumor samples of which four are from *M. trossulus* hosts (Skazina et al. 2020) and three are from *M. chilensis* hosts (red filled circles), a 100 000 years old (Tx101A, blue open circles) and contemporary *M. trossulus* mussels. (B) Subtree of MtrBTN2 mtCOI haplotypes with the two closest relatives from healthy mussels. Synonymous and non-synonymous mutations are reported with slashed traits and stars respectively. Host species and geographical origins are indicated on the right side of the tree.

## 4. Discussion

We investigated the presence of transmissible cancer in samples of European *M. edulis* and *M. galloprovincialis* mussels and their hybrids. We used SNP KASP genotyping to screen a large number of mussels in order to detect transmissible cancers and at the same time studied the genotypes of the infected hosts and the genotypes of tumors. We detected MtrBTN2, a previously described transmissible cancer that originated in a *M. trossulus* host, but no other transmissible cancers have been found with our screening approach in the inspected area. The MtrBTN2 was known to affect *M. edulis*, (Riquet et al. 2017, Burioli et al. 2019), *M. chilensis* (Yonemitsu et al. 2019) and *M. trossulus* (Skazina et al. 2020) but we also found cases in *M. galloprovincialis* and hybrid individuals, thus extending the known spreading capacity of MtrBTN2 to an additional species of the complex. Despite an overall low prevalence (0.42%), it appeared more prevalent in *M. edulis* hosts (0.78%) than *M. galloprovincialis* (0.07%) and hybrids (0.26%) hosts. We also found neoplastic mussels without genetic chimerism suggesting the presence of conventional non-transmissible cancers. The analysis of the genotype of 6 tumors from *M. edulis* hosts confirmed the *M. trossulus* origin of MtrBTN2 but with an excess of *M. edulis*-state SNP alleles compared to MtrBTN1 and the *M. trossulus* population we analysed, or a deficit compared to the highly introgressed *M. trossulus* population of the Baltic Sea. We surprisingly observed a high amount of nuclear and mitochondrial polymorphism in MtrBTN2 samples which defines two sub-lineages that co-exist and spread in the same populations, as it has been observed in other transmissible cancers (CTVT : Baez-Ortega et al. 2019, DFTD: Kwon et al. 2020, Patton et al. 2020).

The MtrBTN2 prevalences found in our study were consistent with DN prevalences previously reported in *M. edulis* (around 1%, for review see Carballal et al. 2015, Muttray et al. 2018) and *M. galloprovincialis* mussels (rarer and when detected <1%, for review see Carballal et al. 2015, Matozzo et al. 2018), but slightly lower than the prevalence of *M. trossulus* BTN found in Brittany (3.4%, Burioli et al. 2019). It is likely that we missed very early stages of the disease. Indeed, MtFF estimates the excess of *M. trossulus* allele fluorescence and when the proportion of cancer cells is too low, the host genotype overshines the signal. The two other indexes we used (PUF and 2TD) did not help to find additional tumors, with the exception of one sample (CamaretA_61) that was infected by a few neoplastic cells and showed a MtFF value at the limit of the outlier threshold (Figure 4A, green triangle-up). Although the prevalence of MtrBTN2 could be slightly underestimated, the detection limit is similar for *M. edulis*, *M. galloprovincialis* and their hybrids.

The higher prevalence in *M. edulis* than *M. galloprovincialis* and their hybrids in between could reflect a lower sensitivity (higher efficient immune response) of *M. galloprovincialis* to MtrBTN2. Indeed, Fuentes et al. (2002) found a higher DN prevalence in hybrids from lab crosses (*edulis* x *galloprovincialis*) than *M. galloprovincialis* ones (*galloprovincialis* x *galloprovincialis*), but unfortunately no *M. edulis* lab crosses (*edulis* x *edulis*) were reported. Moreover, it has been shown that *M. galloprovincialis* is less affected by Pea crab and Bucephalid trematode parasitism than *M. edulis* living in the same region (Seed 1969, Coustau et al. 1991a). Also, a study of mussel hemocytes transcriptomes shows higher sequence numbers related to immune genes in *M. galloprovincialis* compared to *M. edulis* after in vitro and in vivo stimulation with pathogen associated molecular patterns and heat-inactivated *Vibrio anguillarum*, respectively (Moreira et al. 2018). Beyond that, *M. galloprovincialis* is a worldwide invader (Popovic et al. 2020) with a fitness advantage in natural hybrid zones (Skibinski 1983; Coustau et al. 1991b; Hilbish et al. 1994; reviewed in Gardner 1994) and in lab crosses (Bierne 2006) which, in different ways, appears to be more efficient than *M. edulis*.

The presence of eight non-transmissible neoplasia is not surprising because it is the prerequisite for transmissible cancer to evolve in a founder host. Cases of conventional non-transmissible cancer have also been described in Chile where two tumors without transmissible cancer hallmarks were found in the same area with two MtrBTN2 tumors (Yonemitsu et *al*. 2019). The eight tumors (type B) were collected at the same date (April 2017), in the bay of Brest, where level of lead (Pb) was found above the regulatory threshold for human food (Charles et al. 2020). Environmental contaminants have previously been proposed as an etiological cause of DN (Lowe and Moore 1978, Hillman 1993), such as biotoxins (Landsberg 1996) and viral factors (Elston et *al*. 1988, Moore et al. 1991, Rasmussen 1986), which could be involved in the high prevalence of type B diseased individuals in this specific area and date. However, we also sampled mussels from bay of Brest in February where Pb level was also high (Charles et al. 2020), but no type B cancerous mussels were found. The coexistence of a transmissible cancer with non-transmissible cancers, warn for systematic use of molecular markers to verify the transmissible nature of DN. More generally, wildlife cancer research needs to consider a more general use of molecular markers to address the issue (Leathlobhair et al. 2017).

Polymorphism in transmissible cancer at SNPs ascertained in host populations can suggest multiple emergences, or a single emergence with somatic back mutations, structural mutations or with a deviation from clonal propagation (horizontal gene transfer). Comparison of the 6 tumors with MtrBTN2 mtCOI sequences (Yonemitsu et *al*. 2019, Skazina et al. 2020), and the grouping of the six tumors with 54 fixed SNPs seems to suggest a single emergence is the most parsimonious interpretation. However, we cannot rigorously reject the hypothesis of a twin co-emergence in an ancestral (or contemporary unsampled) *M. trossulus* population with a different genetic composition than contemporaneous *M. trossulus* populations. The recent independent emergence of two transmissible facial-tumour lineages (DFTD1 and DFTD2) in Tasmanian devils suggests that when the narrow windows of favorable condition are met, multiple emergences can succeed in a short time period (Ujvari et al. 2016). Interestingly, we found *M. edulis*-state alleles on some SNPs (Figure 5C, red arrowhead) which could suggest an ancient origin, in a *M. trossulus* population which was less divergent than the actual population, or an origin in a recombinant hybrid between *M. trossulus* and *M. edulis* (Figure 5B, H1 and H2). However, it is also likely that somatic null alleles caused by structural variations or point mutations in primer-binding sites lead to enhanced detection of the host alleles at these SNPs.

Given that we analysed only 76 nucleotide positions and that the mtCOI divergence is not very strong, we believe polymorphism observed among tumors could more likely be due to structural rather than point variations, such as insertion-deletion and chromosomal losses or gains, or recombination leading to loss of heterozygosity, that are widespread in cancer (McClelland 2017, Nichols et al. 2020) and have been reported in other transmissible cancers (e.g. in DFT1 and DFT2, Kwon et al. 2020). According to the genotyping method used (competition of the two allele fluorescences), partial or complete chromosomal losses or gains results in a modification of the ratio of fluorescences of the two alleles of hyperploid MtrBTN2 tumors and of the ratio of tumor and host genotypes. We observed that most somatic variations that could be oriented resulted in gain of heterozygosity rather than loss (10 gains mostly of *M. edulis*-allele state, 2 losses), supporting the hypothesis that the variation is due to somatic null alleles leading to enhanced detection of the host *M. edulis* alleles. The same hypothesis would also explain the divergence between the two sublineages (H1 in Figure 5B). Another, although somewhat less likely, hypothesis could be horizontal gene transfer from host to tumors. This hypothesis would be supported by 5 SNPs for which the host allele state does not explain the heterozygous state of tumors (Figure 5C, black star). The same hypothesis would also explain the divergence between the two sub-lineages under the hypothesis of a hybrid founder genotype (H2 in Figure 5B). Indeed, spontaneous fusion between tumor and host cells has been shown to bring tumor heterogeneity which increases invasion and migratory capacities (reviewed in Noubissi and Ogle 2016). Recombination can also occur after cancer cell fusion (Miroshnychenko et al. 2021). Fusion between two BTN lineages or between BTN and conventional cancers could explain hyperpoliploïdy and variable VAFFs observed. However, in the absence of further evidence, somatic null alleles and mutations remains the most parsimonious explanation to the diversity observed.

Finally, one of the 6 tumors, L9_3, had a different genotype at the two mitochondrial SNPs analysed (Figure 5C, see mitochondrial) and the mtCOI tree showed this sequence is divergent to the other mtCOI sequences of MtrBTN2 tumors sampled in *M. edulis* and *M. chilensis* hosts (Figure 6). The mismatch between the nuclear and the mitochondrial trees suggests the existence of mitochondrial capture or recombination events in MtrBTN2. This phenomenon has been described several times in CTVT where mitogenomes reveal multiple independent horizontal transfers of mitochondria from hosts (Strakova et al. 2016, 2020). In addition, recombinant mitochondria have been observed in two MtrBTN2 tumors from Chile (Yonemitsu et al. 2019). Here however, the capture/recombination would have happened from another MtrBTN2 tumor, possibly by cellular fusion as explained above.

## 5. Conclusion

Our study revealed that MtrBTN2, a bivalve transmissible neoplasia that originated in an *M. trossulus* host, affects European *M. edulis* at a low prevalence and also *M. galloprovincialis* and hybrids although at a much lower rate. No other transmissible cancer has been identified with our approach, but conventional non-transmissible cancers have been observed, which suggests molecular markers should be used in order to diagnose the transmissible status of disseminated neoplasia. Somewhat surprisingly we found polymorphism among MtrBTN2 tumors, likely due to somatic structural variations leading to null alleles. The origin of this polymorphism will need to be further investigated with alternative sequencing methods. The coexistence of sublineages in the same geographic region suggests a long history of MtrBTN2 diversification with a complex biogeography as also reported in CTVT and DFTD (Baez-Ortega et al. 2019, Know et al. 2020, Patton et al.2020).

## Acknowledgements

This work was supported by Montpellier Université d’Excellence (BLUECANCER project), Agence Nationale de la Recherche (TRANSCAN project, ANR-18-CE35-0009), the French DPMA (Direction des pêches maritimes et de l’aquaculture, DPMA-2017-MORBLEU Conventions N°: 16/1212569 & 17/1212952) and Conseil Régional de Normandie, LabEx CeMED (Junior Research Team funding).

## Author Contributions

M.H., A.S., M.J.M and N.B. designed the project. All authors contributed to sampling, sample preparation and data curation. M.H., A.S., C.A. conducted the molecular work with guidance from G.C., D.D-G and N.B., and A.V., E.A.V.B. and A.B. performed histology, cytology and flow cytometry. M.H., A.S., J.J.W. and N.B. analyzed the data. M.H. and N.B. wrote the initial manuscript. All authors contributed insights about interpretation of results, and critically reviewed and edited the manuscript.

## Supplementary Files

**Table S1:**
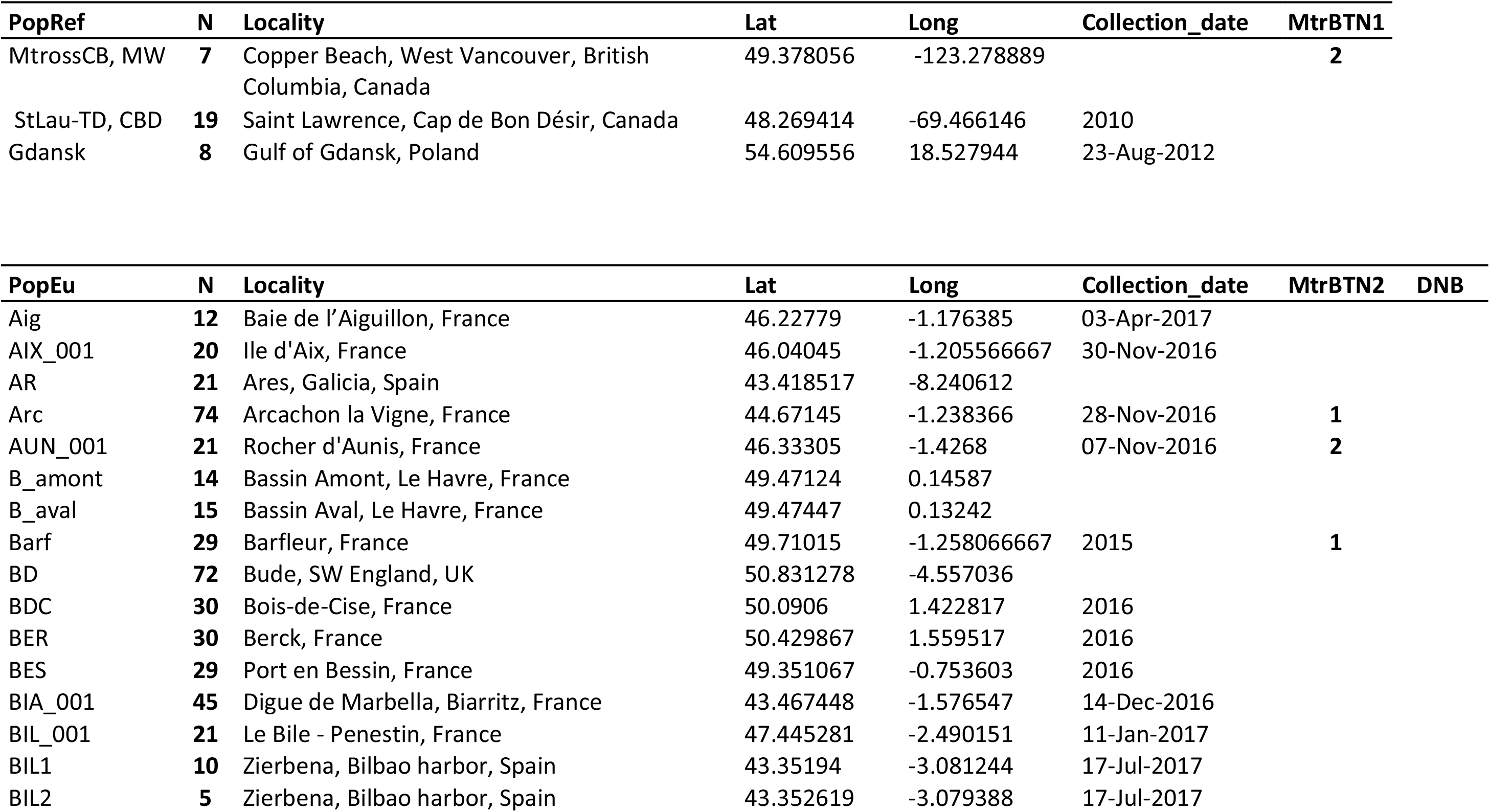
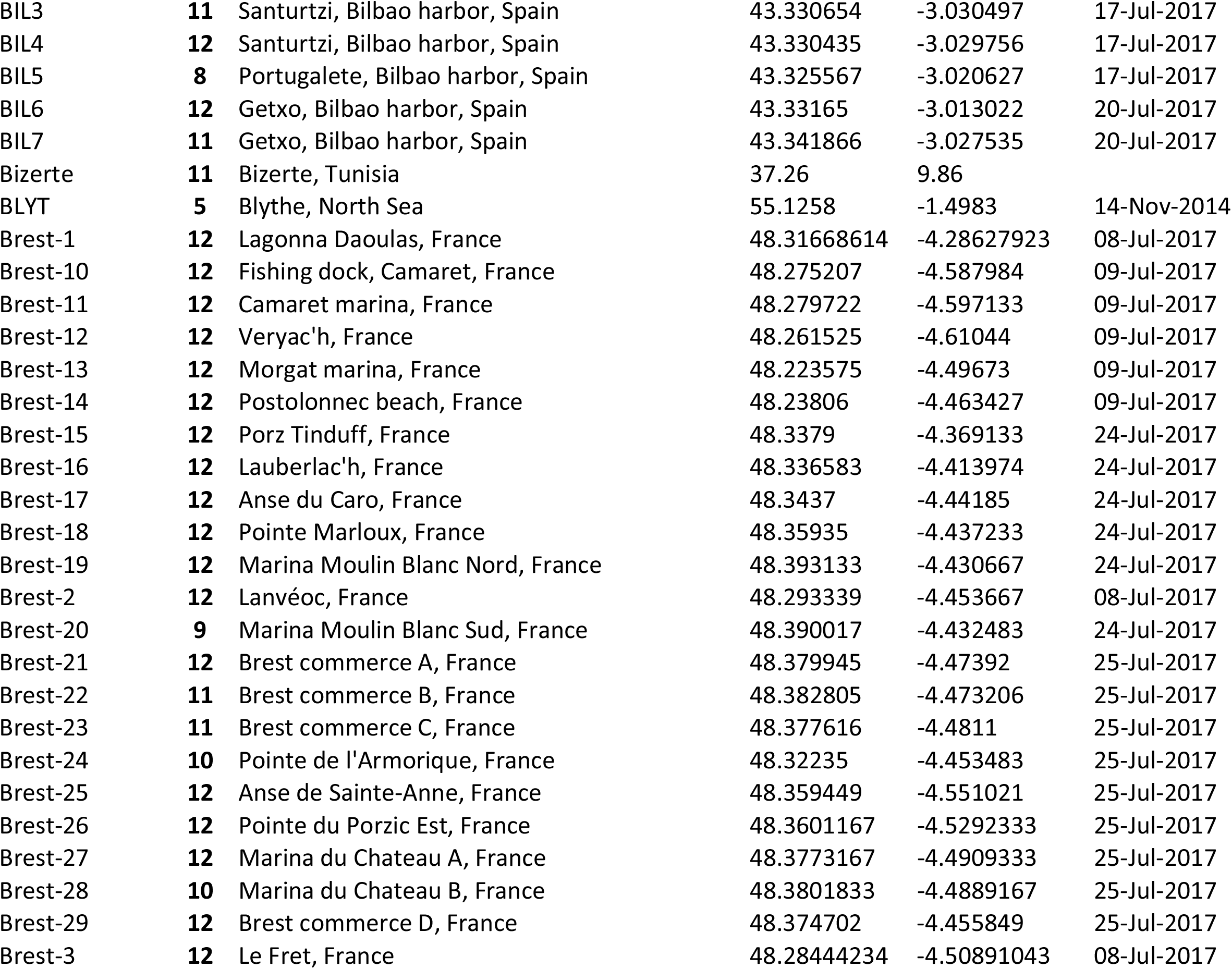
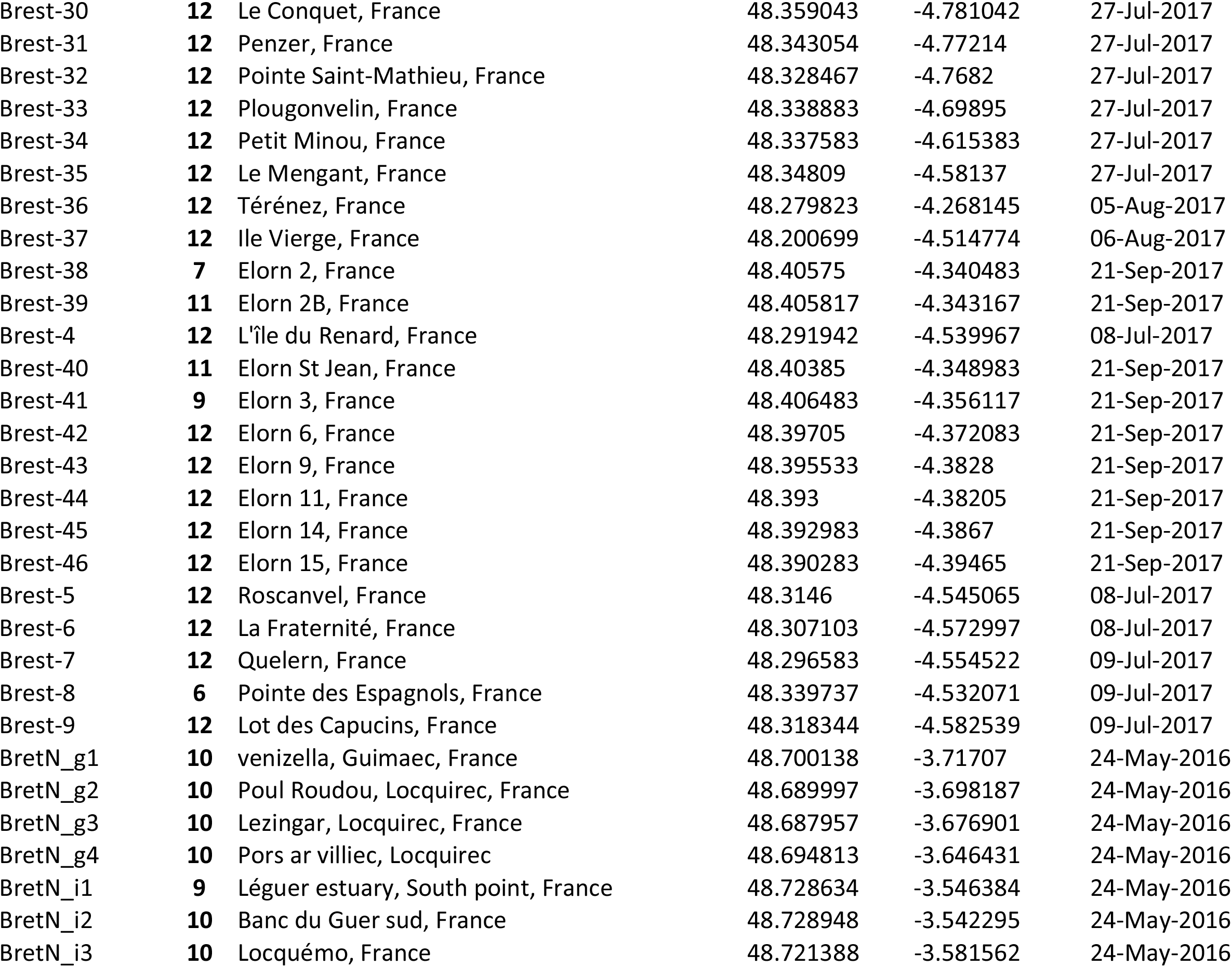
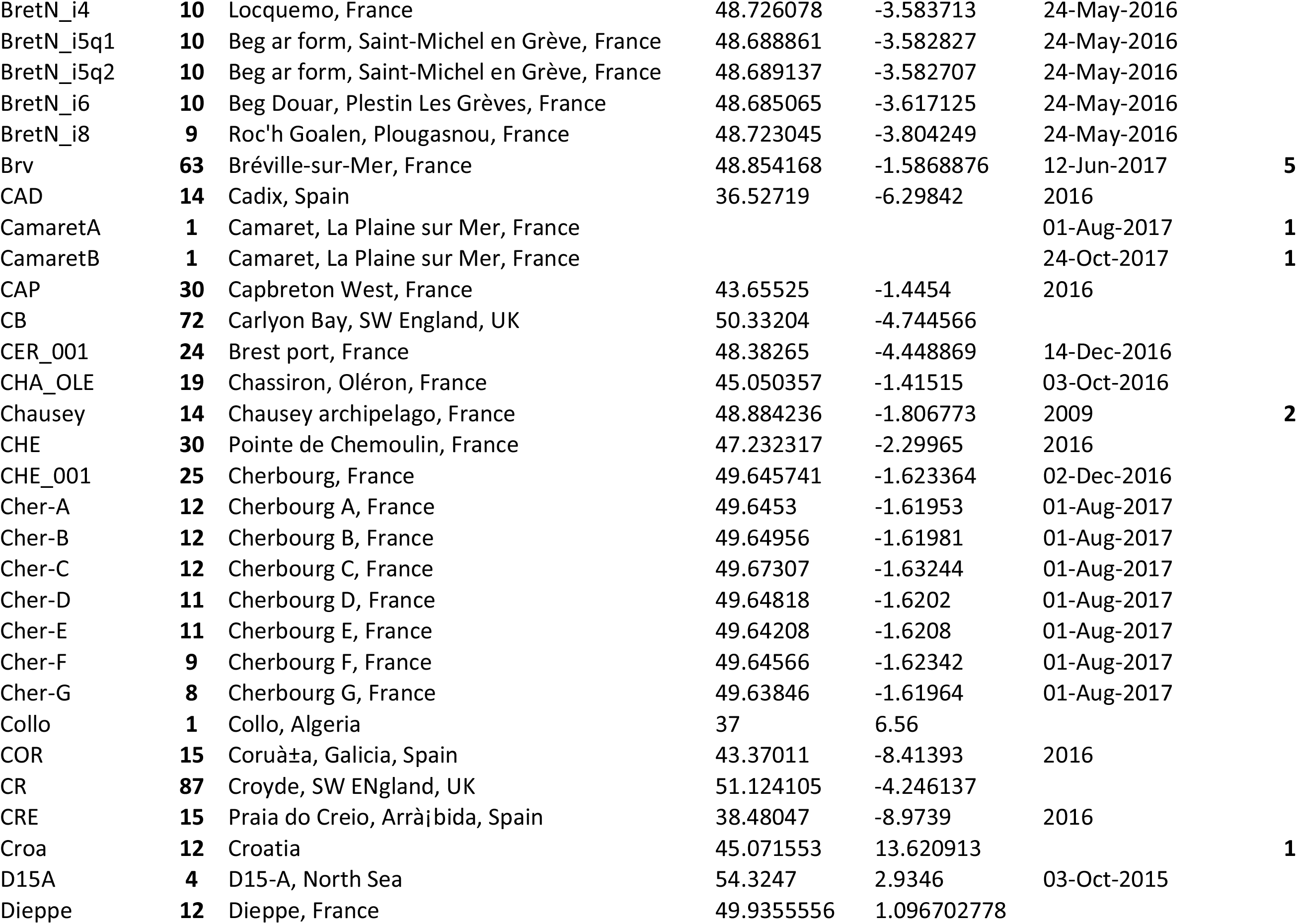
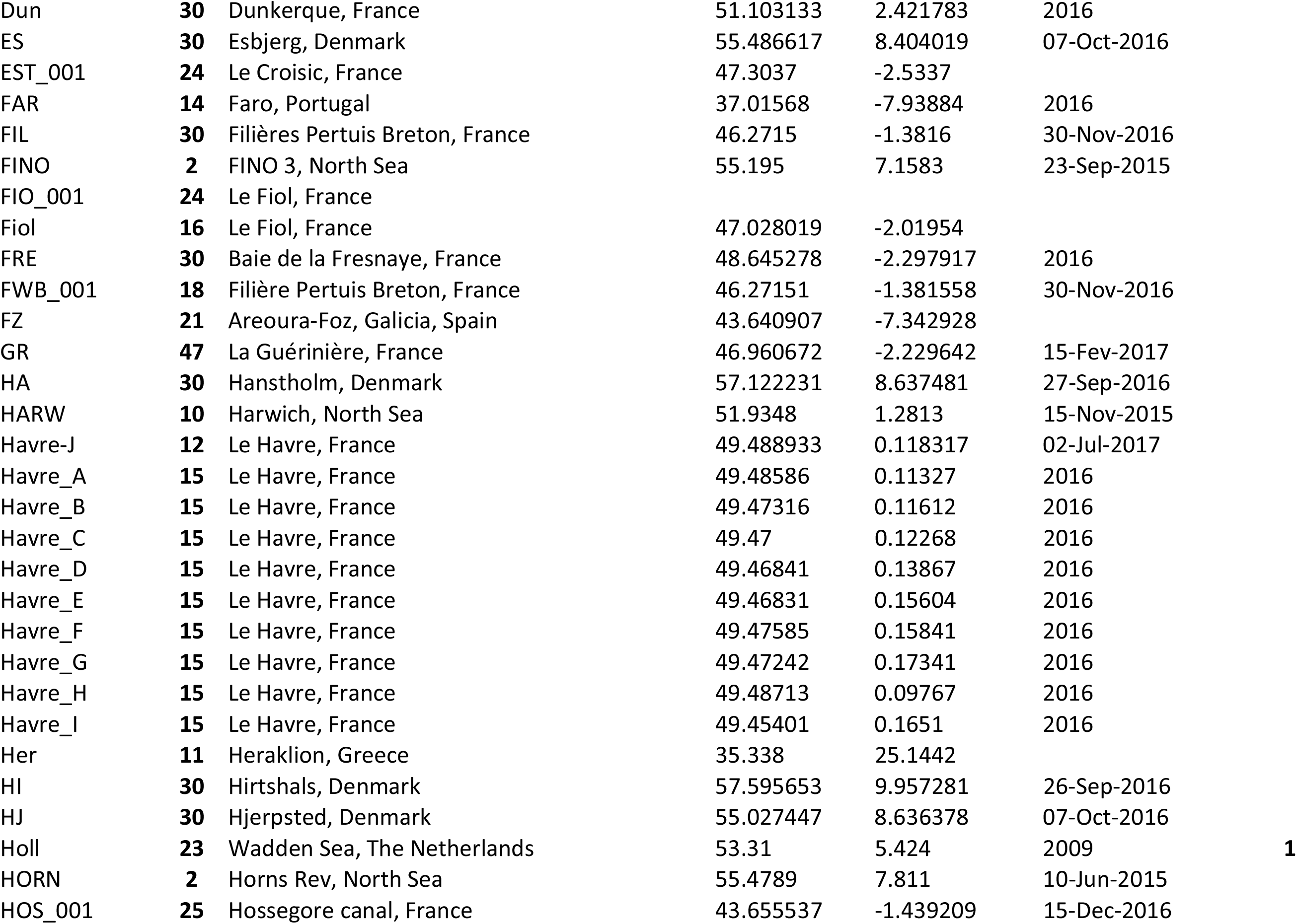
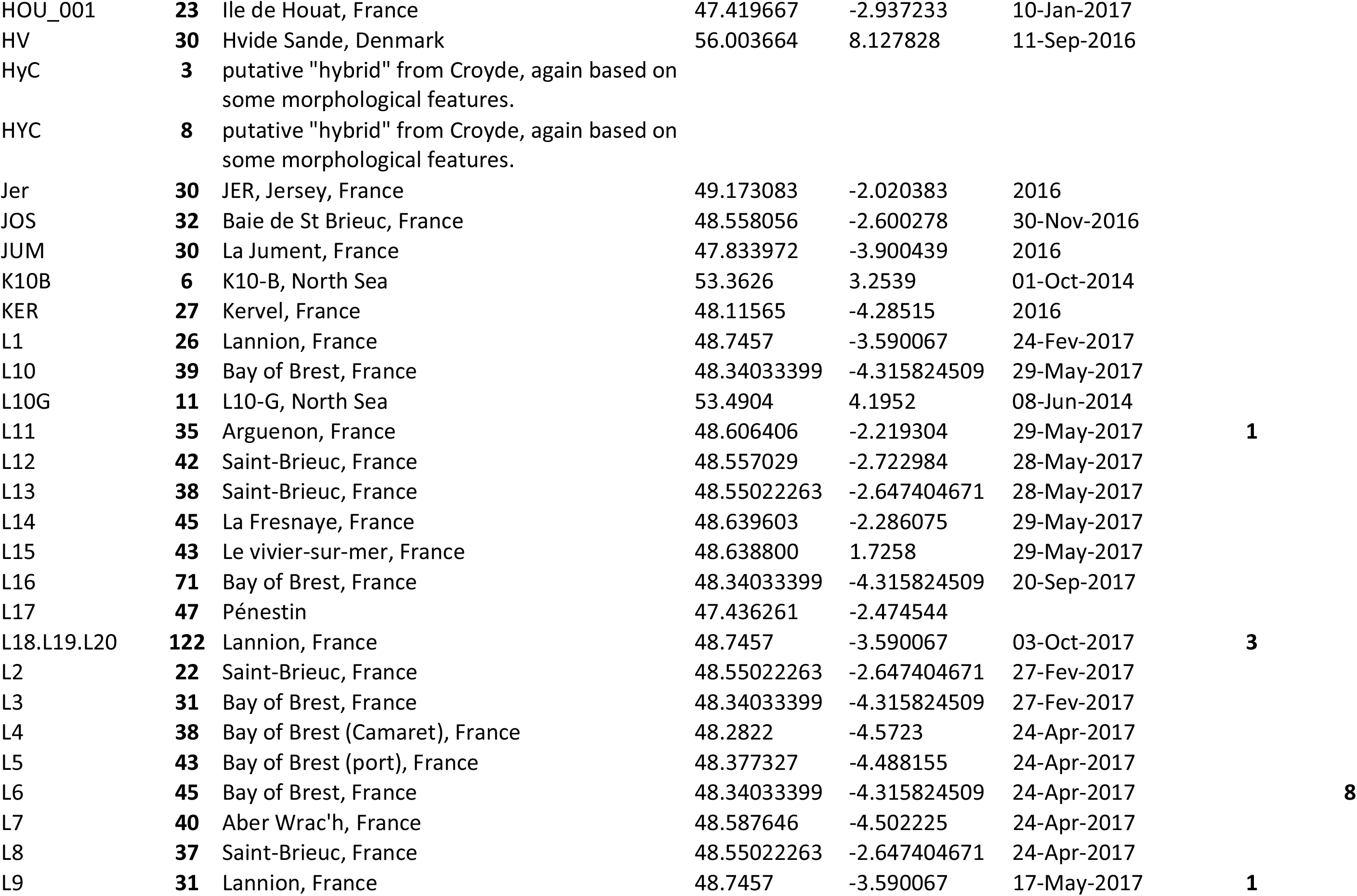
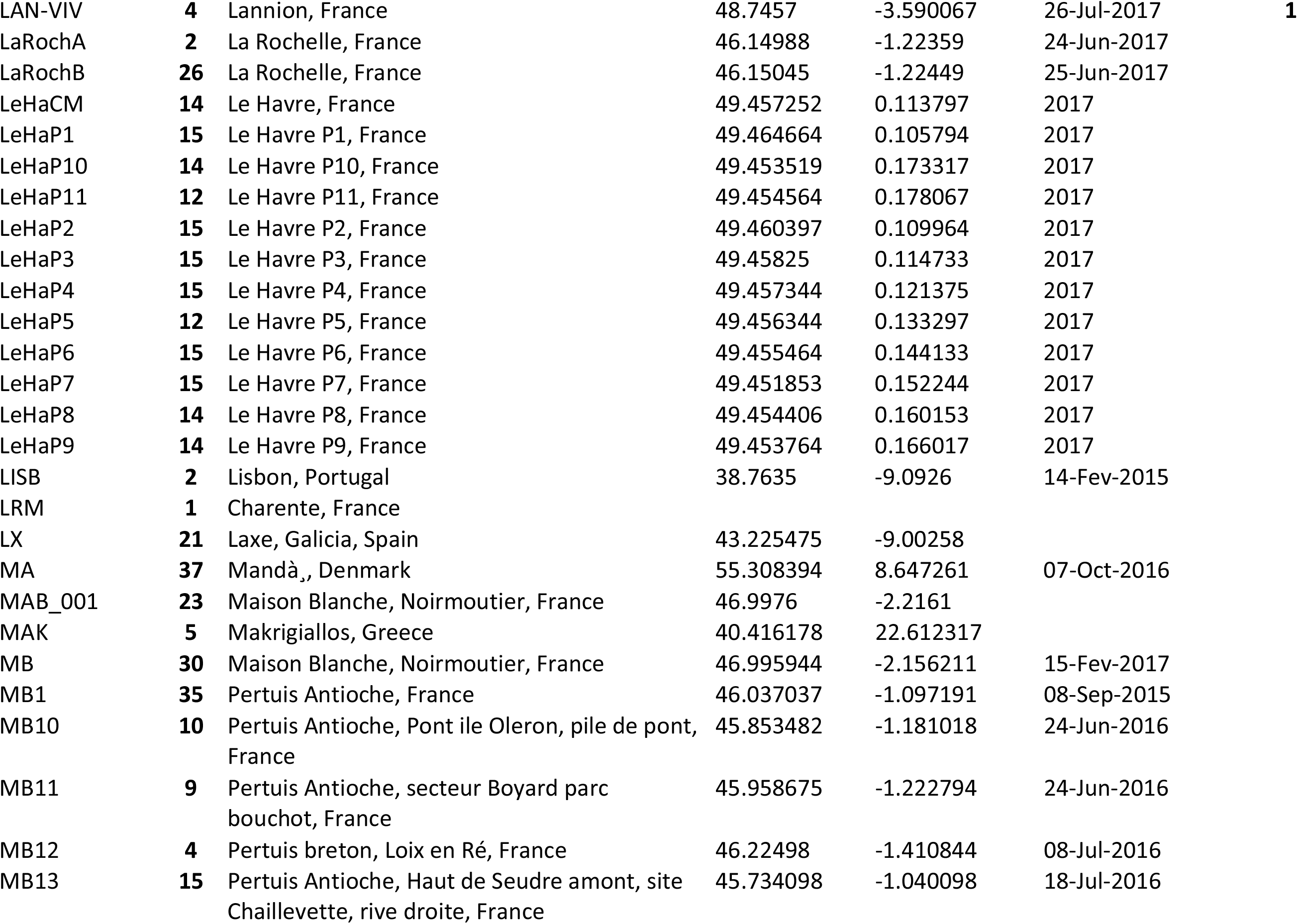
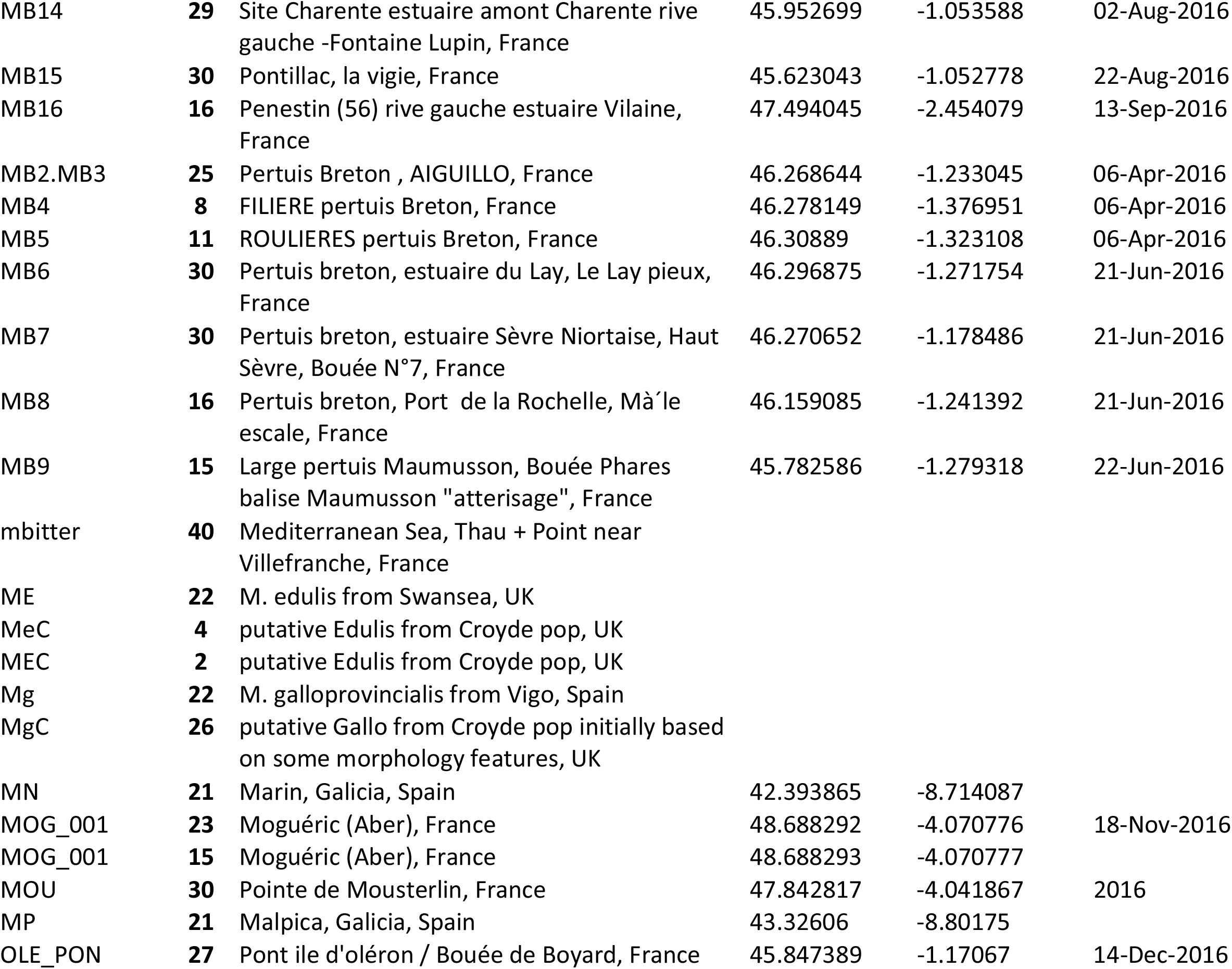
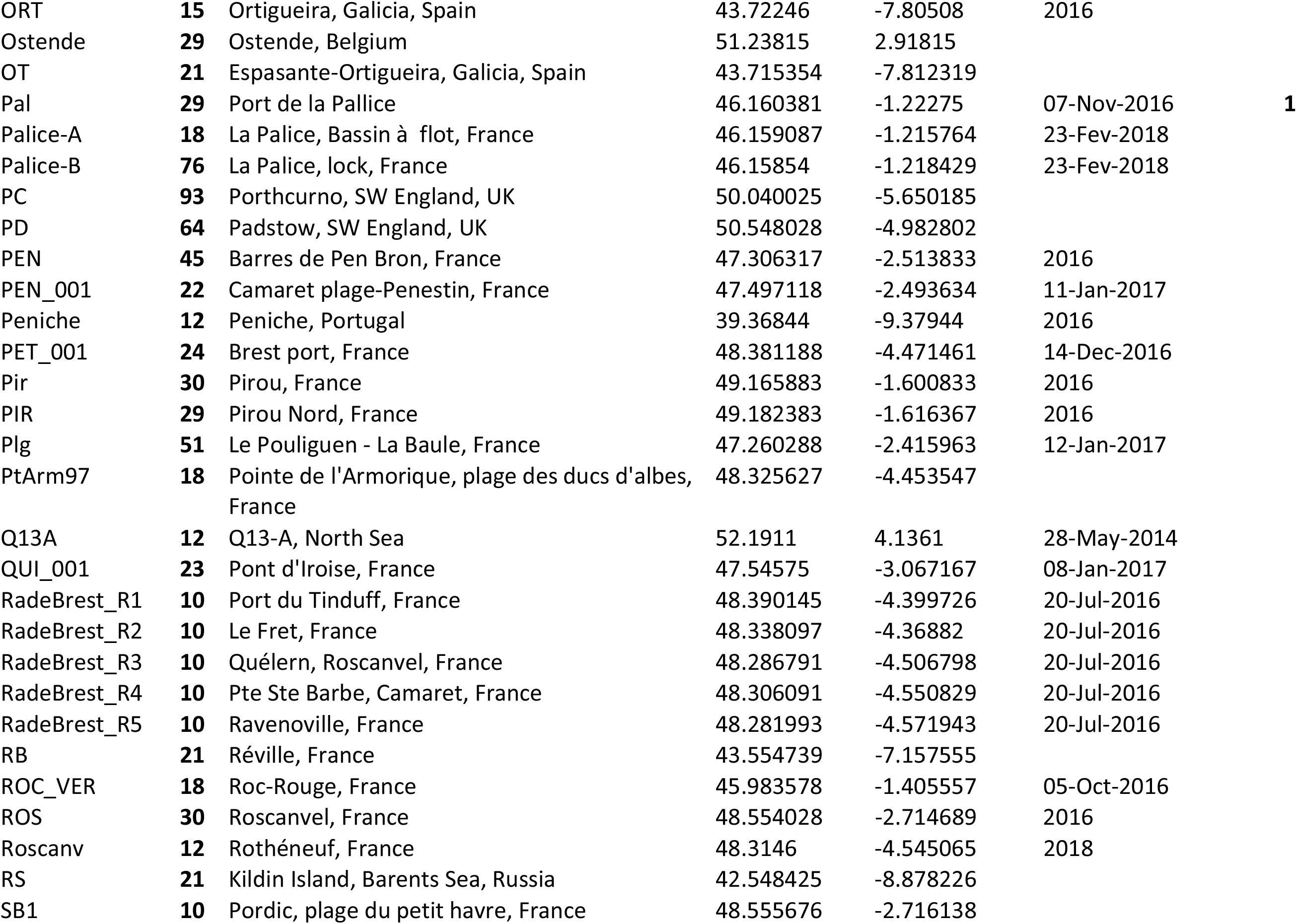
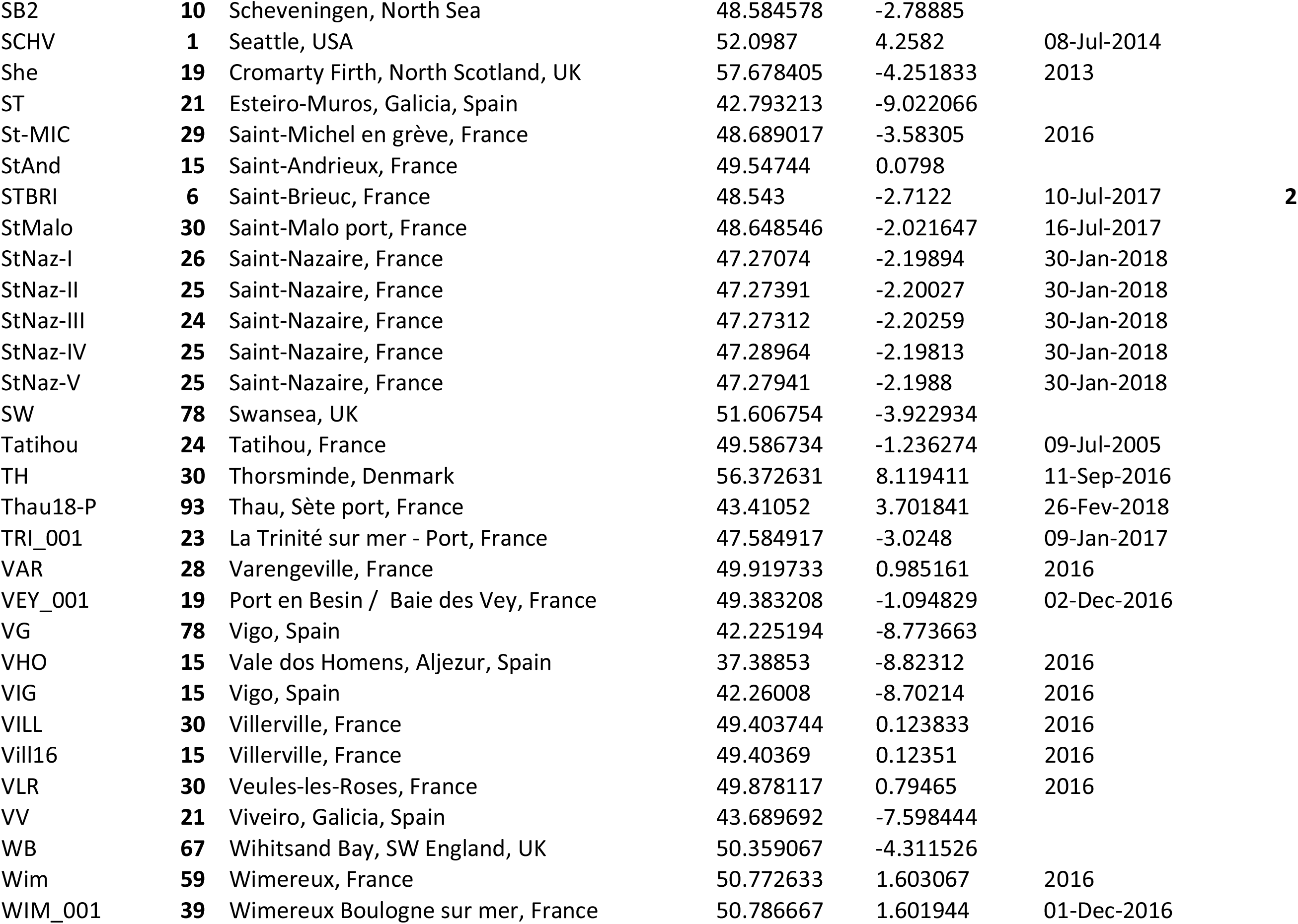
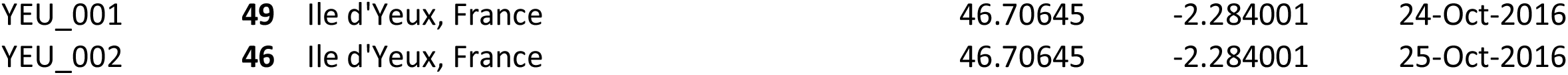
Description of the sampled locations of *M. trossulus* references (PopRef) and *M. edulis* and *M. galloprovincialis* individuals (PopEu). Pop: population name; N: number of individuals sampled; Lat: latitude; Long: longitude; MtrBTN1: number of individuals with MtrBTN1 tumor; MtrBTN2: number of individuals with MtrBTN2, the tumor detected in this study; DNB: number of individuals with disseminated neoplasia type B (non-transmissible cancer).

**Table S2:**
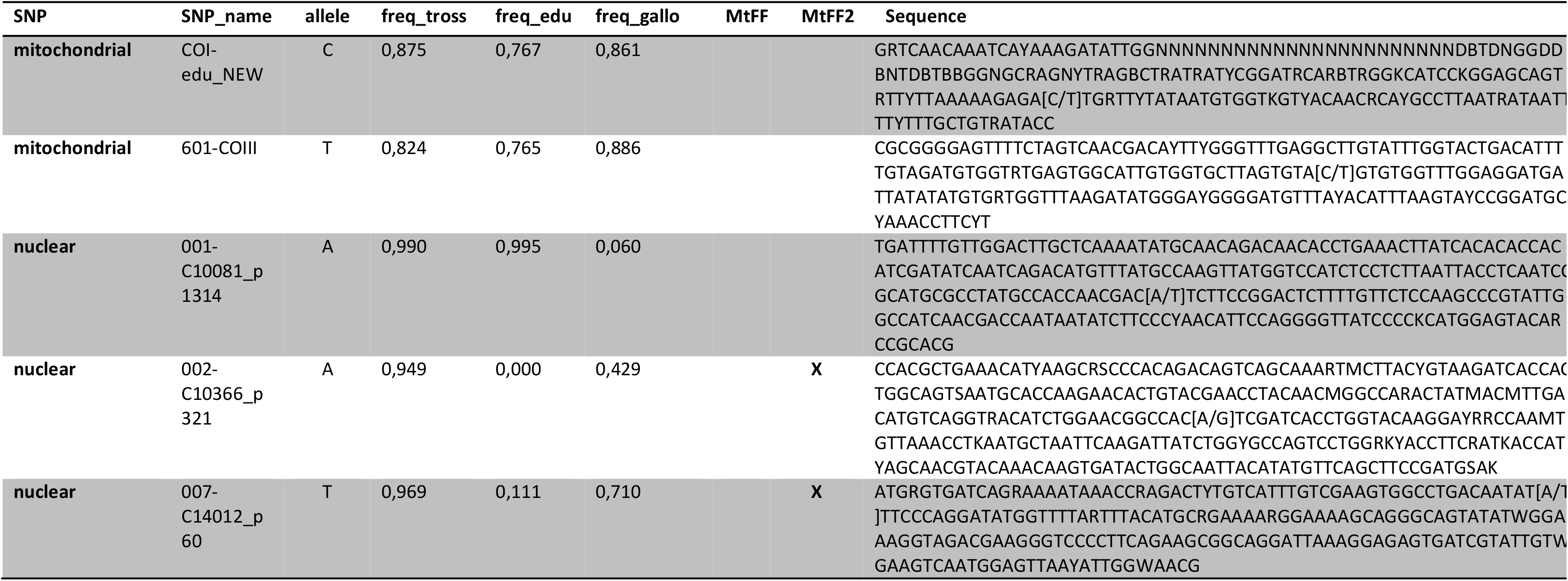
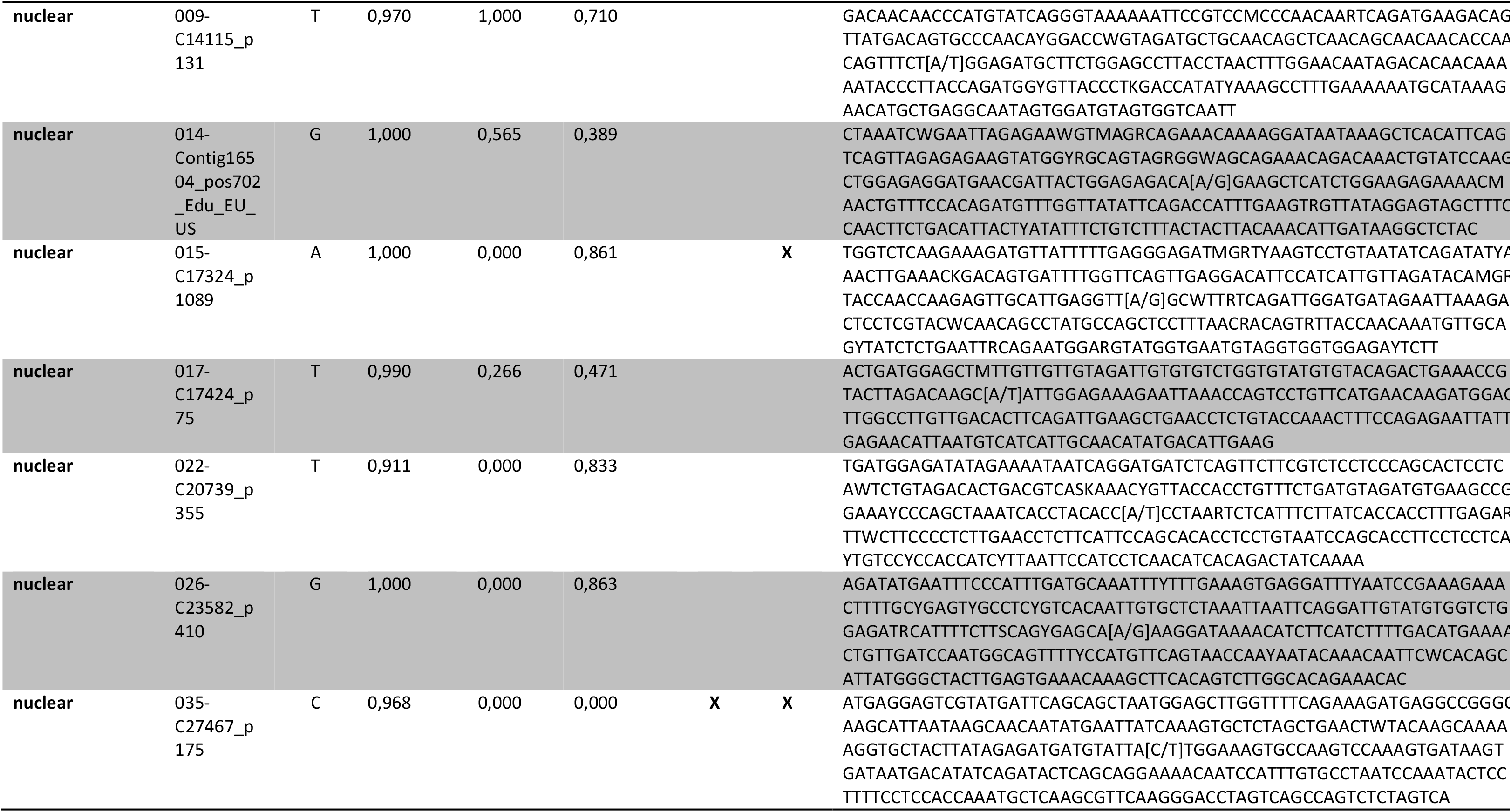
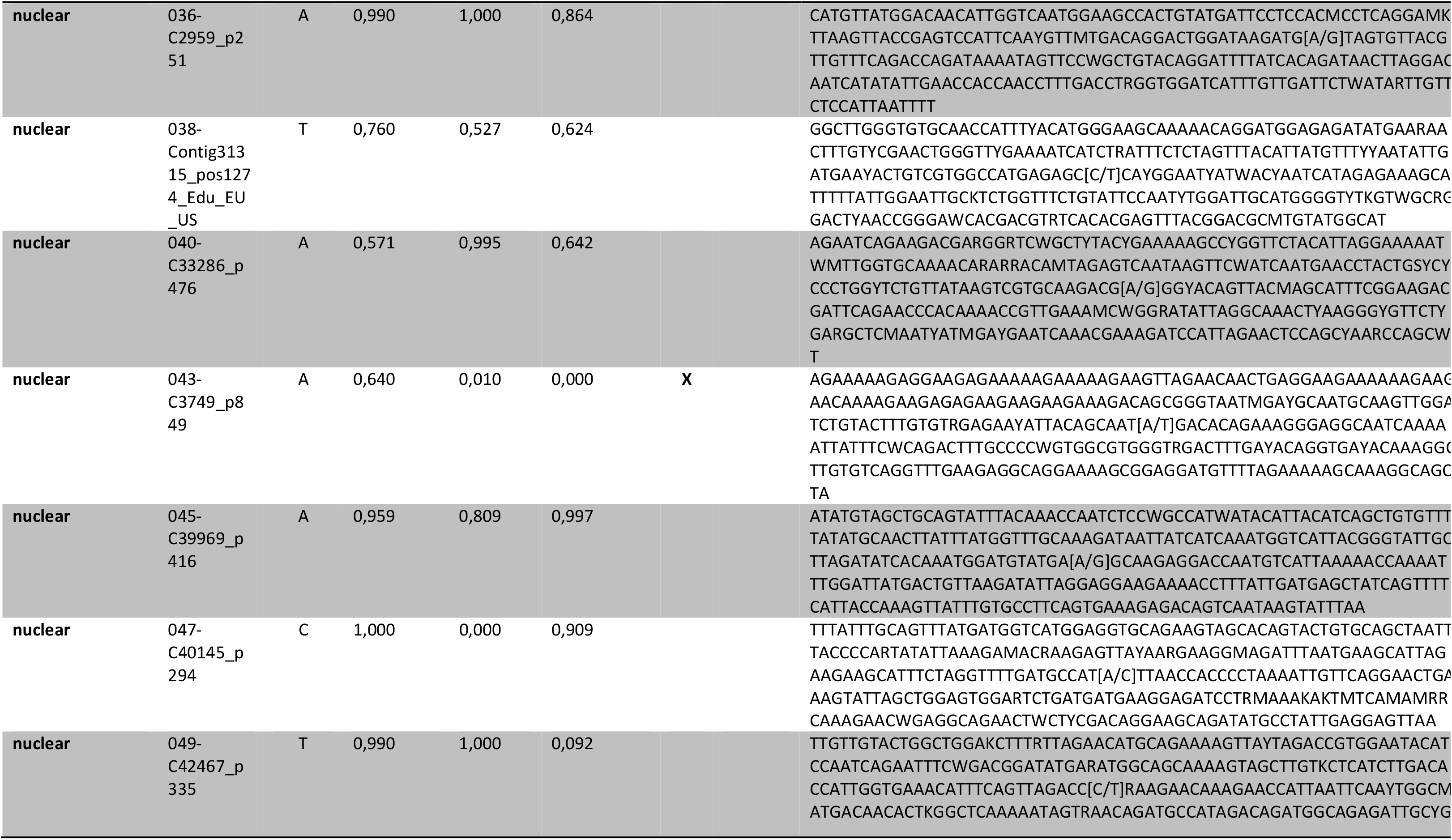
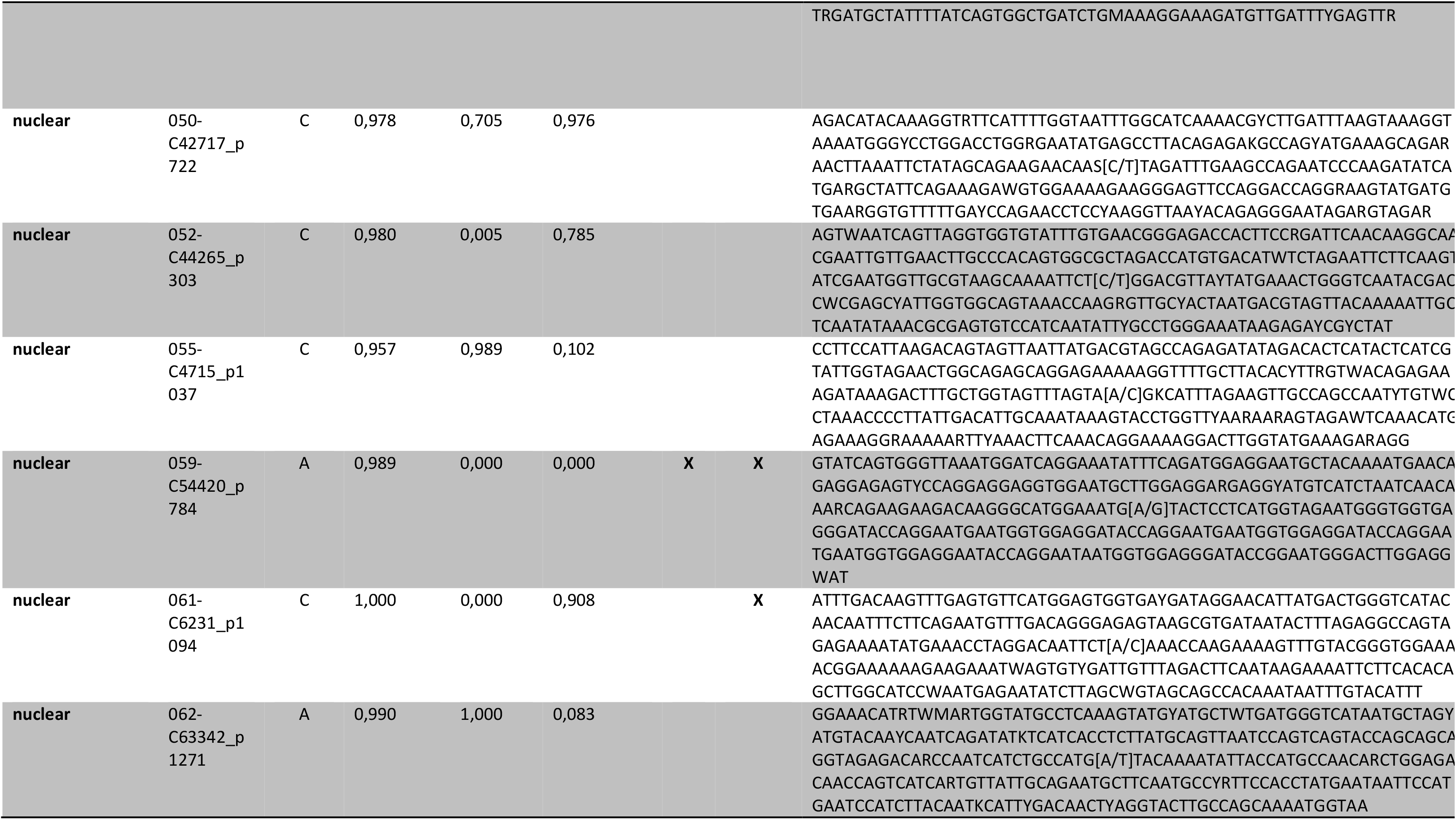
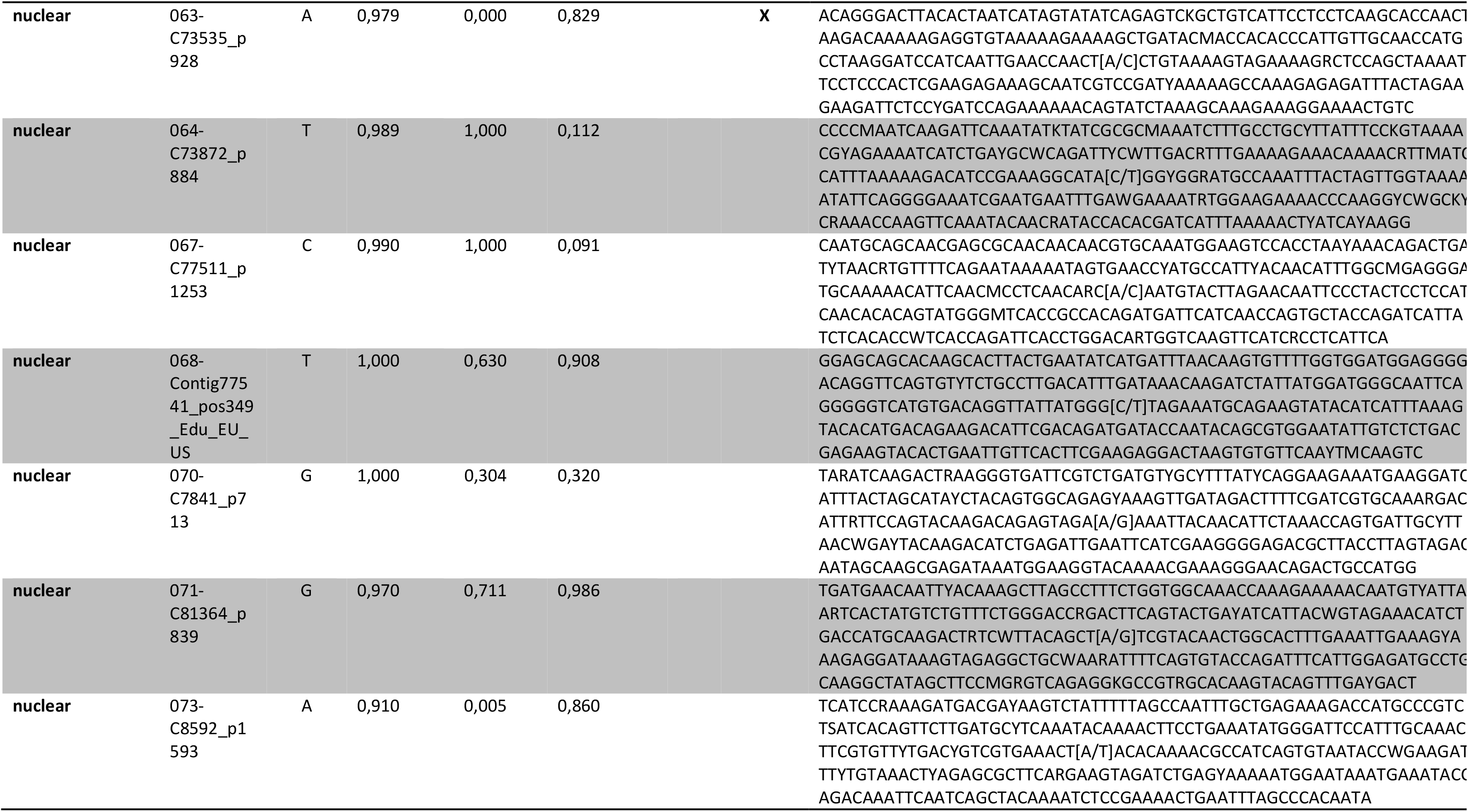
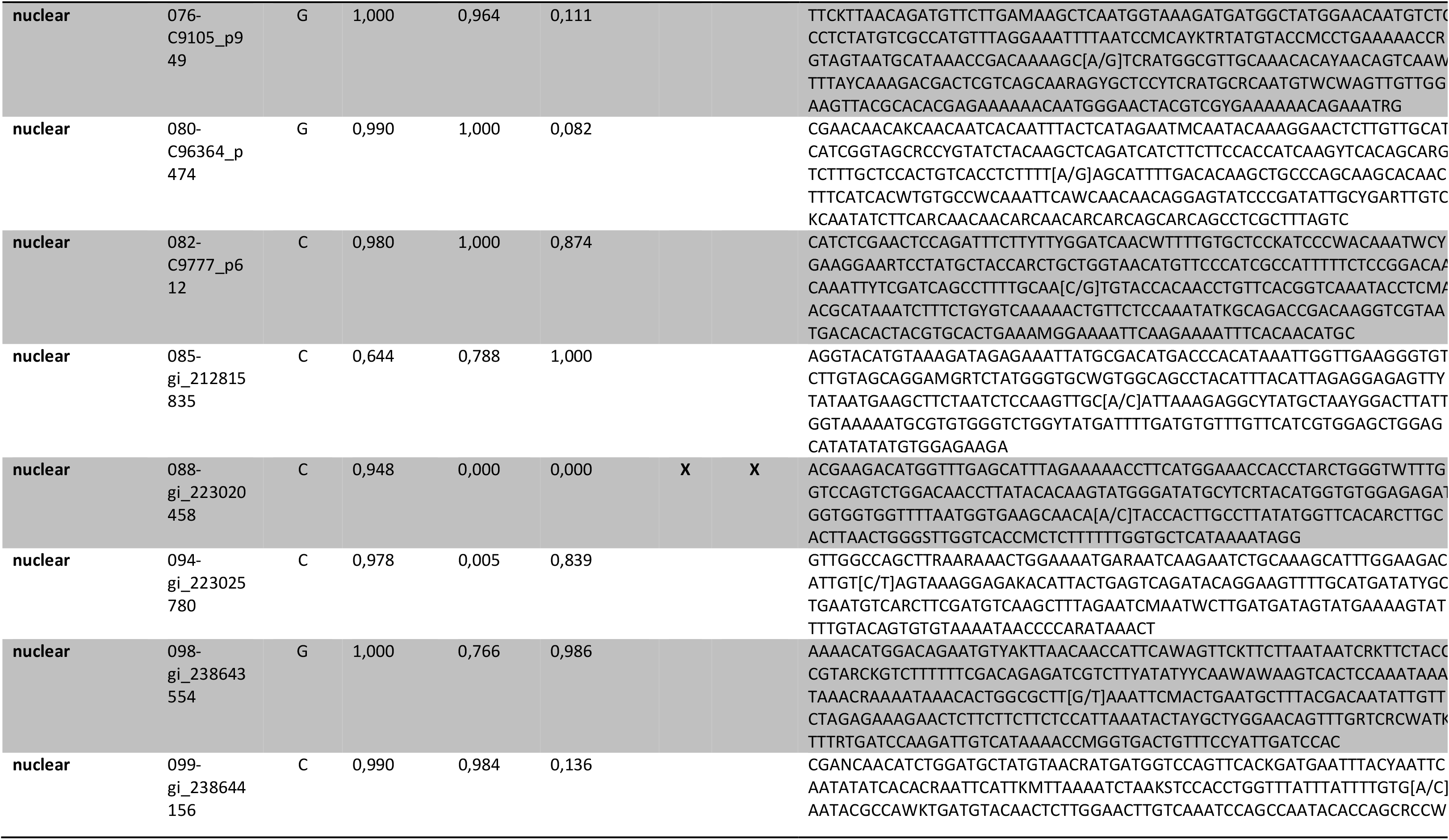
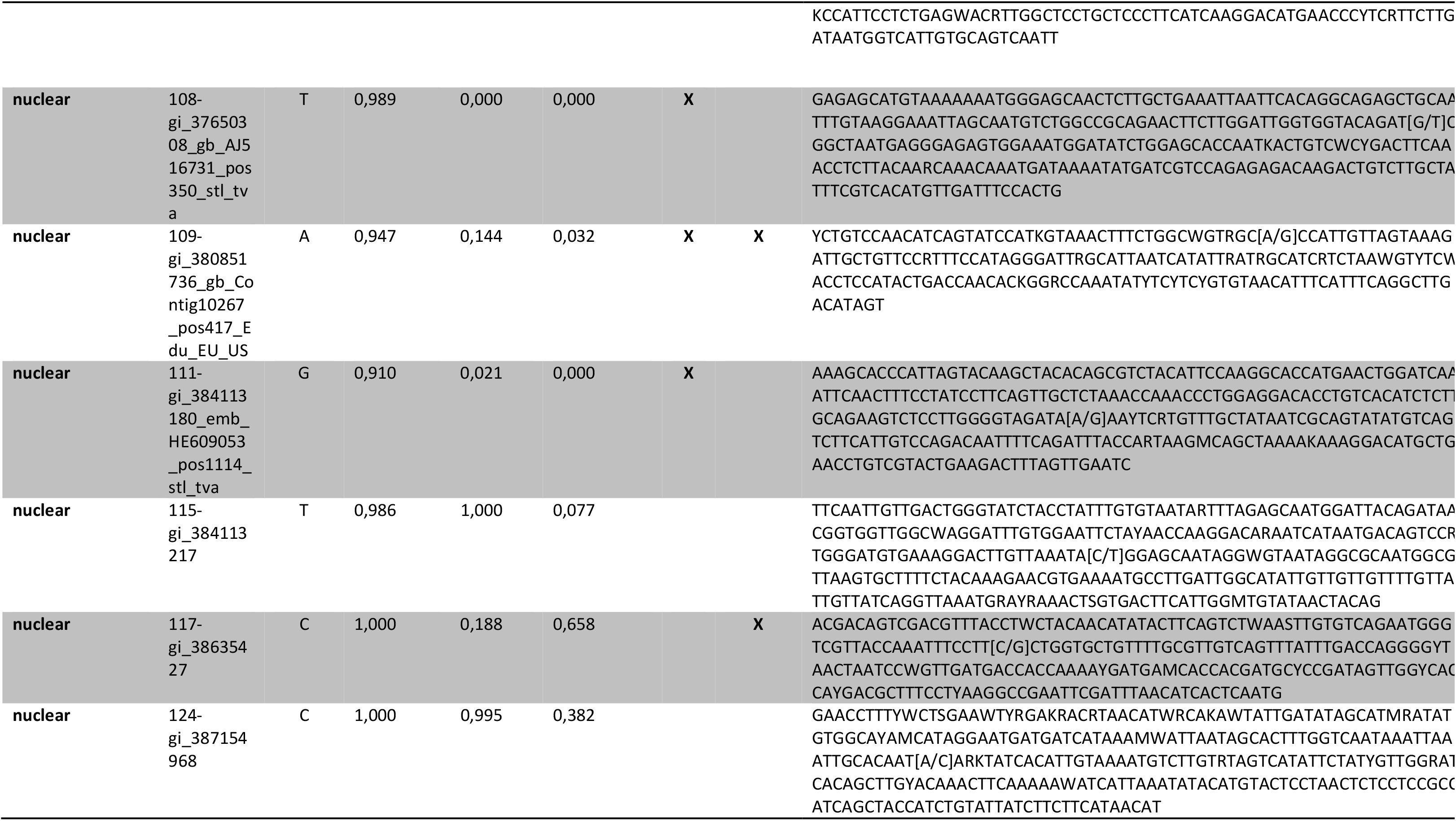
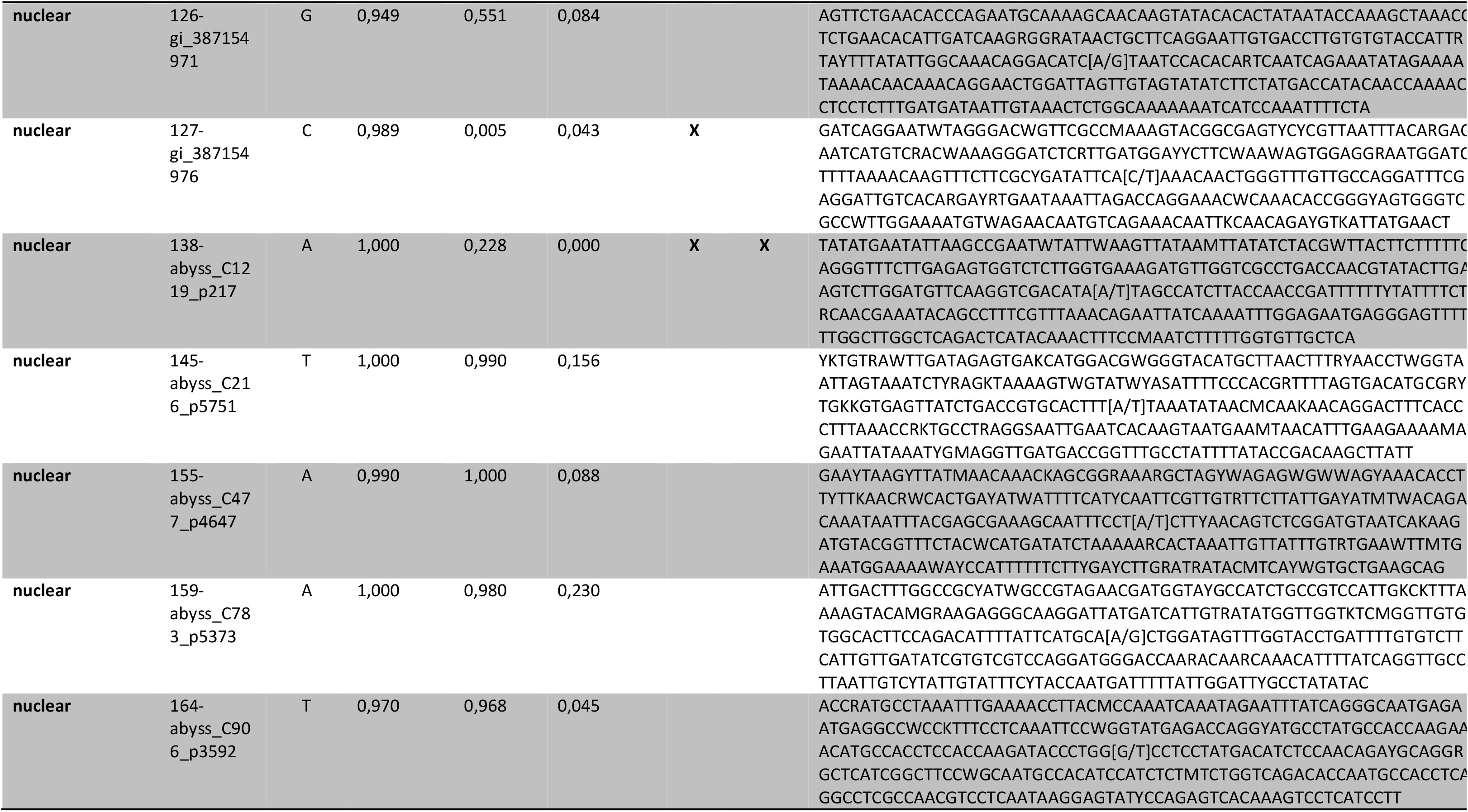
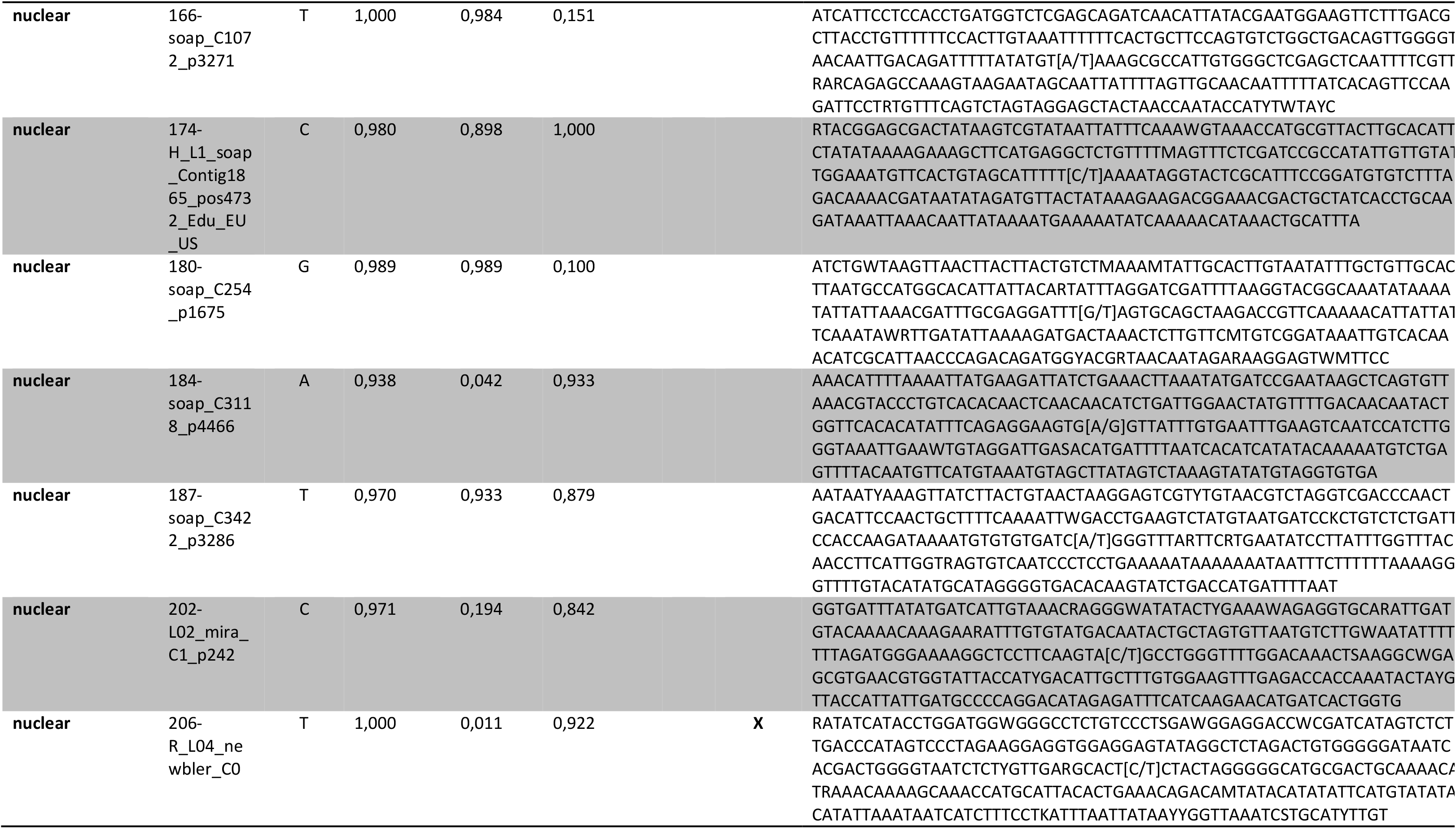
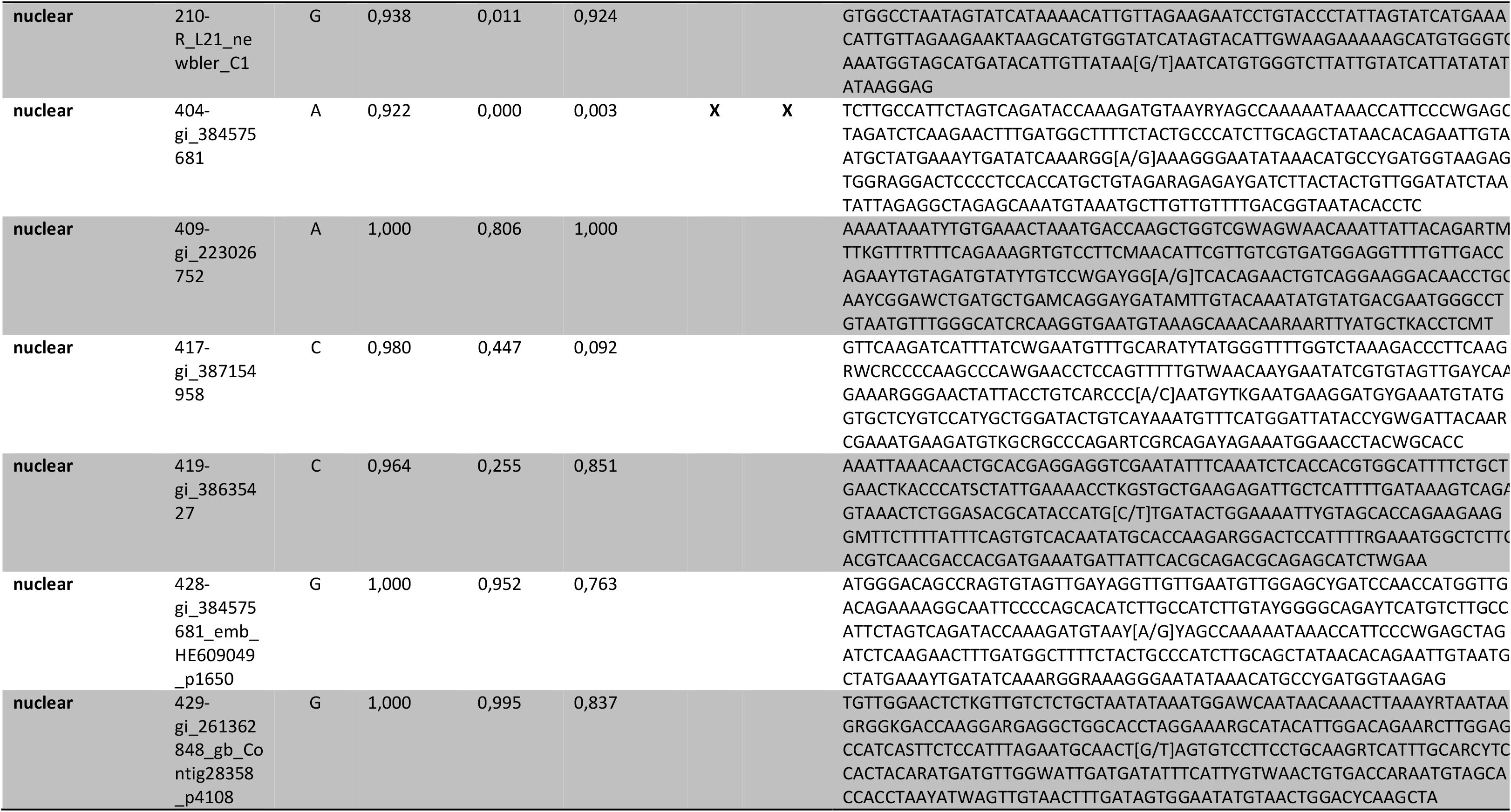
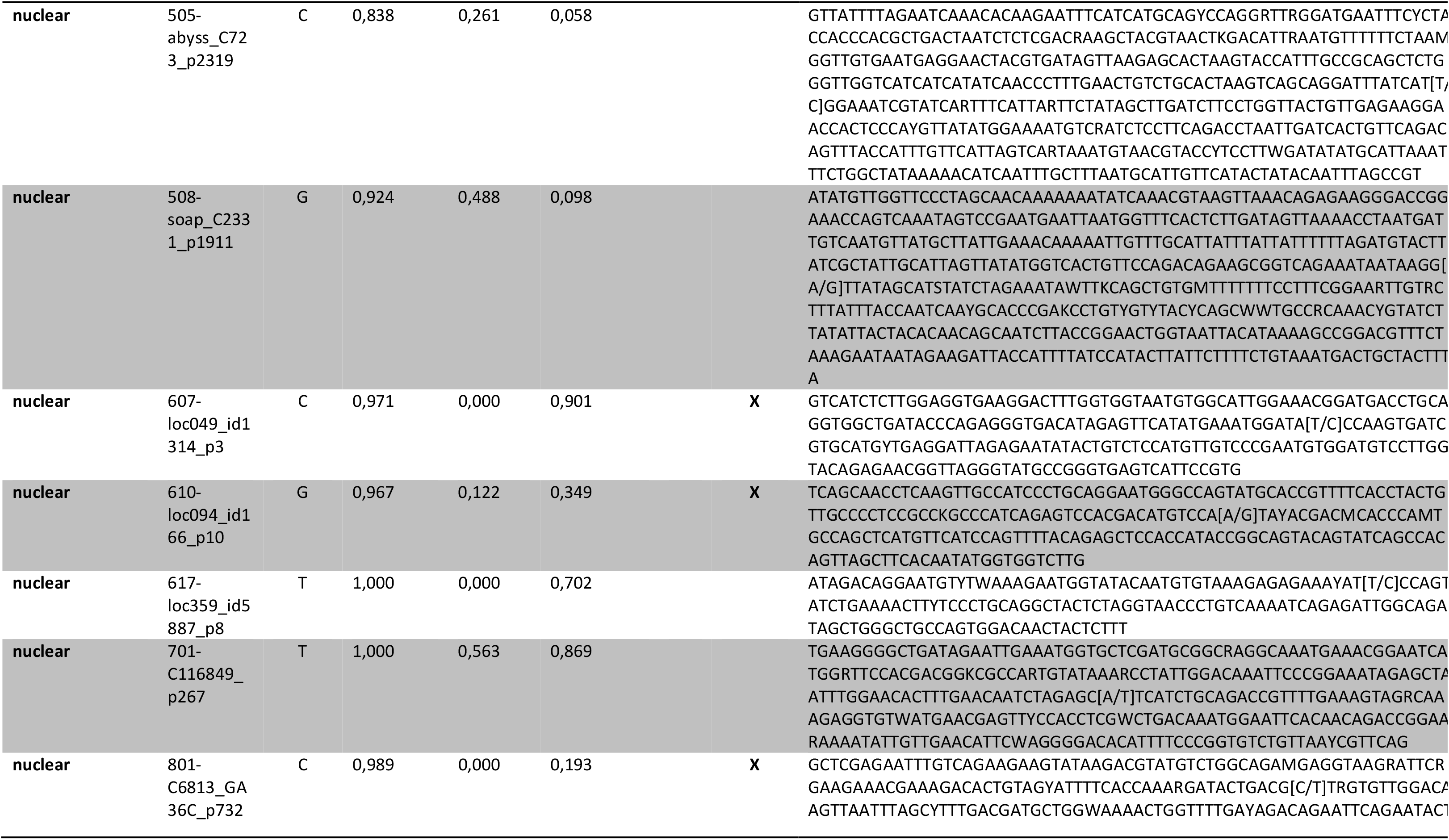
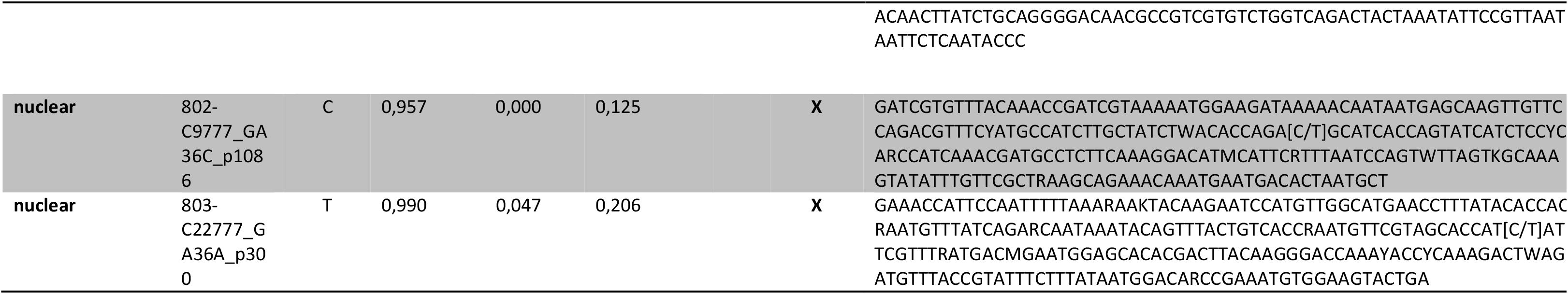
Bi-allelic SNPs informations. Freq_tross, freq_edu, freq_gallo correpsond to allele frequence in references populations; Allele: allele the most frequent in *M. trossulus* references populations; MtFF: 10 diagnostics SNPs used to estimate *M. trossulus* fluorescence fraction; MtFF2: 17 diagnostics SNPs used to estimate the proportion of cancer cells in the 6 MtrBTN2 tumors from *M. edulis* hosts; Sequence: sequence around the SNP targeted in KASP experiment, alleles are represented [allele1/allele2].

**Table S3:**
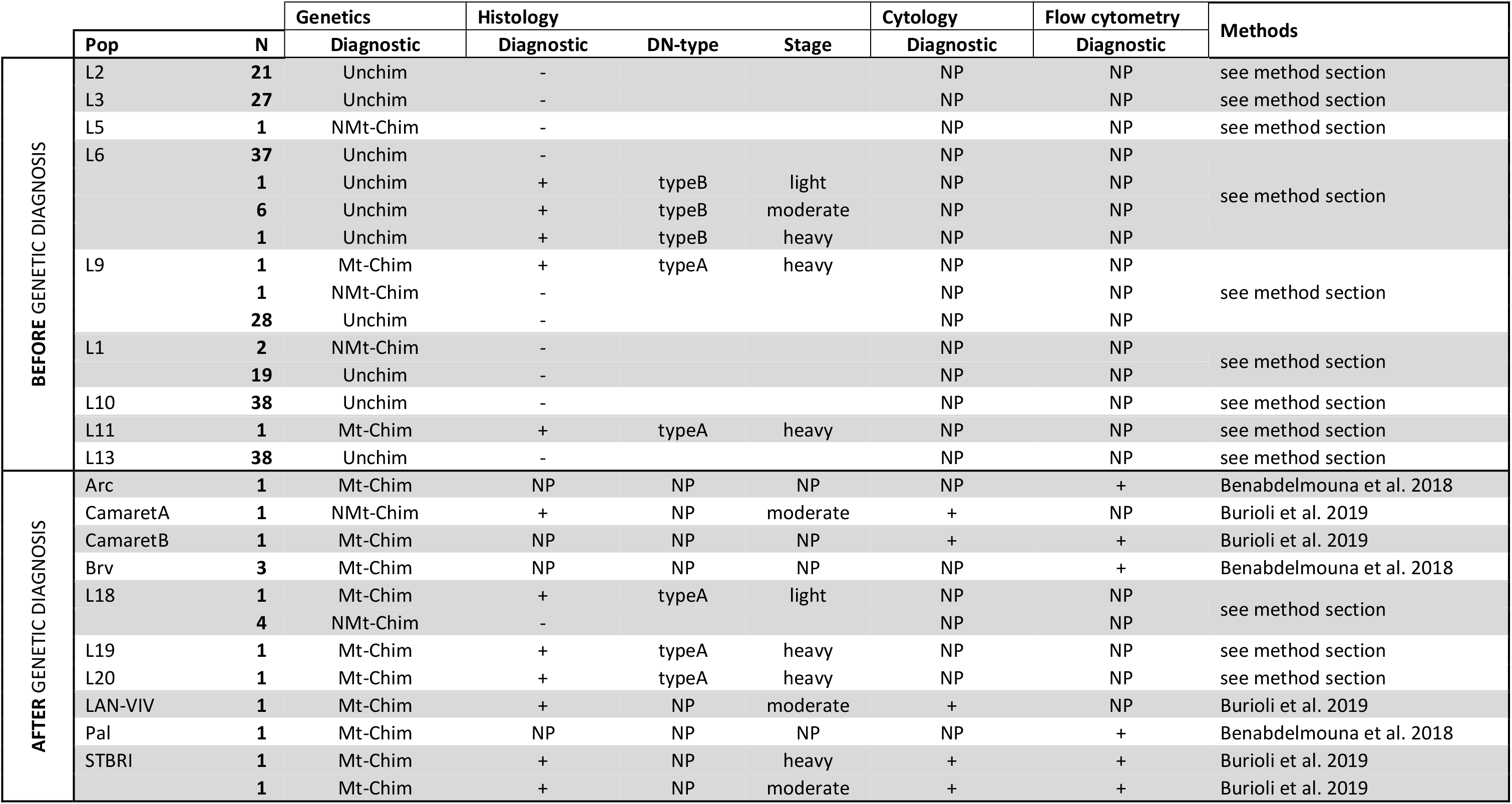
Results of additional diagnosis methods. Pop: population. N: number of individuals. Unchim: diagnosed as non-chimeric. NMt-Chim: diagnosed as non-trossulus chimeric (not due to cancer). Mt-Chim: diagnosed as trossulus chimeric (transmissible cancer, MtrBTN2). DN-type: cell type described in results section. NP: not performed.

**Figure S1: Graphical inference of the 6 MtrBTN2 tumors genotypes at 76 SNPs. (A)** KASP fluorescence data plot where genotypes are indicated by colored points. **(B)** Correlation plot between y*’_i_* fluorescence signal and MtFF2 (*M. trossulus* fluorescence fraction 2, proxy of the proportion of cancerous cell in the sample). *M. edulis* samples are represented in red (MtFF2 value close to 0) and *M. trossulus* samples are represented in yellow (MtFF2 value close to 1). In both panels, the 6 tumors are represented in black: triangle filled correspond to hemolymph samples and open triangle to mantle/gill samples (samples from the same individual are related by a pointed line).

**Figure S2:**
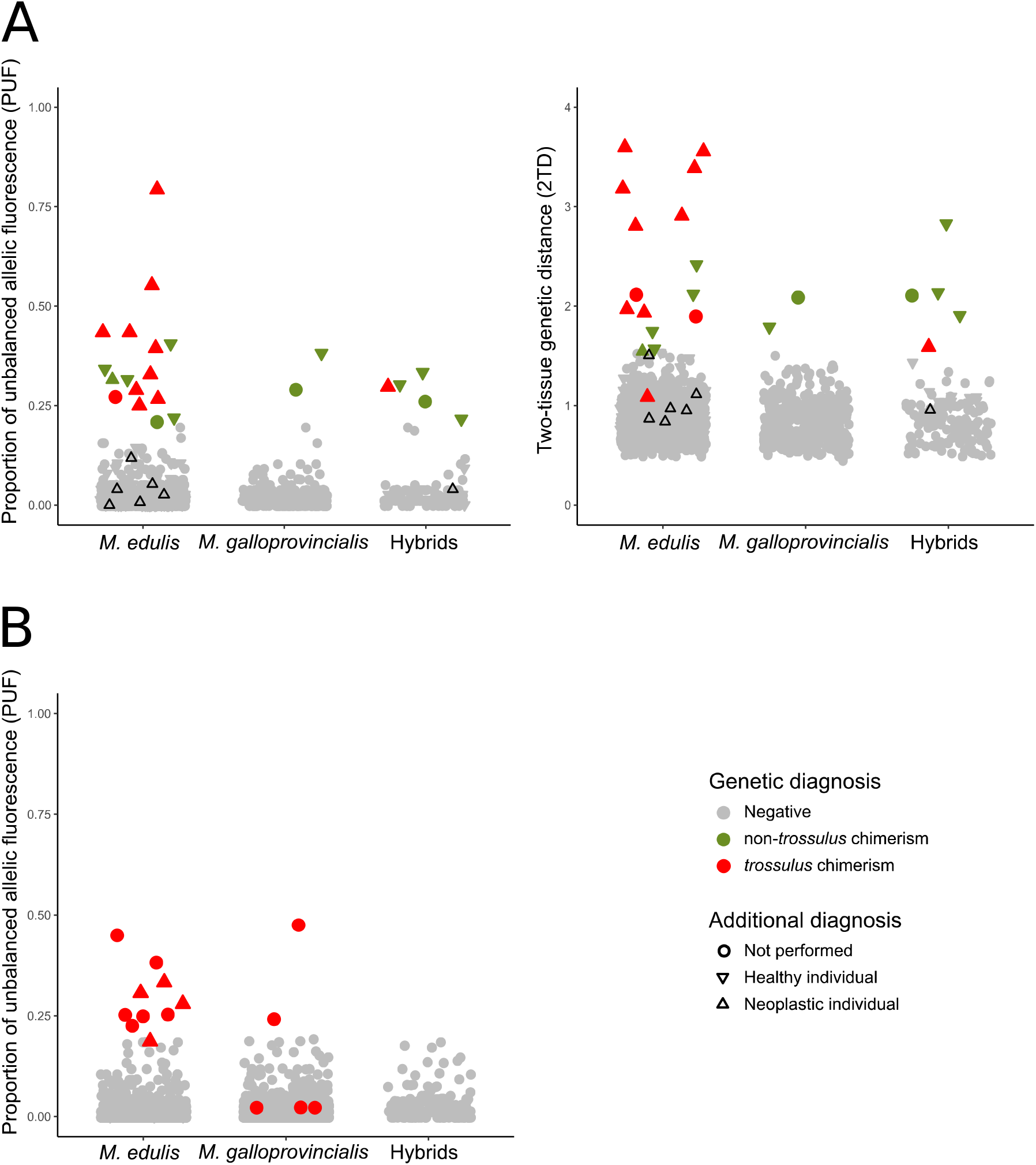
Proportion of unbalanced allelic Fluorescence (PUF) and two-tissue genetic distance (2TD) used for genetic diagnosis. **(A)** PUF (left) and 2TD (right) values for individuals sampled at two tissues. Color corresponds to genetic diagnosis: red for trossulus chimeric individuals (positive to MtF, PUF and mostly 2TD), green for non-trossulus chimeric individuals (positive to PUF and 2TD indexes) and grey for non-chimeric individuals (negative to all indexes). The symbol form corresponds to phenotypic diagnosis: triangle pointing up for neoplastic individuals, triangle pointing down for healthy individuals and dot for a not performed phenotype diagnosis. Black triangles pointing up represent neoplastic individuals of type B which are not chimeric (negative to all indexes). **(B)** PUF values for single-sampled individuals. Color corresponds to genetic diagnosis: red for trossulus chimerism (positive to MtFF) and non-chimeric individuals (negative to MtFF). The symbol form corresponds to phenotypic diagnosis: triangle pointing up for neoplastic individuals, triangle pointing down for healthy individuals and dot for a not performed phenotype diagnosis.

**Figure S3:**
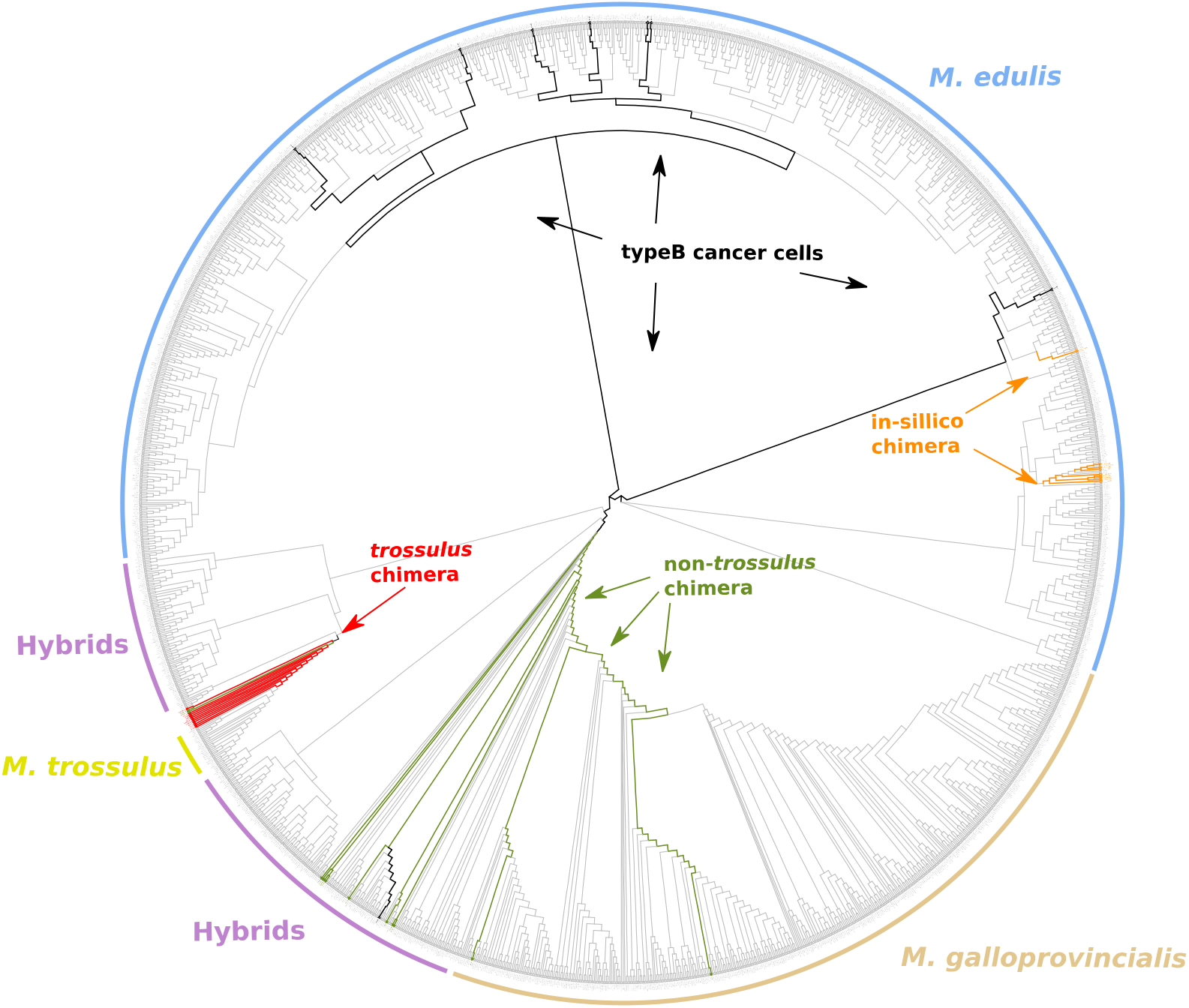
Neighbor-joining Tree estimated on all SNPs fluorescence values (*y’_i_*). Color corresponds to genetic diagnosis: red for trossulus chimerism (MtrBTN2), green for non-trossulus chimerism, grey for non-chimeric individuals and black for non-chimeric individuals with neoplastic type B cells. Orange branch corresponds to in silico chimera generated by mixing one sample (“cancer”) with 9 others samples (“hosts”) at 40%, 60% or 80% (3 of each).

**Figure S4:**
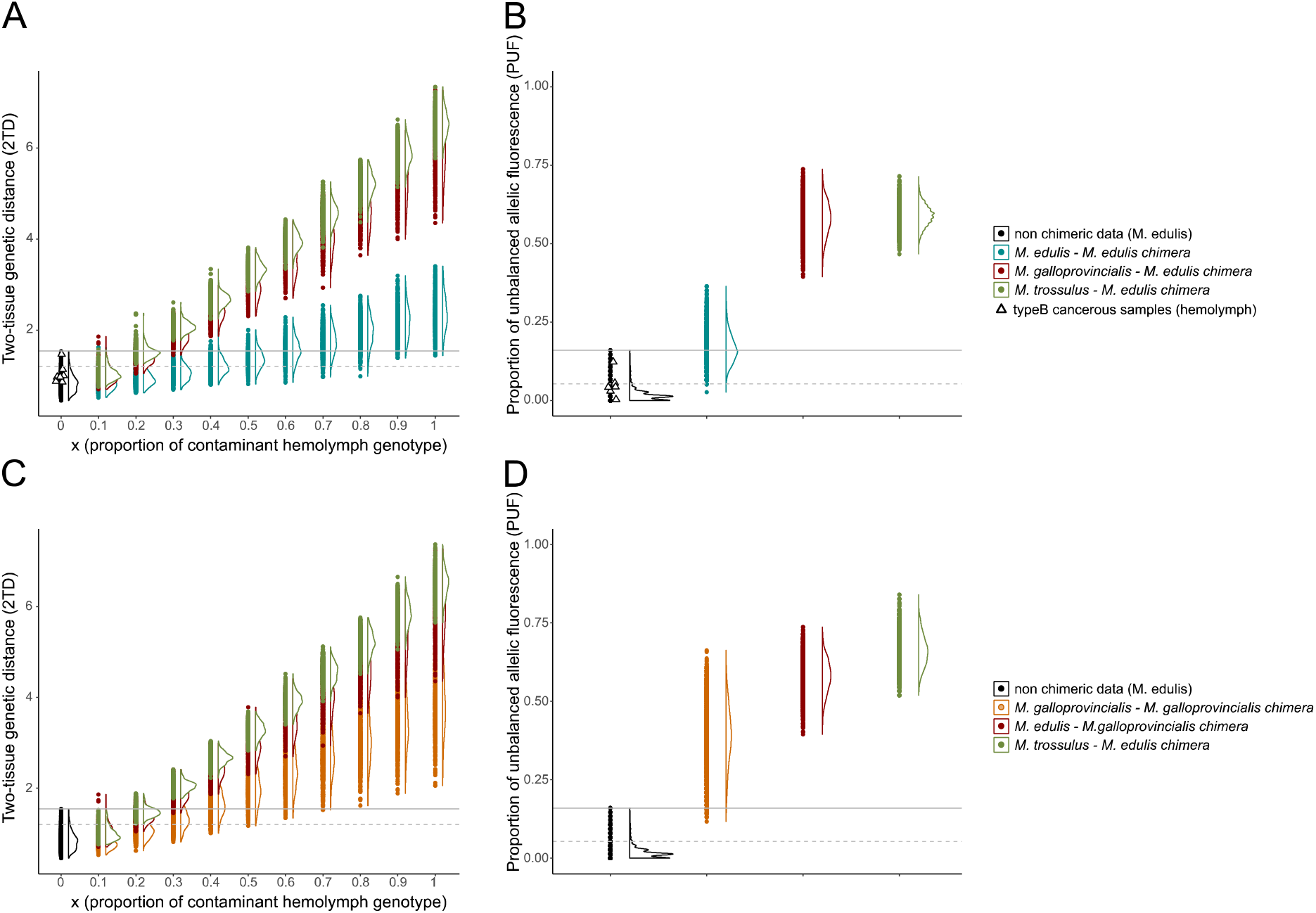
PUF and 2TD estimation of in silico chimera. Panels (A) and (B) correspond to PUF and 2TD values (respectively) estimated for in silico chimera in *M. edulis* « hosts », while panels (C) and (D) correspond to PUF and 2TD values (respectively) estimated for in-silico chimera in *M. galloprovincialis* « hosts ». Colored circles correspond to intraspecific (blue and orange circles) and interspecific (green and red circles) mixed genotypes, black circles corresponds to non-chimeric mussels performed of this study and typeB cancerous samples are reported by black triangle. Curves indicate the distribution of values and grey lines correspond to the maximum (continuous line) and percentile 95% (dotted line) of PUF and 2TD in non-chimeric individuals.

**Figure S5:**
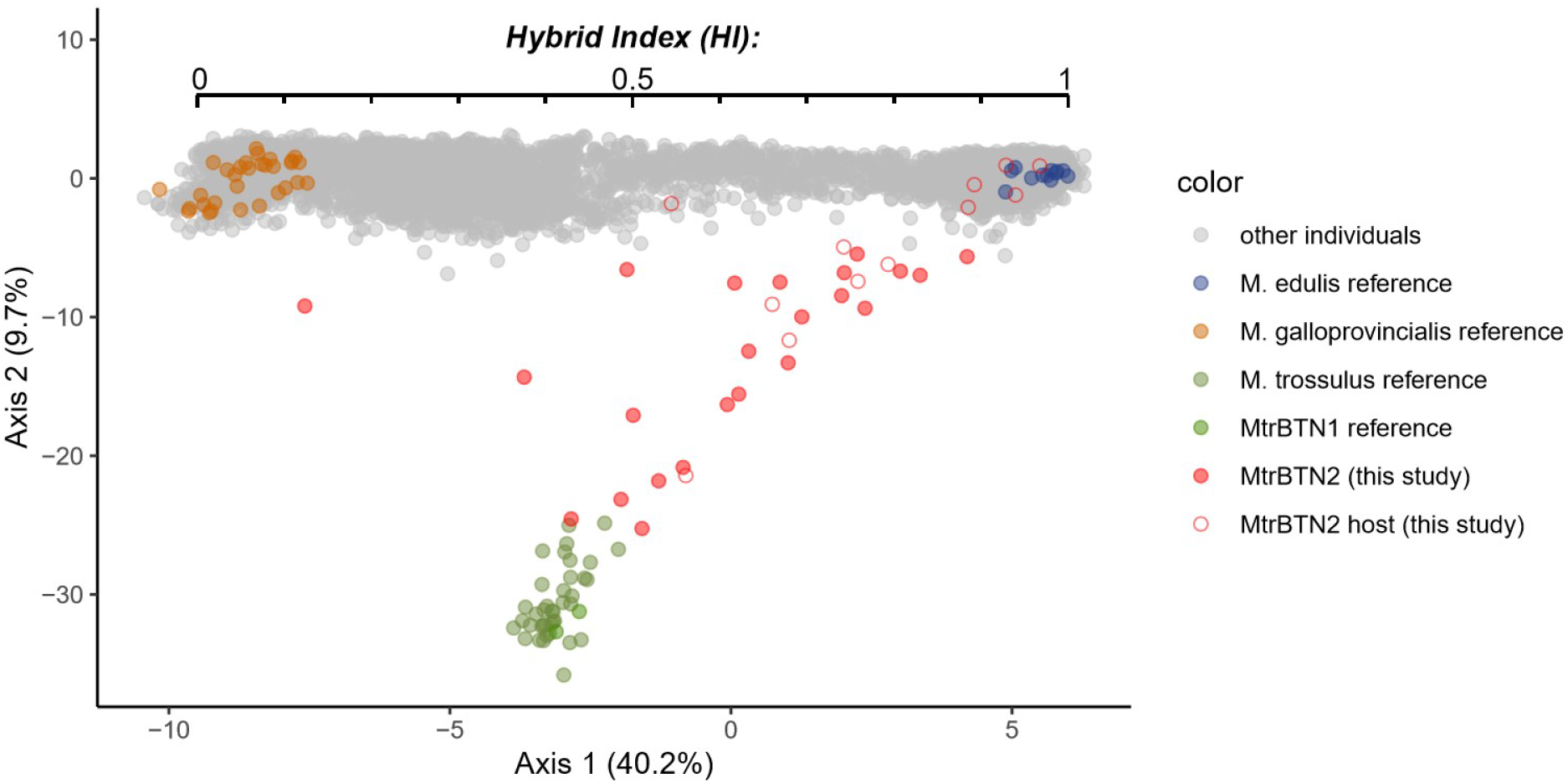
Mussels classification as *M. edulis*, *M. galloprovincialis* or hybrid. PCA on all SNP *y’_i_* values (VAFF). The first axis discriminates *M. edulis* (blue) from *M. galloprovincialis* (orange). We use it to infer a hybrid index and assign the European mussels to *M. edulis* (Hitalic>0.85), *M. galloprovincialis* (HI< 0.4) or hybrids (0.4< HI<0.85) genetic backgrounds.

**Figure S6:**
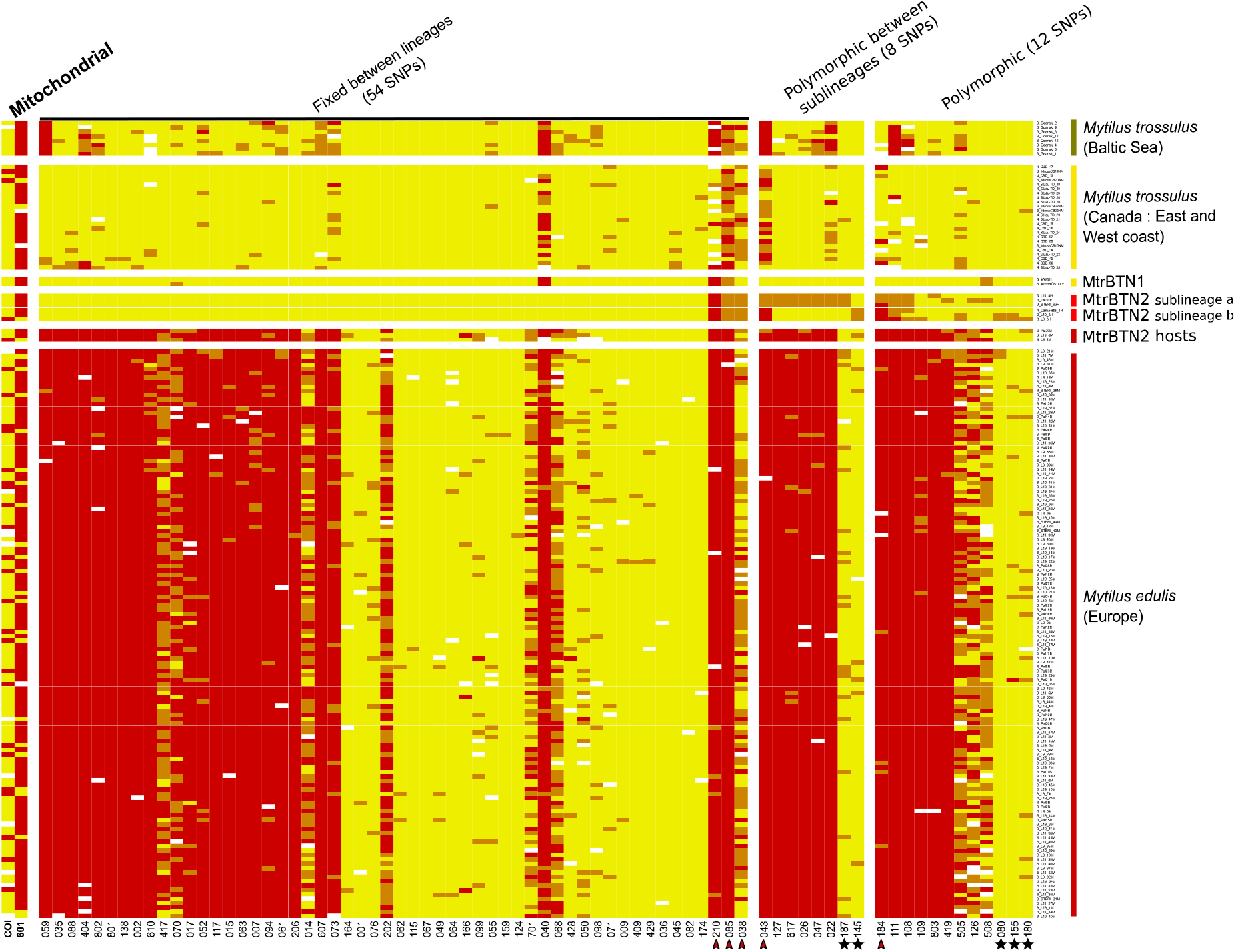
Plot of genotypes for all mitochondrial and nuclear SNPs in column and individuals in rows. Red: homozygote of *edulis*-state allele; yellow: homozygote of trossulus-state allele; orange: heterozygote. The first two columns correspond to mitochondrial SNPs, the twelve next to a subset of nuclear SNPs fixed in the 6 MtrBTN2 tumors, then eight SNPs fixed between the two sublineages and then 12 SNPs revealing polymorphism within each sublineages. Red arrowhead pointed the five SNPs with *M. edulis*-state allele excess in the 6 tumors. Black star highlights the five SNPs for which *M.edulis*-state allele in tumors are not explained by the null allele hypothesis.

