## Supplementary Figure 1 for "Prevalence and polymorphism of a mussel transmissible cancer in Europe"

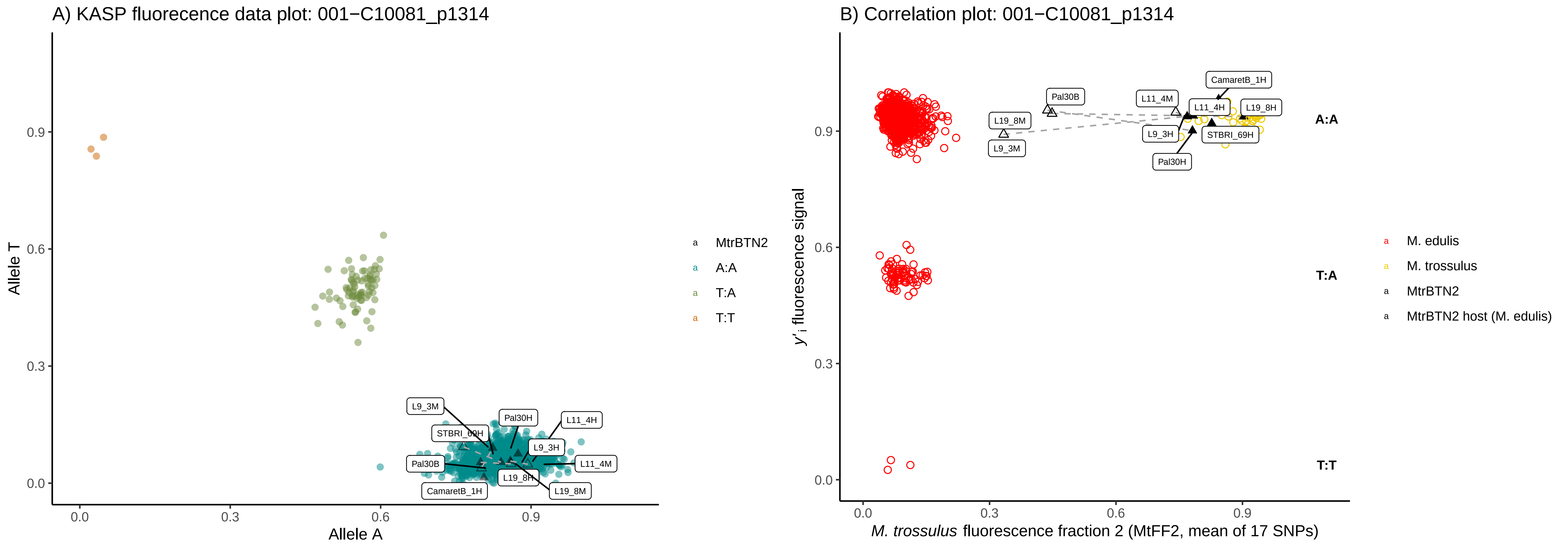
A) KASP fluorecence data plot: 001−C10081_p1314
Allele T
Allele A
0.0
0.3
0.6
0.9
L9_3M
Pal30H
L11_4H
STBRI_69H
L9_3H
Pal30B
L11_4M
L19_8H
CamaretB_1H
L19_8M
MtrBTN2
A:A
T:A
T:T
B) Correlation plot: 001−C10081_p1314
y'i fluorescence signal
M. trossulus fluorescence fraction 2 (MtFF2, mean of 17 SNPs)
CamaretB_1H
Pal30B
L11_4M
L11_4H
L19_8H
L19_8M
L9_3H
STBRI_69H
L9_3M
Pal30H
A:A
T:A
T:T
M. edulis
M. trossulus
MtrBTN2
MtrBTN2 host (M. edulis)

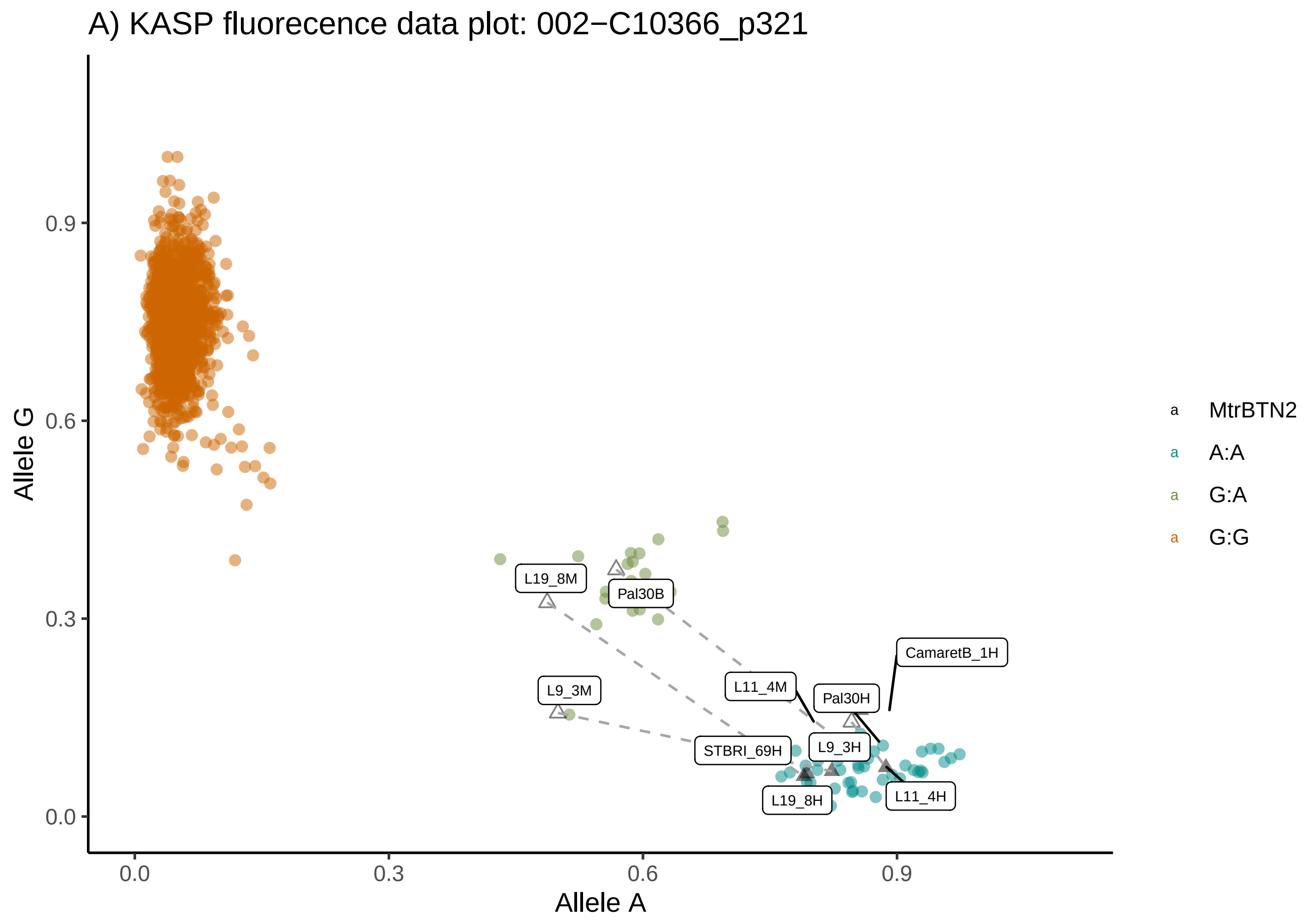
A) KASP fluorecence data plot: 002−C10366_p321
Allele G
Allele A
0.0
0.3
0.6
0.9
L19_8M
Pal30B
L9_3M
L11_4M
Pal30H
CamaretB_1H
STBRI_69H
L9_3H
L19_8H
L11_4H
MtrBTN2
A:A
G:A
G:G

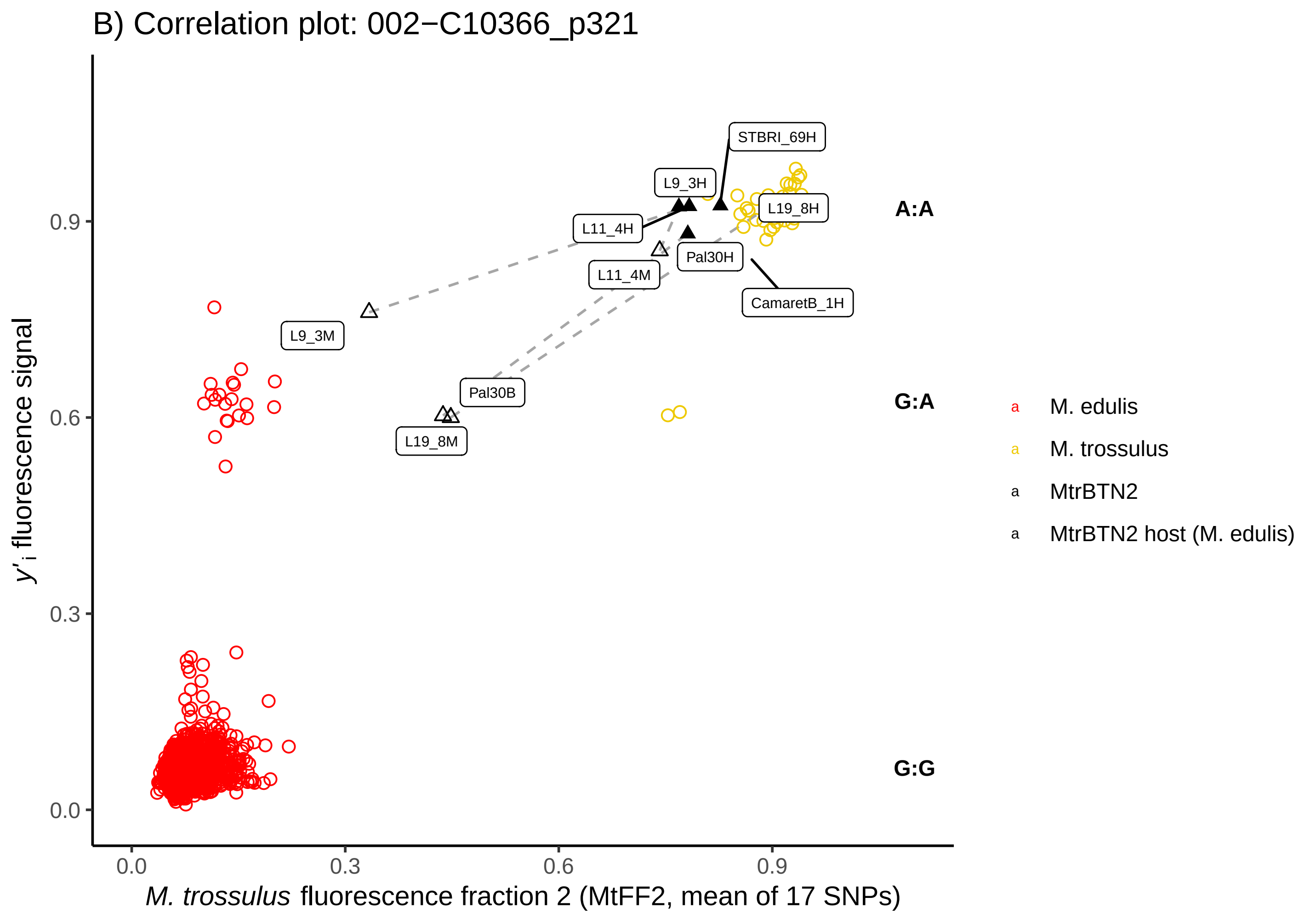
B) Correlation plot: 002−C10366_p321
y'i fluorescence signal
M. trossulus fluorescence fraction 2 (MtFF2, mean of 17 SNPs)
0.0
0.3
0.6
0.9
STBRI_69H
L9_3H
L19_8H
L11_4H
Pal30H
L11_4M
CamaretB_1H
L9_3M
Pal30B
L19_8M
A:A
G:A
G:G
M. edulis
M. trossulus
MtrBTN2
MtrBTN2 host (M. edulis)

A) KASP fluorecence data plot: 007−C14012_p60

B) Correlation plot: 007−C14012_p60

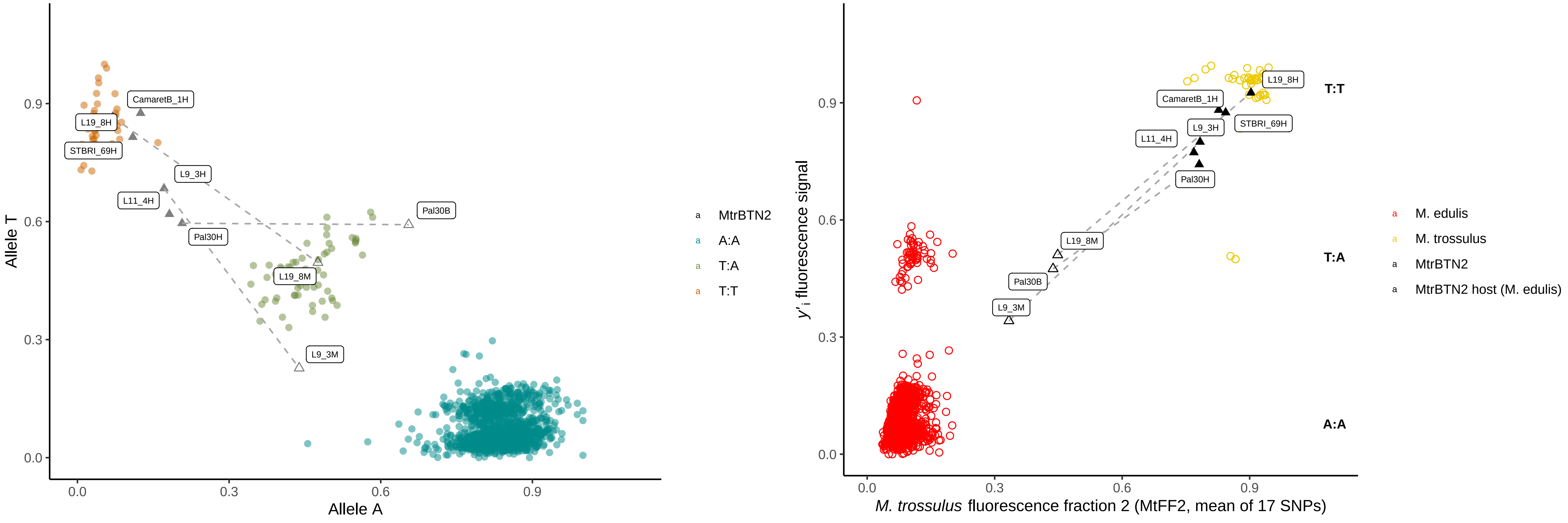

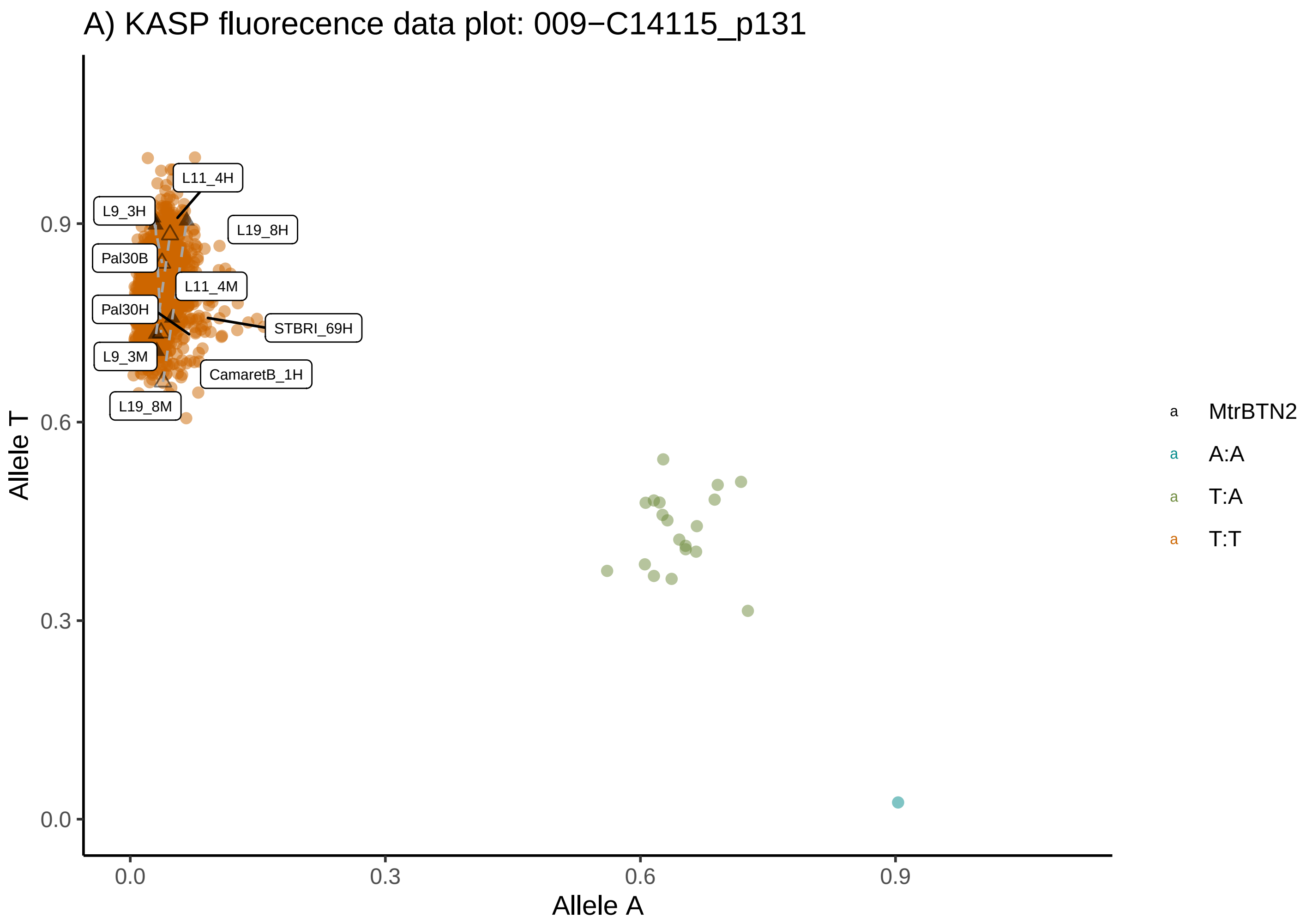
A) KASP fluorecence data plot: 009−C14115_p131
Allele T
Allele A
0.0
0.3
0.6
0.9
L11_4H
L9_3H
L19_8H
Pal30B
L11_4M
Pal30H
STBRI_69H
L9_3M
CamaretB_1H
L19_8M
MtrBTN2
A:A
T:A
T:T

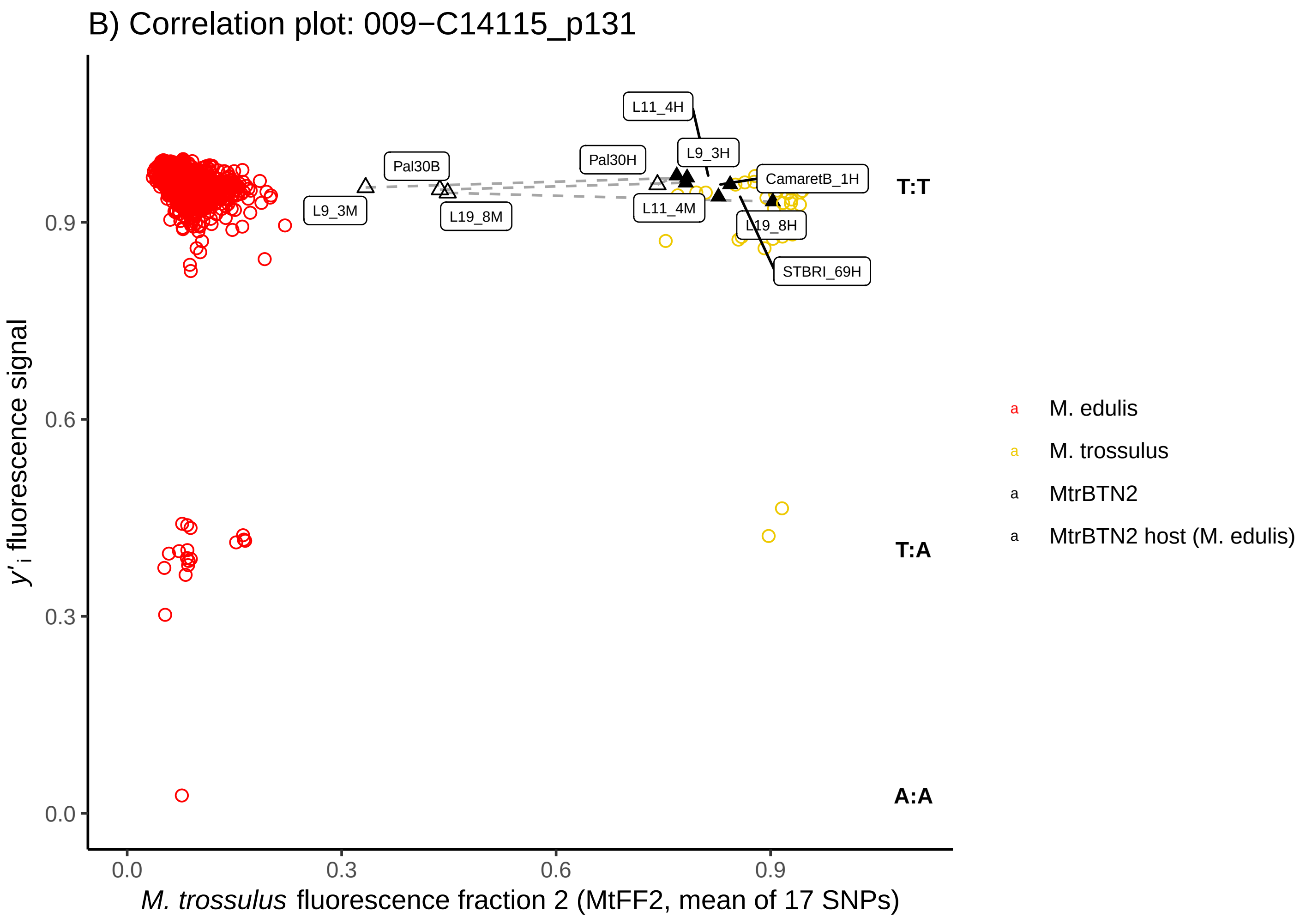
B) Correlation plot: 009−C14115_p131
y'i fluorescence signal
M. trossulus fluorescence fraction 2 (MtFF2, mean of 17 SNPs)
0.0
0.3
0.6
0.9
L11_4H
L9_3H
Pal30H
Pal30B
CamaretB_1H
T:T
L9_3M
L19_8M
L11_4M
L19_8H
STBRI_69H
T:A
A:A
M. edulis
M. trossulus
MtrBTN2
MtrBTN2 host (M. edulis)

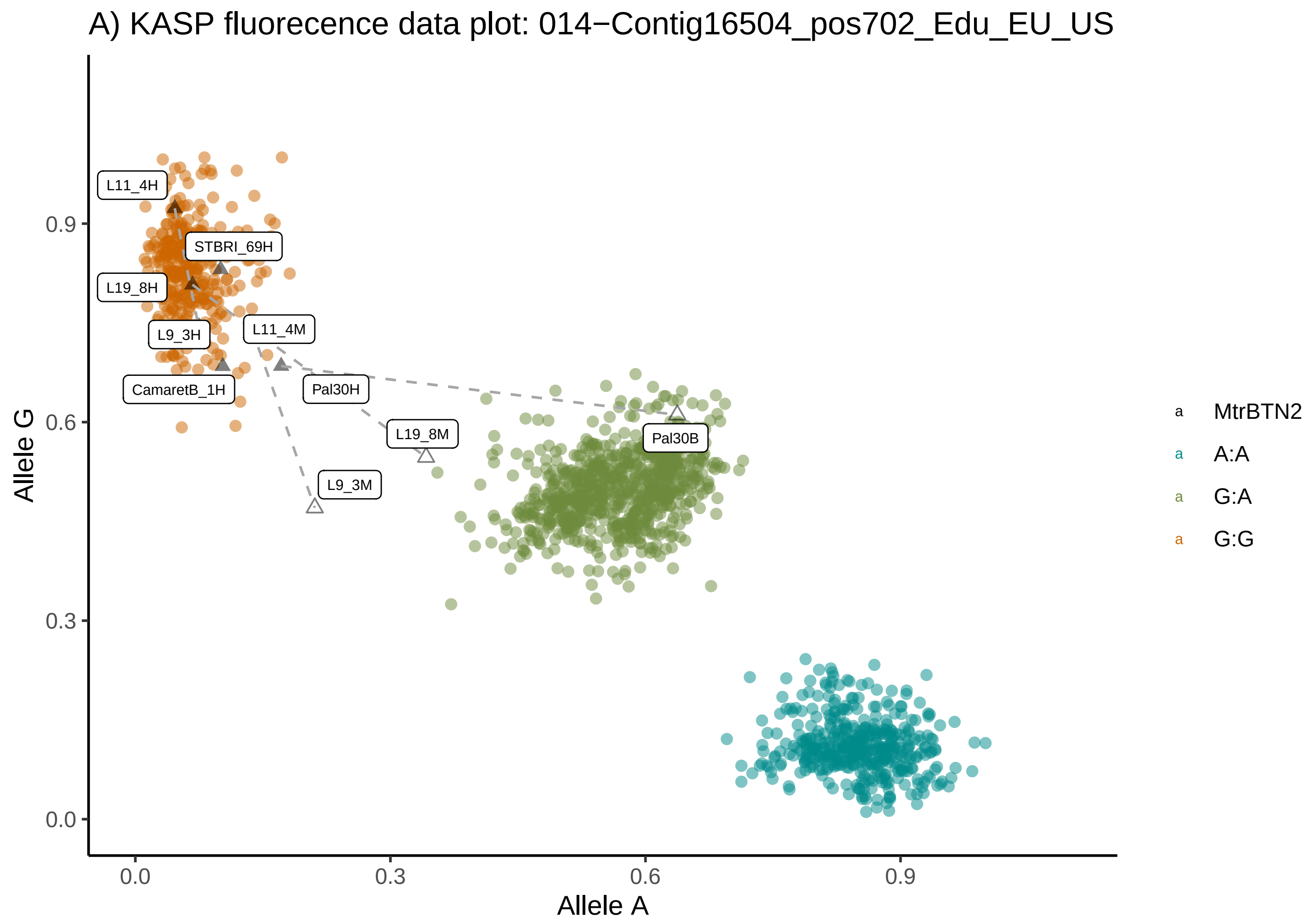
A) KASP fluorecence data plot: 014−Contig16504_pos702_Edu_EU_US
Allele G
Allele A
0.0
0.3
0.6
0.9
L11_4H
STBRI_69H
L19_8H
L9_3H
L11_4M
CamaretB_1H
Pal30H
L19_8M
L9_3M
Pal30B
MtrBTN2
A:A
G:A
G:G

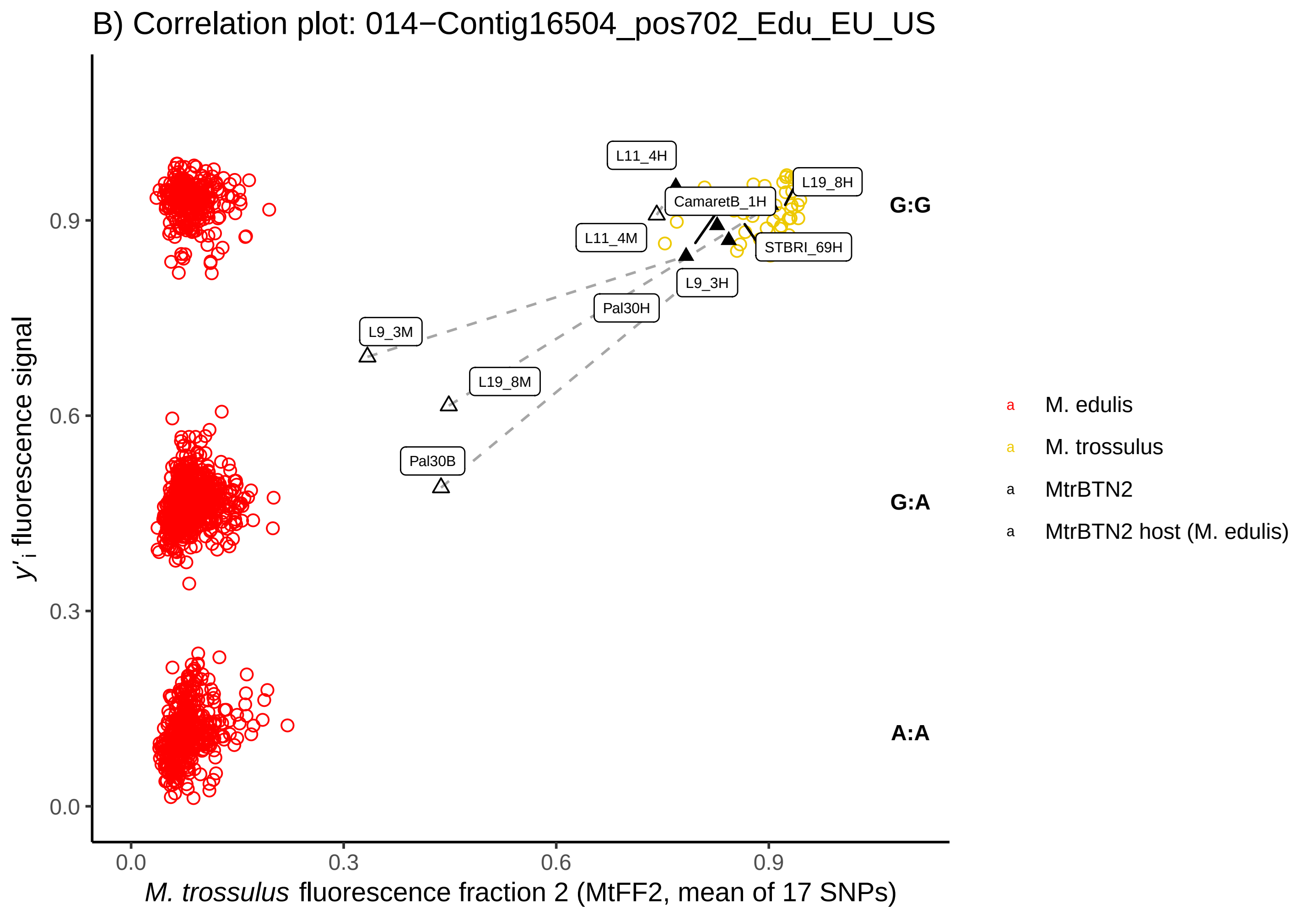
B) Correlation plot: 014−Contig16504_pos702_Edu_EU_US
y'i fluorescence signal
M. trossulus fluorescence fraction 2 (MtFF2, mean of 17 SNPs)
0.0
0.3
0.6
0.9
L11_4H
L19_8H
CamaretB_1H
L11_4M
STBRI_69H
L9_3H
Pal30H
L9_3M
L19_8M
Pal30B
G:G
G:A
A:A
M. edulis
M. trossulus
MtrBTN2
MtrBTN2 host (M. edulis)

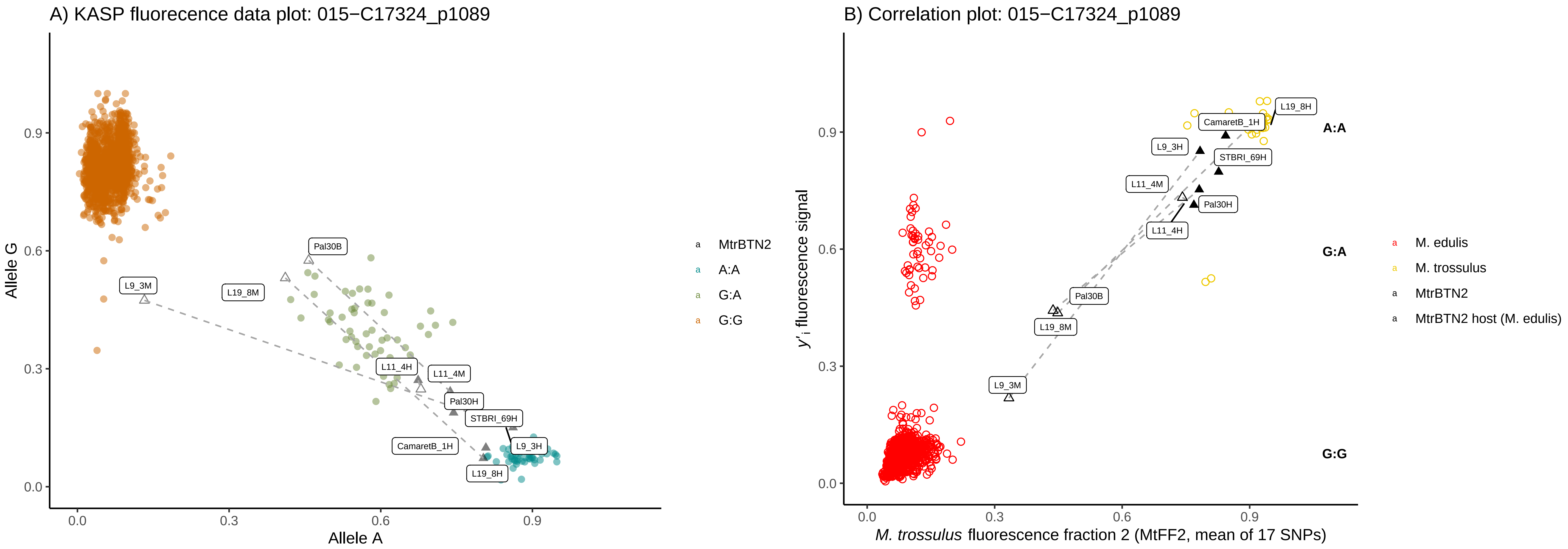
A) KASP fluorecence data plot: 015−C17324_p1089
Allele G
Allele A
0.0
0.3
0.6
0.9
Pal30B
L9_3M
L19_8M
L11_4H
L11_4M
Pal30H
STBRI_69H
CamaretB_1H
L9_3H
L19_8H
MtrBTN2
A:A
G:A
G:G
B) Correlation plot: 015−C17324_p1089
y'i fluorescence signal
M. trossulus fluorescence fraction 2 (MtFF2, mean of 17 SNPs)
L19_8H
CamaretB_1H
L9_3H
STBRI_69H
L11_4M
Pal30H
L11_4H
Pal30B
L19_8M
L9_3M
A:A
G:A
G:G
M. edulis
M. trossulus
MtrBTN2
MtrBTN2 host (M. edulis)

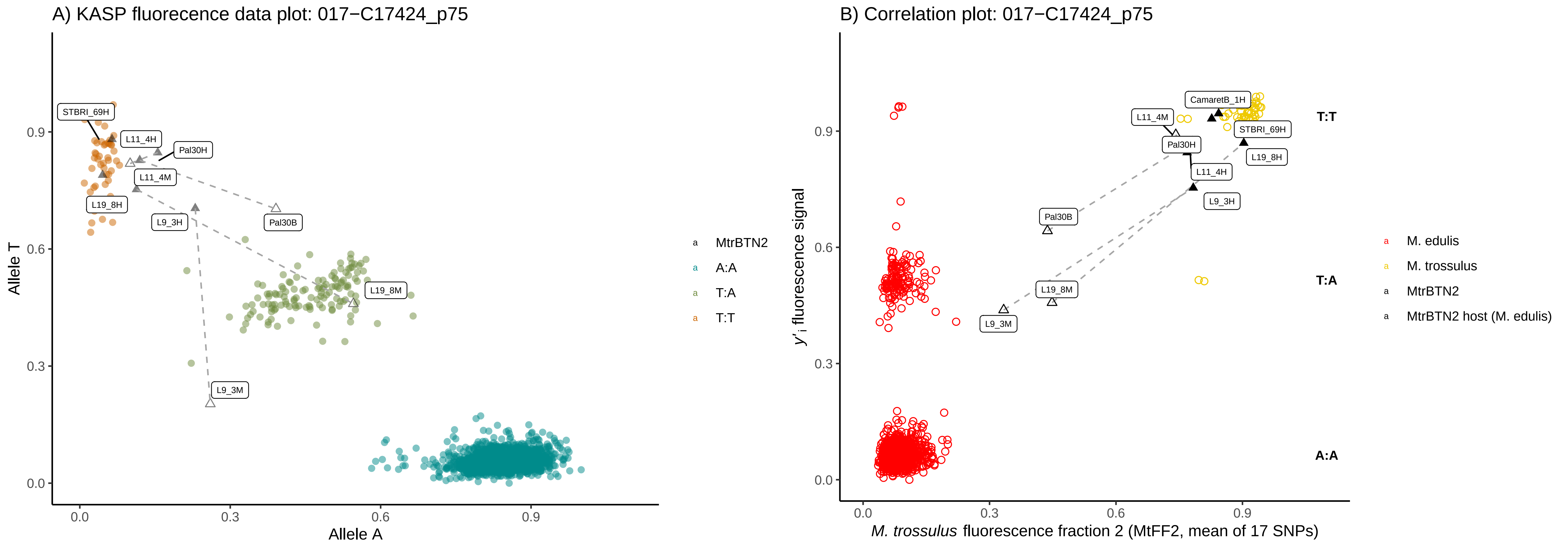
A) KASP fluorecence data plot: 017−C17424_p75
Allele T
Allele A
0.0
0.3
0.6
0.9
STBRI_69H
L11_4H
Pal30H
L11_4M
L19_8H
L9_3H
Pal30B
L19_8M
L9_3M
MtrBTN2
A:A
T:A
T:T
B) Correlation plot: 017−C17424_p75
y'i fluorescence signal
M. trossulus fluorescence fraction 2 (MtFF2, mean of 17 SNPs)
CamaretB_1H
L11_4M
STBRI_69H
Pal30H
L19_8H
L11_4H
L9_3H
Pal30B
L19_8M
L9_3M
T:T
T:A
A:A
M. edulis
M. trossulus
MtrBTN2
MtrBTN2 host (M. edulis)

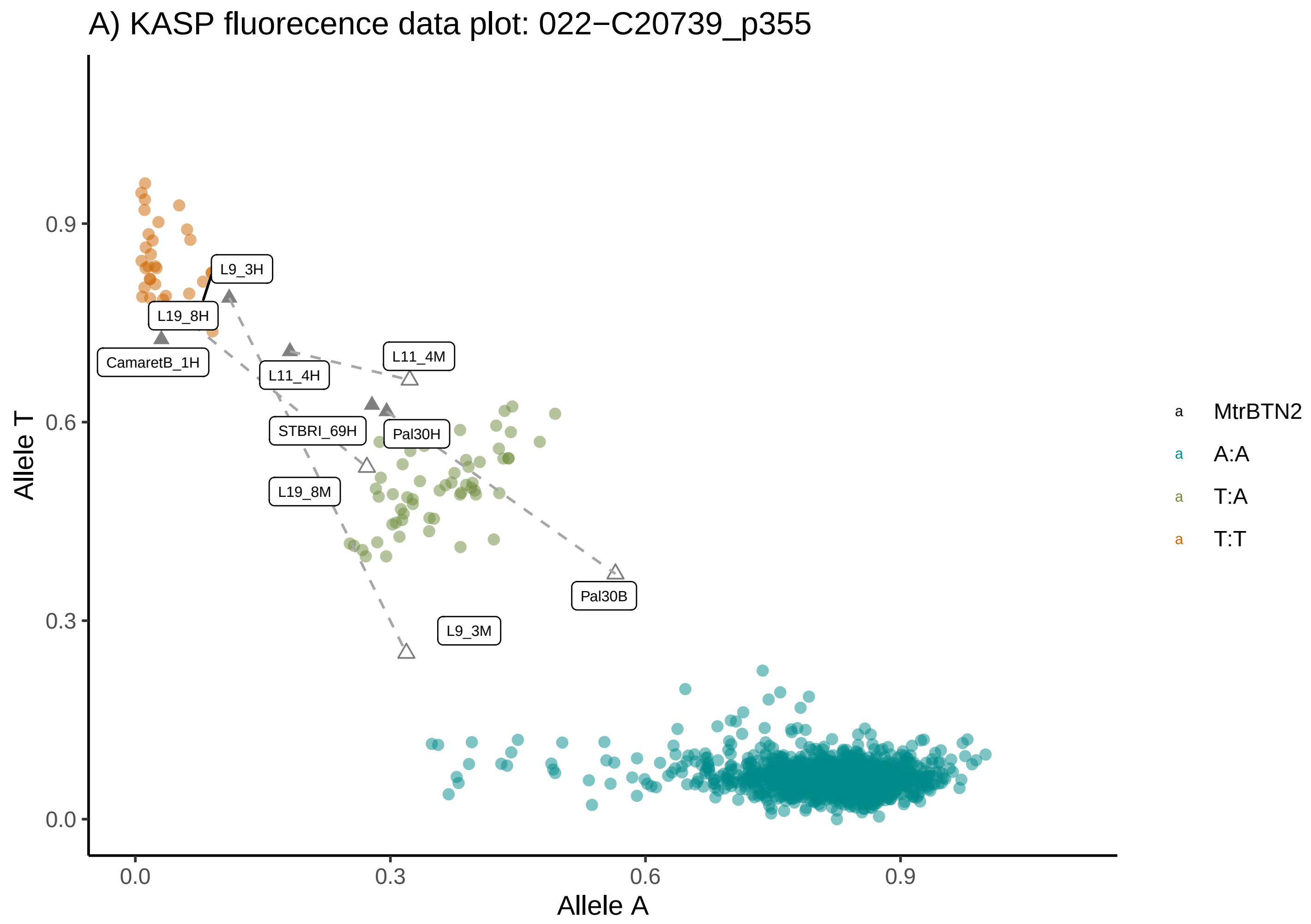
A) KASP fluorecence data plot: 022−C20739_p355
Allele T
Allele A
0.0
0.3
0.6
0.9
CamaretB_1H
L19_8H
L9_3H
L11_4H
L11_4M
STBRI_69H
Pal30H
L19_8M
Pal30B
L9_3M
MtrBTN2
A:A
T:A
T:T

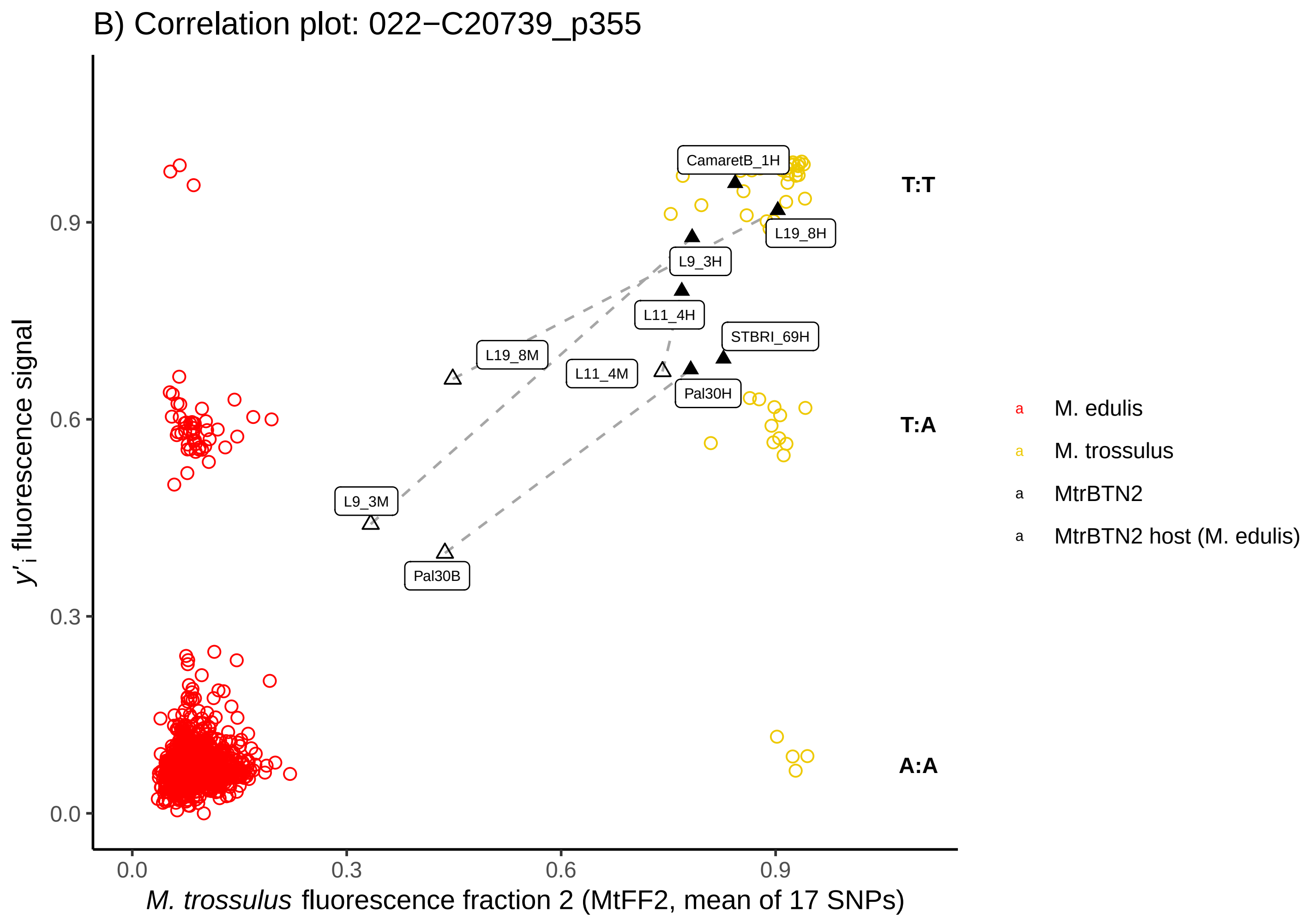
B) Correlation plot: 022−C20739_p355
y'i fluorescence signal
M. trossulus fluorescence fraction 2 (MtFF2, mean of 17 SNPs)
0.0
0.3
0.6
0.9
CamaretB_1H
L19_8H
L9_3H
L11_4H
STBRI_69H
L19_8M
L11_4M
Pal30H
L9_3M
Pal30B
T:T
T:A
A:A
M. edulis
M. trossulus
MtrBTN2
MtrBTN2 host (M. edulis)

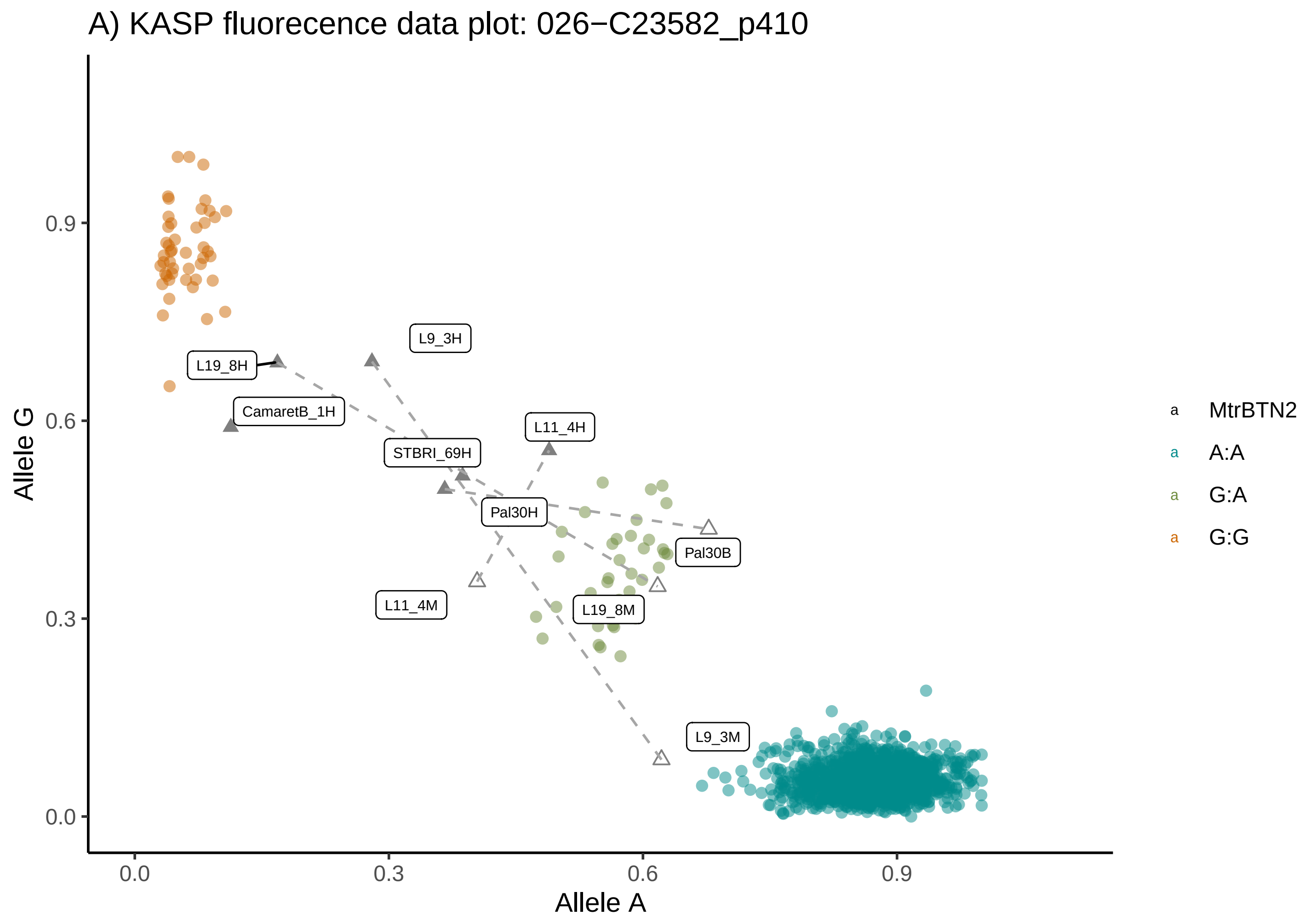
A) KASP fluorecence data plot: 026−C23582_p410
Allele G
Allele A
0.0
0.3
0.6
0.9
L9_3H
L19_8H
CamaretB_1H
L11_4H
STBRI_69H
Pal30H
Pal30B
L11_4M
L19_8M
L9_3M
MtrBTN2
A:A
G:A
G:G

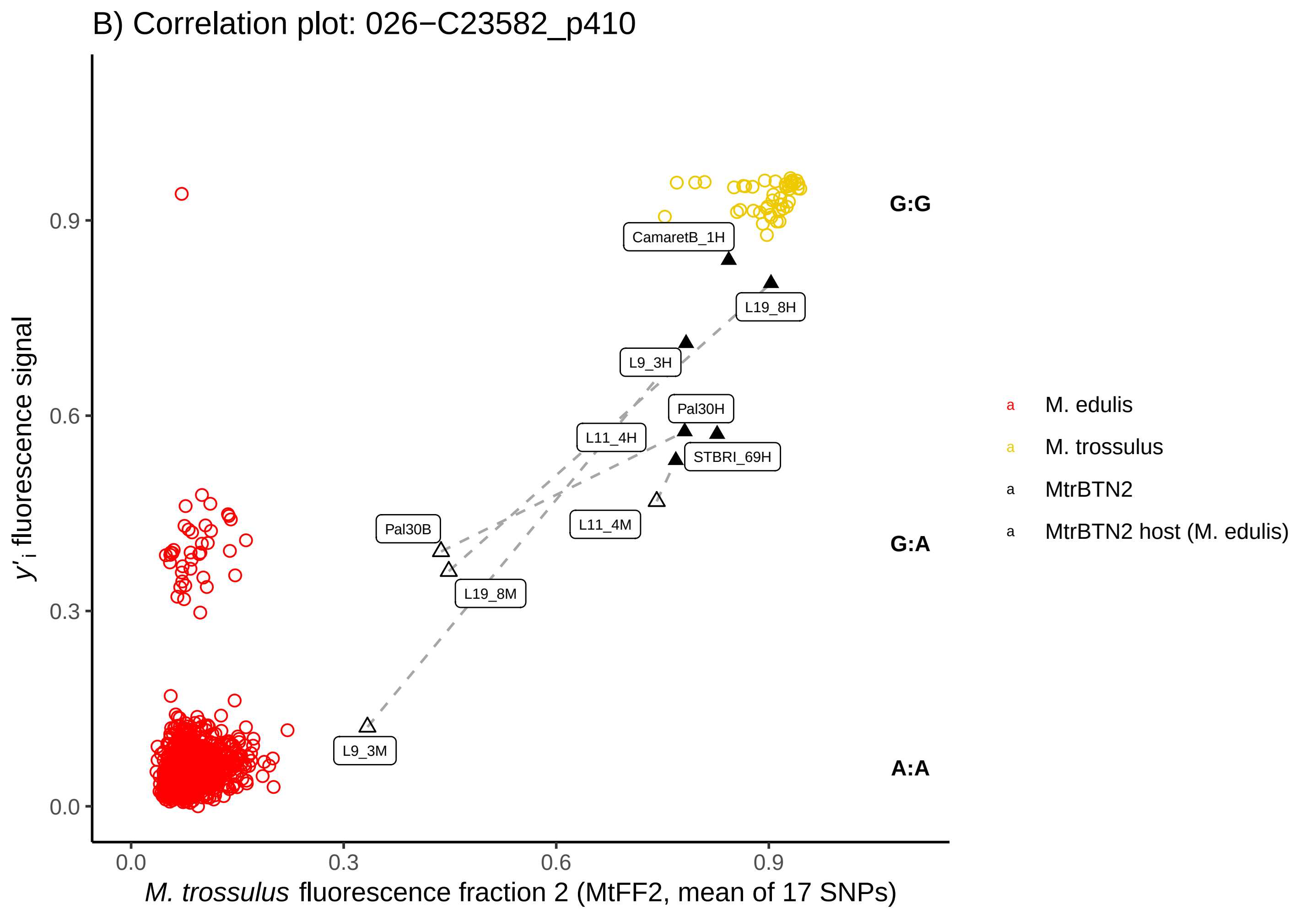
B) Correlation plot: 026−C23582_p410
y'i fluorescence signal
M. trossulus fluorescence fraction 2 (MtFF2, mean of 17 SNPs)
0.0
0.3
0.6
0.9
G:G
G:A
A:A
CamaretB_1H
L19_8H
L9_3H
Pal30H
L11_4H
STBRI_69H
Pal30B
L11_4M
L19_8M
L9_3M
M. edulis
M. trossulus
MtrBTN2
MtrBTN2 host (M. edulis)

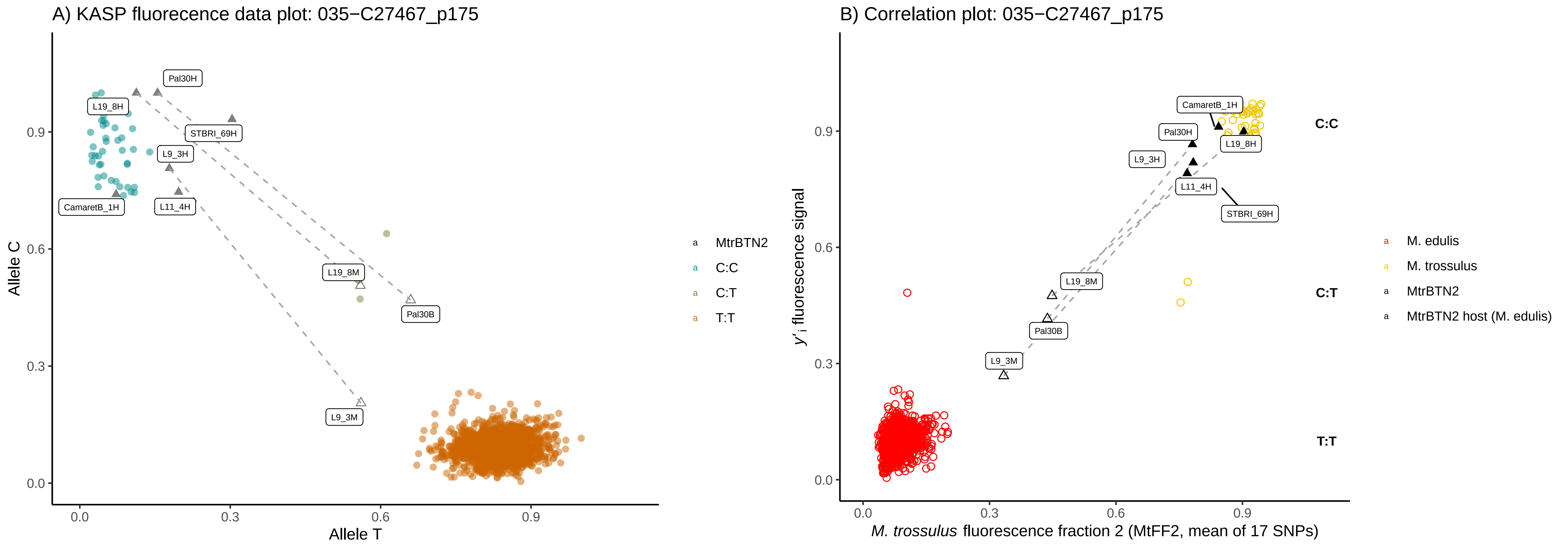

A) KASP fluorecence data plot: 035−C27467_p175
Allele C
Allele T
Pal30H
L19_8H
STBRI_69H
L9_3H
CamaretB_1H
L11_4H
L19_8M
Pal30B
L9_3M
MtrBTN2
C:C
C:T
T:T
B) Correlation plot: 035−C27467_p175
y'i fluorescence signal
M. trossulus fluorescence fraction 2 (MtFF2, mean of 17 SNPs)
CamaretB_1H
Pal30H
L19_8H
L9_3H
L11_4H
STBRI_69H
L19_8M
Pal30B
L9_3M
C:C
C:T
T:T
M. edulis
M. trossulus
MtrBTN2
MtrBTN2 host (M. edulis)

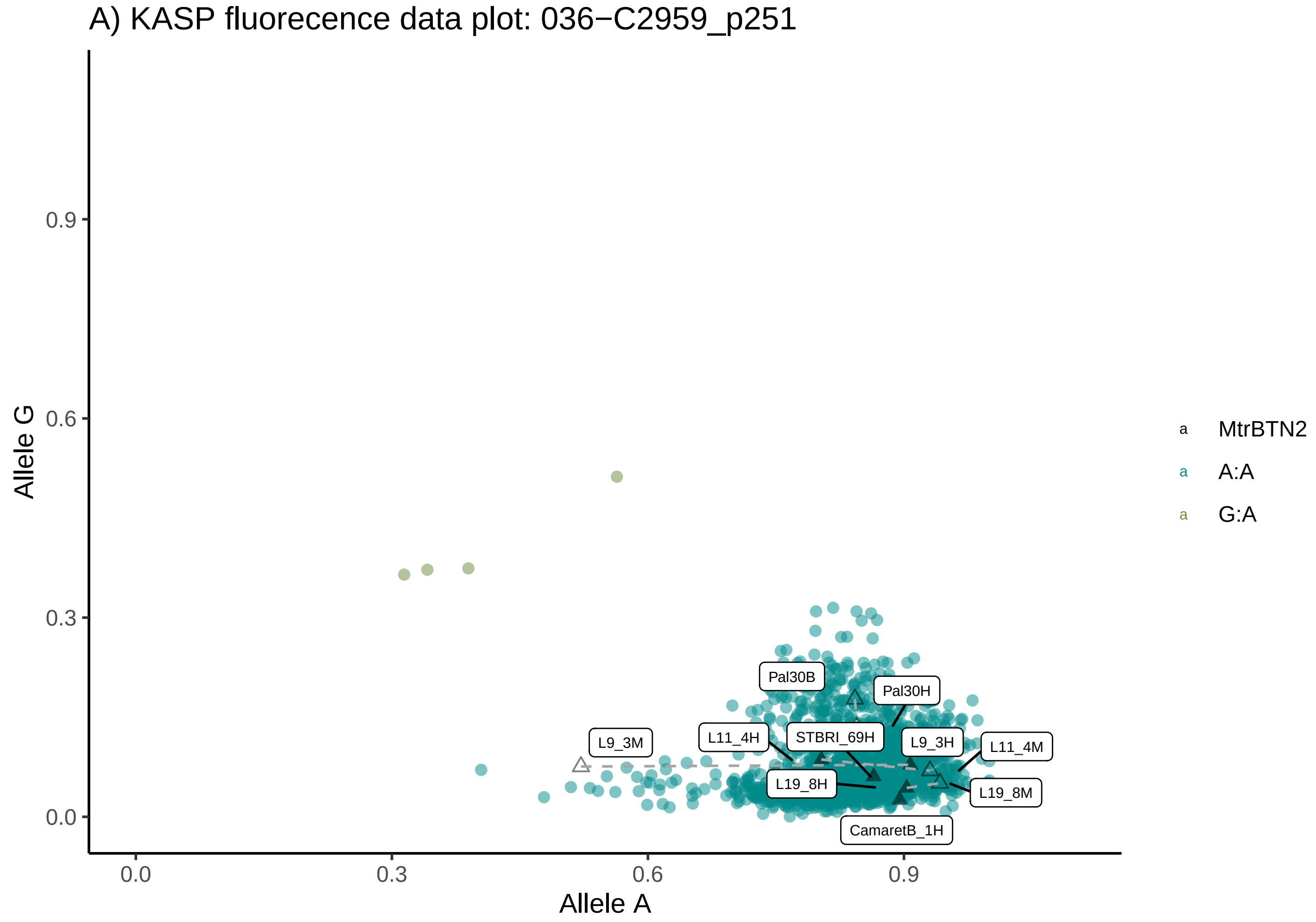
A) KASP fluorecence data plot: 036−C2959_p251
Allele G
Allele A
0.0
0.3
0.6
0.9
Pal30B
Pal30H
L9_3M
L11_4H
STBRI_69H
L9_3H
L11_4M
L19_8H
L19_8M
CamaretB_1H
MtrBTN2
A:A
G:A

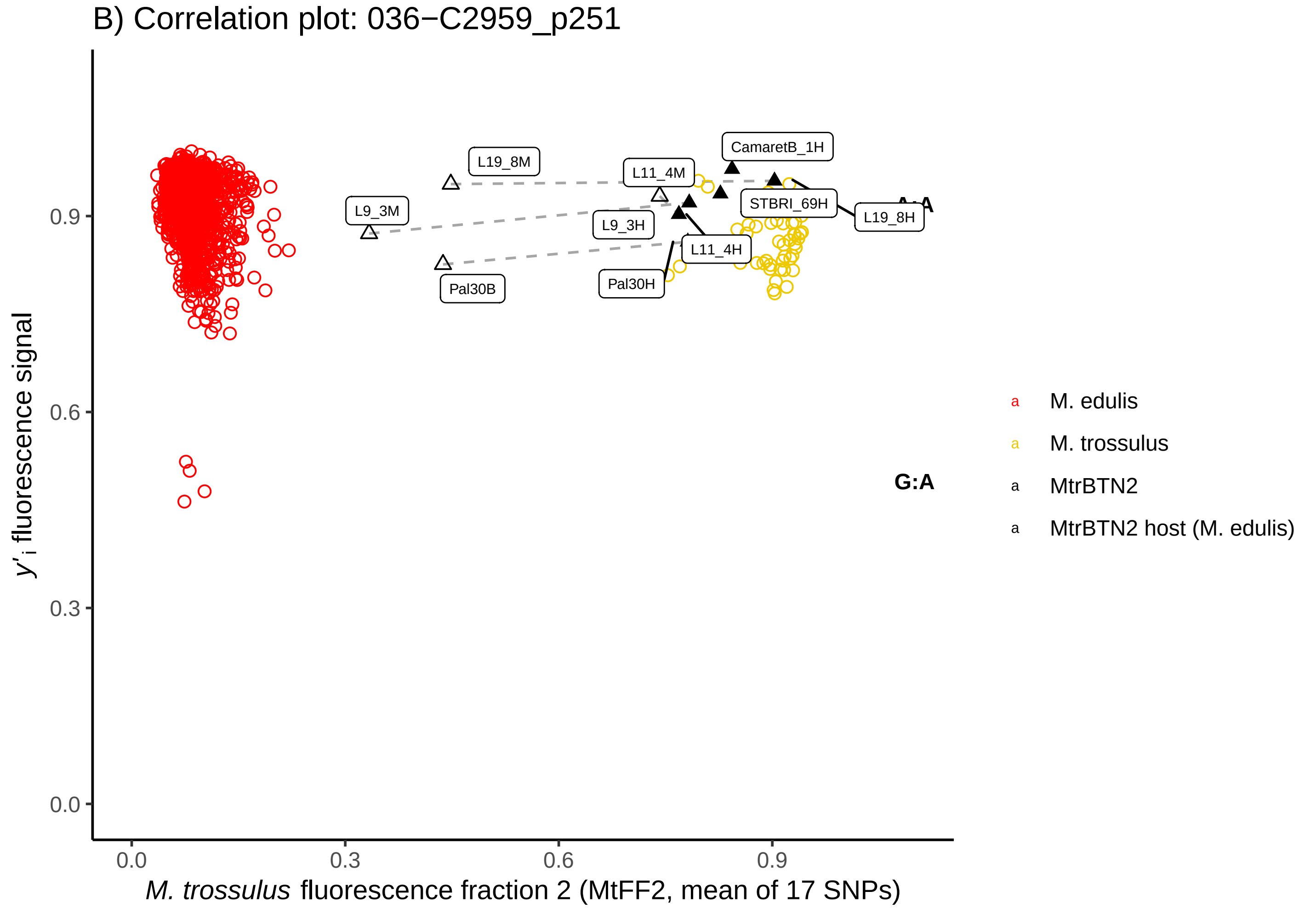
B) Correlation plot: 036−C2959_p251
y'i fluorescence signal
M. trossulus fluorescence fraction 2 (MtFF2, mean of 17 SNPs)
0.0
0.3
0.6
0.9
L19_8M
CamaretB_1H
L11_4M
L9_3M
STBRI_69H
L19_8H
L9_3H
L11_4H
Pal30B
Pal30H
A:A
G:A
M. edulis
M. trossulus
MtrBTN2
MtrBTN2 host (M. edulis)

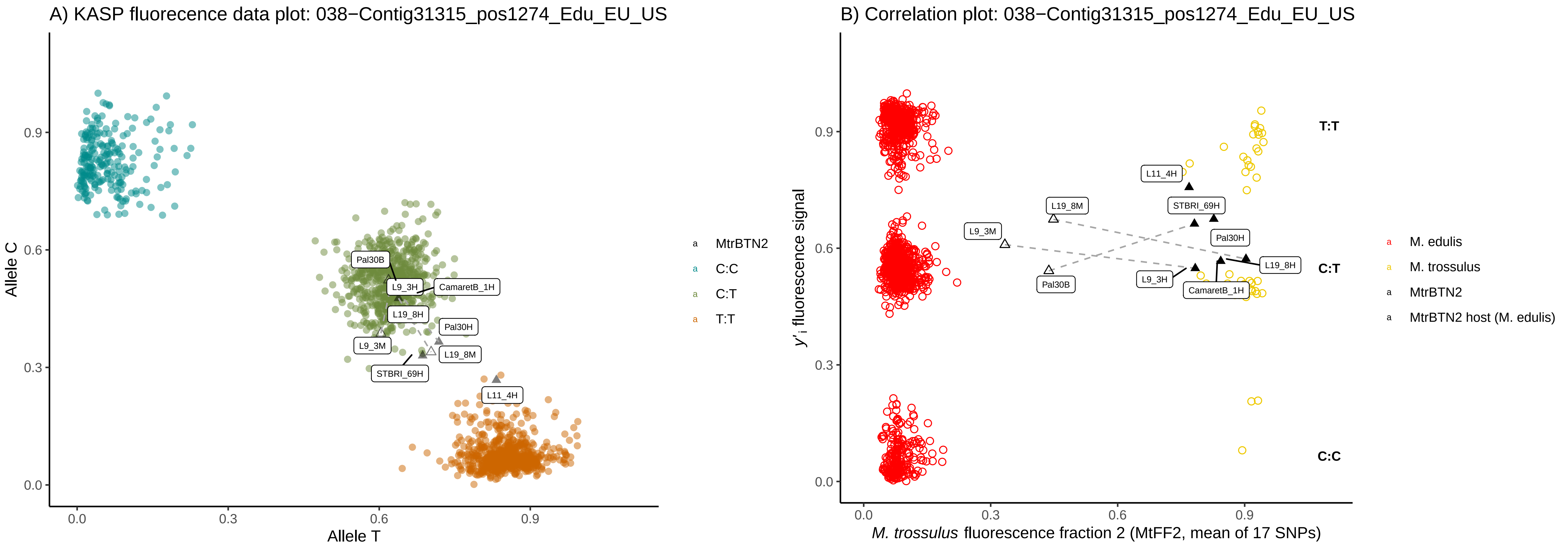
A) KASP fluorecence data plot: 038−Contig31315_pos1274_Edu_EU_US
Allele C
Allele T
0.0
0.3
0.6
0.9
Pal30B
L9_3H
CamaretB_1H
L19_8H
Pal30H
L9_3M
L19_8M
STBRI_69H
L11_4H
MtrBTN2
C:C
C:T
T:T
B) Correlation plot: 038−Contig31315_pos1274_Edu_EU_US
y'i fluorescence signal
M. trossulus fluorescence fraction 2 (MtFF2, mean of 17 SNPs)
T:T
C:T
C:C
L11_4H
L19_8M
STBRI_69H
L9_3M
Pal30H
L19_8H
Pal30B
L9_3H
CamaretB_1H
M. edulis
M. trossulus
MtrBTN2
MtrBTN2 host (M. edulis)

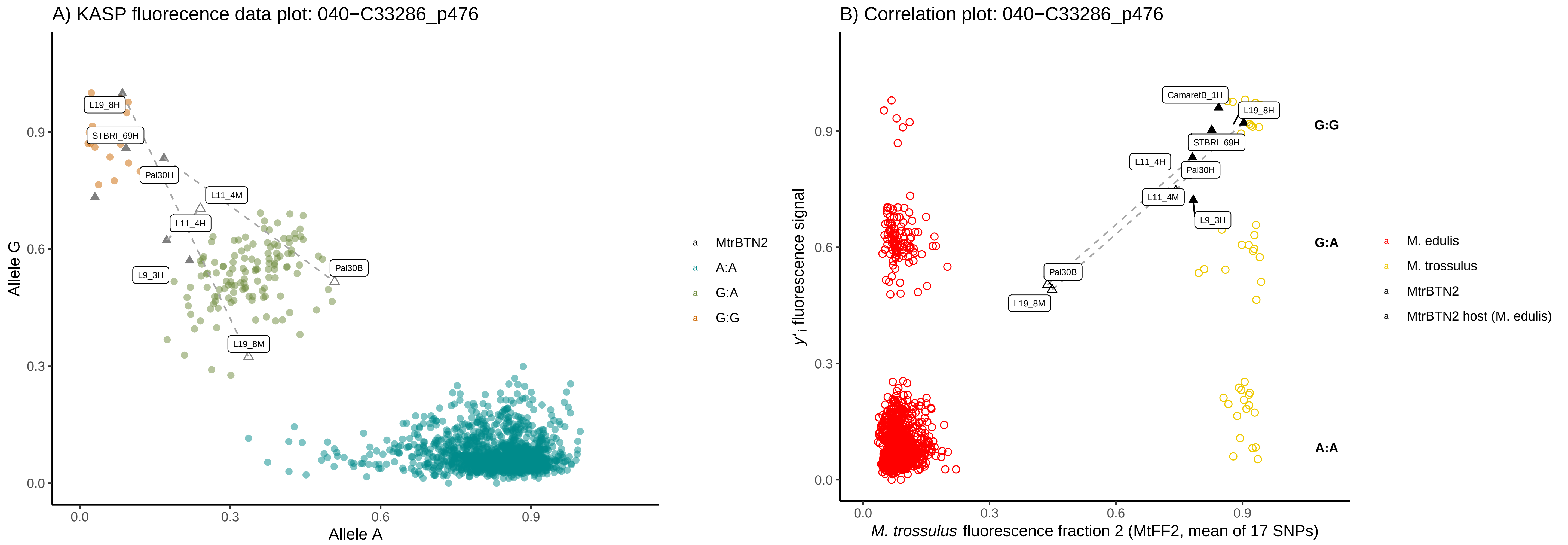
A) KASP fluorecence data plot: 040−C33286_p476
Allele G
Allele A
0.0
0.3
0.6
0.9
L19_8H
STBRI_69H
Pal30H
L11_4M
L11_4H
L9_3H
Pal30B
L19_8M
MtrBTN2
A:A
G:A
G:G
B) Correlation plot: 040−C33286_p476
y'i fluorescence signal
M. trossulus fluorescence fraction 2 (MtFF2, mean of 17 SNPs)
CamaretB_1H
L19_8H
STBRI_69H
L11_4H
Pal30H
L11_4M
L9_3H
Pal30B
L19_8M
G:G
G:A
A:A
M. edulis
M. trossulus
MtrBTN2
MtrBTN2 host (M. edulis)

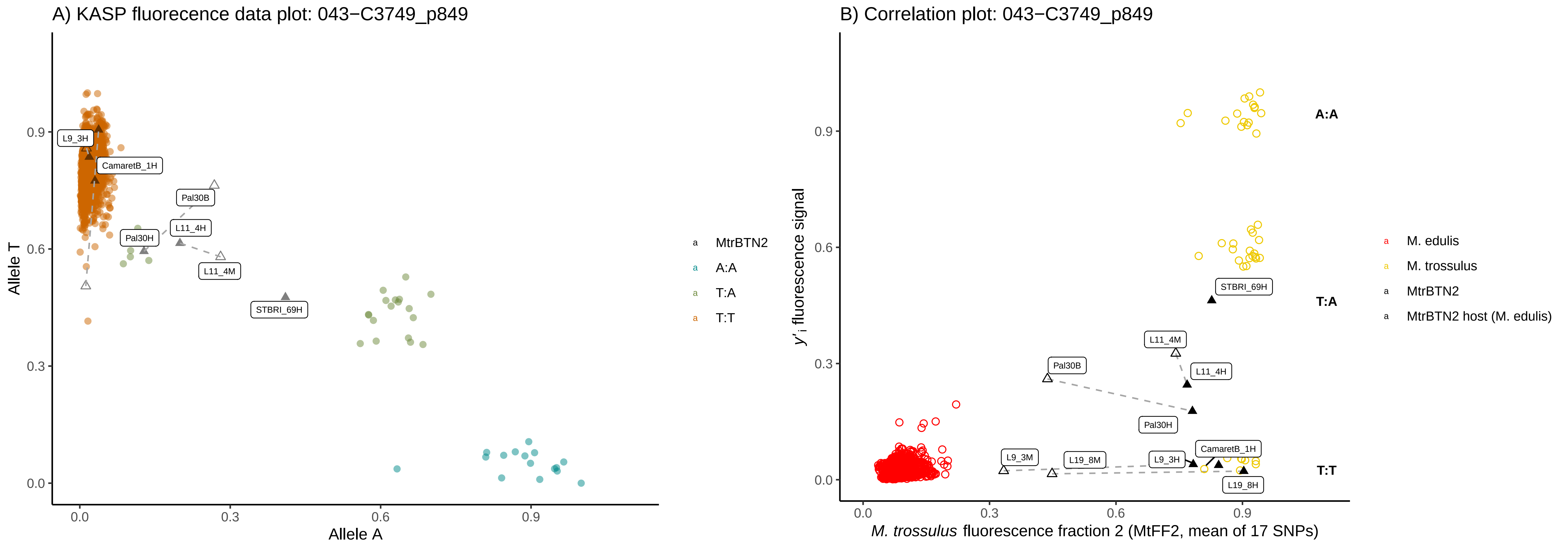
A) KASP fluorecence data plot: 043−C3749_p849
Allele T
Allele A
0.0
0.3
0.6
0.9
L9_3H
CamaretB_1H
Pal30B
L11_4H
Pal30H
L11_4M
STBRI_69H
MtrBTN2
A:A
T:A
T:T
B) Correlation plot: 043−C3749_p849
y'i fluorescence signal
M. trossulus fluorescence fraction 2 (MtFF2, mean of 17 SNPs)
A:A
T:A
T:T
STBRI_69H
L11_4M
L11_4H
Pal30B
Pal30H
L9_3M
L19_8M
L9_3H
CamaretB_1H
L19_8H
M. edulis
M. trossulus
MtrBTN2
MtrBTN2 host (M. edulis)

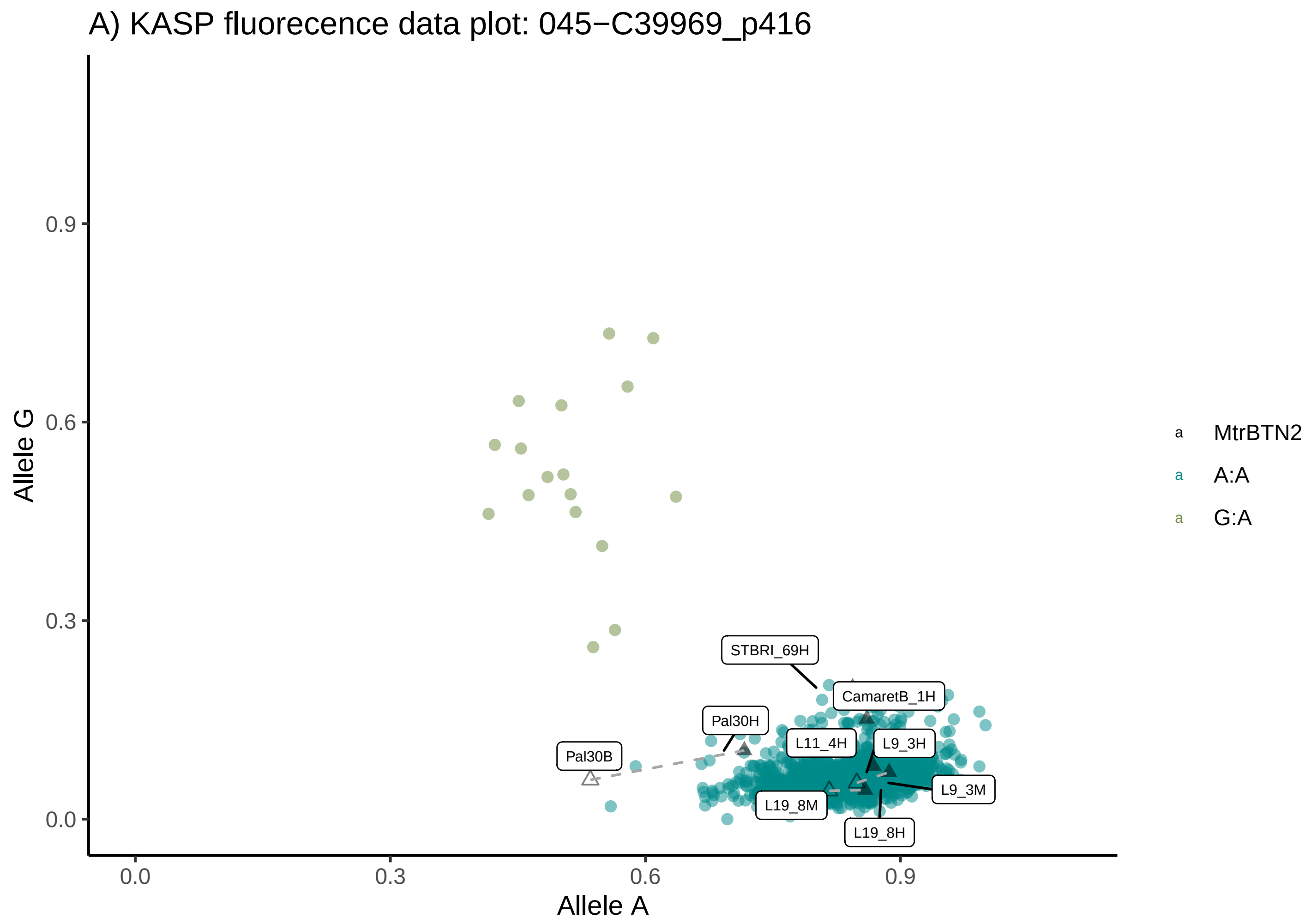
A) KASP fluorecence data plot: 045−C39969_p416
Allele G
Allele A
0.0
0.3
0.6
0.9
STBRI_69H
CamaretB_1H
Pal30H
L11_4H
L9_3H
Pal30B
L9_3M
L19_8M
L19_8H
MtrBTN2
A:A
G:A

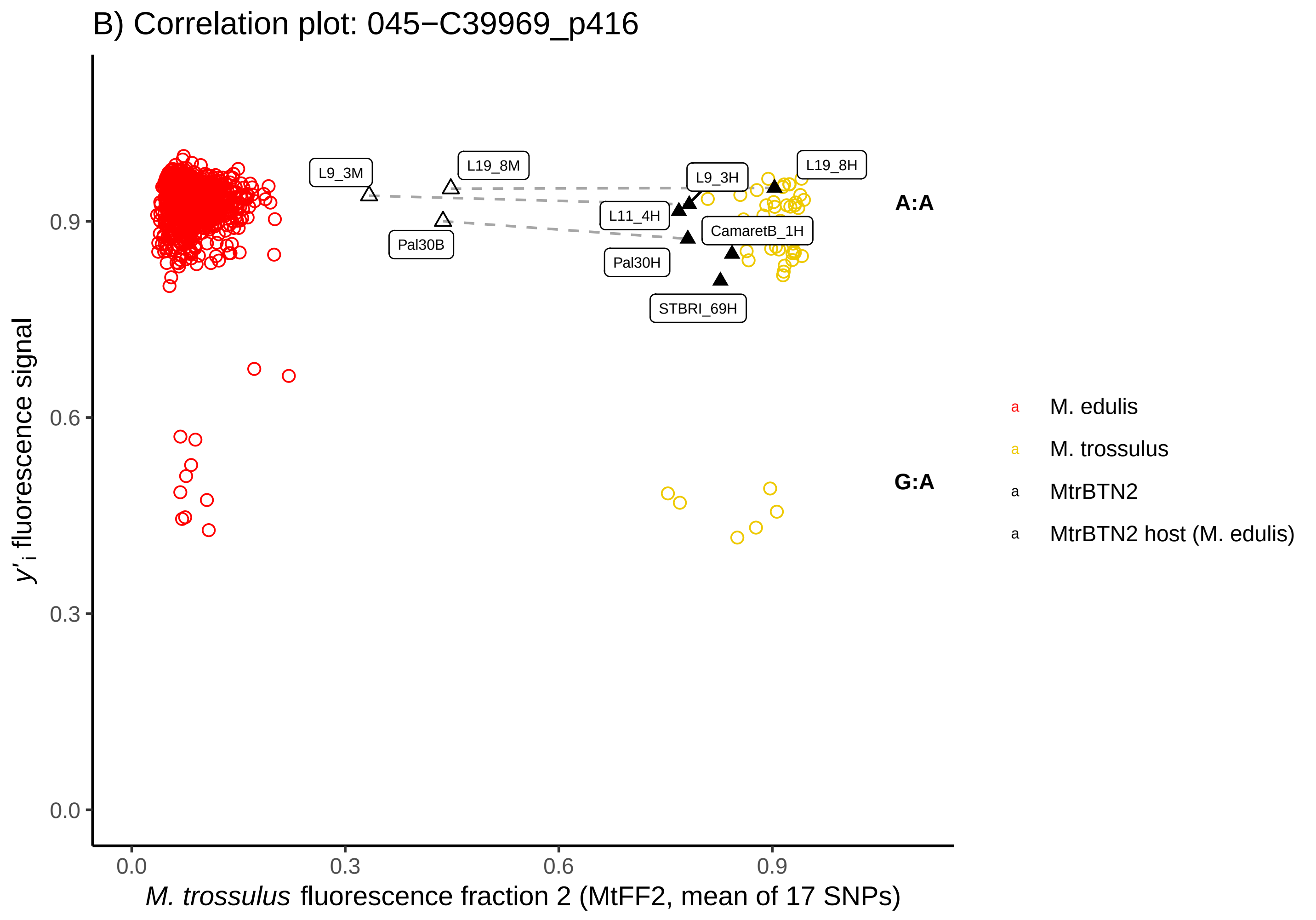
B) Correlation plot: 045−C39969_p416
y'i fluorescence signal
M. trossulus fluorescence fraction 2 (MtFF2, mean of 17 SNPs)
0.0
0.3
0.6
0.9
L9_3M
L19_8M
L9_3H
L19_8H
L11_4H
CamaretB_1H
Pal30B
Pal30H
STBRI_69H
A:A
G:A
M. edulis
M. trossulus
MtrBTN2
MtrBTN2 host (M. edulis)

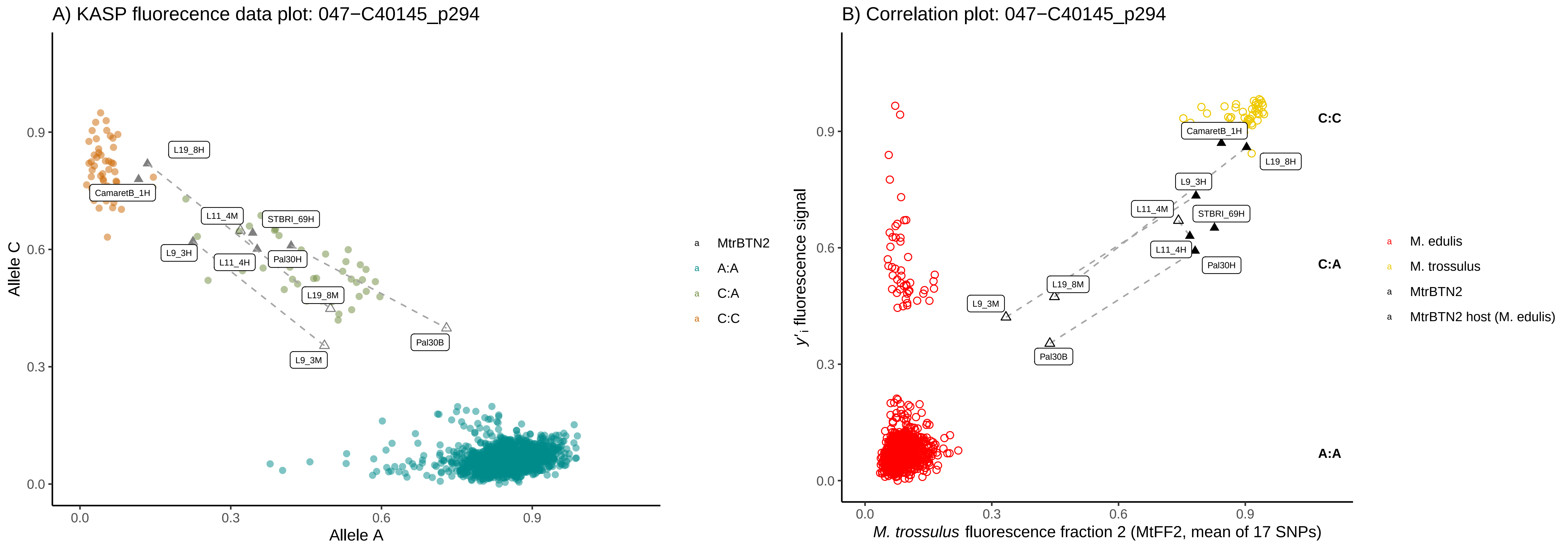
A) KASP fluorecence data plot: 047−C40145_p294
Allele C
Allele A
0.0
0.3
0.6
0.9
L19_8H
CamaretB_1H
L11_4M
STBRI_69H
L9_3H
L11_4H
Pal30H
L19_8M
Pal30B
L9_3M
MtrBTN2
A:A
C:A
C:C
B) Correlation plot: 047−C40145_p294
y'i fluorescence signal
M. trossulus fluorescence fraction 2 (MtFF2, mean of 17 SNPs)
C:C
C:A
A:A
M. edulis
M. trossulus
MtrBTN2
MtrBTN2 host (M. edulis)

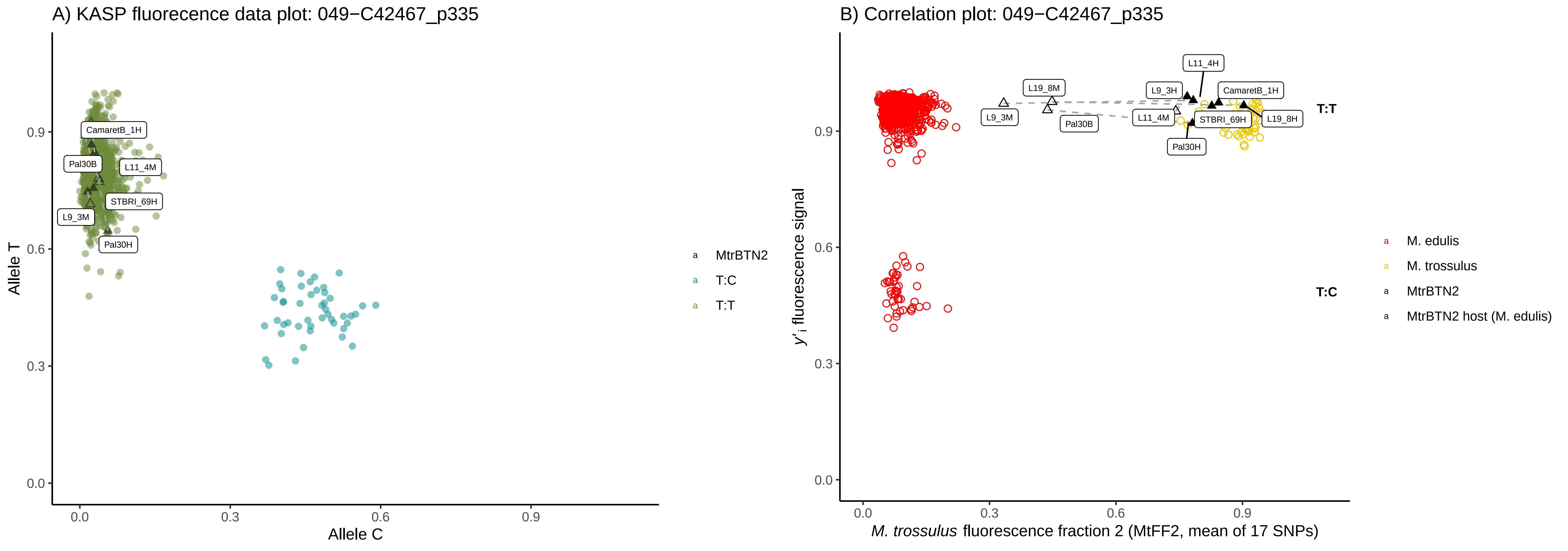
A) KASP fluorecence data plot: 049−C42467_p335
Allele T
Allele C
0.0
0.3
0.6
0.9
CamaretB_1H
Pal30B
L11_4M
STBRI_69H
L9_3M
Pal30H
MtrBTN2
T:C
T:T
B) Correlation plot: 049−C42467_p335
y'i fluorescence signal
M. trossulus fluorescence fraction 2 (MtFF2, mean of 17 SNPs)
L11_4H
L19_8M
L9_3H
CamaretB_1H
L9_3M
Pal30B
L11_4M
STBRI_69H
L19_8H
Pal30H
T:T
T:C
M. edulis
M. trossulus
MtrBTN2
MtrBTN2 host (M. edulis)

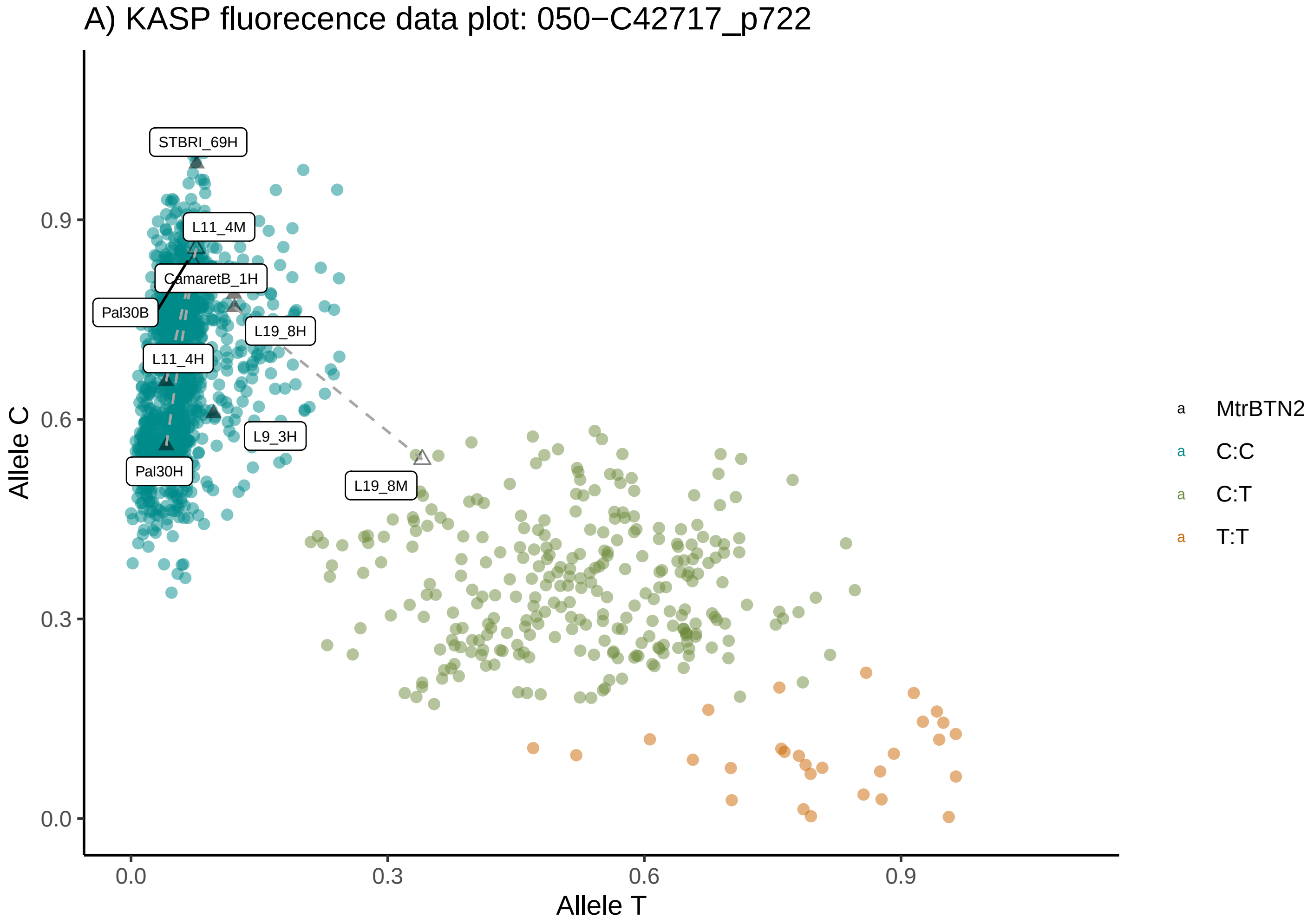
A) KASP fluorecence data plot: 050−C42717_p722
Allele C
Allele T
STBRI_69H
L11_4M
CamaretB_1H
Pal30B
L19_8H
L11_4H
L9_3H
Pal30H
L19_8M
MtrBTN2
C:C
C:T
T:T

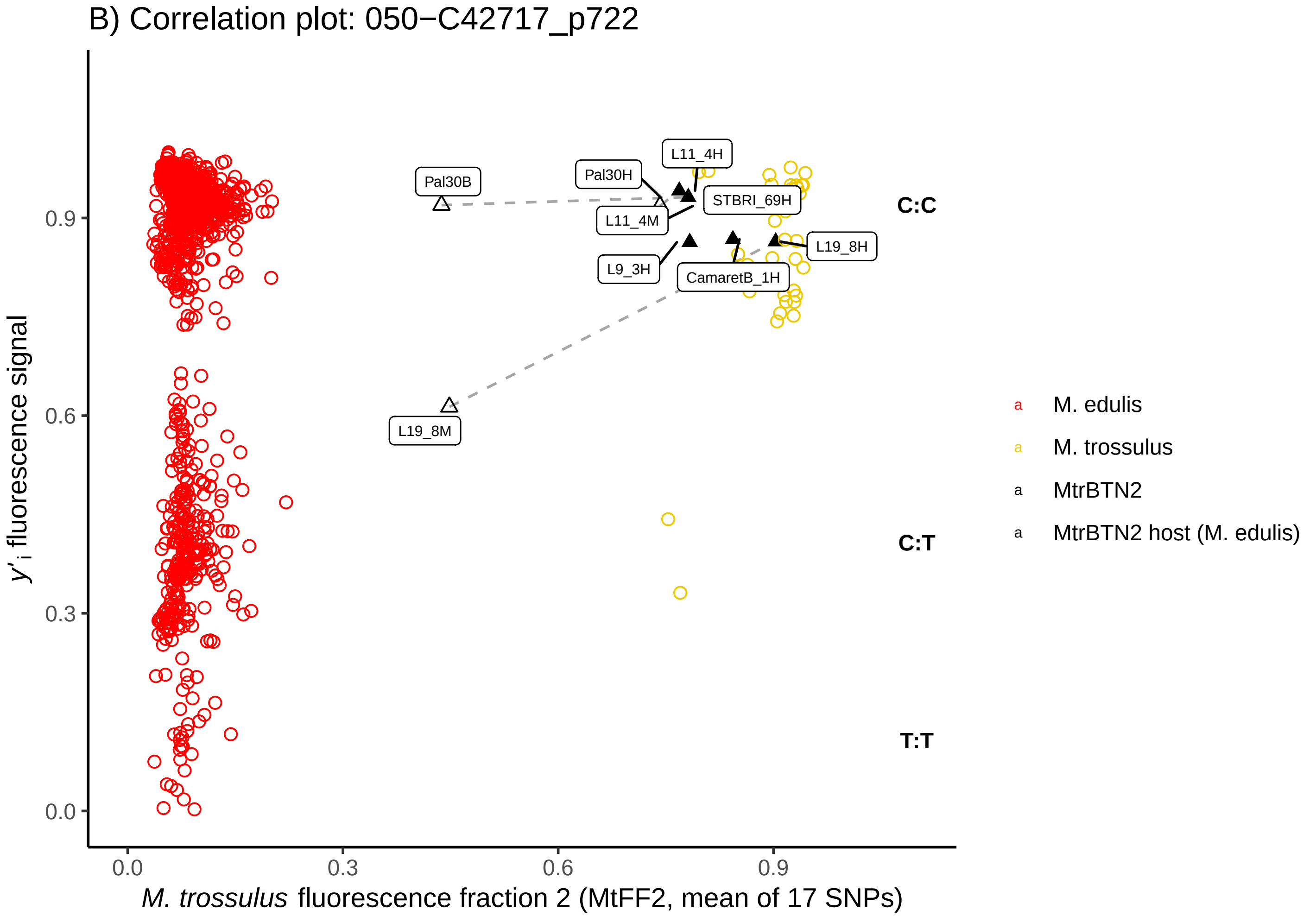
B) Correlation plot: 050−C42717_p722
y'i fluorescence signal
M. trossulus fluorescence fraction 2 (MtFF2, mean of 17 SNPs)
Pal30B
Pal30H
L11_4H
STBRI_69H
L11_4M
L9_3H
CamaretB_1H
L19_8H
L19_8M
C:C
C:T
T:T
M. edulis
M. trossulus
MtrBTN2
MtrBTN2 host (M. edulis)

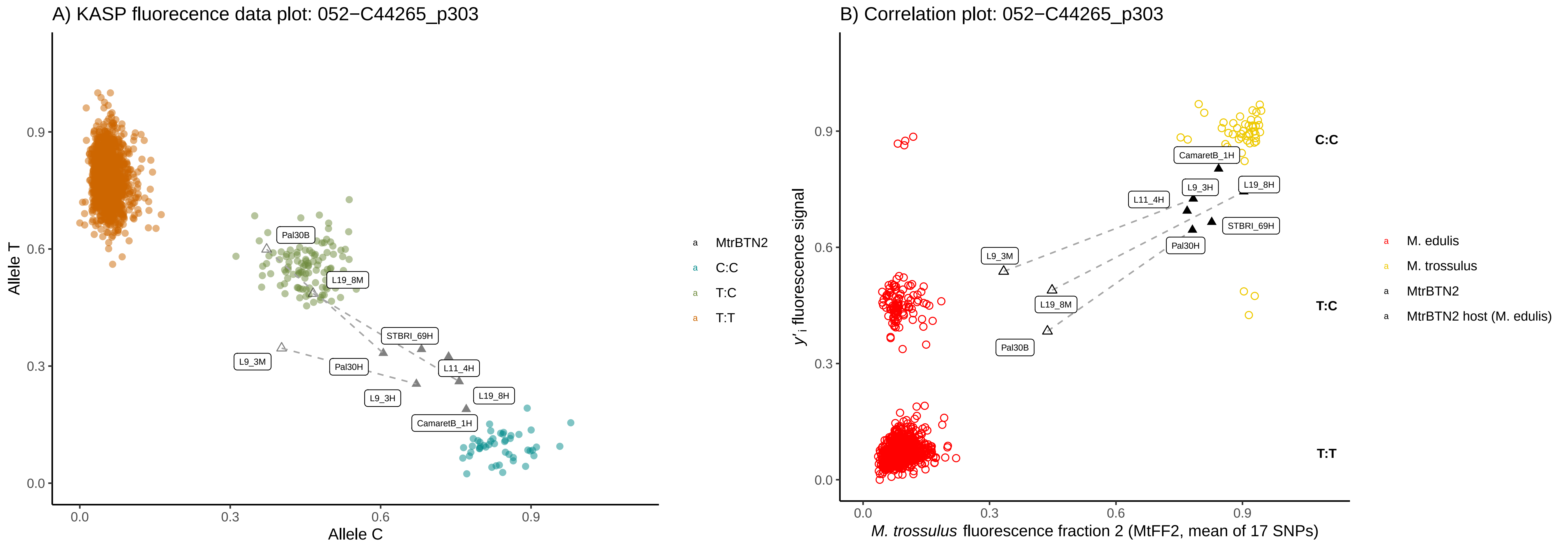

A) KASP fluorecence data plot: 052−C44265_p303
Allele T
Allele C
0.0
0.3
0.6
0.9
Pal30B
L19_8M
STBRI_69H
L9_3M
Pal30H
L11_4H
L9_3H
L19_8H
CamaretB_1H
MtrBTN2
C:C
T:C
T:T
B) Correlation plot: 052−C44265_p303
y'i fluorescence signal
M. trossulus fluorescence fraction 2 (MtFF2, mean of 17 SNPs)
CamaretB_1H
L9_3H
L19_8H
L11_4H
STBRI_69H
Pal30H
L9_3M
L19_8M
Pal30B
C:C
T:C
T:T
M. edulis
M. trossulus
MtrBTN2
MtrBTN2 host (M. edulis)

A) KASP fluorecence data plot: 055−C4715_p1037

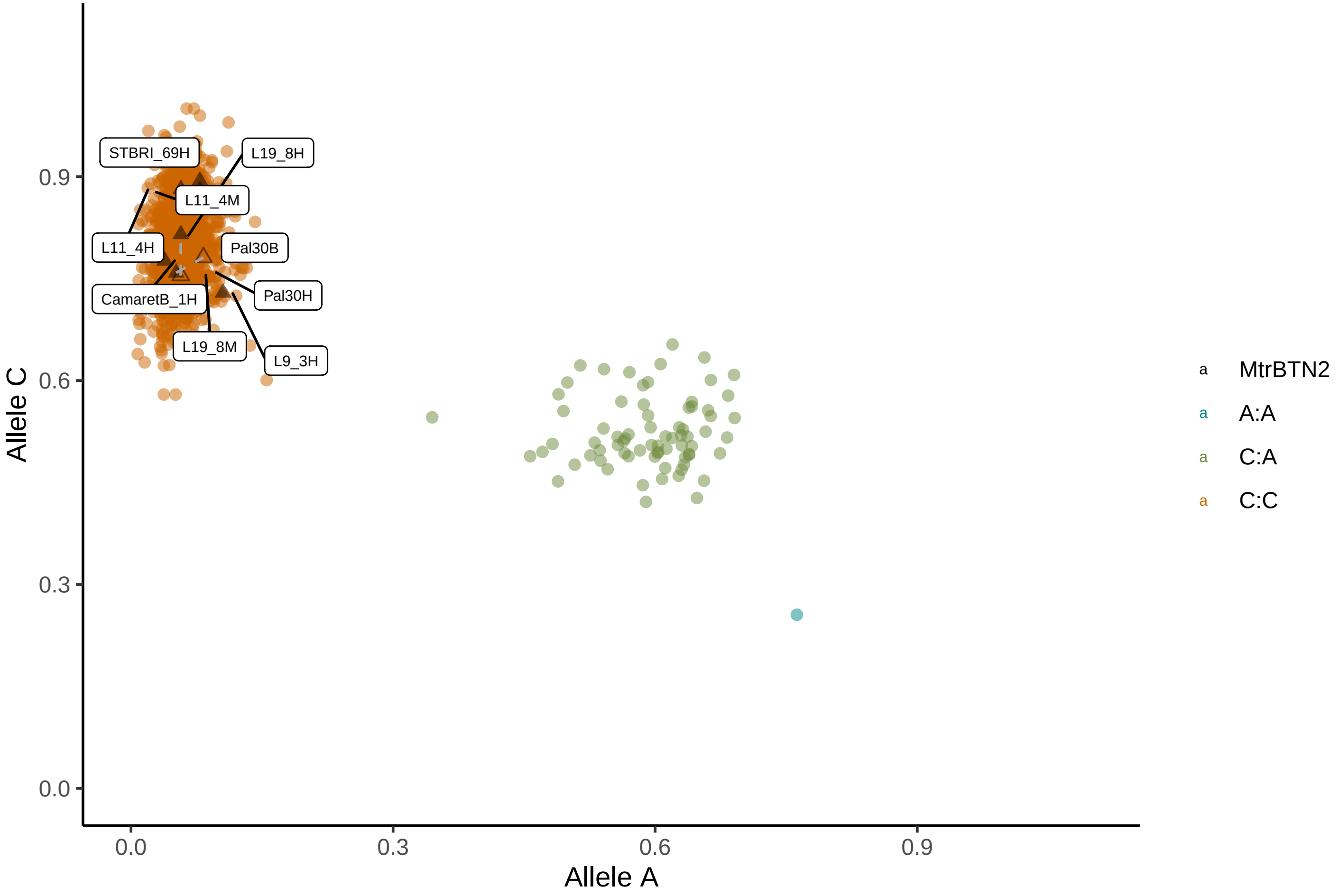


B) Correlation plot: 055−C4715_p1037

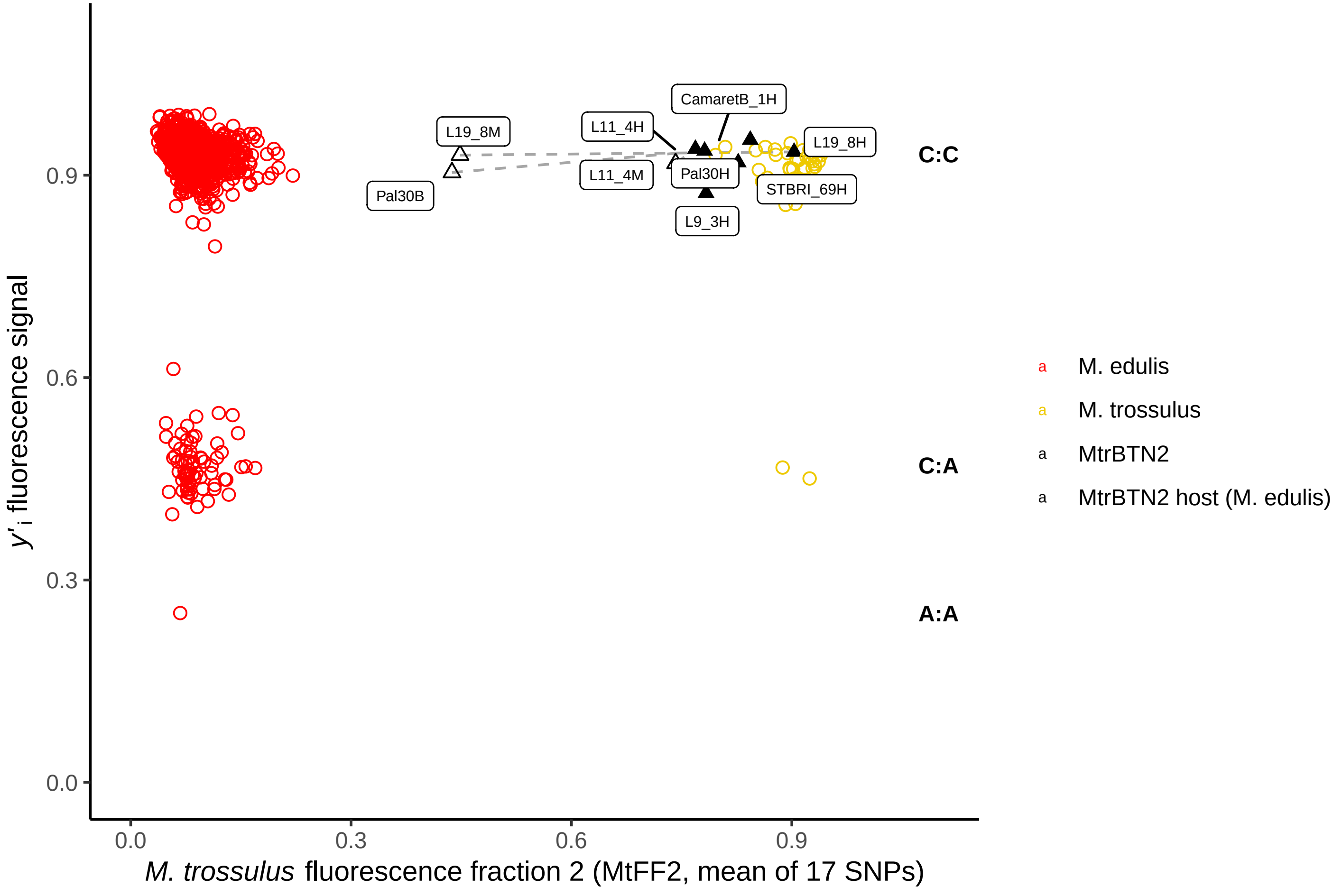

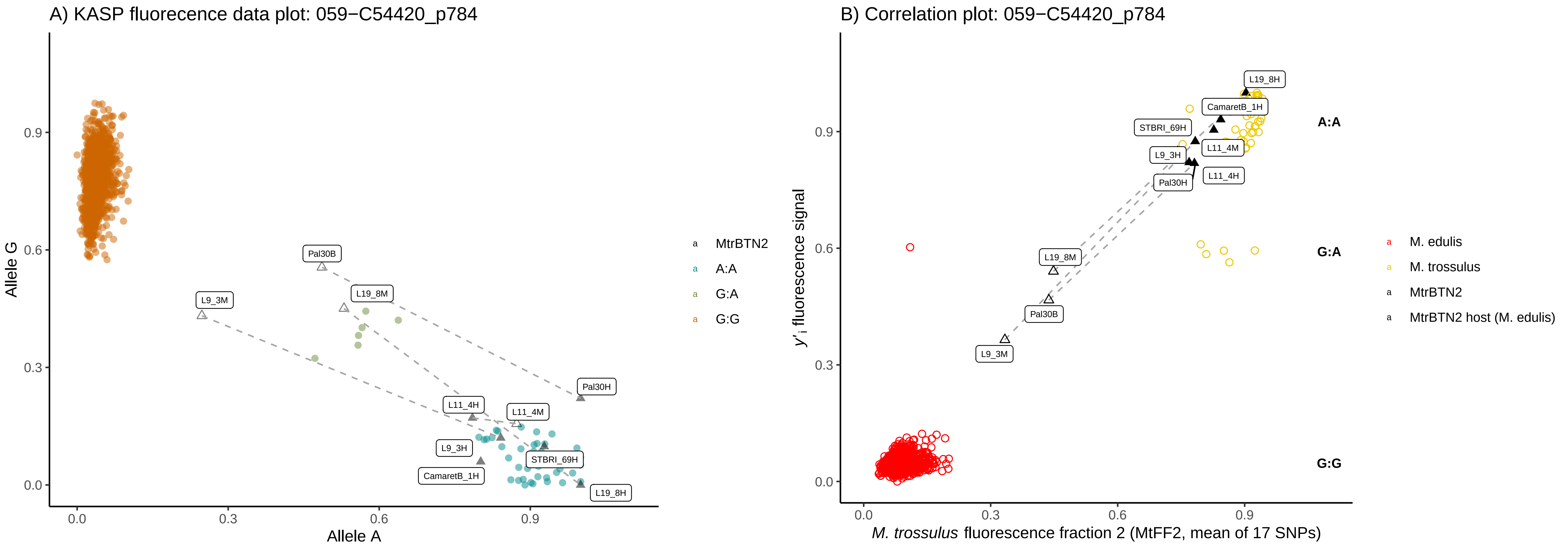
A) KASP fluorecence data plot: 059−C54420_p784
Allele G
Allele A
0.0
0.3
0.6
0.9
Pal30B
L9_3M
L19_8M
Pal30H
L11_4H
L11_4M
L9_3H
STBRI_69H
CamaretB_1H
L19_8H
MtrBTN2
A:A
G:A
G:G
B) Correlation plot: 059−C54420_p784
y'i fluorescence signal
M. trossulus fluorescence fraction 2 (MtFF2, mean of 17 SNPs)
L19_8H
CamaretB_1H
STBRI_69H
L11_4M
L9_3H
L11_4H
Pal30H
L19_8M
Pal30B
L9_3M
A:A
G:A
G:G
M. edulis
M. trossulus
MtrBTN2
MtrBTN2 host (M. edulis)

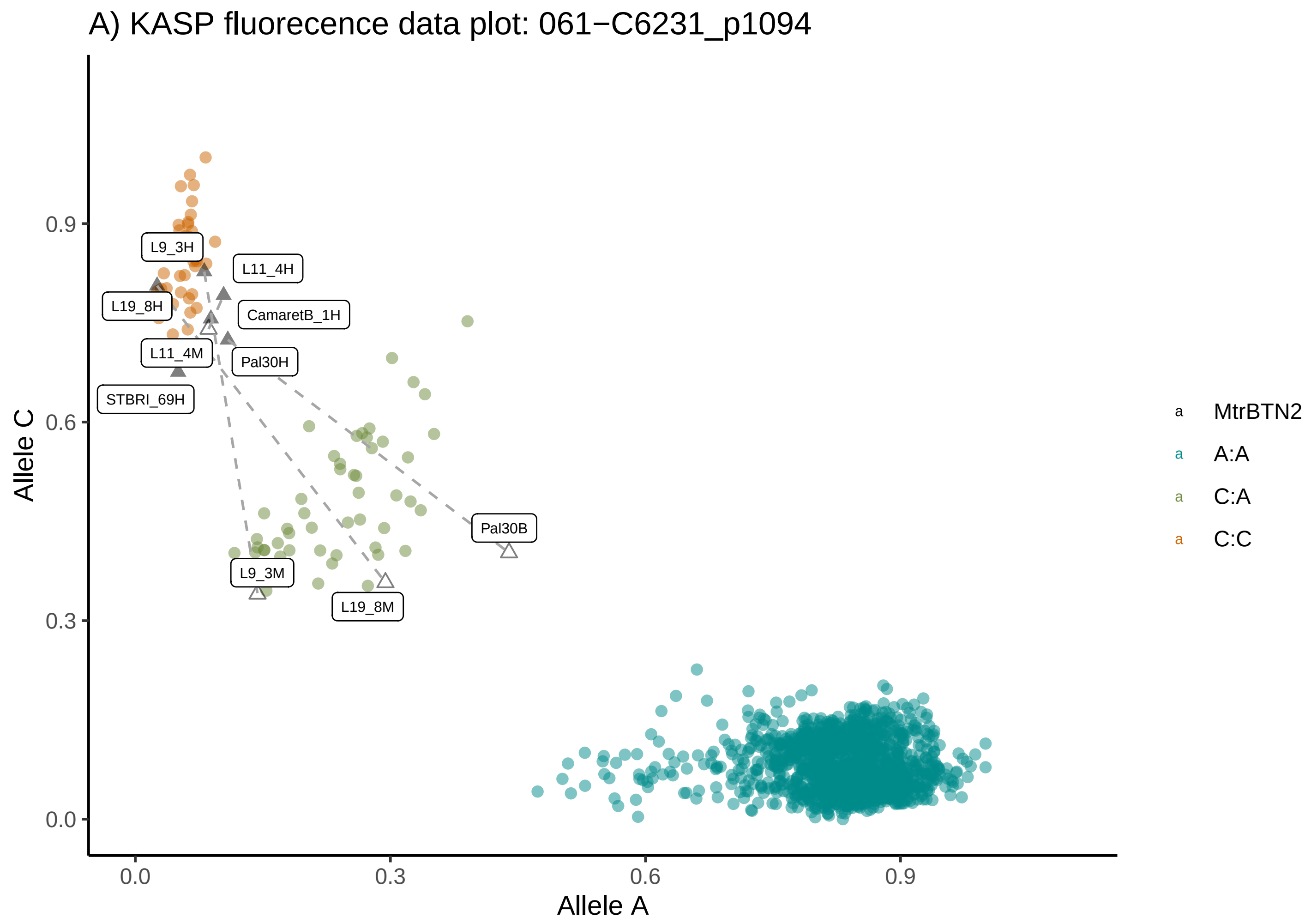

A) KASP fluorecence data plot: 061−C6231_p1094
Allele C
Allele A
0.0
0.3
0.6
0.9
L9_3H
L11_4H
L19_8H
CamaretB_1H
L11_4M
Pal30H
STBRI_69H
Pal30B
L9_3M
L19_8M
MtrBTN2
A:A
C:A
C:C


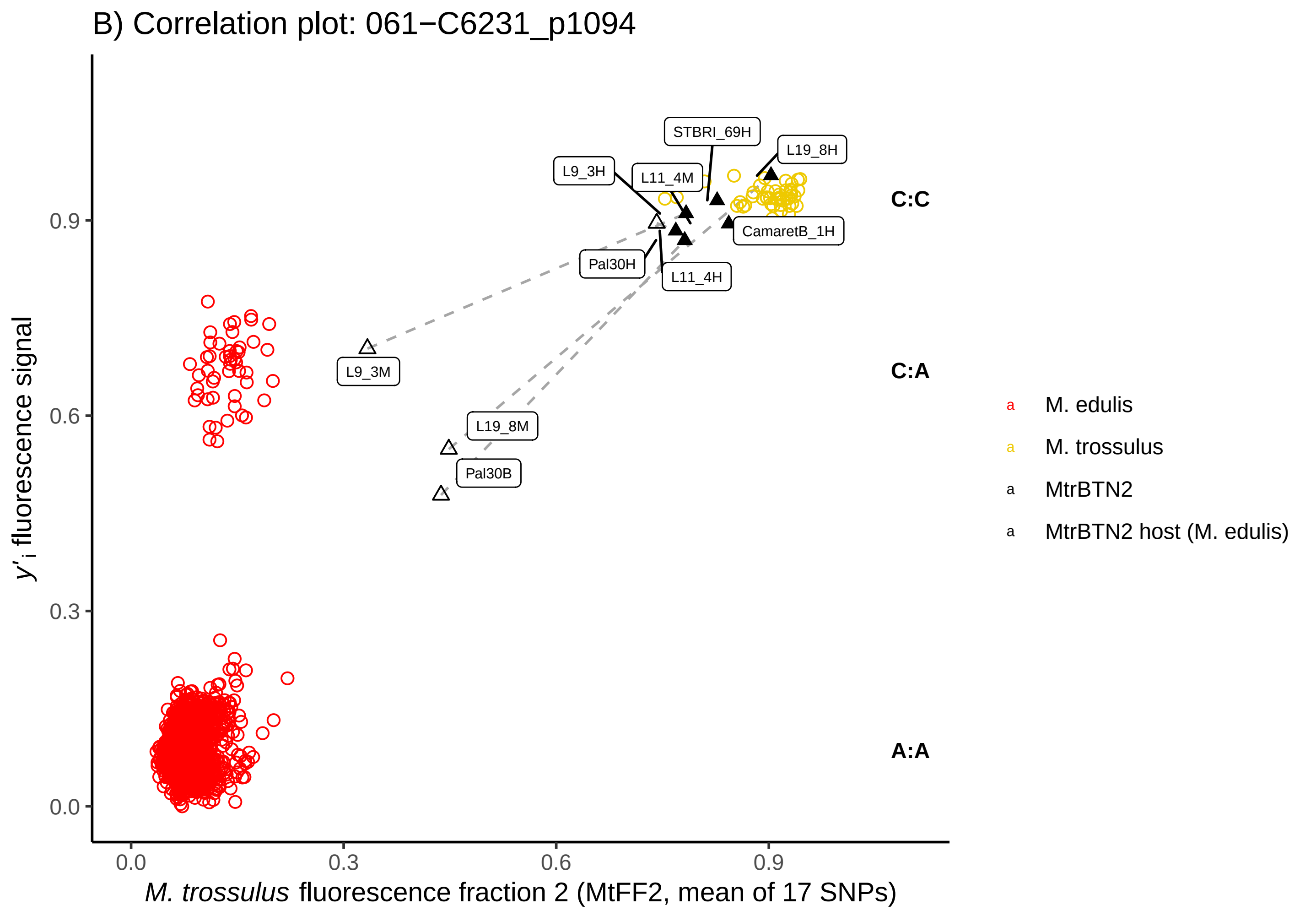

B) Correlation plot: 061−C6231_p1094
y'i fluorescence signal
M. trossulus fluorescence fraction 2 (MtFF2, mean of 17 SNPs)
0.0
0.3
0.6
0.9
STBRI_69H
L19_8H
L9_3H
L11_4M
CamaretB_1H
Pal30H
L11_4H
L9_3M
L19_8M
Pal30B
C:C
C:A
A:A
M. edulis
M. trossulus
MtrBTN2
MtrBTN2 host (M. edulis)

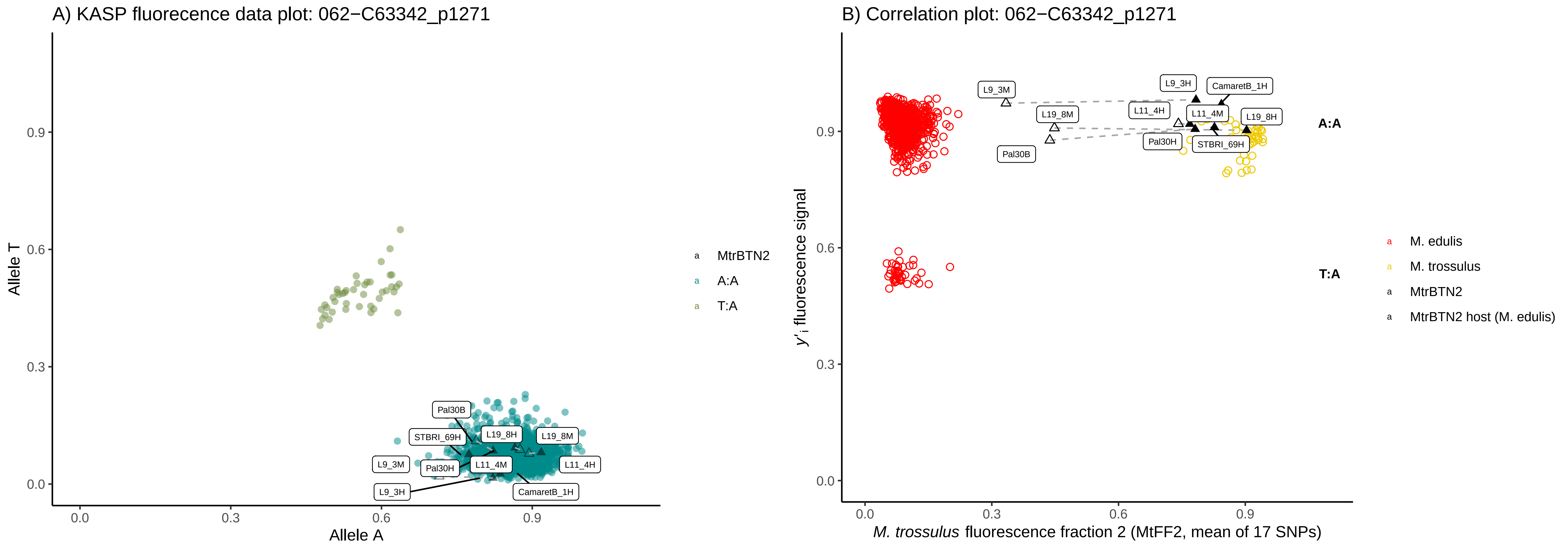
A) KASP fluorecence data plot: 062−C63342_p1271
Allele T
Allele A
0.0
0.3
0.6
0.9
Pal30B
STBRI_69H
L19_8H
L19_8M
L9_3M
Pal30H
L11_4M
L11_4H
L9_3H
CamaretB_1H
MtrBTN2
A:A
T:A
B) Correlation plot: 062−C63342_p1271
y'i fluorescence signal
M. trossulus fluorescence fraction 2 (MtFF2, mean of 17 SNPs)
L9_3M
L19_8M
Pal30B
L9_3H
CamaretB_1H
L11_4H
L11_4M
L19_8H
Pal30H
STBRI_69H
A:A
T:A
M. edulis
M. trossulus
MtrBTN2
MtrBTN2 host (M. edulis)

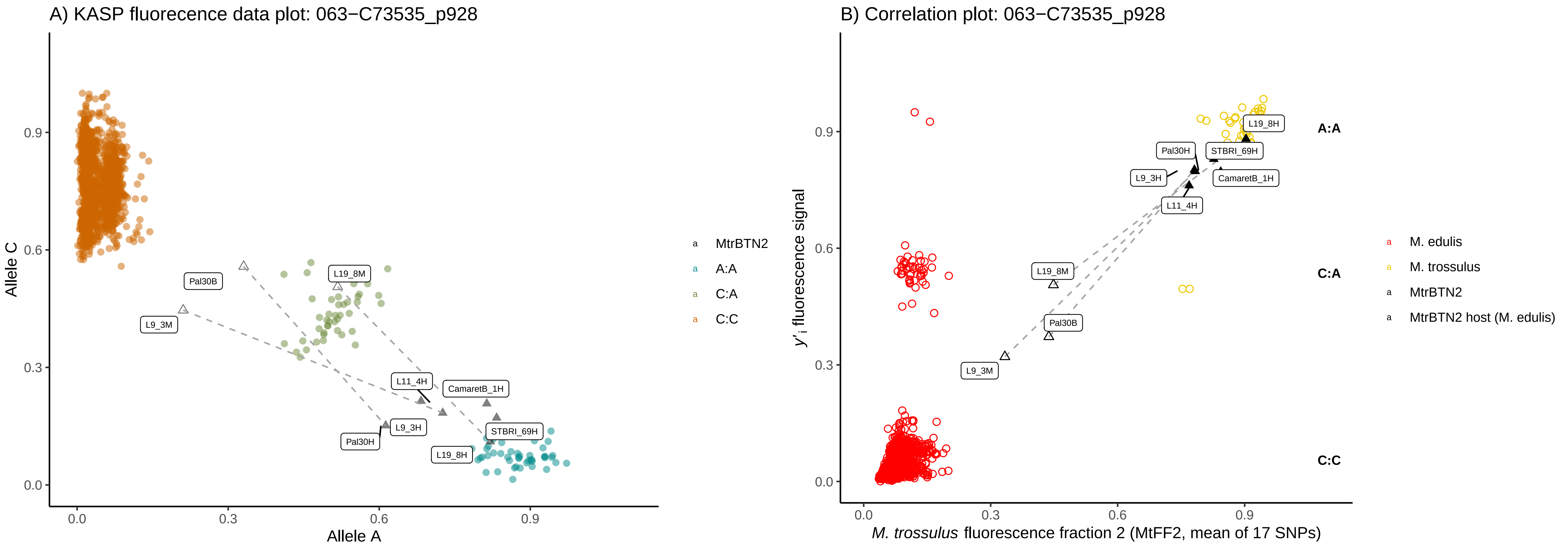

A) KASP fluorecence data plot: 063−C73535_p928
Allele C
Allele A
0.0
0.3
0.6
0.9
Pal30B
L9_3M
L19_8M
L11_4H
CamaretB_1H
L9_3H
Pal30H
STBRI_69H
L19_8H
MtrBTN2
A:A
C:A
C:C
B) Correlation plot: 063−C73535_p928
y'i fluorescence signal
M. trossulus fluorescence fraction 2 (MtFF2, mean of 17 SNPs)
A:A
C:A
C:C
Pal30H
STBRI_69H
L9_3H
CamaretB_1H
L11_4H
L19_8H
L19_8M
Pal30B
L9_3M
M. edulis
M. trossulus
MtrBTN2
MtrBTN2 host (M. edulis)

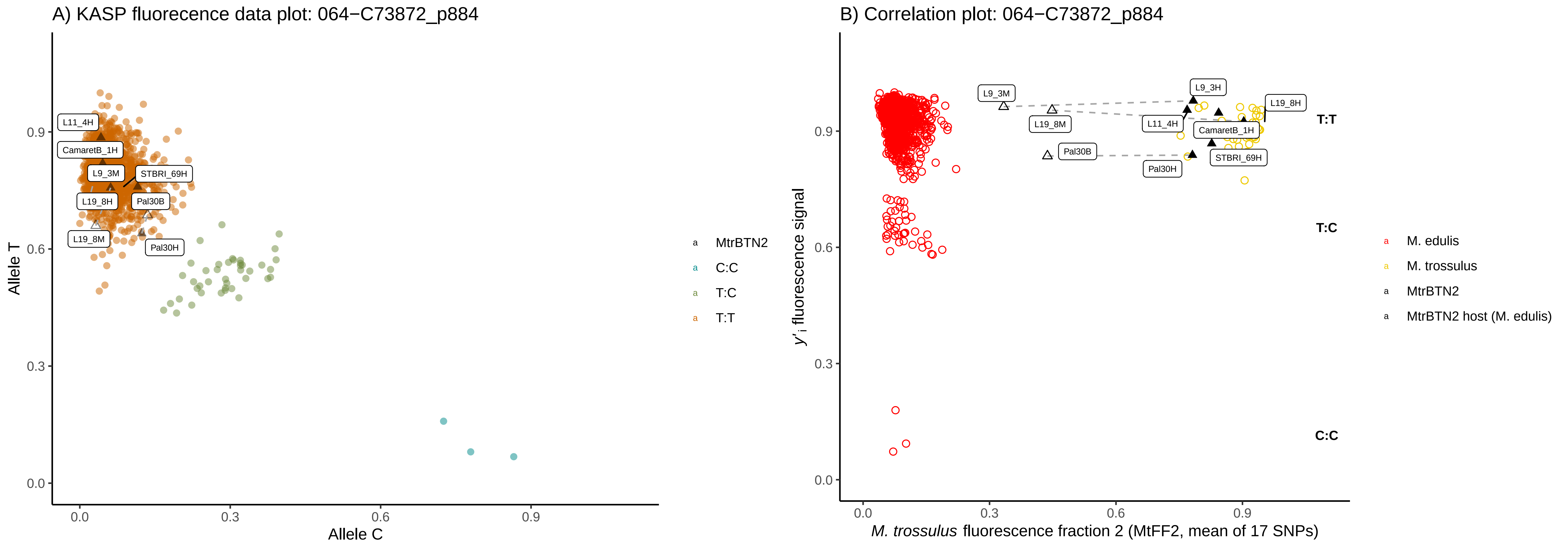
A) KASP fluorecence data plot: 064−C73872_p884
Allele T
Allele C
0.0
0.3
0.6
0.9
L11_4H
CamaretB_1H
L9_3M
STBRI_69H
L19_8H
Pal30B
L19_8M
Pal30H
MtrBTN2
C:C
T:C
T:T
B) Correlation plot: 064−C73872_p884
y'i fluorescence signal
M. trossulus fluorescence fraction 2 (MtFF2, mean of 17 SNPs)
L9_3M
L9_3H
L19_8H
L19_8M
L11_4H
CamaretB_1H
Pal30B
STBRI_69H
Pal30H
T:T
T:C
C:C
M. edulis
M. trossulus
MtrBTN2
MtrBTN2 host (M. edulis)

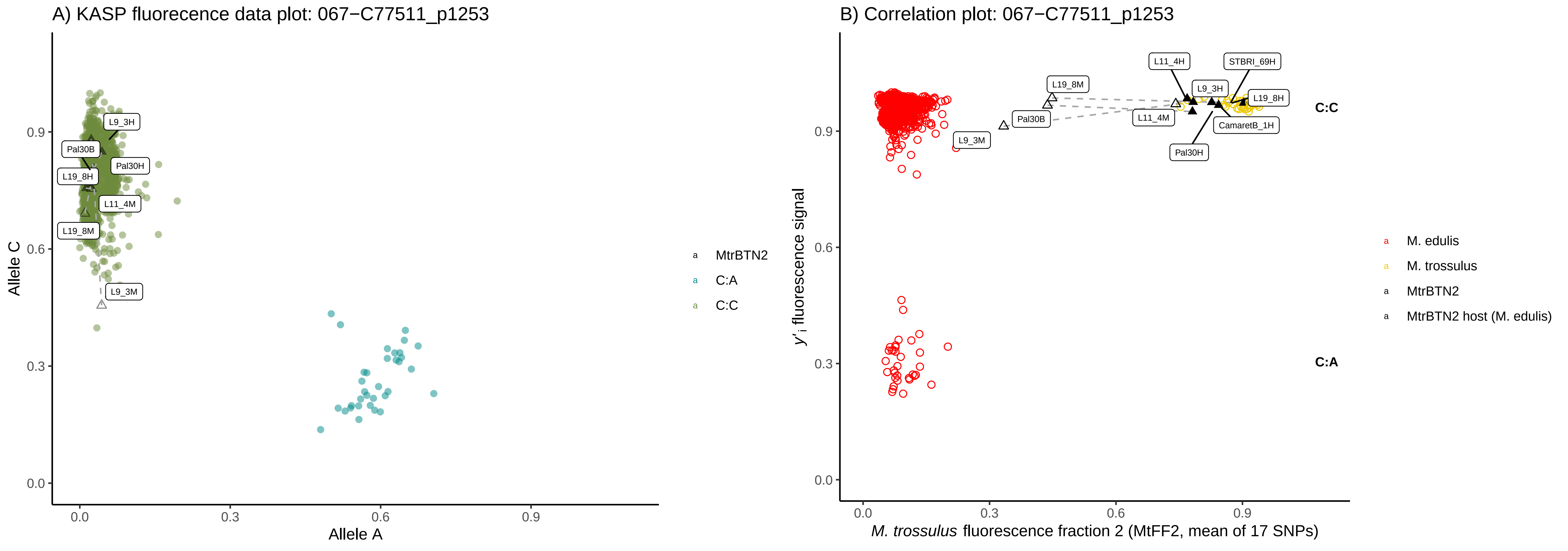
A) KASP fluorecence data plot: 067−C77511_p1253
Allele C
Allele A
0.0
0.3
0.6
0.9
L9_3H
Pal30B
Pal30H
L19_8H
L11_4M
L19_8M
L9_3M
MtrBTN2
C:A
C:C
B) Correlation plot: 067−C77511_p1253
y'i fluorescence signal
M. trossulus fluorescence fraction 2 (MtFF2, mean of 17 SNPs)
L11_4H
STBRI_69H
L19_8M
L9_3H
L19_8H
Pal30B
L11_4M
CamaretB_1H
L9_3M
Pal30H
C:C
C:A
M. edulis
M. trossulus
MtrBTN2
MtrBTN2 host (M. edulis)

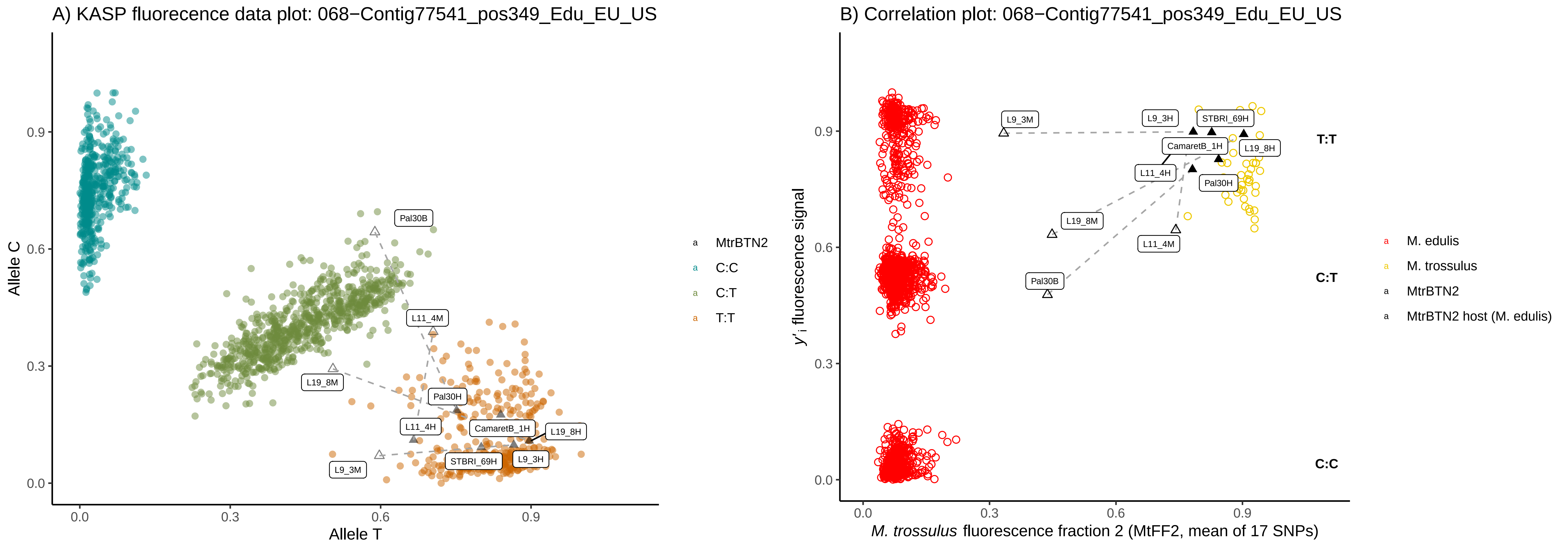
A) KASP fluorecence data plot: 068−Contig77541_pos349_Edu_EU_US
Allele C
Allele T
0.0
0.3
0.6
0.9
Pal30B
L11_4M
L19_8M
Pal30H
L11_4H
CamaretB_1H
L19_8H
STBRI_69H
L9_3H
L9_3M
MtrBTN2
C:C
C:T
T:T
B) Correlation plot: 068−Contig77541_pos349_Edu_EU_US
y'i fluorescence signal
M. trossulus fluorescence fraction 2 (MtFF2, mean of 17 SNPs)
L9_3M
L9_3H
STBRI_69H
CamaretB_1H
L19_8H
L11_4H
Pal30H
L19_8M
L11_4M
Pal30B
T:T
C:T
C:C
M. edulis
M. trossulus
MtrBTN2
MtrBTN2 host (M. edulis)

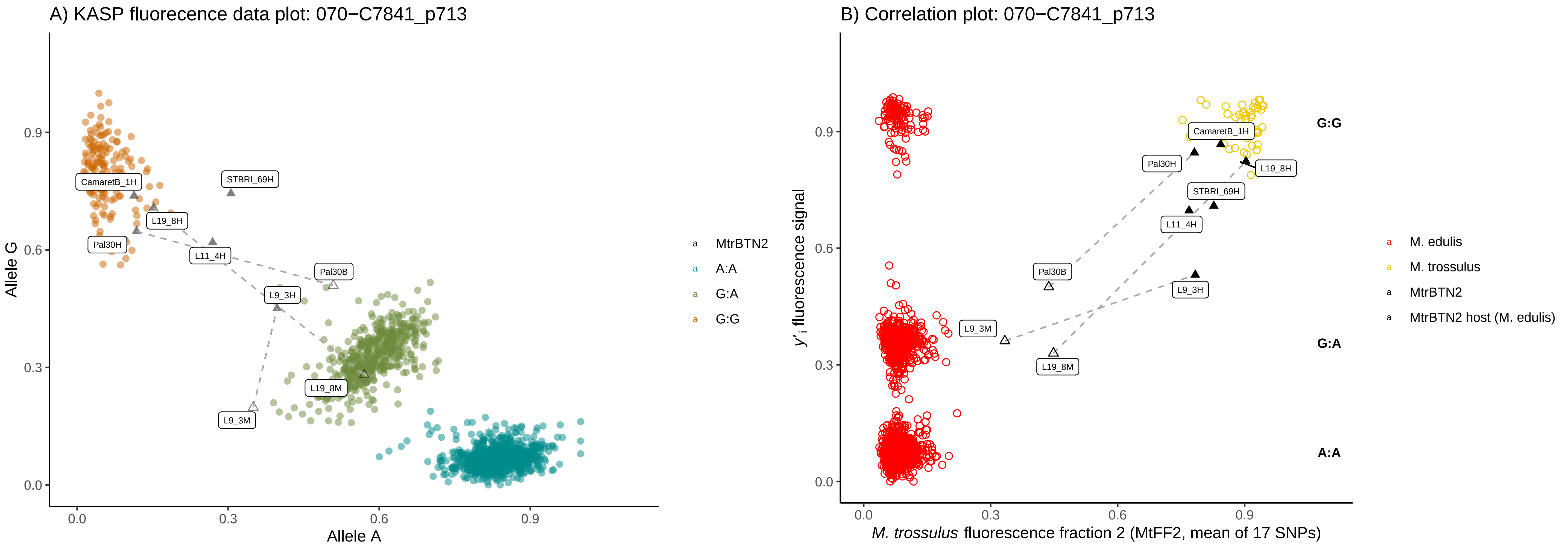
A) KASP fluorecence data plot: 070−C7841_p713
Allele G
Allele A
0.0
0.3
0.6
0.9
CamaretB_1H
STBRI_69H
L19_8H
Pal30H
L11_4H
Pal30B
L9_3H
L19_8M
L9_3M
MtrBTN2
A:A
G:A
G:G
B) Correlation plot: 070−C7841_p713
y'i fluorescence signal
M. trossulus fluorescence fraction 2 (MtFF2, mean of 17 SNPs)
G:G
G:A
A:A
M. edulis
M. trossulus
MtrBTN2
MtrBTN2 host (M. edulis)

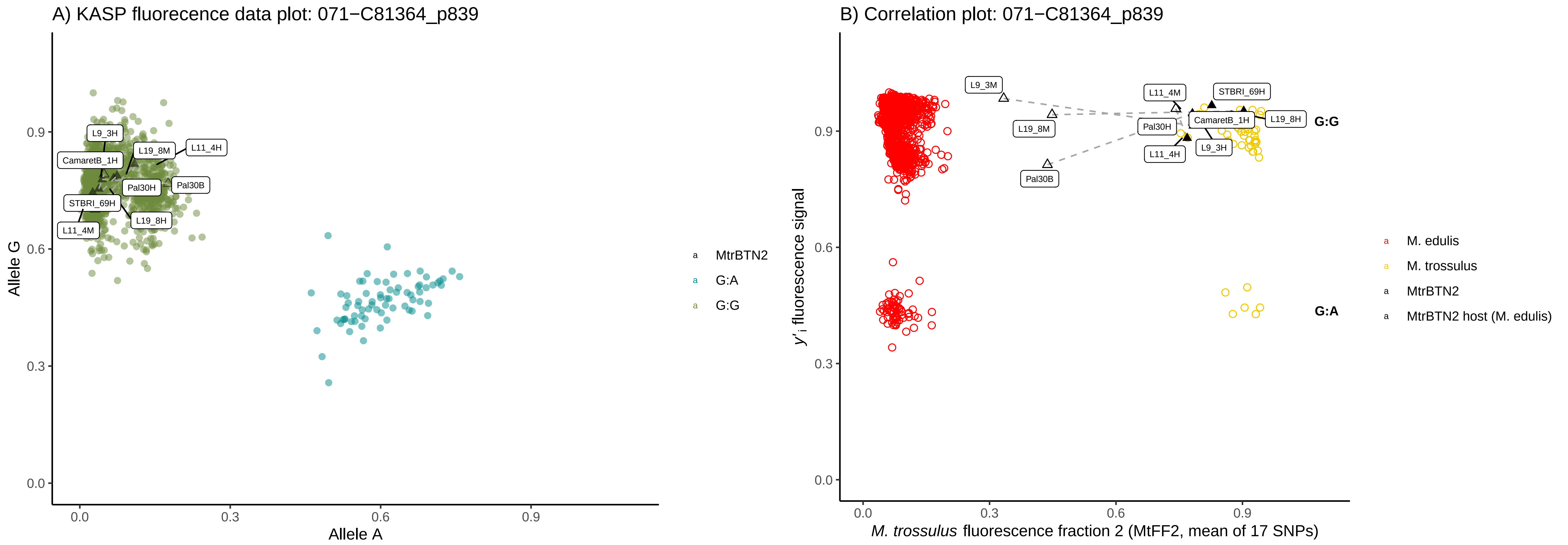

A) KASP fluorecence data plot: 071−C81364_p839
Allele G
Allele A
0.0
0.3
0.6
0.9
L9_3H
CamaretB_1H
L19_8M
L11_4H
Pal30H
Pal30B
STBRI_69H
L19_8H
L11_4M
MtrBTN2
G:A
G:G
B) Correlation plot: 071−C81364_p839
y'i fluorescence signal
M. trossulus fluorescence fraction 2 (MtFF2, mean of 17 SNPs)
L9_3M
L19_8M
Pal30B
L11_4M
STBRI_69H
Pal30H
CamaretB_1H
L19_8H
L11_4H
L9_3H
G:G
G:A
M. edulis
M. trossulus
MtrBTN2
MtrBTN2 host (M. edulis)

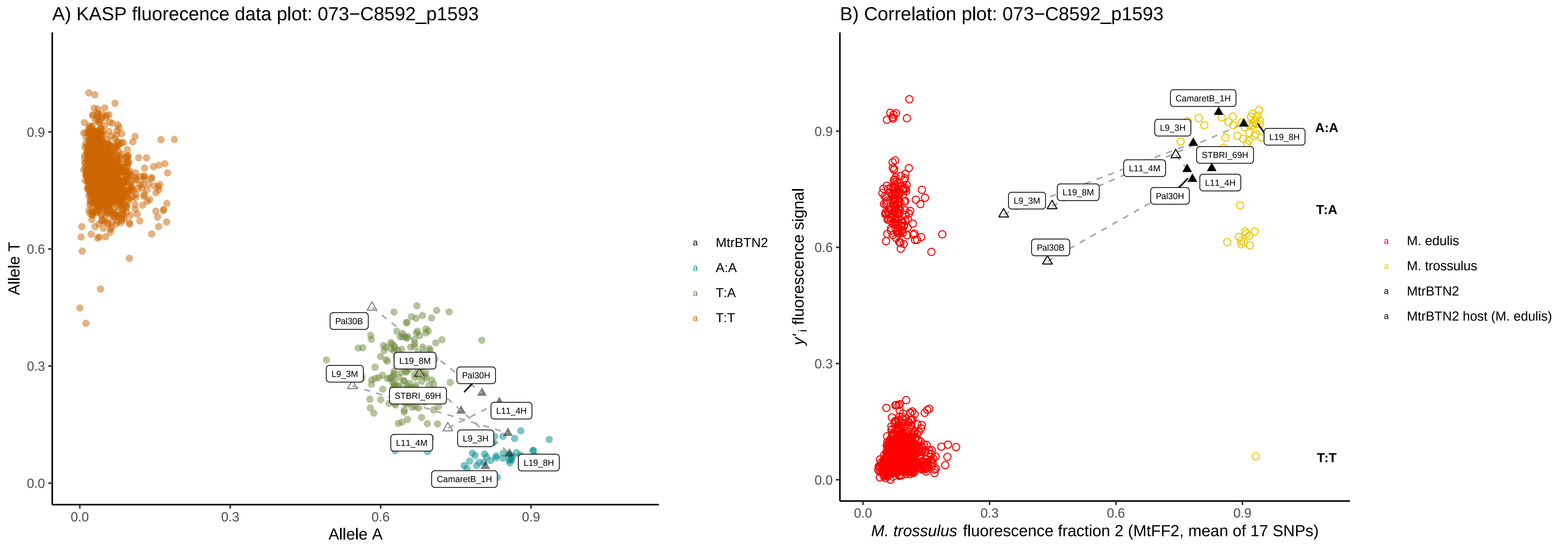
A) KASP fluorecence data plot: 073−C8592_p1593
Allele T
Allele A
0.0
0.3
0.6
0.9
Pal30B
L19_8M
L9_3M
Pal30H
STBRI_69H
L11_4H
L9_3H
L11_4M
L19_8H
CamaretB_1H
MtrBTN2
A:A
T:A
T:T
B) Correlation plot: 073−C8592_p1593
y'i fluorescence signal
M. trossulus fluorescence fraction 2 (MtFF2, mean of 17 SNPs)
CamaretB_1H
L9_3H
L19_8H
STBRI_69H
L11_4M
L11_4H
Pal30H
L19_8M
L9_3M
Pal30B
A:A
T:A
T:T
M. edulis
M. trossulus
MtrBTN2
MtrBTN2 host (M. edulis)

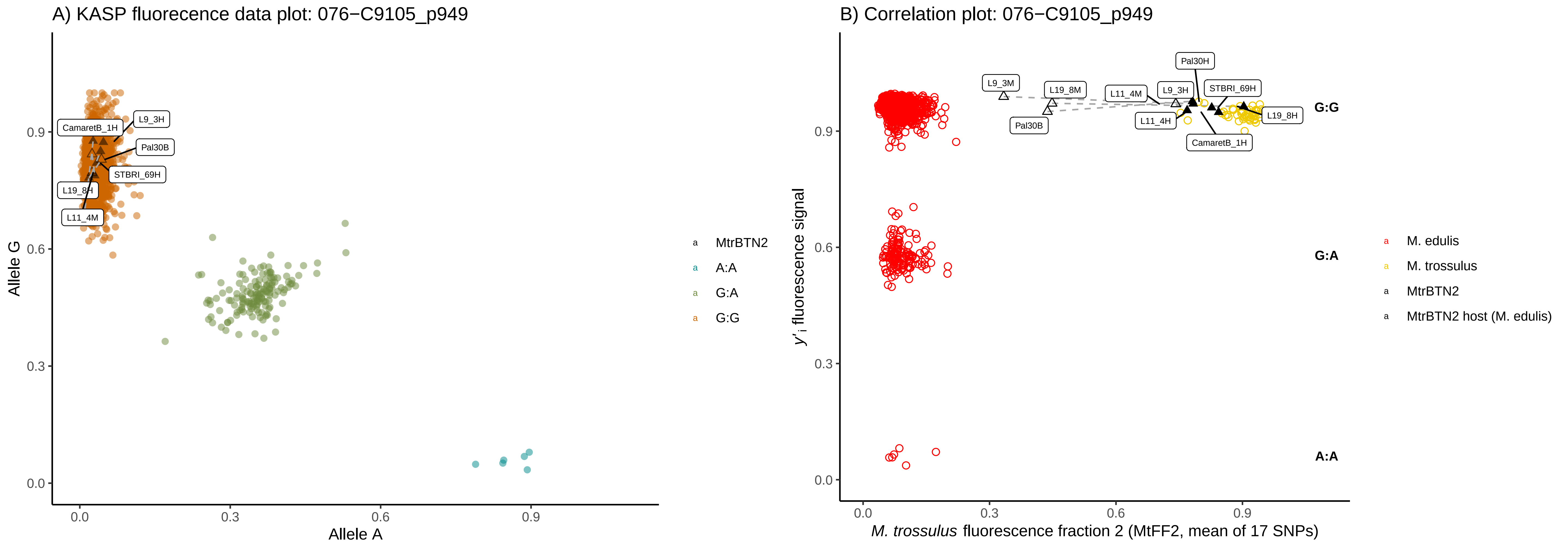

A) KASP fluorecence data plot: 076−C9105_p949
Allele G
Allele A
CamaretB_1H
L9_3H
Pal30B
STBRI_69H
L19_8H
L11_4M
MtrBTN2
A:A
G:A
G:G
B) Correlation plot: 076−C9105_p949
y'i fluorescence signal
M. trossulus fluorescence fraction 2 (MtFF2, mean of 17 SNPs)
Pal30H
L9_3M
L19_8M
L11_4M
L9_3H
STBRI_69H
Pal30B
L11_4H
L19_8H
CamaretB_1H
G:G
G:A
A:A
M. edulis
M. trossulus
MtrBTN2
MtrBTN2 host (M. edulis)

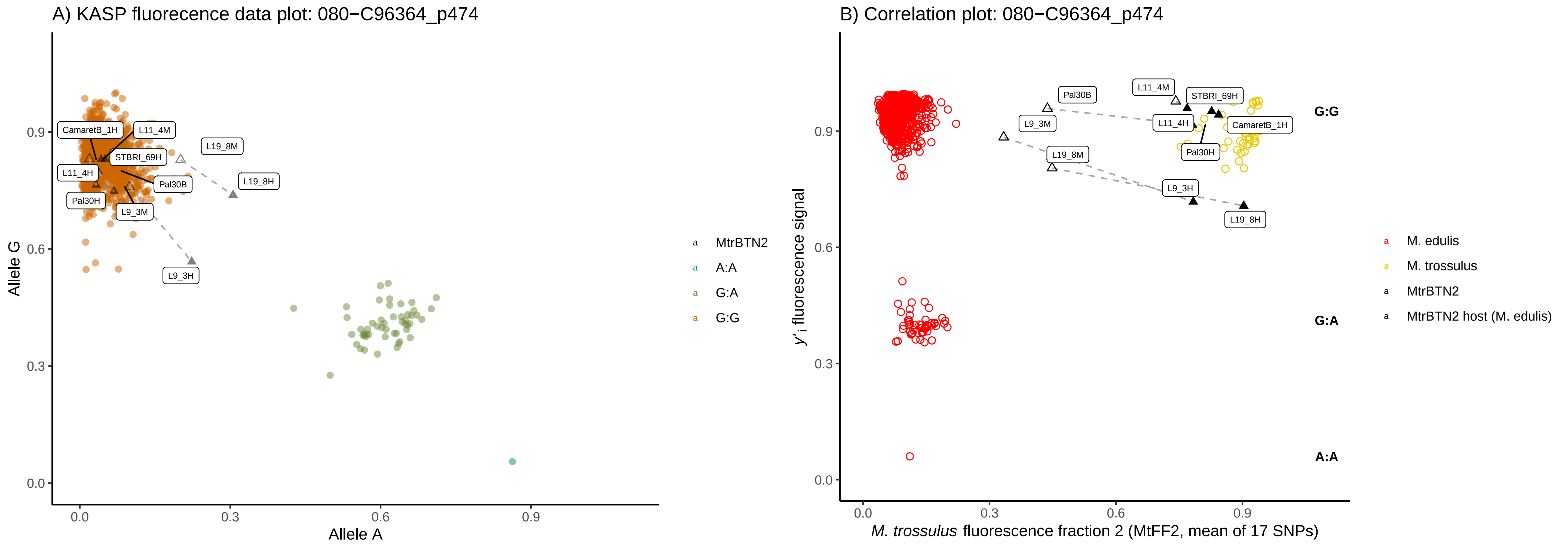
A) KASP fluorecence data plot: 080−C96364_p474
Allele G
Allele A
0.0
0.3
0.6
0.9
CamaretB_1H
L11_4M
L19_8M
STBRI_69H
L11_4H
Pal30B
L19_8H
Pal30H
L9_3M
L9_3H
MtrBTN2
A:A
G:A
G:G
B) Correlation plot: 080−C96364_p474
y'i fluorescence signal
M. trossulus fluorescence fraction 2 (MtFF2, mean of 17 SNPs)
G:G
G:A
A:A
M. edulis
M. trossulus
MtrBTN2
MtrBTN2 host (M. edulis)

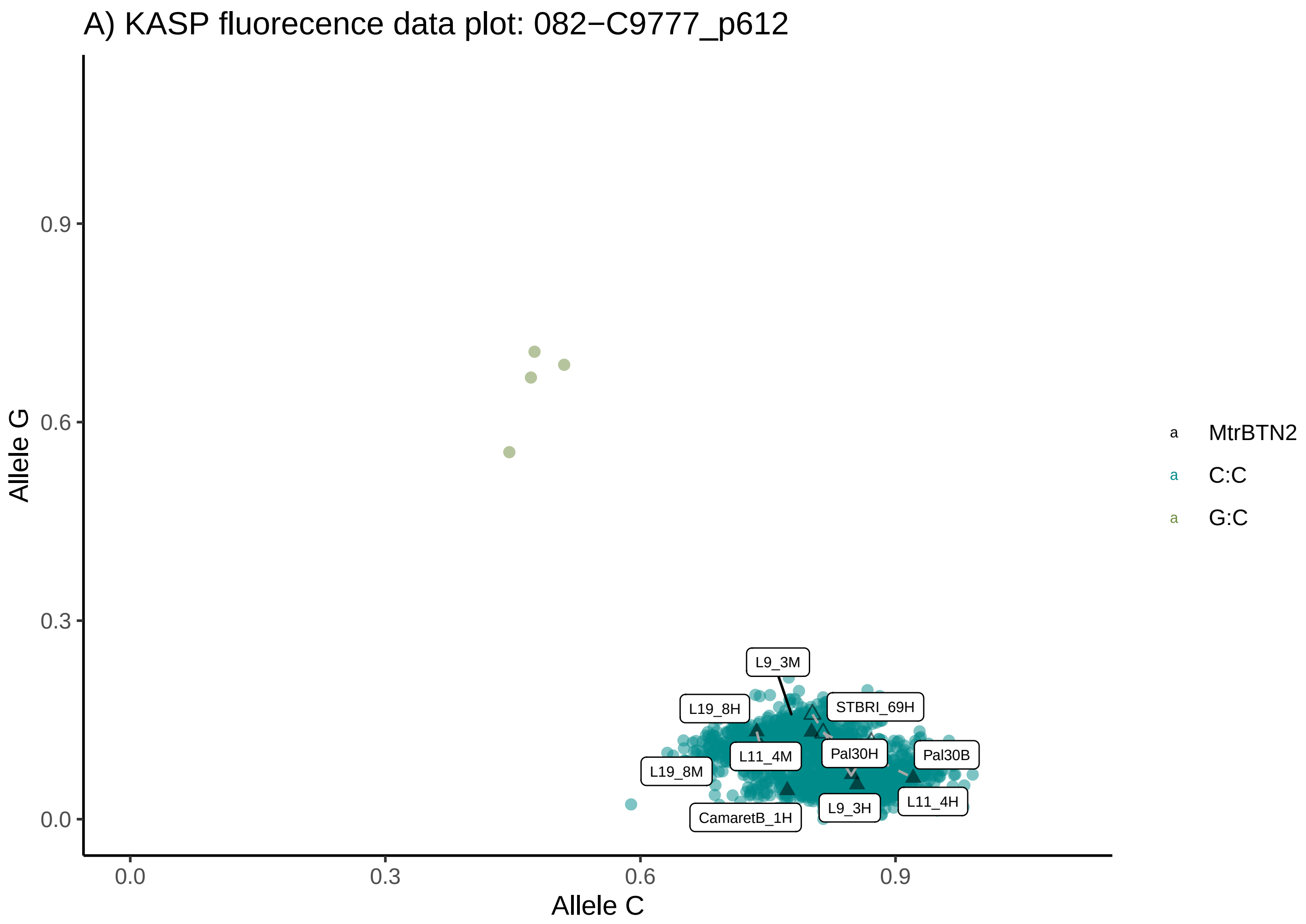
A) KASP fluorecence data plot: 082−C9777_p612
Allele G
Allele C
0.0
0.3
0.6
0.9
L9_3M
L19_8H
STBRI_69H
L11_4M
Pal30H
Pal30B
L19_8M
CamaretB_1H
L9_3H
L11_4H
MtrBTN2
C:C
G:C

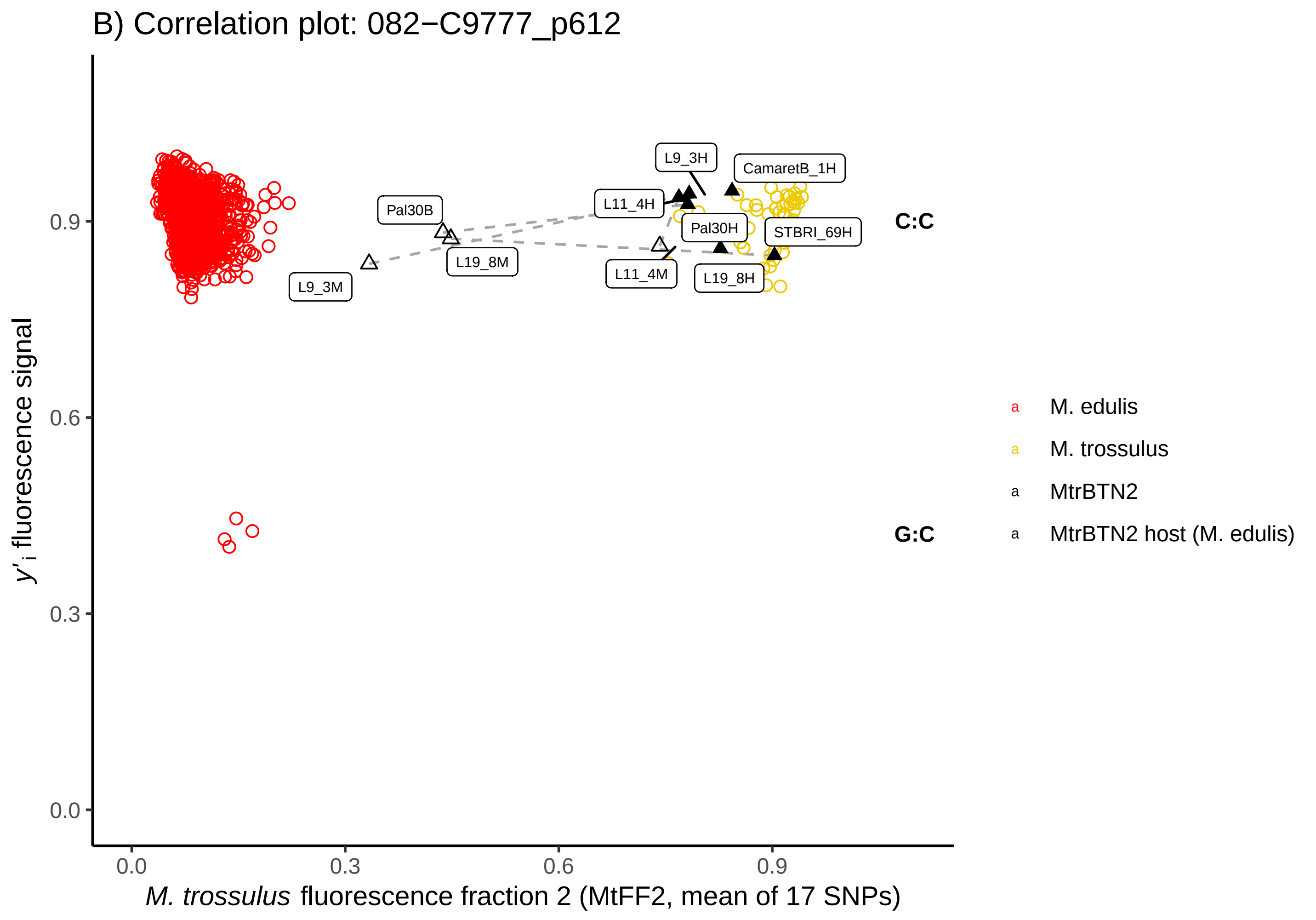
B) Correlation plot: 082−C9777_p612
y'i fluorescence signal
M. trossulus fluorescence fraction 2 (MtFF2, mean of 17 SNPs)
0.0
0.3
0.6
0.9
L9_3H
CamaretB_1H
Pal30B
L11_4H
Pal30H
STBRI_69H
L19_8M
L9_3M
L11_4M
L19_8H
C:C
G:C
M. edulis
M. trossulus
MtrBTN2
MtrBTN2 host (M. edulis)

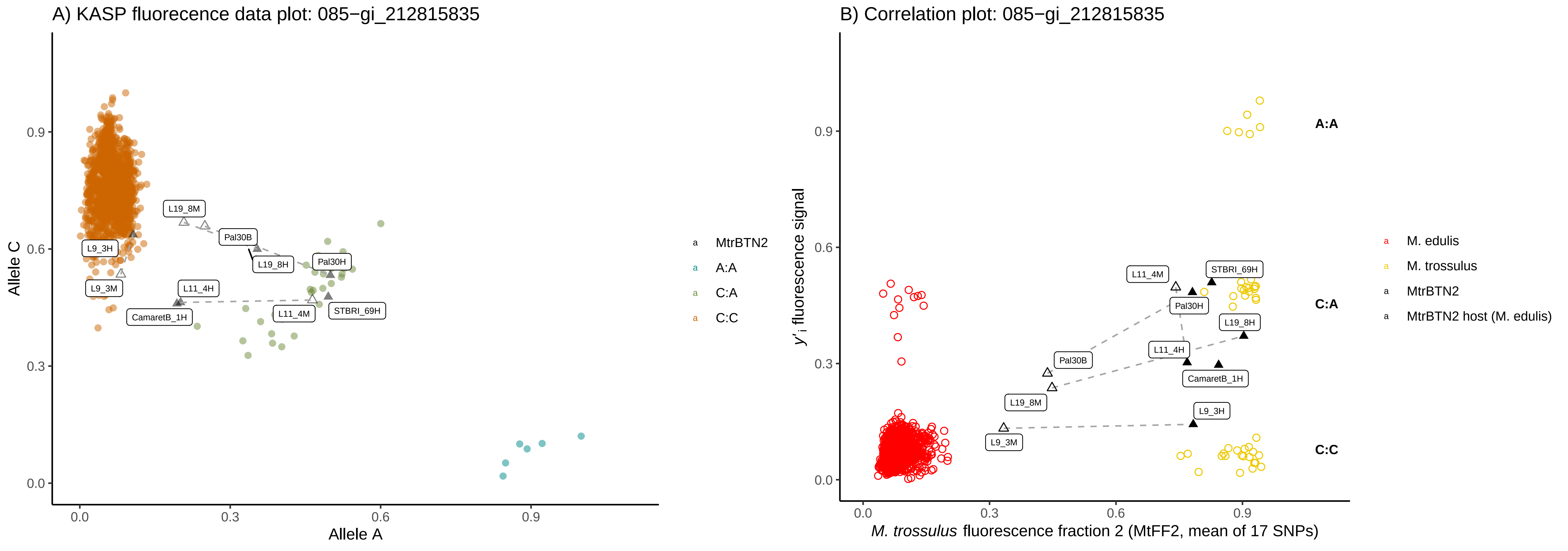
A) KASP fluorecence data plot: 085−gi_212815835
Allele C
Allele A
0.0
0.3
0.6
0.9
L19_8M
Pal30B
L19_8H
Pal30H
L9_3H
L9_3M
L11_4H
L11_4M
STBRI_69H
CamaretB_1H
MtrBTN2
A:A
C:A
C:C
B) Correlation plot: 085−gi_212815835
y'i fluorescence signal
M. trossulus fluorescence fraction 2 (MtFF2, mean of 17 SNPs)
A:A
C:A
C:C
M. edulis
M. trossulus
MtrBTN2
MtrBTN2 host (M. edulis)

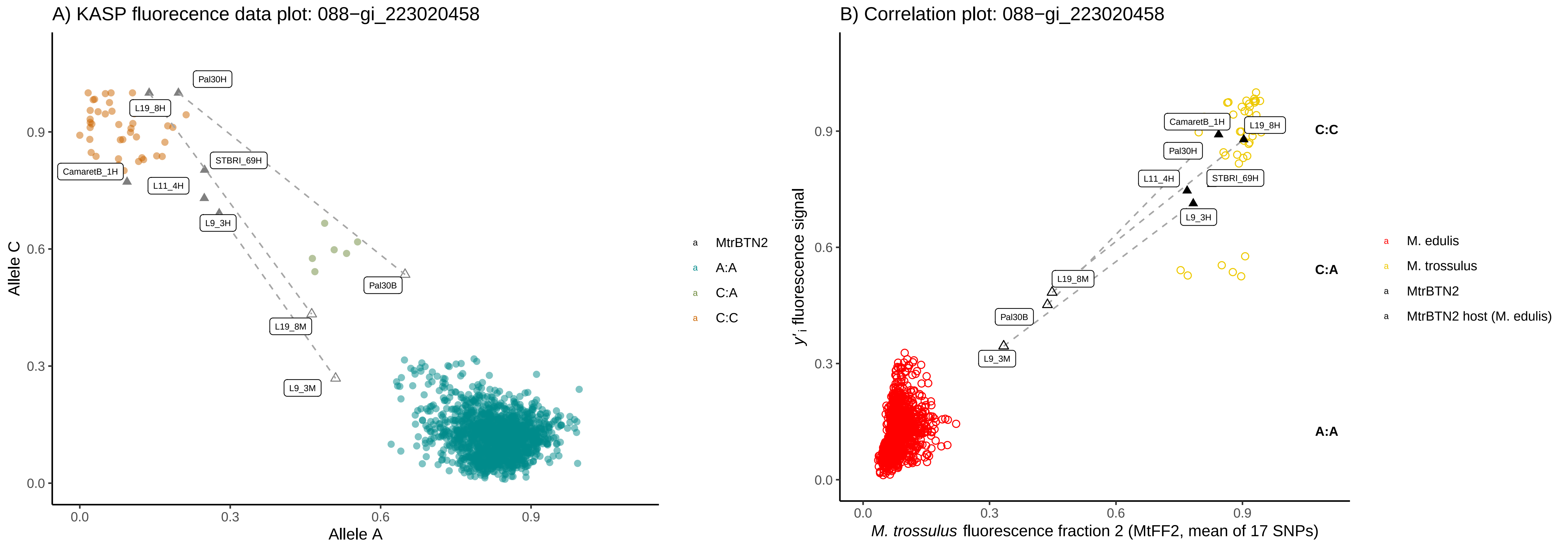
A) KASP fluorecence data plot: 088−gi_223020458
Allele C
Allele A
0.0
0.3
0.6
0.9
Pal30H
L19_8H
STBRI_69H
CamaretB_1H
L11_4H
L9_3H
Pal30B
L19_8M
L9_3M
MtrBTN2
A:A
C:A
C:C
B) Correlation plot: 088−gi_223020458
y'i fluorescence signal
M. trossulus fluorescence fraction 2 (MtFF2, mean of 17 SNPs)
CamaretB_1H
L19_8H
Pal30H
L11_4H
STBRI_69H
L9_3H
L19_8M
Pal30B
L9_3M
C:C
C:A
A:A
M. edulis
M. trossulus
MtrBTN2
MtrBTN2 host (M. edulis)

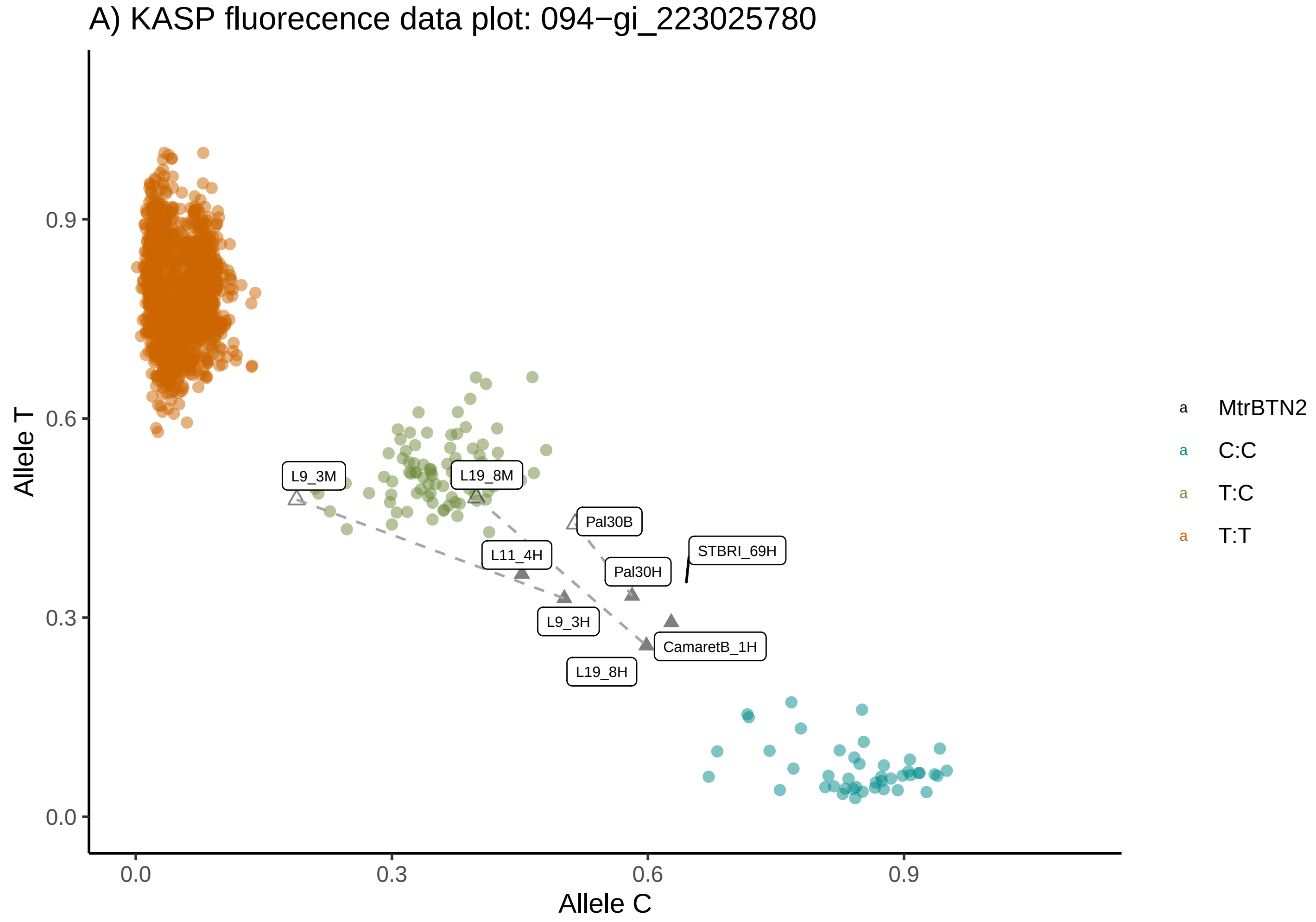
A) KASP fluorecence data plot: 094−gi_223025780
Allele T
Allele C
0.0
0.3
0.6
0.9
L9_3M
L19_8M
Pal30B
L11_4H
STBRI_69H
Pal30H
L9_3H
CamaretB_1H
L19_8H
MtrBTN2
C:C
T:C
T:T

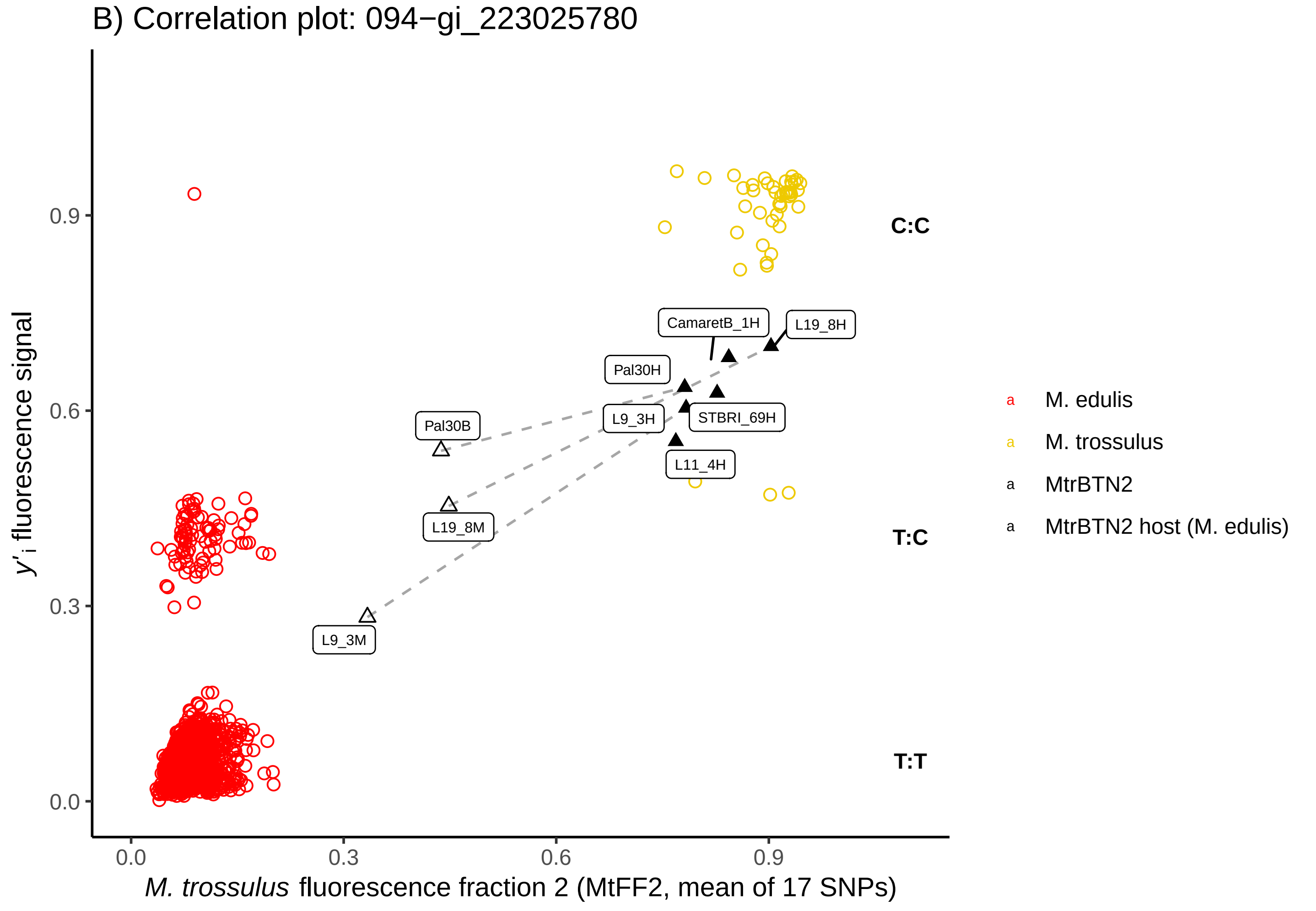
B) Correlation plot: 094−gi_223025780
y'i fluorescence signal
M. trossulus fluorescence fraction 2 (MtFF2, mean of 17 SNPs)
0.0
0.3
0.6
0.9
C:C
T:C
T:T
CamaretB_1H
L19_8H
Pal30H
Pal30B
L9_3H
STBRI_69H
L11_4H
L19_8M
L9_3M
M. edulis
M. trossulus
MtrBTN2
MtrBTN2 host (M. edulis)

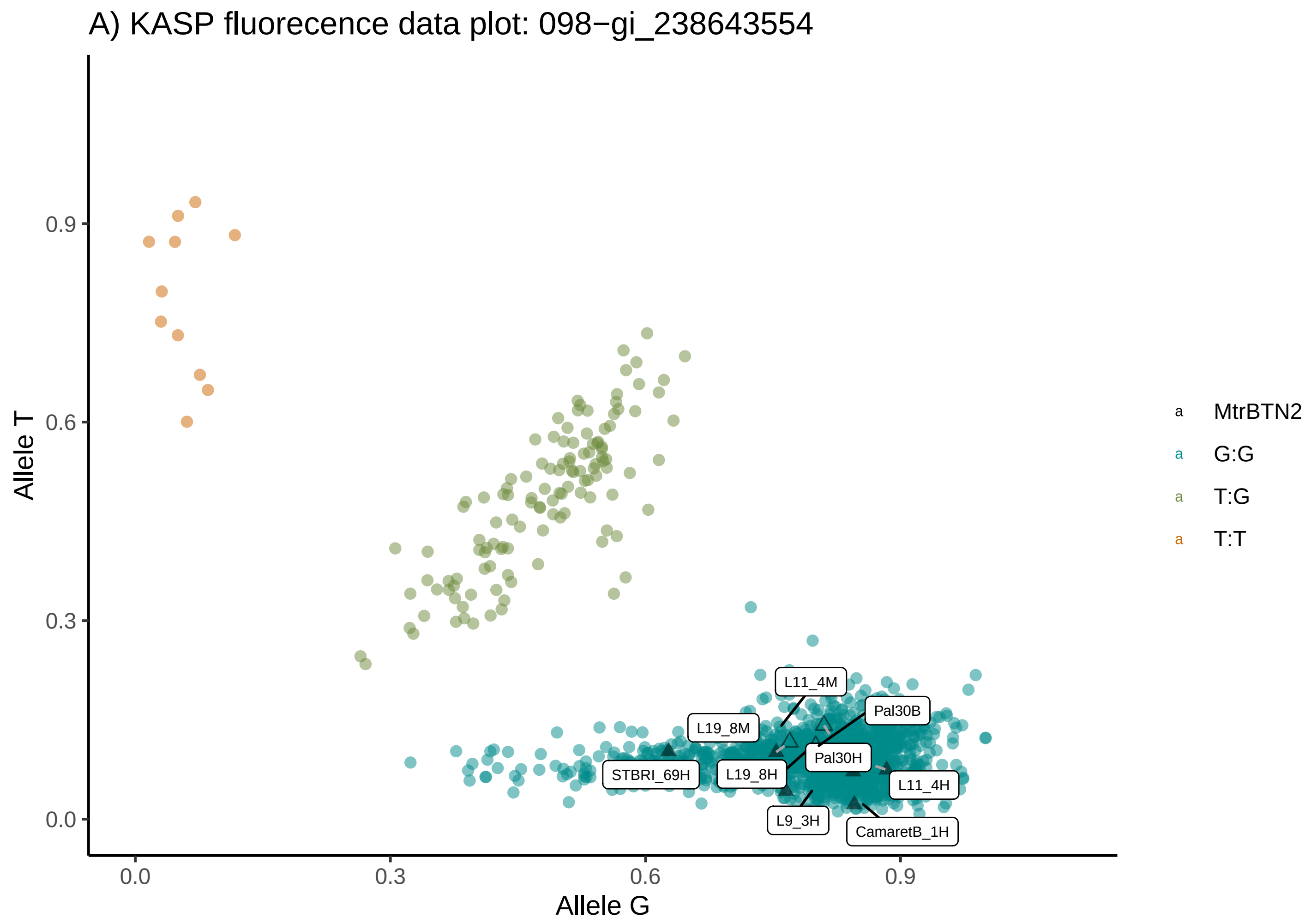
A) KASP fluorecence data plot: 098−gi_238643554
Allele T
Allele G
0.0
0.3
0.6
0.9
L11_4M
L19_8M
Pal30B
Pal30H
STBRI_69H
L19_8H
L11_4H
L9_3H
CamaretB_1H
MtrBTN2
G:G
T:G
T:T

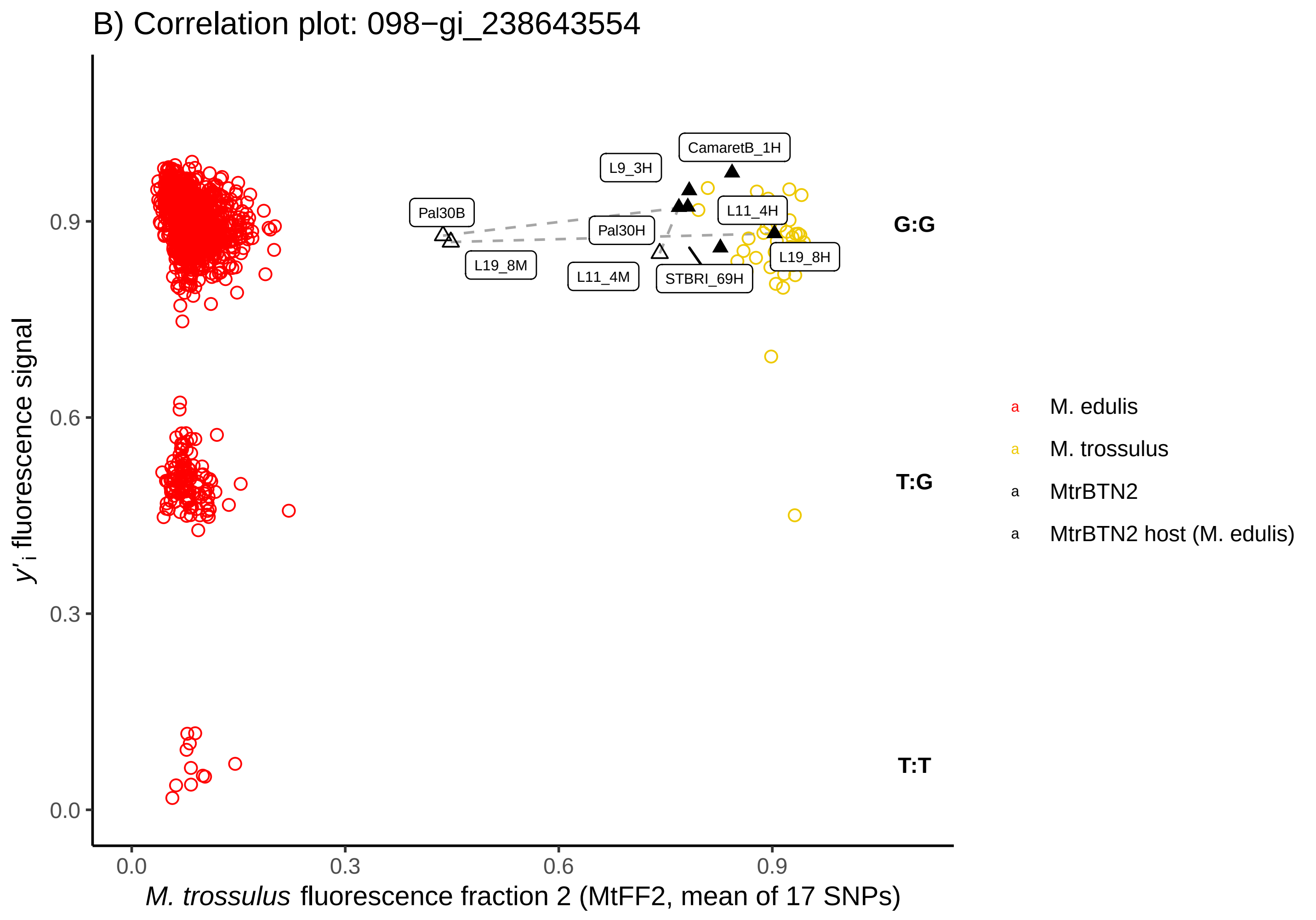
B) Correlation plot: 098−gi_238643554
y'i fluorescence signal
M. trossulus fluorescence fraction 2 (MtFF2, mean of 17 SNPs)
0.0
0.3
0.6
0.9
CamaretB_1H
L9_3H
Pal30B
L11_4H
Pal30H
L19_8M
L19_8H
L11_4M
STBRI_69H
G:G
T:G
T:T
M. edulis
M. trossulus
MtrBTN2
MtrBTN2 host (M. edulis)

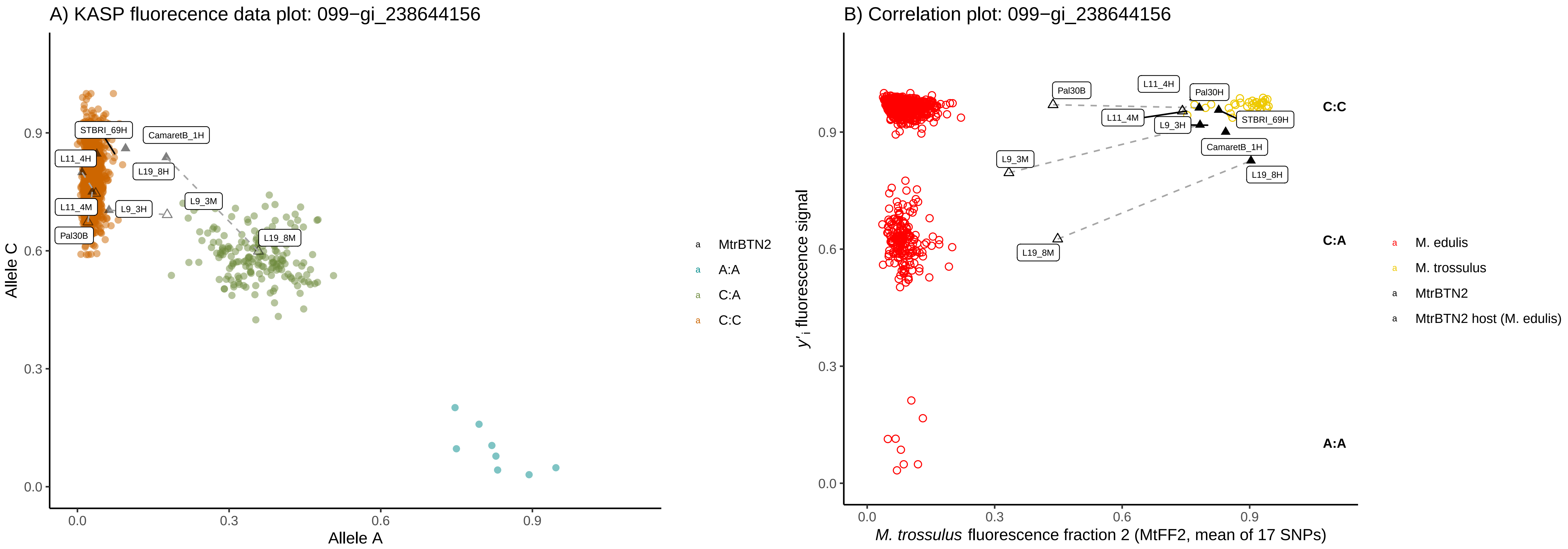

A) KASP fluorecence data plot: 099−gi_238644156
Allele C
Allele A
STBRI_69H
CamaretB_1H
L11_4H
L19_8H
L9_3M
L11_4M
L9_3H
Pal30B
L19_8M
MtrBTN2
A:A
C:A
C:C
B) Correlation plot: 099−gi_238644156
y'i fluorescence signal
M. trossulus fluorescence fraction 2 (MtFF2, mean of 17 SNPs)
Pal30B
L11_4H
Pal30H
L11_4M
L9_3H
STBRI_69H
CamaretB_1H
L9_3M
L19_8H
L19_8M
C:C
C:A
A:A
M. edulis
M. trossulus
MtrBTN2
MtrBTN2 host (M. edulis)

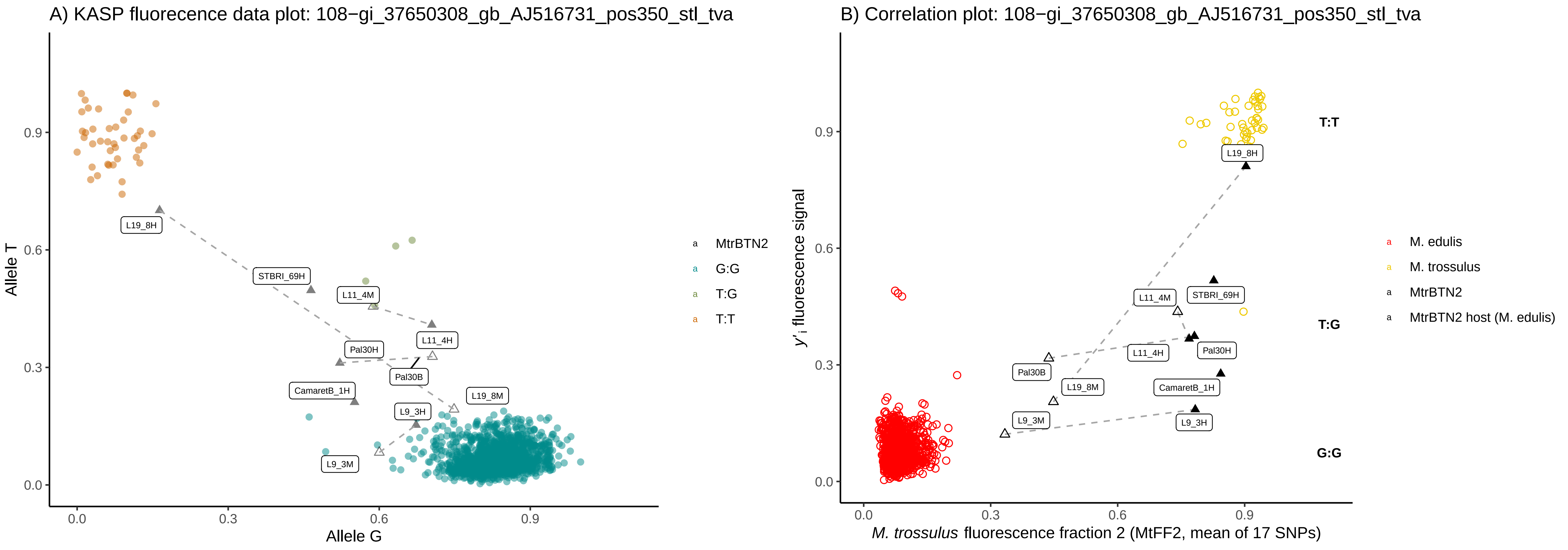
A) KASP fluorecence data plot: 108−gi_37650308_gb_AJ516731_pos350_stl_tva
Allele T
Allele G
0.0
0.3
0.6
0.9
L19_8H
STBRI_69H
L11_4M
L11_4H
Pal30H
Pal30B
CamaretB_1H
L19_8M
L9_3H
L9_3M
MtrBTN2
G:G
T:G
T:T
B) Correlation plot: 108−gi_37650308_gb_AJ516731_pos350_stl_tva
y'i fluorescence signal
M. trossulus fluorescence fraction 2 (MtFF2, mean of 17 SNPs)
T:T
T:G
G:G
L19_8H
L11_4M
STBRI_69H
L11_4H
Pal30H
Pal30B
L19_8M
CamaretB_1H
L9_3M
L9_3H
M. edulis
M. trossulus
MtrBTN2
MtrBTN2 host (M. edulis)

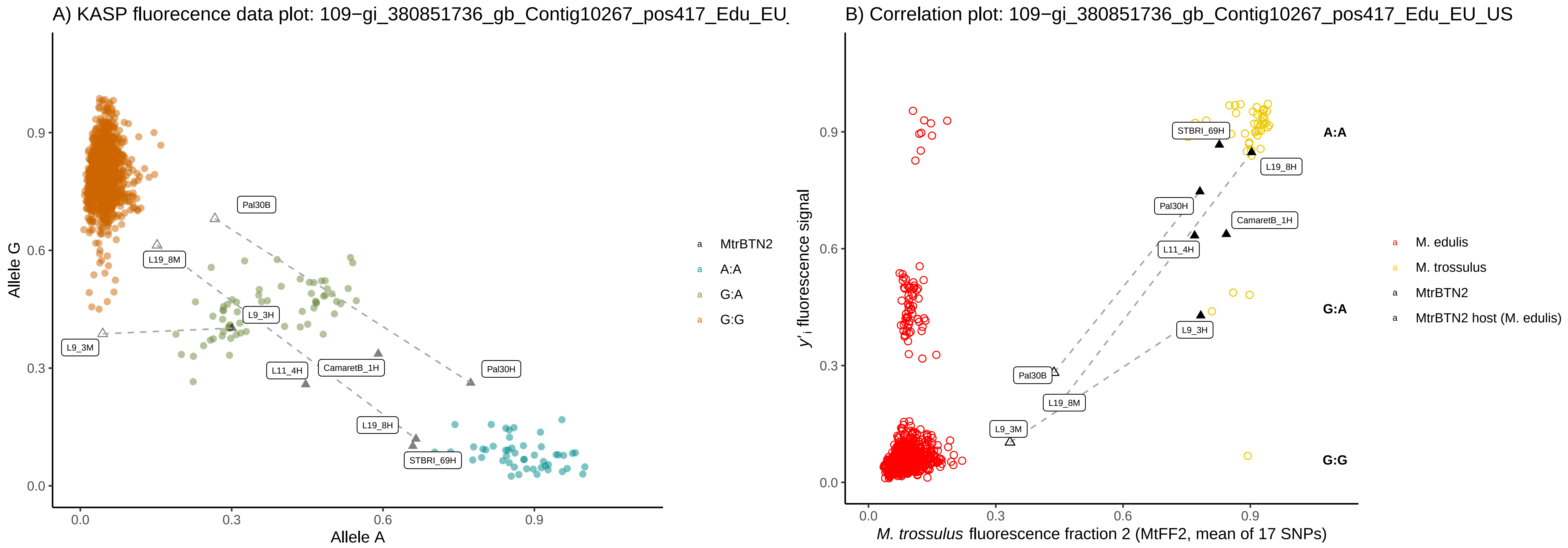

A) KASP fluorecence data plot: 109−gi_380851736_gb_Contig10267_pos417_Edu_EU_
Allele G
Allele A
0.0
0.3
0.6
0.9
Pal30B
L19_8M
L9_3H
L9_3M
L11_4H
CamaretB_1H
Pal30H
L19_8H
STBRI_69H
MtrBTN2
A:A
G:A
G:G
B) Correlation plot: 109−gi_380851736_gb_Contig10267_pos417_Edu_EU_US
y'i fluorescence signal
M. trossulus fluorescence fraction 2 (MtFF2, mean of 17 SNPs)
A:A
G:A
G:G
M. edulis
M. trossulus
MtrBTN2
MtrBTN2 host (M. edulis)

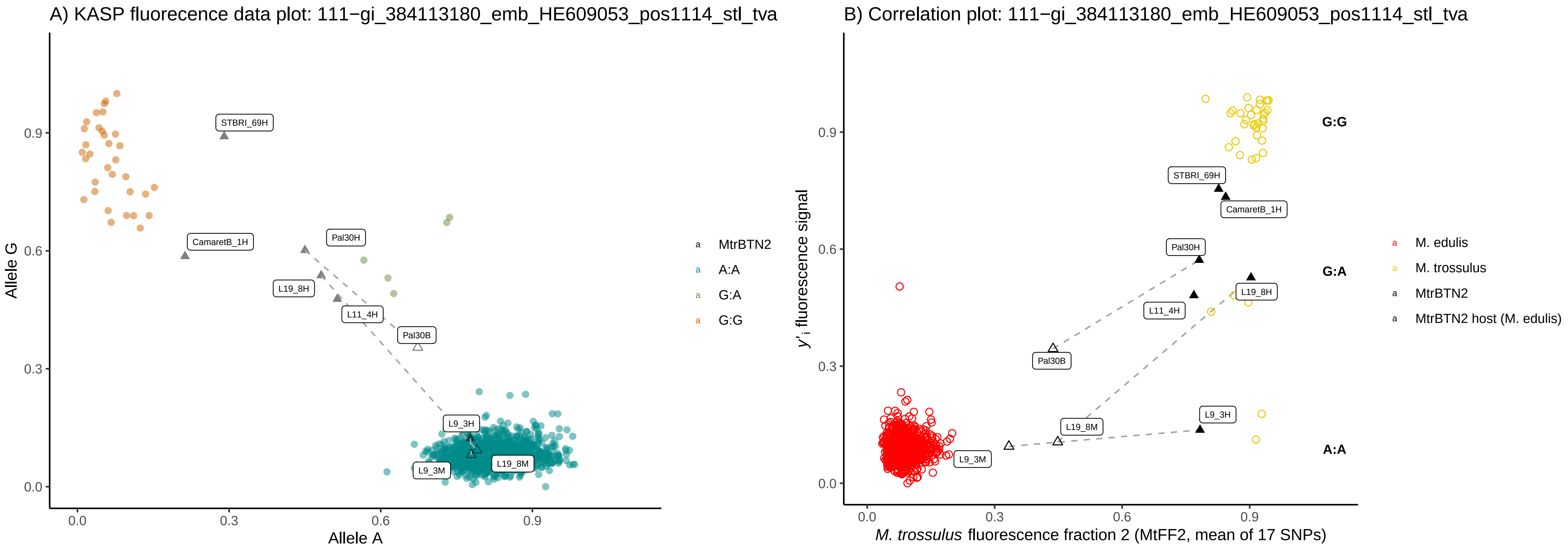
A) KASP fluorecence data plot: 111−gi_384113180_emb_HE609053_pos1114_stl_tva
Allele G
Allele A
0.0
0.3
0.6
0.9
STBRI_69H
CamaretB_1H
Pal30H
L19_8H
L11_4H
Pal30B
L9_3H
L9_3M
L19_8M
MtrBTN2
A:A
G:A
G:G
B) Correlation plot: 111−gi_384113180_emb_HE609053_pos1114_stl_tva
y'i fluorescence signal
M. trossulus fluorescence fraction 2 (MtFF2, mean of 17 SNPs)
G:G
G:A
A:A
M. edulis
M. trossulus
MtrBTN2
MtrBTN2 host (M. edulis)

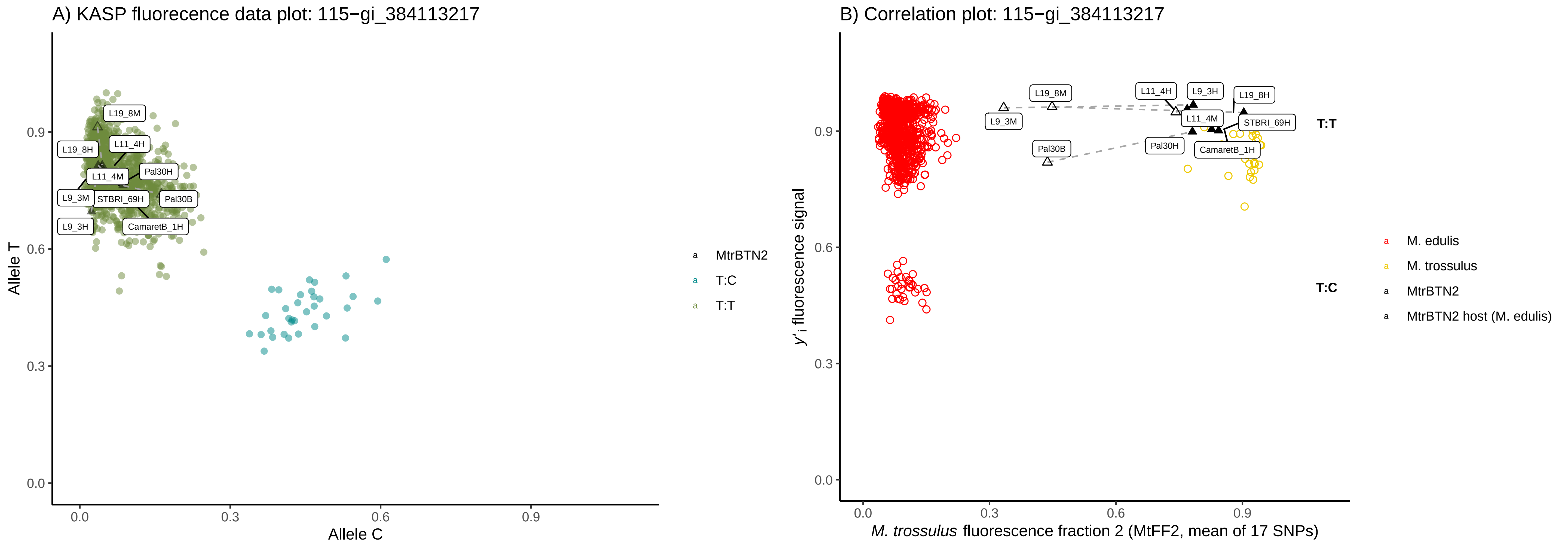
A) KASP fluorecence data plot: 115−gi_384113217
Allele T
Allele C
0.0
0.3
0.6
0.9
L19_8M
L19_8H
L11_4H
L11_4M
Pal30H
L9_3M
STBRI_69H
Pal30B
L9_3H
CamaretB_1H
MtrBTN2
T:C
T:T
B) Correlation plot: 115−gi_384113217
y'i fluorescence signal
M. trossulus fluorescence fraction 2 (MtFF2, mean of 17 SNPs)
L19_8M
L11_4H
L9_3H
L19_8H
L9_3M
L11_4M
STBRI_69H
Pal30B
Pal30H
CamaretB_1H
T:T
T:C
M. edulis
M. trossulus
MtrBTN2
MtrBTN2 host (M. edulis)

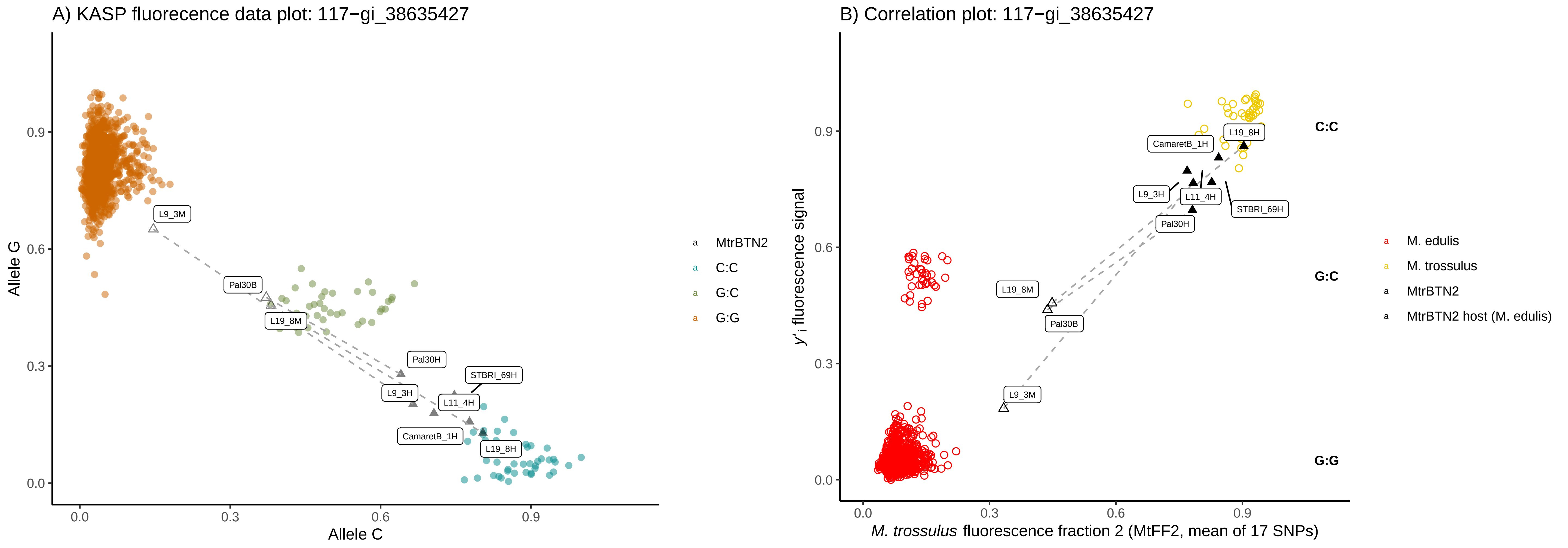
A) KASP fluorecence data plot: 117−gi_38635427
Allele G
Allele C
0.0
0.3
0.6
0.9
L9_3M
Pal30B
L19_8M
Pal30H
STBRI_69H
L9_3H
L11_4H
CamaretB_1H
L19_8H
MtrBTN2
C:C
G:C
G:G
B) Correlation plot: 117−gi_38635427
y'i fluorescence signal
M. trossulus fluorescence fraction 2 (MtFF2, mean of 17 SNPs)
C:C
G:C
G:G
CamaretB_1H
L19_8H
L9_3H
L11_4H
STBRI_69H
Pal30H
L19_8M
Pal30B
L9_3M
M. edulis
M. trossulus
MtrBTN2
MtrBTN2 host (M. edulis)

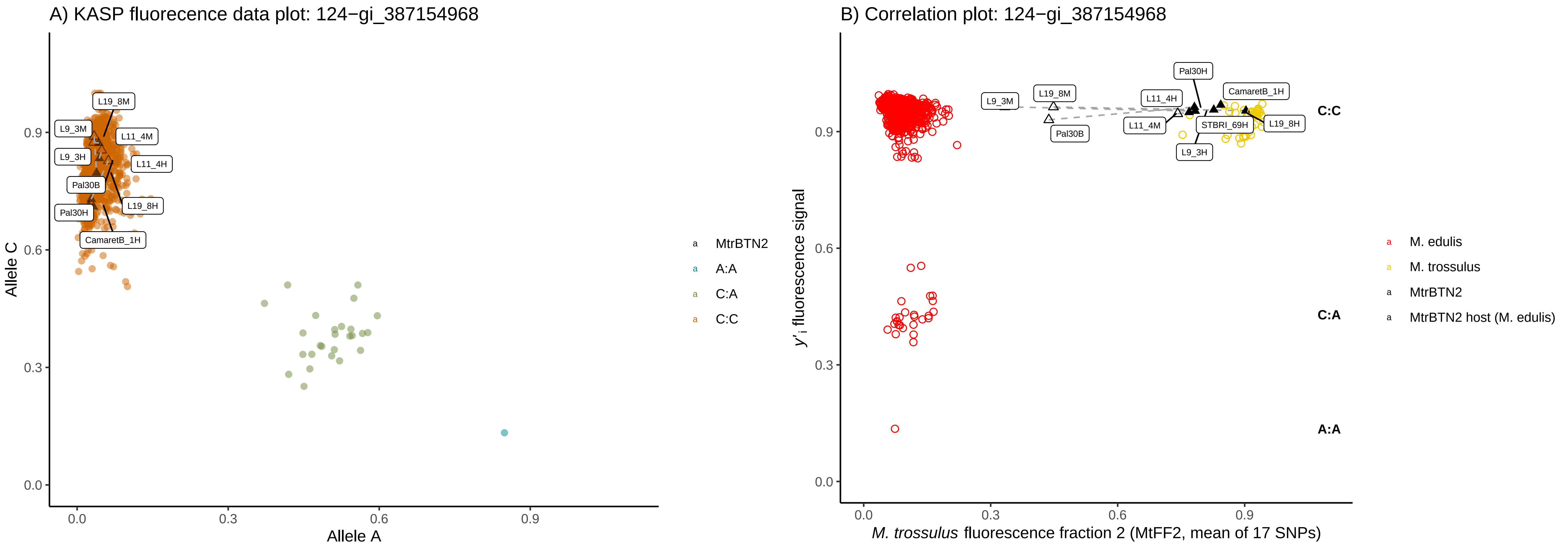
A) KASP fluorecence data plot: 124−gi_387154968
L19_8M
L9_3M
L11_4M
L9_3H
L11_4H
Pal30B
L19_8H
Pal30H
CamaretB_1H
Allele C
Allele A
0.0
0.3
0.6
0.9
MtrBTN2
A:A
C:A
C:C
B) Correlation plot: 124−gi_387154968
Pal30H
CamaretB_1H
L9_3M
L19_8M
L11_4H
L11_4M
STBRI_69H
L19_8H
Pal30B
L9_3H
C:C
C:A
A:A
y'i fluorescence signal
M. trossulus fluorescence fraction 2 (MtFF2, mean of 17 SNPs)
M. edulis
M. trossulus
MtrBTN2
MtrBTN2 host (M. edulis)

A) KASP fluorecence data plot: 126−gi_387154971

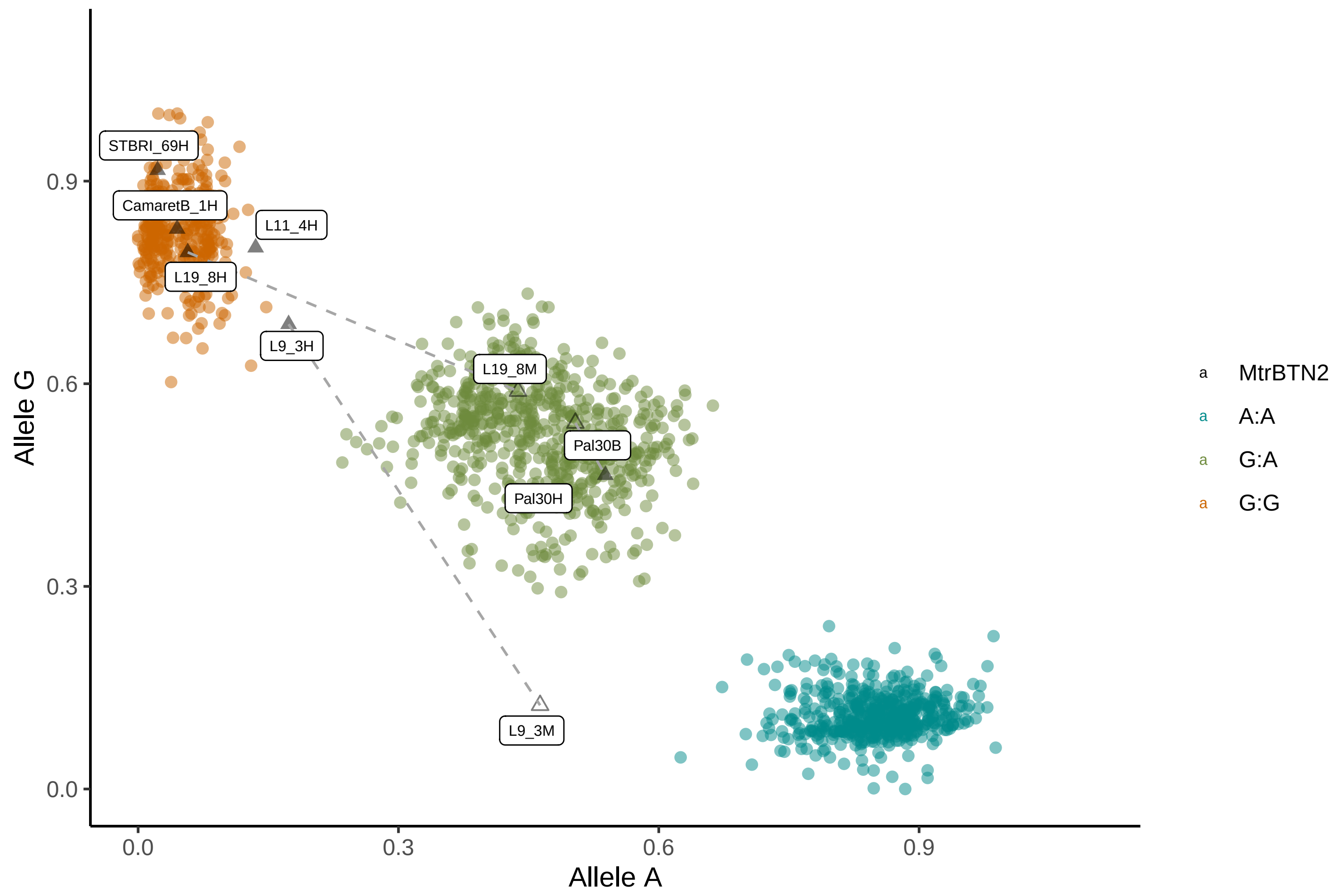


B) Correlation plot: 126−gi_387154971

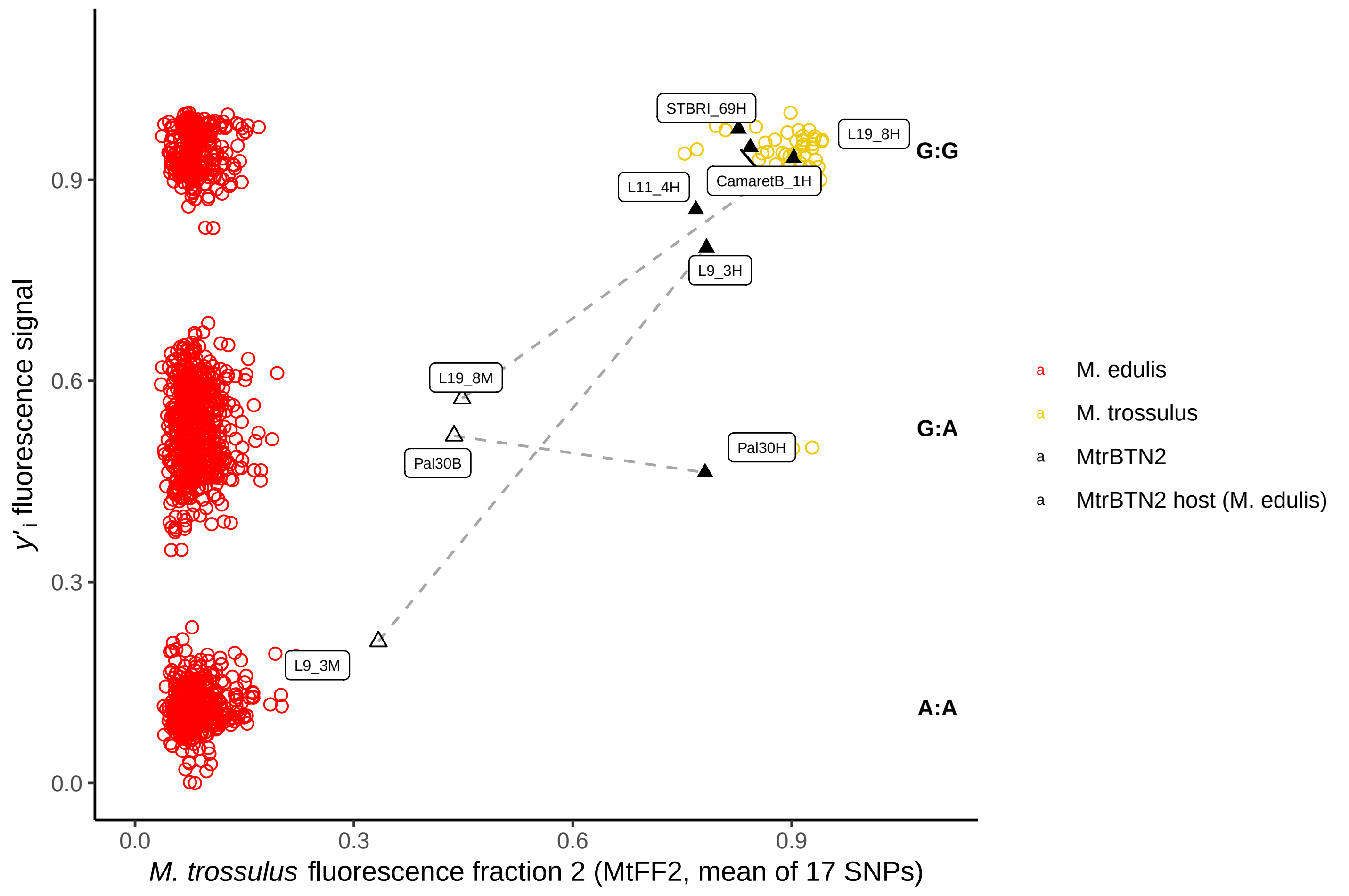

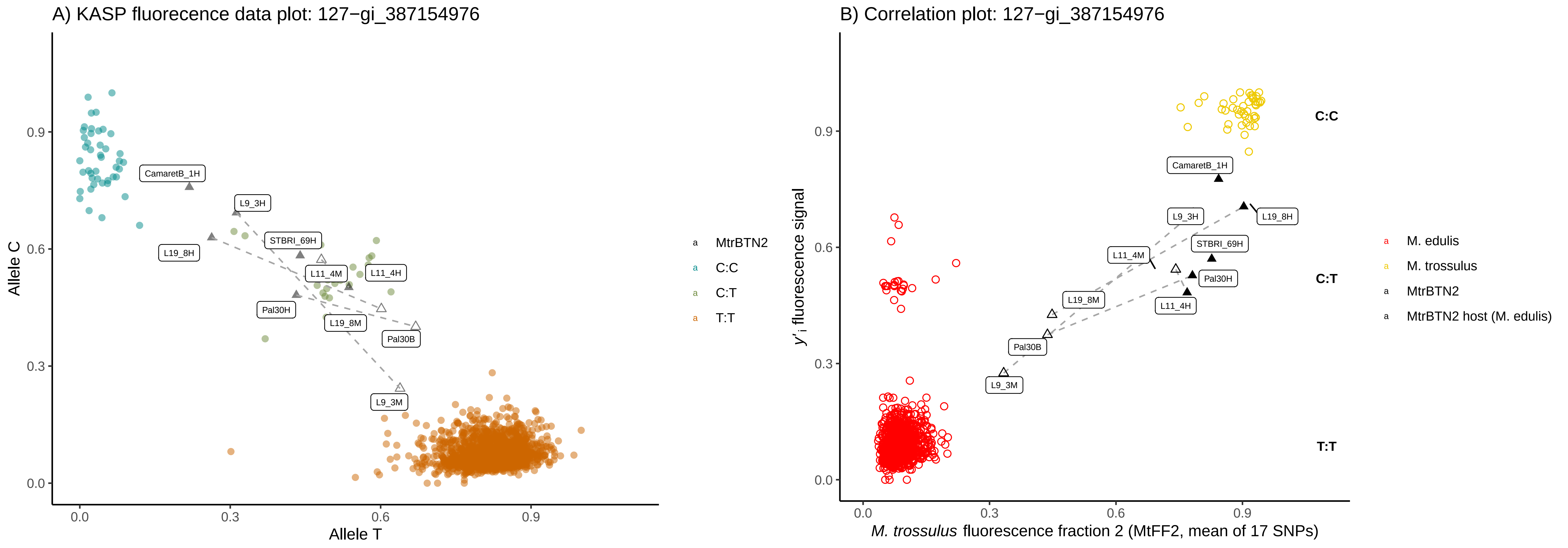
A) KASP fluorecence data plot: 127−gi_387154976
Allele C
Allele T
0.0
0.3
0.6
0.9
CamaretB_1H
L9_3H
STBRI_69H
L19_8H
L11_4M
L11_4H
Pal30H
L19_8M
Pal30B
L9_3M
MtrBTN2
C:C
C:T
T:T
B) Correlation plot: 127−gi_387154976
y'i fluorescence signal
M. trossulus fluorescence fraction 2 (MtFF2, mean of 17 SNPs)
C:C
C:T
T:T
M. edulis
M. trossulus
MtrBTN2
MtrBTN2 host (M. edulis)

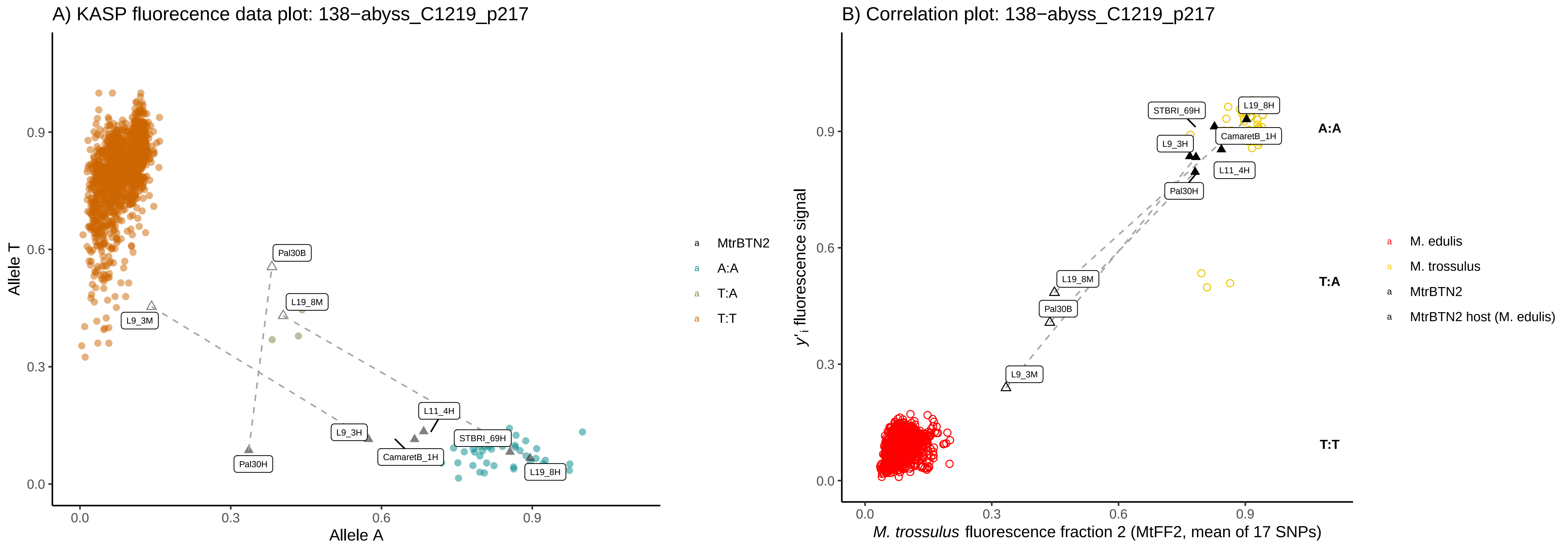

A) KASP fluorecence data plot: 138−abyss_C1219_p217
Allele T
Allele A
0.0
0.3
0.6
0.9
Pal30B
L19_8M
L9_3M
L11_4H
L9_3H
STBRI_69H
CamaretB_1H
Pal30H
L19_8H
MtrBTN2
A:A
T:A
T:T
B) Correlation plot: 138−abyss_C1219_p217
$y'_i$ fluorescence signal
*M. trossulus* fluorescence fraction 2 (MtFF2, mean of 17 SNPs)
STBRI_69H
L19_8H
CamaretB_1H
L9_3H
L11_4H
Pal30H
L19_8M
Pal30B
L9_3M
A:A
T:A
T:T
M. edulis
M. trossulus
MtrBTN2
MtrBTN2 host (M. edulis)

A) KASP fluorecence data plot: 145−abyss_C216_p5751

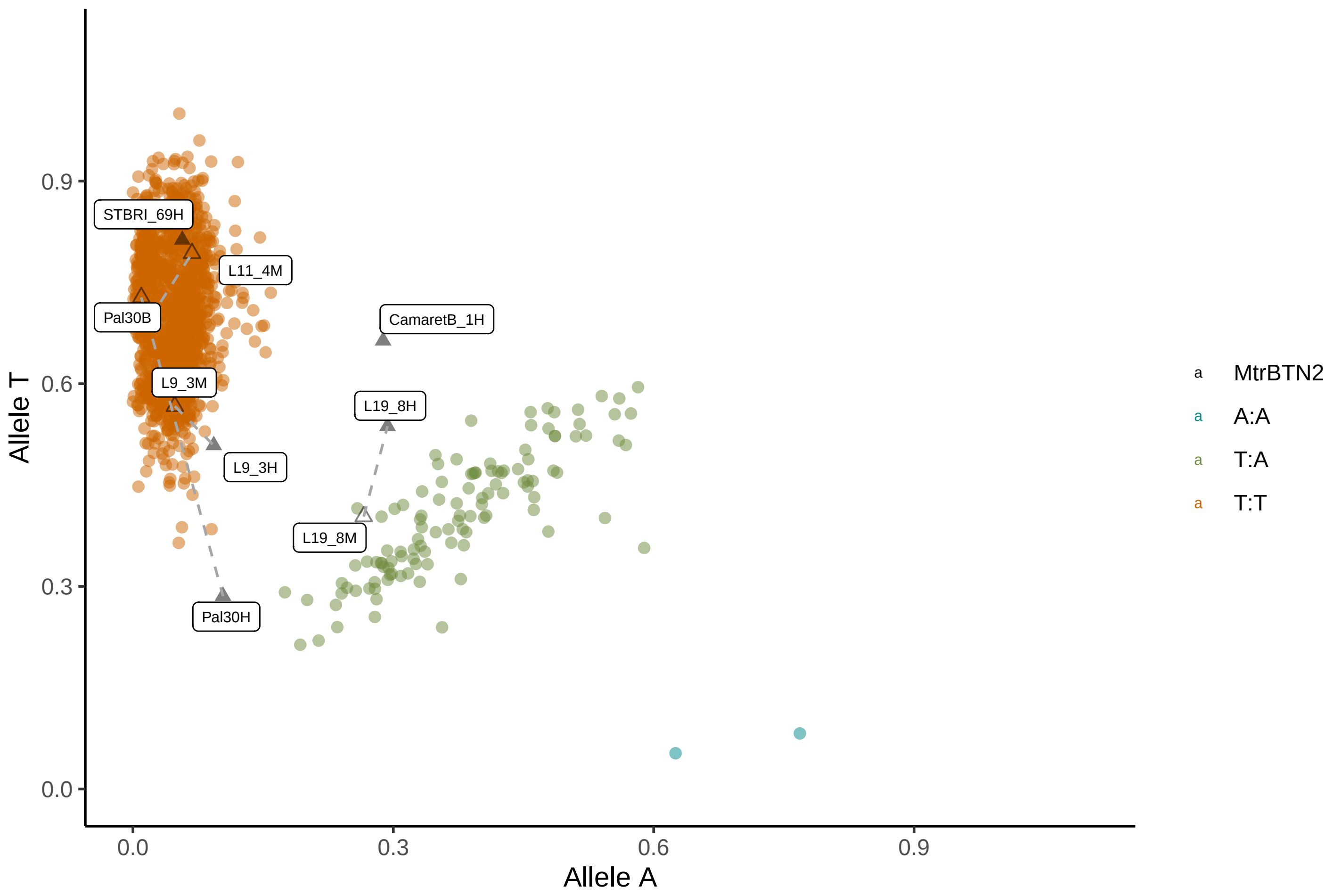


B) Correlation plot: 145−abyss_C216_p5751

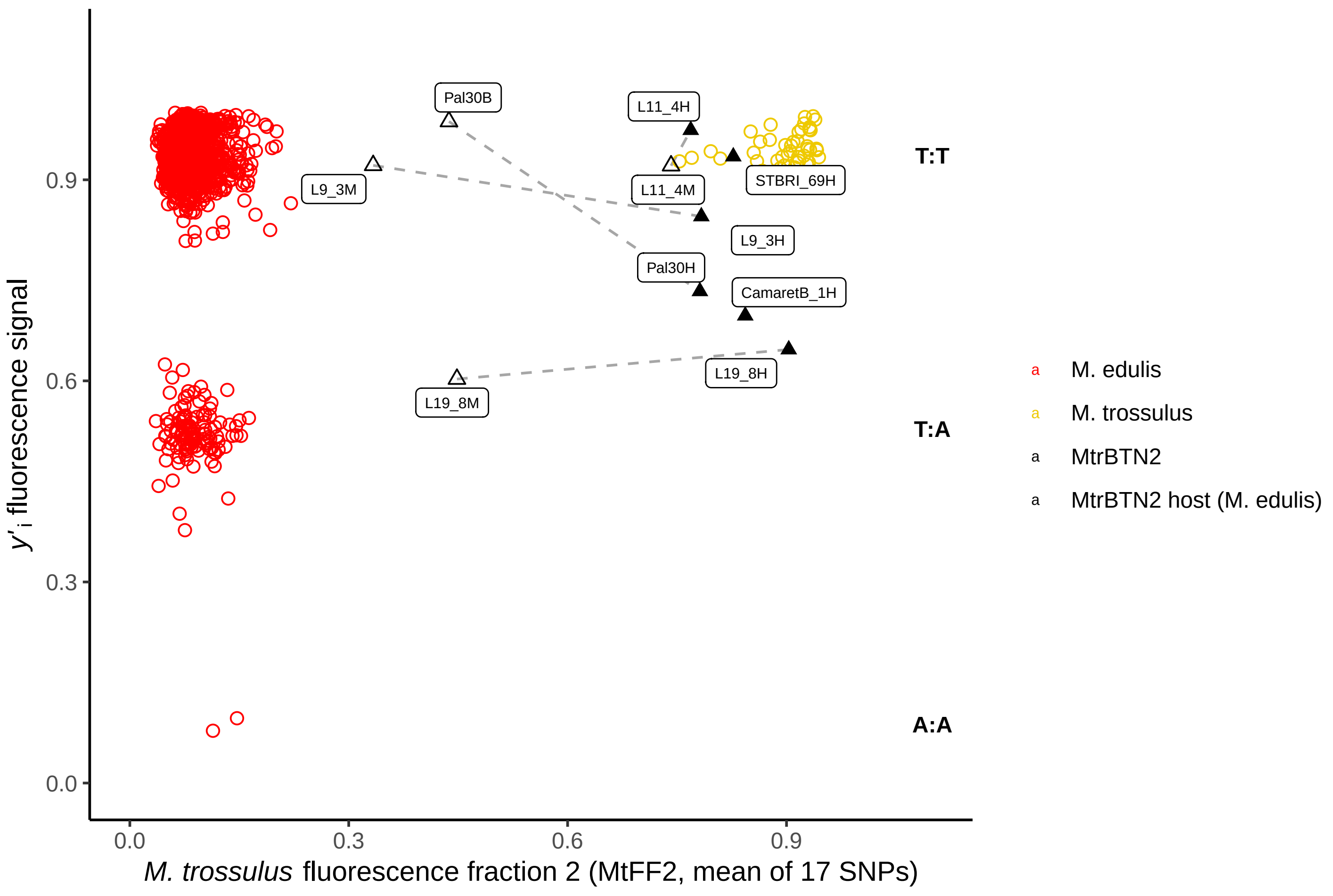

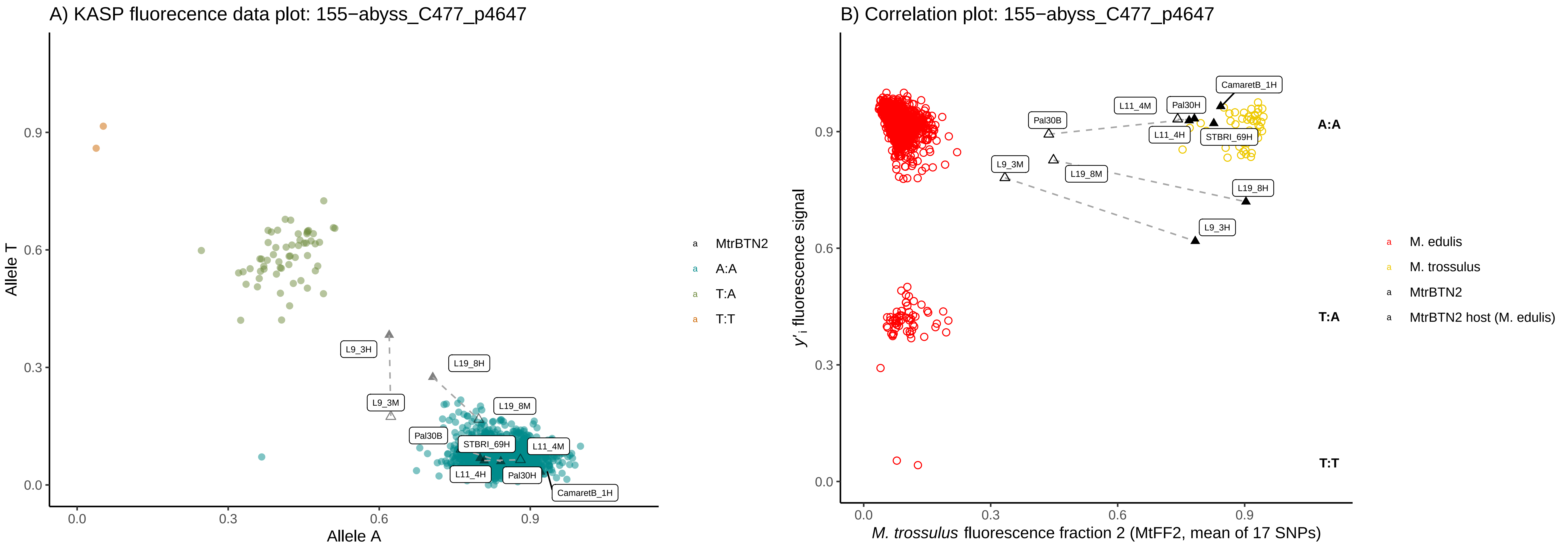
A) KASP fluorecence data plot: 155−abyss_C477_p4647
Allele T
Allele A
0.0
0.3
0.6
0.9
L9_3H
L19_8H
L9_3M
L19_8M
Pal30B
STBRI_69H
L11_4M
L11_4H
Pal30H
CamaretB_1H
MtrBTN2
A:A
T:A
T:T
B) Correlation plot: 155−abyss_C477_p4647
y'i fluorescence signal
M. trossulus fluorescence fraction 2 (MtFF2, mean of 17 SNPs)
CamaretB_1H
L11_4M
Pal30H
Pal30B
L11_4H
STBRI_69H
L9_3M
L19_8M
L19_8H
L9_3H
A:A
T:A
T:T
M. edulis
M. trossulus
MtrBTN2
MtrBTN2 host (M. edulis)

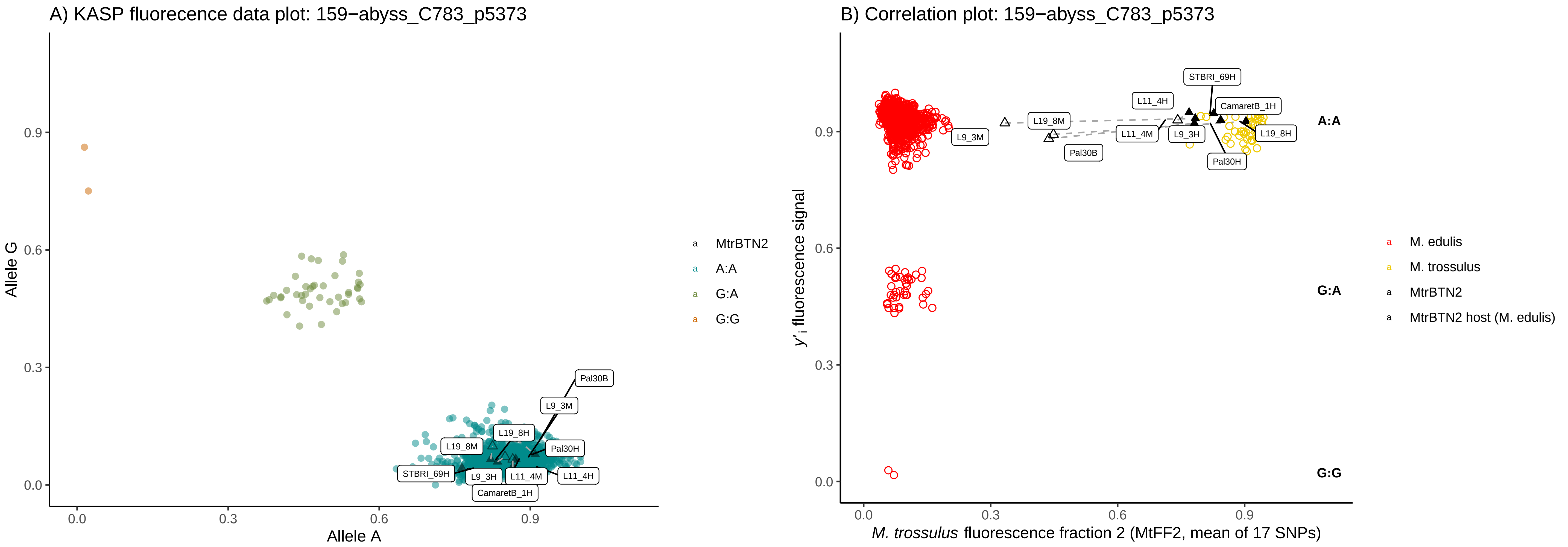

A) KASP fluorecence data plot: 159−abyss_C783_p5373
Allele G
Allele A
0.0
0.3
0.6
0.9
Pal30B
L9_3M
L19_8H
L19_8M
Pal30H
STBRI_69H
L9_3H
L11_4M
L11_4H
CamaretB_1H
MtrBTN2
A:A
G:A
G:G
B) Correlation plot: 159−abyss_C783_p5373
y'i fluorescence signal
M. trossulus fluorescence fraction 2 (MtFF2, mean of 17 SNPs)
STBRI_69H
L11_4H
CamaretB_1H
L19_8M
L11_4M
L9_3H
L19_8H
L9_3M
Pal30B
Pal30H
A:A
G:A
G:G
M. edulis
M. trossulus
MtrBTN2
MtrBTN2 host (M. edulis)

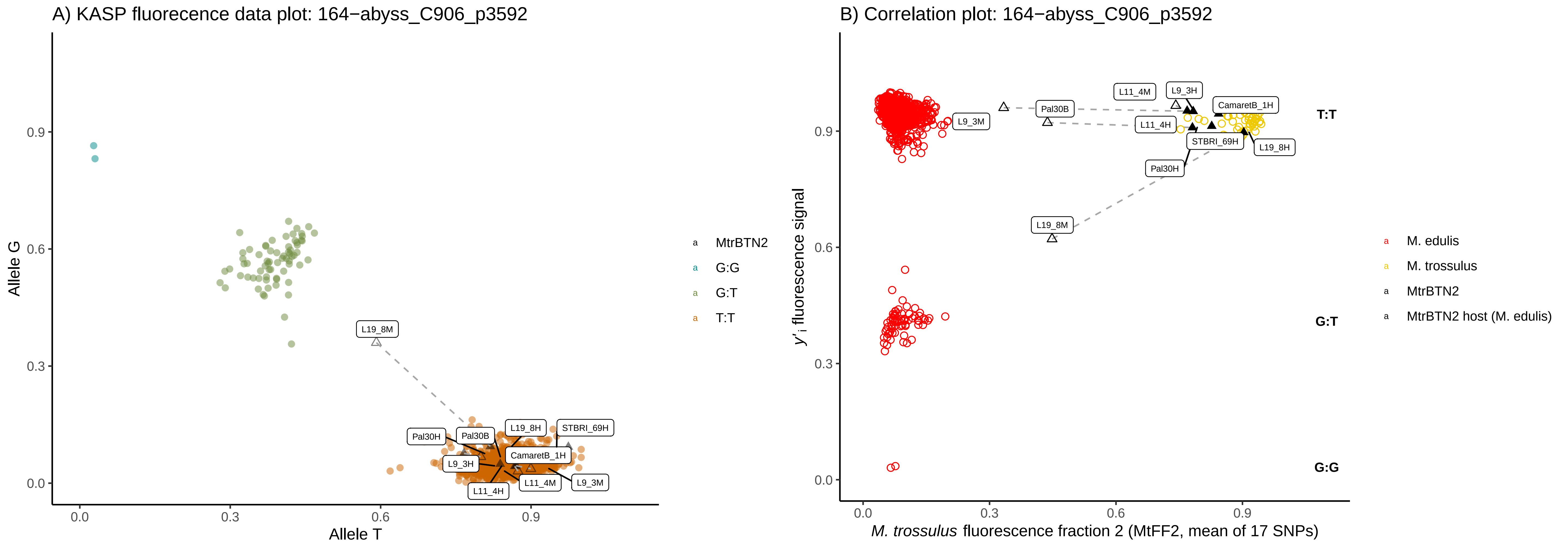
A) KASP fluorecence data plot: 164−abyss_C906_p3592
Allele G
Allele T
0.0
0.3
0.6
0.9
L19_8M
Pal30H
Pal30B
L19_8H
STBRI_69H
L9_3H
CamaretB_1H
L11_4M
L9_3M
L11_4H
MtrBTN2
G:G
G:T
T:T
B) Correlation plot: 164−abyss_C906_p3592
y'i fluorescence signal
M. trossulus fluorescence fraction 2 (MtFF2, mean of 17 SNPs)
L11_4M
L9_3H
CamaretB_1H
Pal30B
L9_3M
L11_4H
STBRI_69H
L19_8H
Pal30H
L19_8M
T:T
G:T
G:G
M. edulis
M. trossulus
MtrBTN2
MtrBTN2 host (M. edulis)

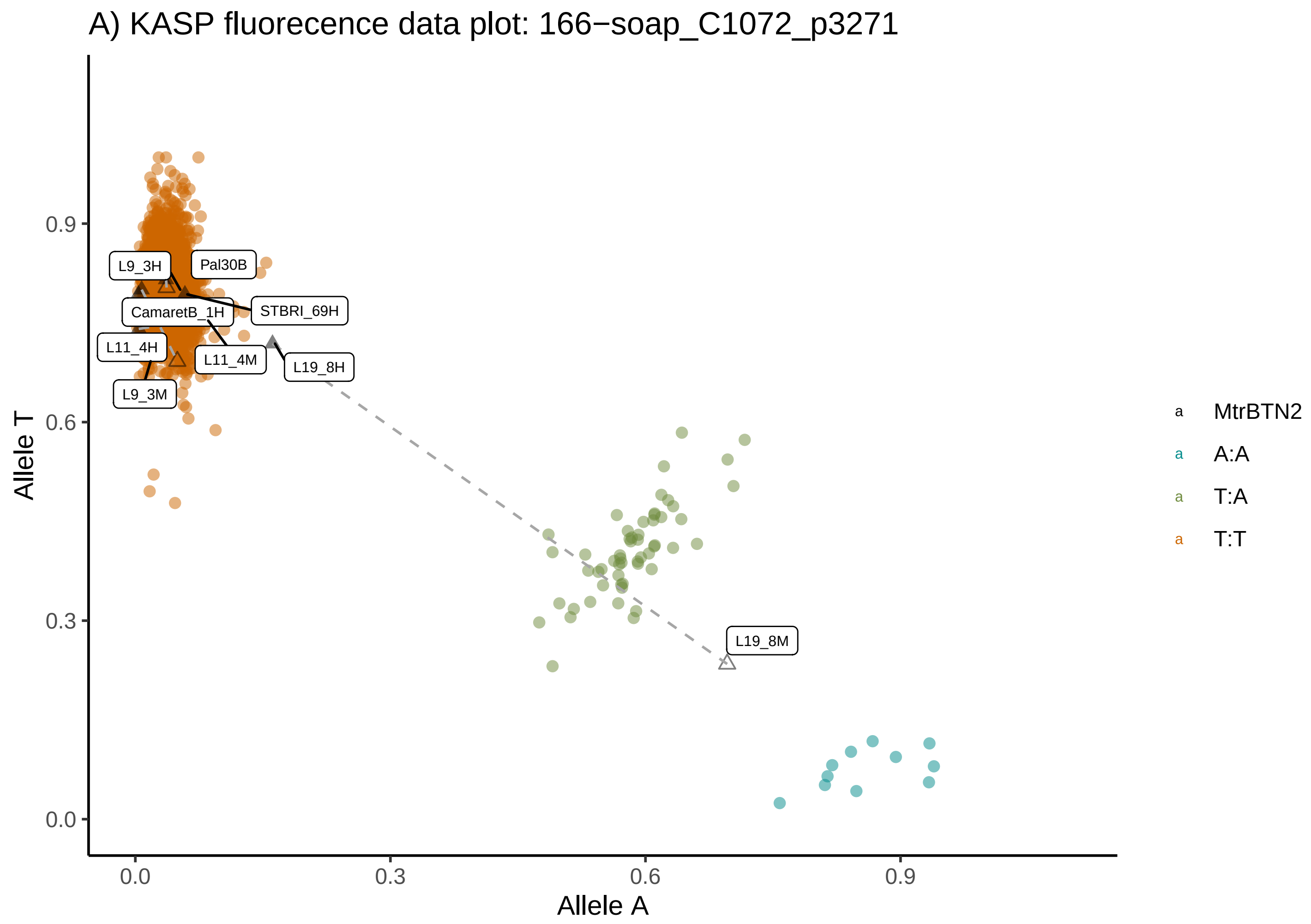
A) KASP fluorecence data plot: 166−soap_C1072_p3271
Allele T
Allele A
0.0
0.3
0.6
0.9
L9_3H
Pal30B
CamaretB_1H
STBRI_69H
L11_4H
L11_4M
L19_8H
L9_3M
L19_8M
MtrBTN2
A:A
T:A
T:T

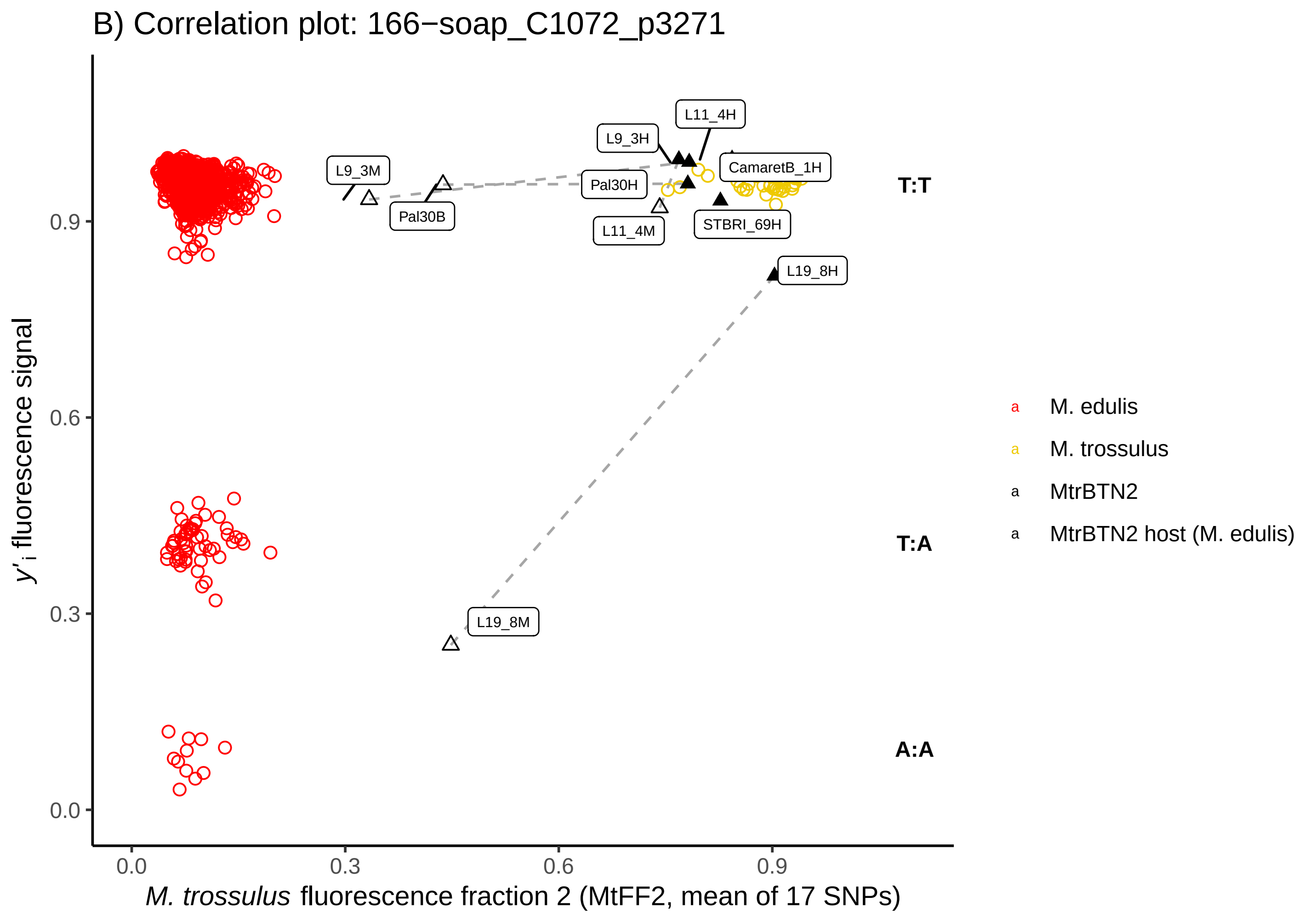
B) Correlation plot: 166−soap_C1072_p3271
y'i fluorescence signal
M. trossulus fluorescence fraction 2 (MtFF2, mean of 17 SNPs)
0.0
0.3
0.6
0.9
L11_4H
L9_3H
L9_3M
CamaretB_1H
Pal30H
Pal30B
L11_4M
STBRI_69H
L19_8H
L19_8M
T:T
T:A
A:A
M. edulis
M. trossulus
MtrBTN2
MtrBTN2 host (M. edulis)

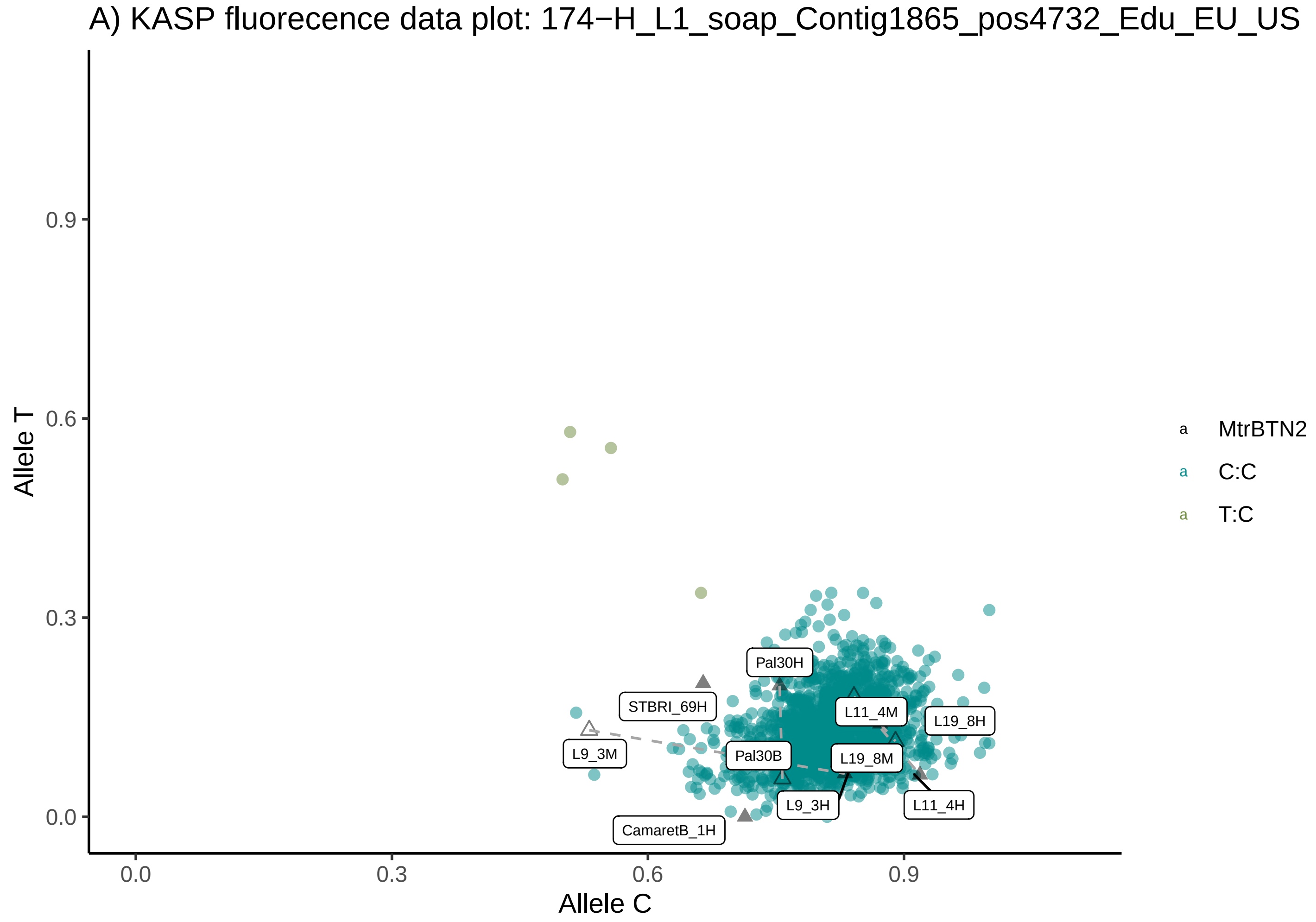
A) KASP fluorecence data plot: 174−H_L1_soap_Contig1865_pos4732_Edu_EU_US
Allele T
Allele C
0.0
0.3
0.6
0.9
Pal30H
STBRI_69H
L11_4M
L19_8H
L9_3M
Pal30B
L19_8M
L9_3H
L11_4H
CamaretB_1H
MtrBTN2
C:C
T:C

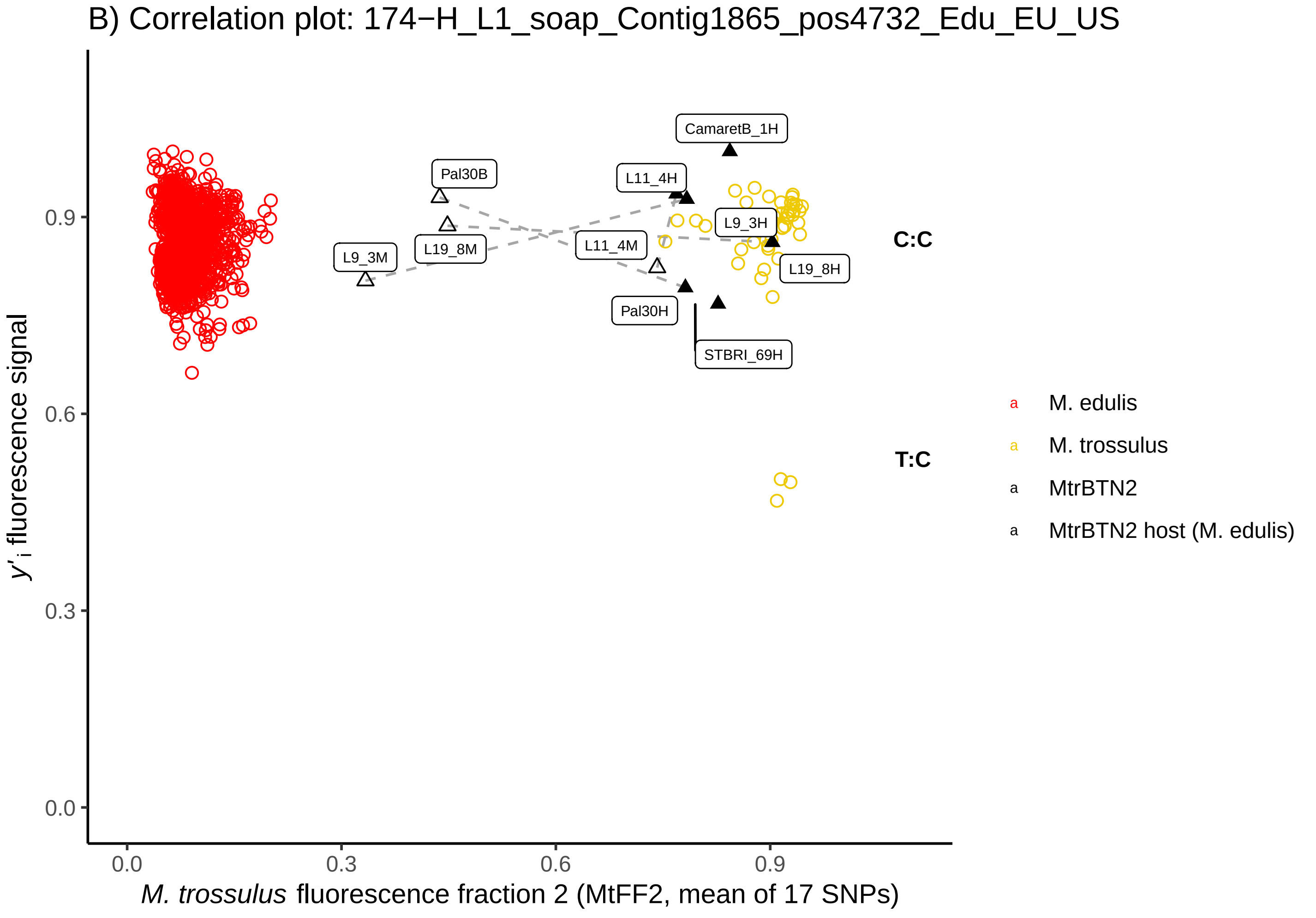
B) Correlation plot: 174−H_L1_soap_Contig1865_pos4732_Edu_EU_US
y'i fluorescence signal
M. trossulus fluorescence fraction 2 (MtFF2, mean of 17 SNPs)
0.0
0.3
0.6
0.9
CamaretB_1H
Pal30B
L11_4H
L9_3H
L11_4M
L9_3M
L19_8M
L19_8H
Pal30H
STBRI_69H
C:C
T:C
M. edulis
M. trossulus
MtrBTN2
MtrBTN2 host (M. edulis)

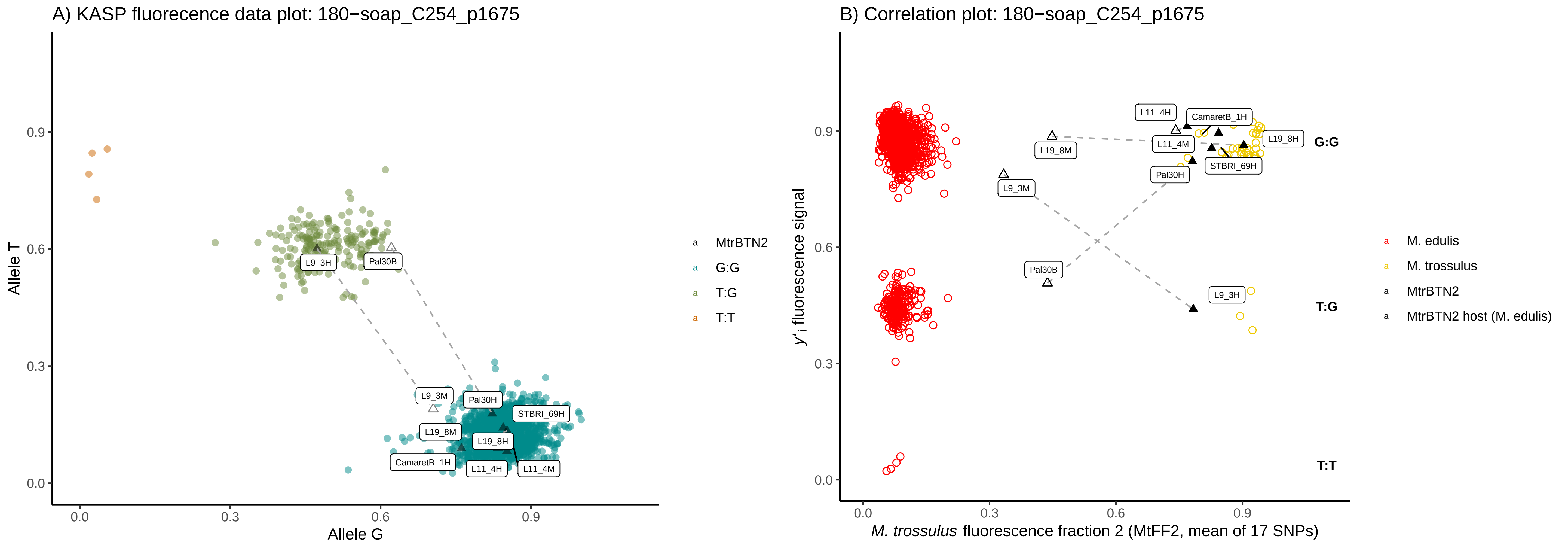
A) KASP fluorecence data plot: 180−soap_C254_p1675
Allele T
Allele G
0.0
0.3
0.6
0.9
L9_3H
Pal30B
L9_3M
Pal30H
STBRI_69H
L19_8M
L19_8H
CamaretB_1H
L11_4H
L11_4M
MtrBTN2
G:G
T:G
T:T
B) Correlation plot: 180−soap_C254_p1675
y'i fluorescence signal
M. trossulus fluorescence fraction 2 (MtFF2, mean of 17 SNPs)
L11_4H
CamaretB_1H
L19_8M
L11_4M
L19_8H
Pal30H
STBRI_69H
L9_3M
Pal30B
L9_3H
G:G
T:G
T:T
M. edulis
M. trossulus
MtrBTN2
MtrBTN2 host (M. edulis)

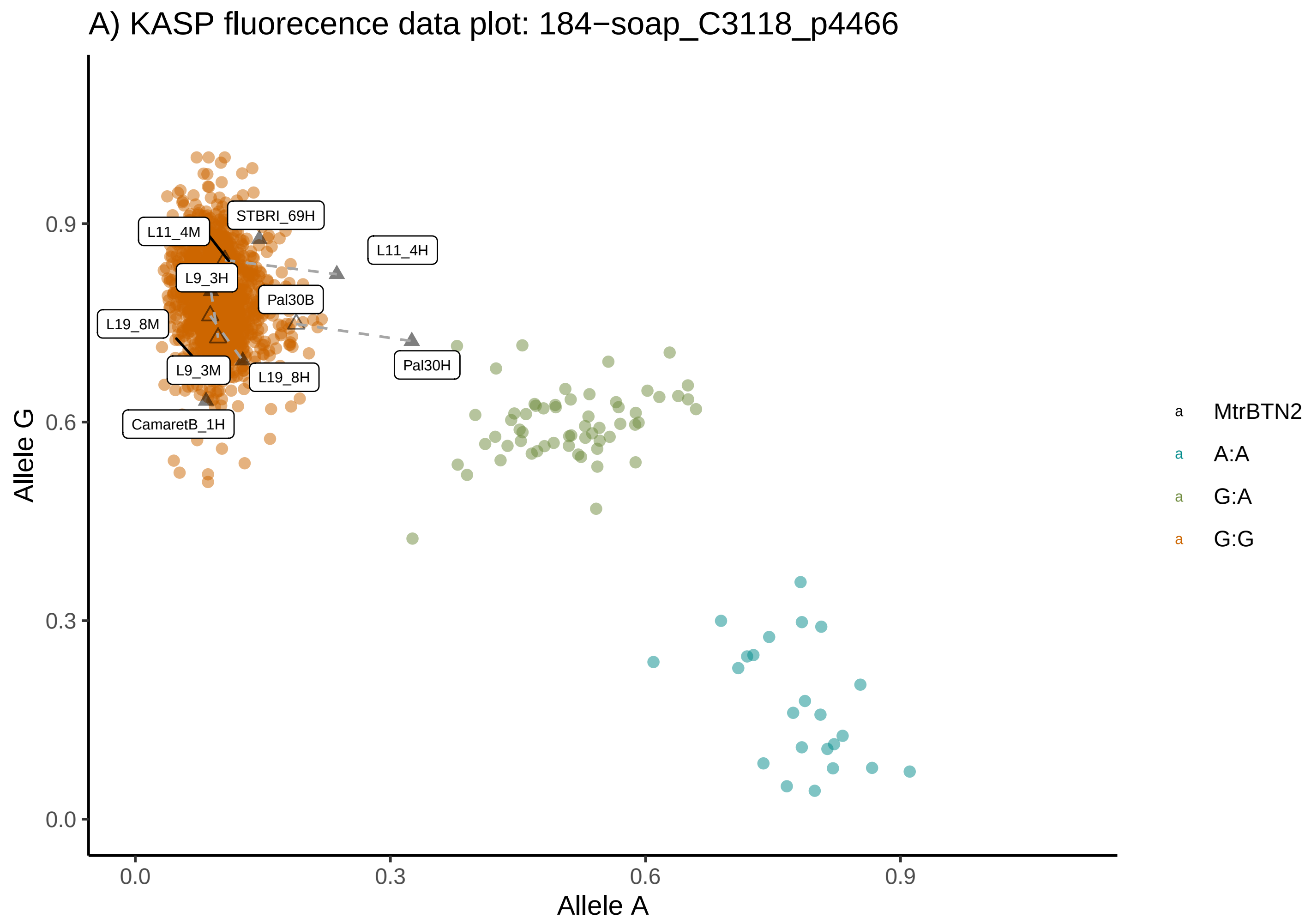
A) KASP fluorecence data plot: 184−soap_C3118_p4466
Allele G
Allele A
0.0
0.3
0.6
0.9
STBRI_69H
L11_4M
L11_4H
L9_3H
Pal30B
L19_8M
Pal30H
L9_3M
L19_8H
CamaretB_1H
MtrBTN2
A:A
G:A
G:G

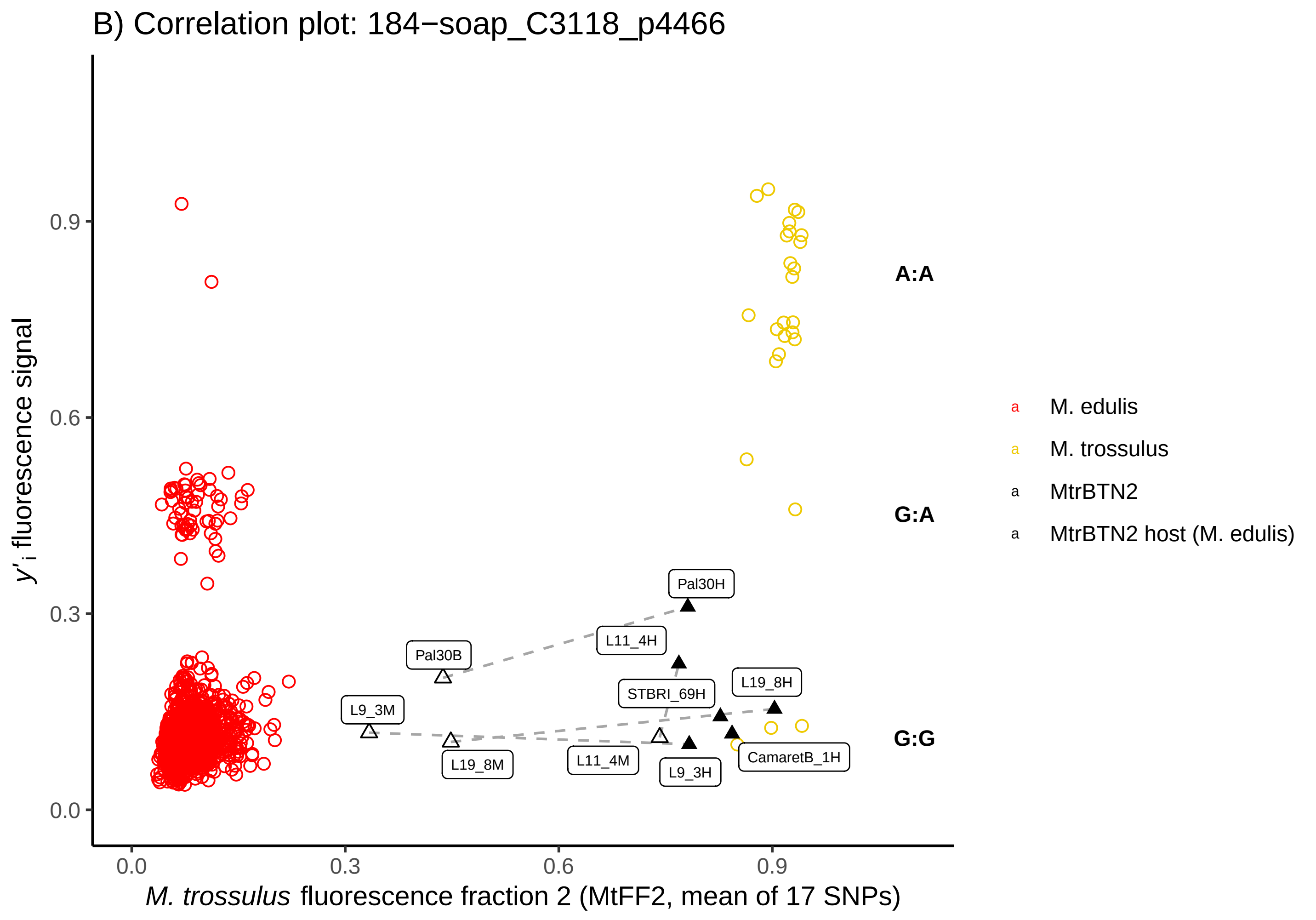
B) Correlation plot: 184−soap_C3118_p4466
y'i fluorescence signal
M. trossulus fluorescence fraction 2 (MtFF2, mean of 17 SNPs)
0.0
0.3
0.6
0.9
A:A
G:A
G:G
Pal30H
L11_4H
Pal30B
STBRI_69H
L19_8H
L9_3M
L19_8M
L11_4M
L9_3H
CamaretB_1H
M. edulis
M. trossulus
MtrBTN2
MtrBTN2 host (M. edulis)

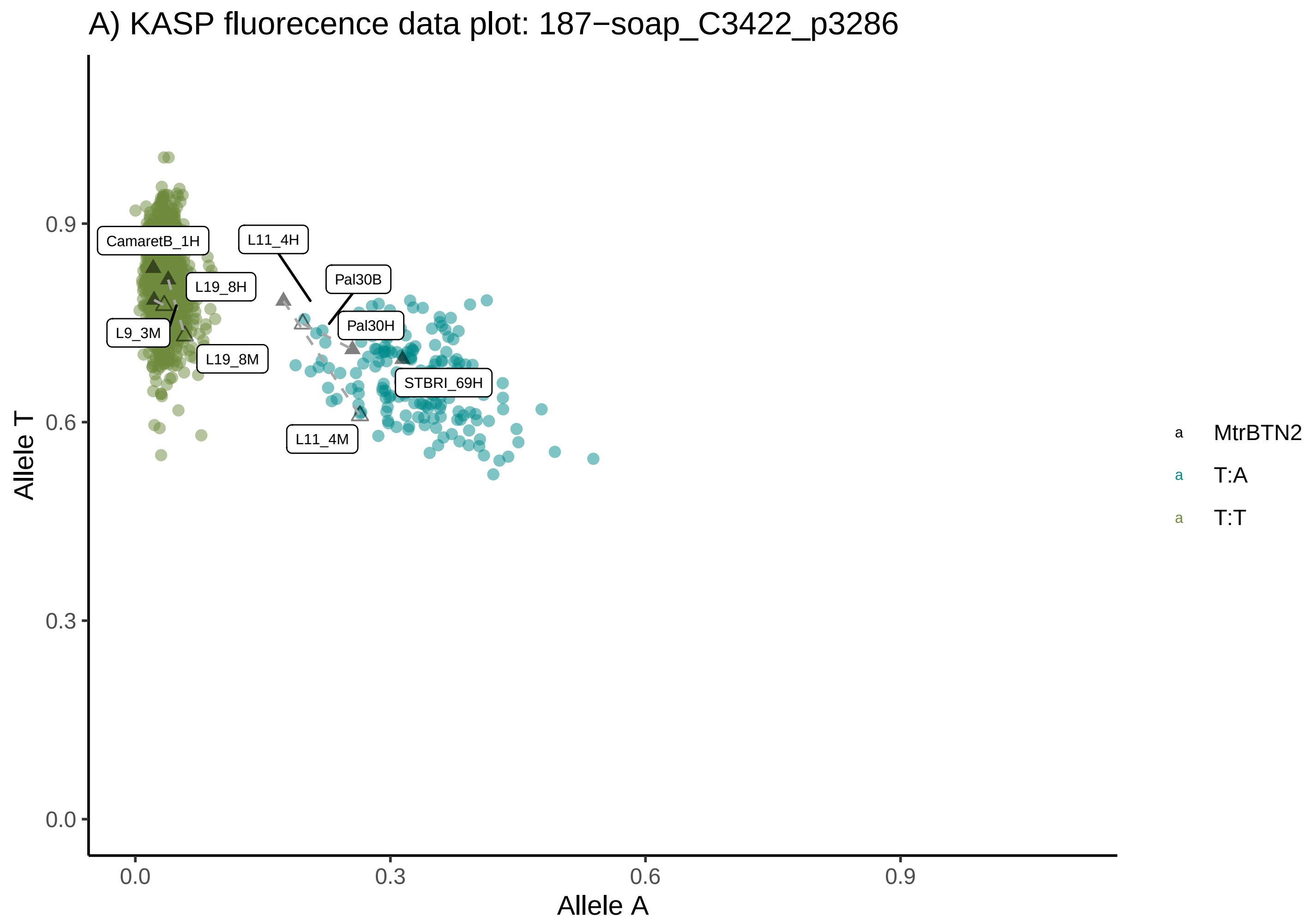
A) KASP fluorecence data plot: 187−soap_C3422_p3286
Allele T
Allele A
0.0
0.3
0.6
0.9
CamaretB_1H
L11_4H
Pal30B
L19_8H
Pal30H
L9_3M
L19_8M
STBRI_69H
L11_4M
MtrBTN2
T:A
T:T

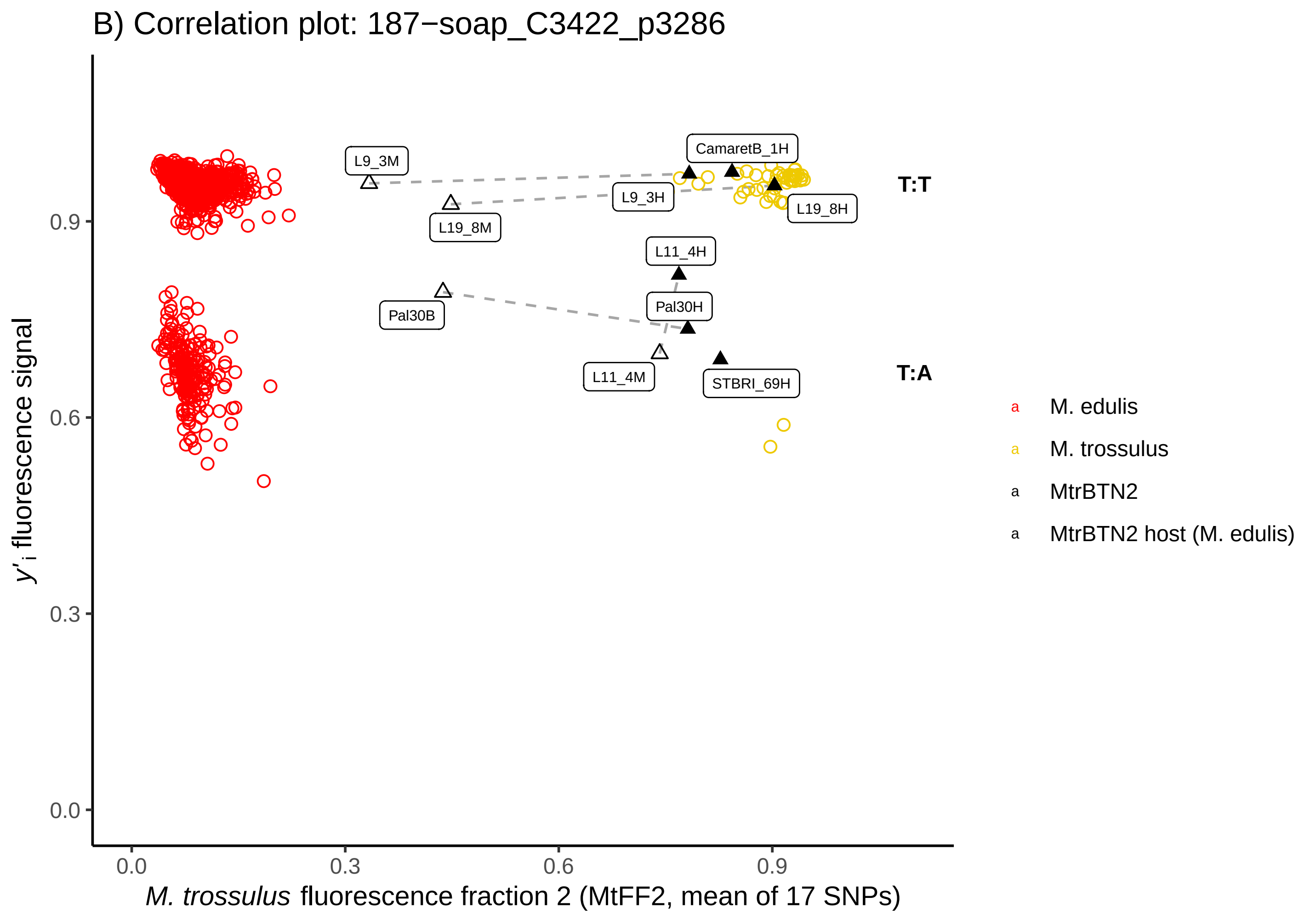
B) Correlation plot: 187−soap_C3422_p3286
y'i fluorescence signal
M. trossulus fluorescence fraction 2 (MtFF2, mean of 17 SNPs)
0.0
0.3
0.6
0.9
L9_3M
CamaretB_1H
L9_3H
L19_8H
L19_8M
L11_4H
Pal30H
Pal30B
L11_4M
STBRI_69H
T:T
T:A
M. edulis
M. trossulus
MtrBTN2
MtrBTN2 host (M. edulis)

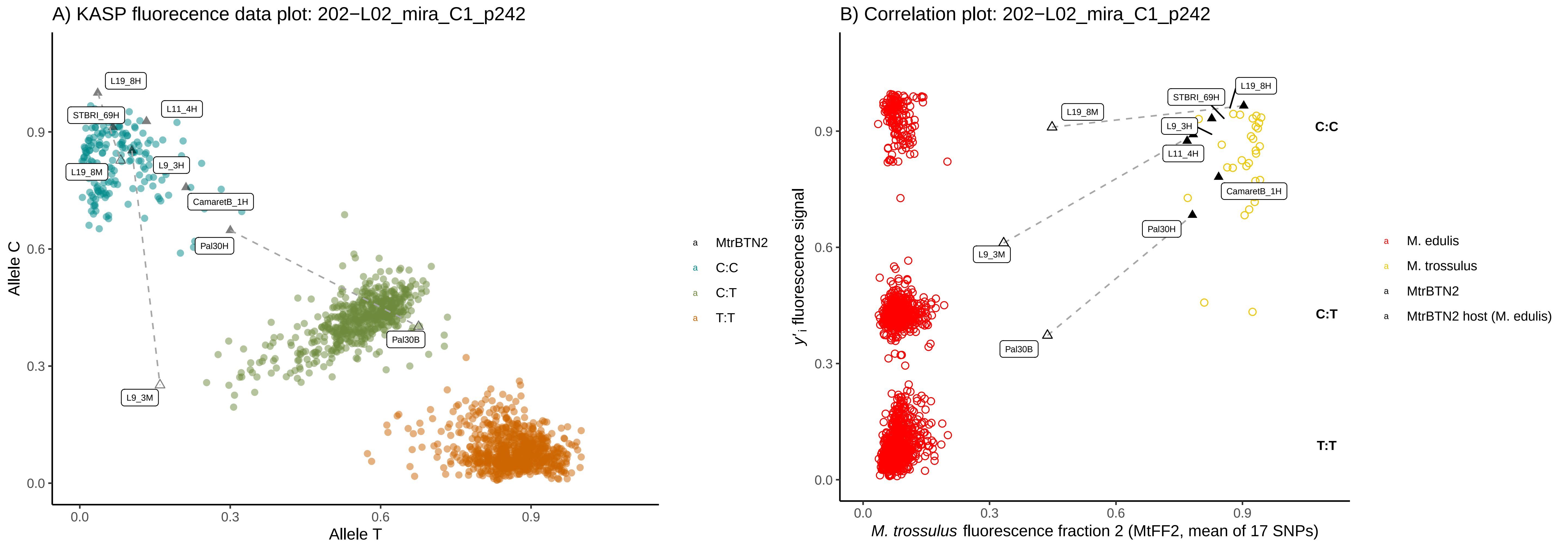
A) KASP fluorecence data plot: 202−L02_mira_C1_p242
Allele C
Allele T
L19_8H
L11_4H
STBRI_69H
L9_3H
L19_8M
CamaretB_1H
Pal30H
Pal30B
L9_3M
MtrBTN2
C:C
C:T
T:T
B) Correlation plot: 202−L02_mira_C1_p242
y'i fluorescence signal
M. trossulus fluorescence fraction 2 (MtFF2, mean of 17 SNPs)
M. edulis
M. trossulus
MtrBTN2
MtrBTN2 host (M. edulis)

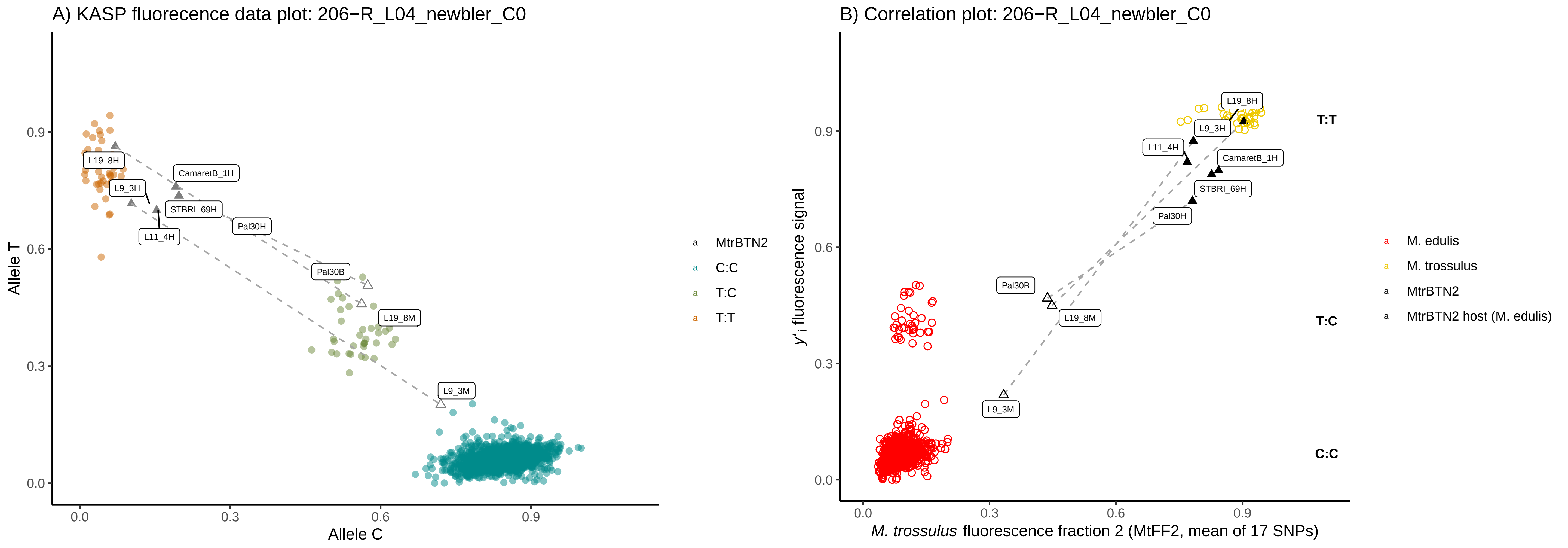
A) KASP fluorecence data plot: 206−R_L04_newbler_C0
Allele T
Allele C
0.0
0.3
0.6
0.9
L19_8H
CamaretB_1H
L9_3H
STBRI_69H
L11_4H
Pal30H
Pal30B
L19_8M
L9_3M
MtrBTN2
C:C
T:C
T:T
B) Correlation plot: 206−R_L04_newbler_C0
y'i fluorescence signal
M. trossulus fluorescence fraction 2 (MtFF2, mean of 17 SNPs)
T:T
T:C
C:C
M. edulis
M. trossulus
MtrBTN2
MtrBTN2 host (M. edulis)

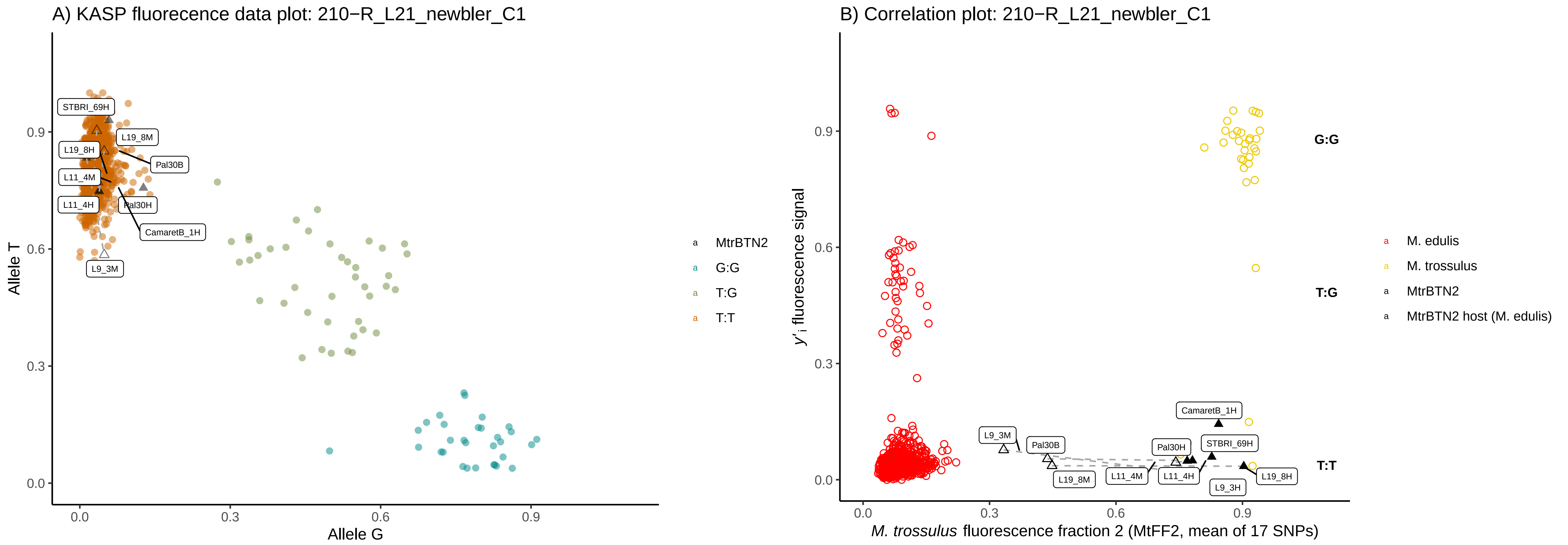
A) KASP fluorecence data plot: 210−R_L21_newbler_C1
Allele T
Allele G
STBRI_69H
L19_8M
L19_8H
Pal30B
L11_4M
L11_4H
Pal30H
CamaretB_1H
L9_3M
MtrBTN2
G:G
T:G
T:T
B) Correlation plot: 210−R_L21_newbler_C1
y'i fluorescence signal
M. trossulus fluorescence fraction 2 (MtFF2, mean of 17 SNPs)
G:G
T:G
T:T
CamaretB_1H
L9_3M
Pal30B
Pal30H
STBRI_69H
L19_8M
L11_4M
L11_4H
L9_3H
L19_8H
M. edulis
M. trossulus
MtrBTN2
MtrBTN2 host (M. edulis)

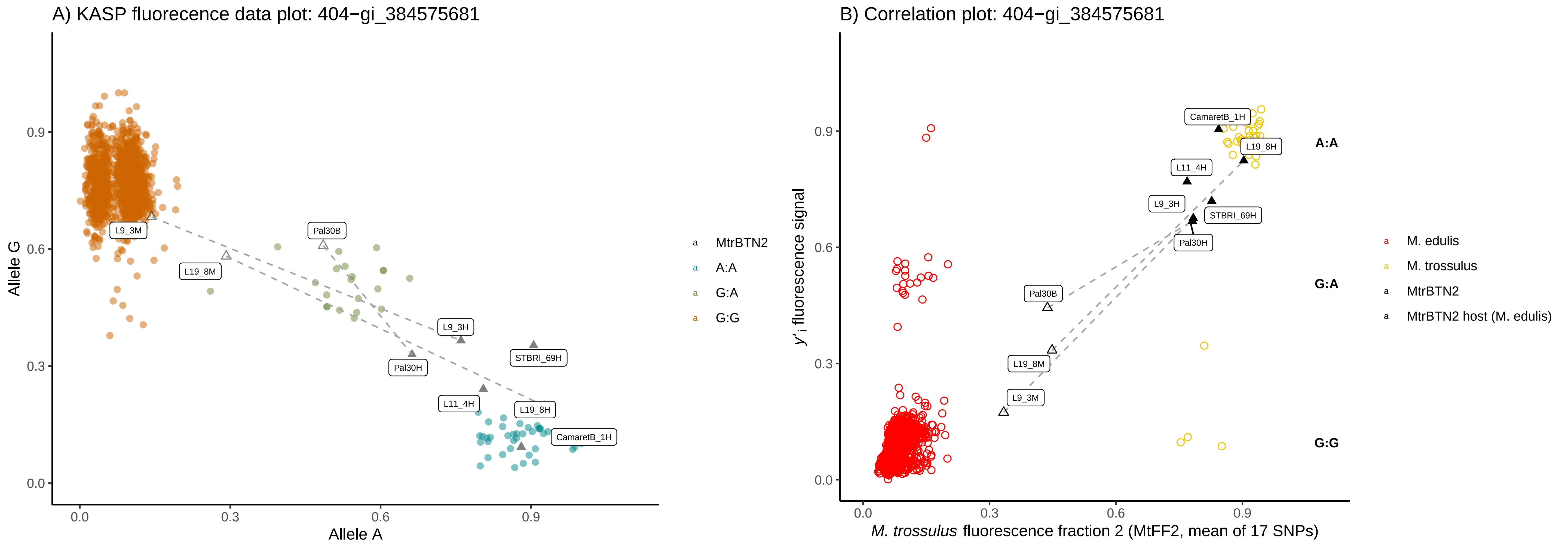
A) KASP fluorecence data plot: 404−gi_384575681
B) Correlation plot: 404−gi_384575681
Allele G
Allele A
0.0
0.3
0.6
0.9
L9_3M
L19_8M
Pal30B
L9_3H
Pal30H
STBRI_69H
L11_4H
L19_8H
CamaretB_1H
MtrBTN2
A:A
G:A
G:G
y'i fluorescence signal
M. trossulus fluorescence fraction 2 (MtFF2, mean of 17 SNPs)
M. edulis
M. trossulus
MtrBTN2 host (M. edulis)

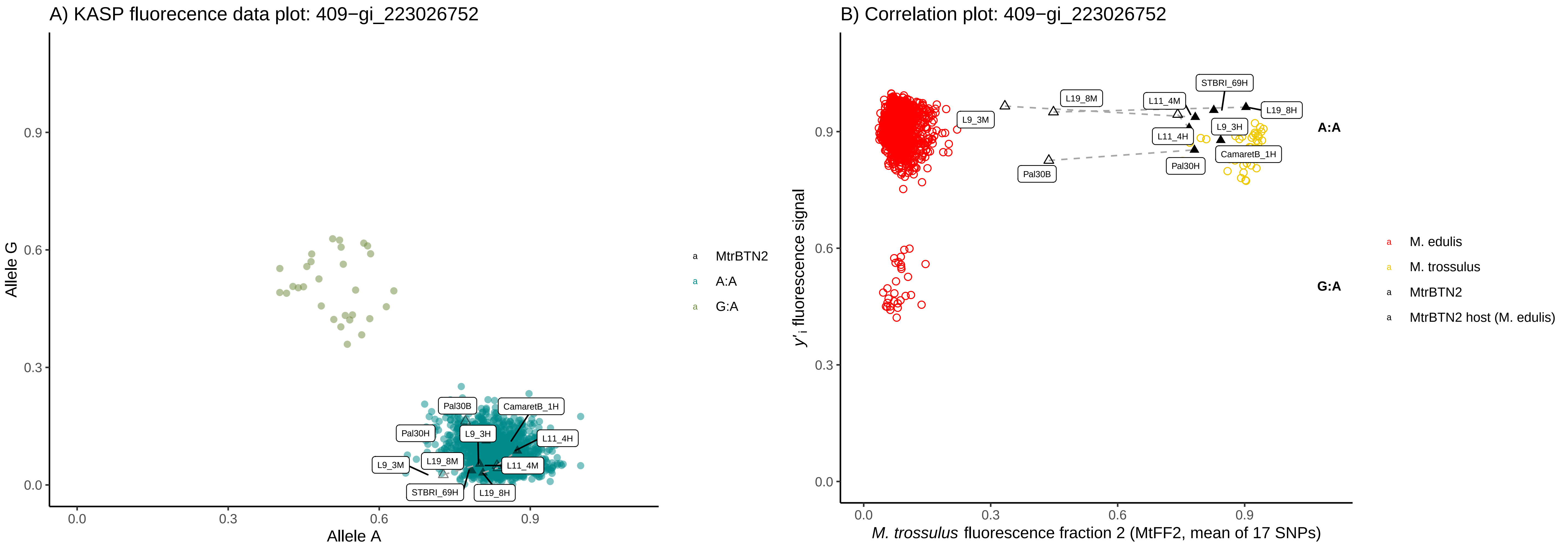
A) KASP fluorecence data plot: 409−gi_223026752
Allele G
Allele A
0.0
0.3
0.6
0.9
Pal30B
CamaretB_1H
Pal30H
L9_3H
L11_4H
L9_3M
L19_8M
L11_4M
STBRI_69H
L19_8H
MtrBTN2
A:A
G:A
B) Correlation plot: 409−gi_223026752
$y'_i$ fluorescence signal
*M. trossulus* fluorescence fraction 2 (MtFF2, mean of 17 SNPs)
STBRI_69H
L19_8M
L11_4M
L19_8H
L9_3M
L9_3H
L11_4H
CamaretB_1H
Pal30H
Pal30B
A:A
G:A
M. edulis
M. trossulus
MtrBTN2
MtrBTN2 host (M. edulis)

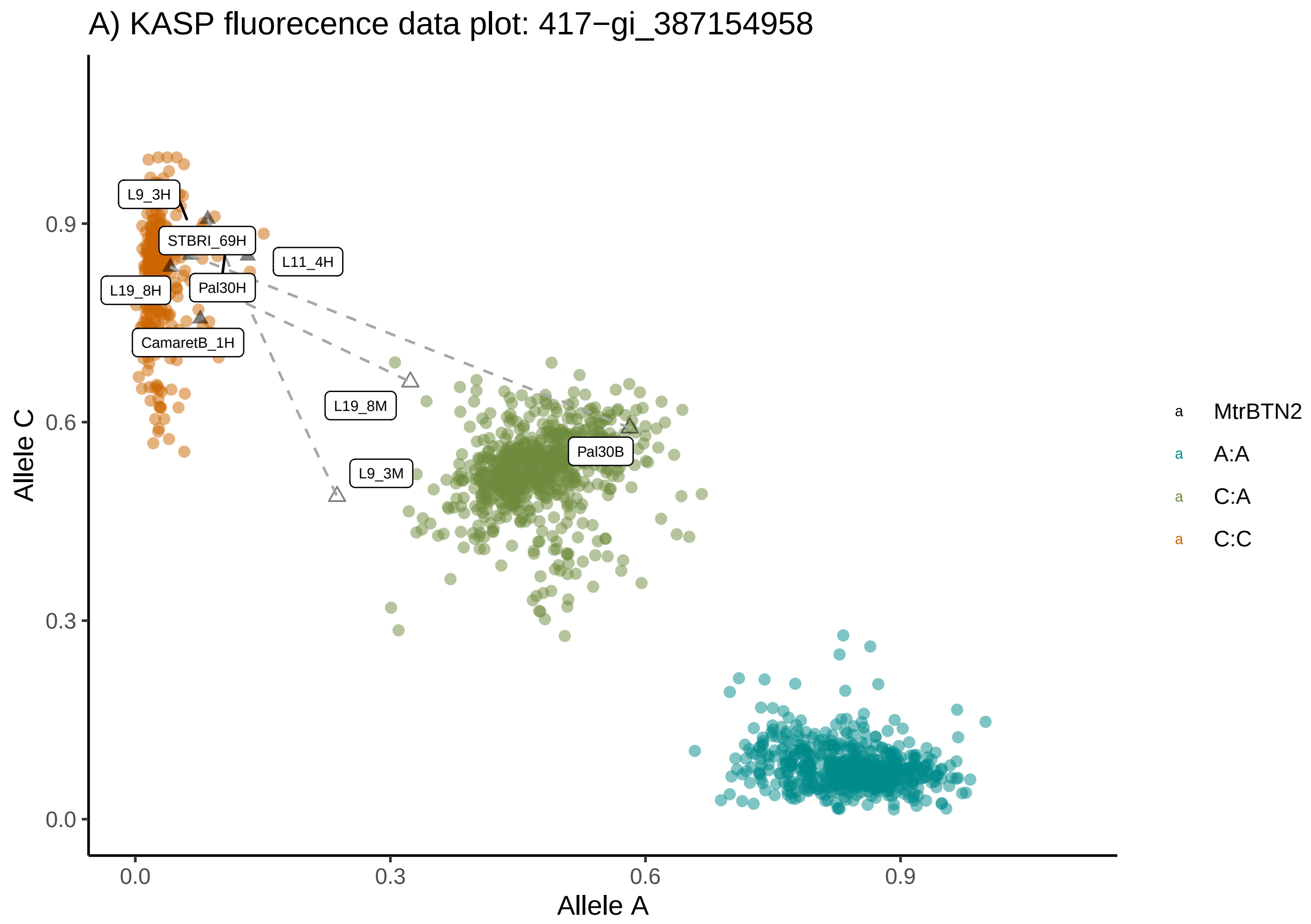
A) KASP fluorecence data plot: 417−gi_387154958
Allele C
Allele A
0.0
0.3
0.6
0.9
L9_3H
STBRI_69H
L11_4H
L19_8H
Pal30H
CamaretB_1H
L19_8M
L9_3M
Pal30B
MtrBTN2
A:A
C:A
C:C

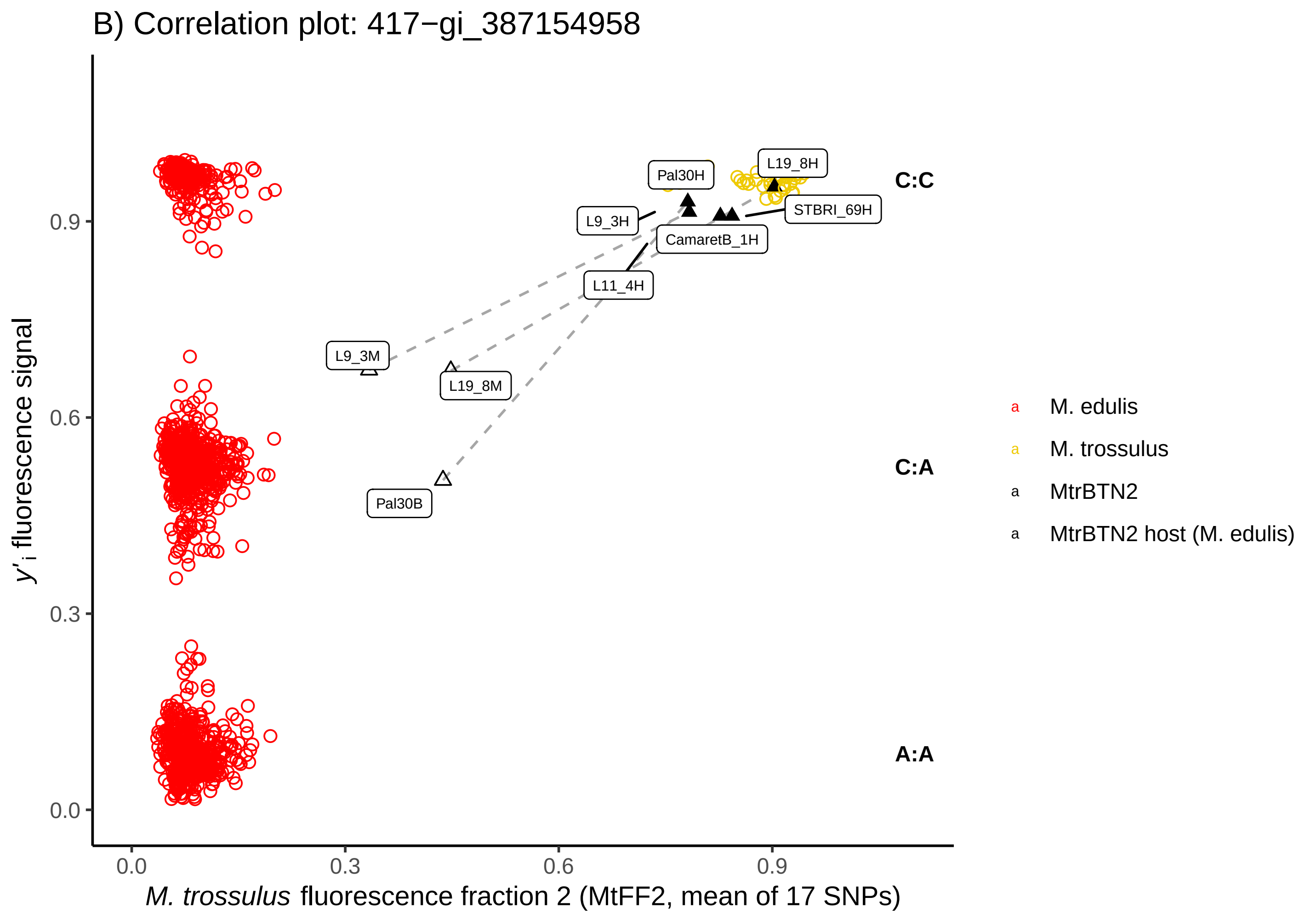
B) Correlation plot: 417−gi_387154958
y'i fluorescence signal
M. trossulus fluorescence fraction 2 (MtFF2, mean of 17 SNPs)
0.0
0.3
0.6
0.9
Pal30H
L19_8H
L9_3H
STBRI_69H
CamaretB_1H
L11_4H
L9_3M
L19_8M
Pal30B
C:C
C:A
A:A
M. edulis
M. trossulus
MtrBTN2
MtrBTN2 host (M. edulis)

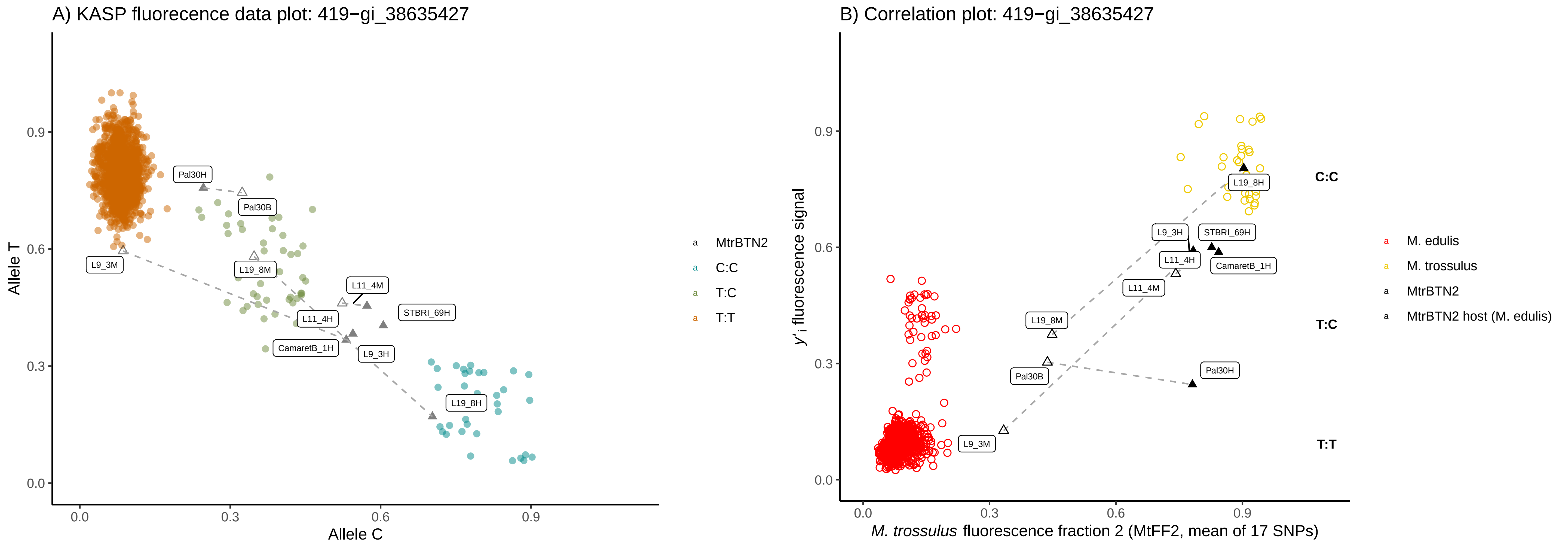

A) KASP fluorecence data plot: 419−gi_38635427
Allele T
Allele C
Pal30H
Pal30B
L9_3M
L19_8M
L11_4M
L11_4H
STBRI_69H
CamaretB_1H
L9_3H
L19_8H
MtrBTN2
C:C
T:C
T:T
B) Correlation plot: 419−gi_38635427
y'i fluorescence signal
M. trossulus fluorescence fraction 2 (MtFF2, mean of 17 SNPs)
C:C
T:C
T:T
M. edulis
M. trossulus
MtrBTN2
MtrBTN2 host (M. edulis)

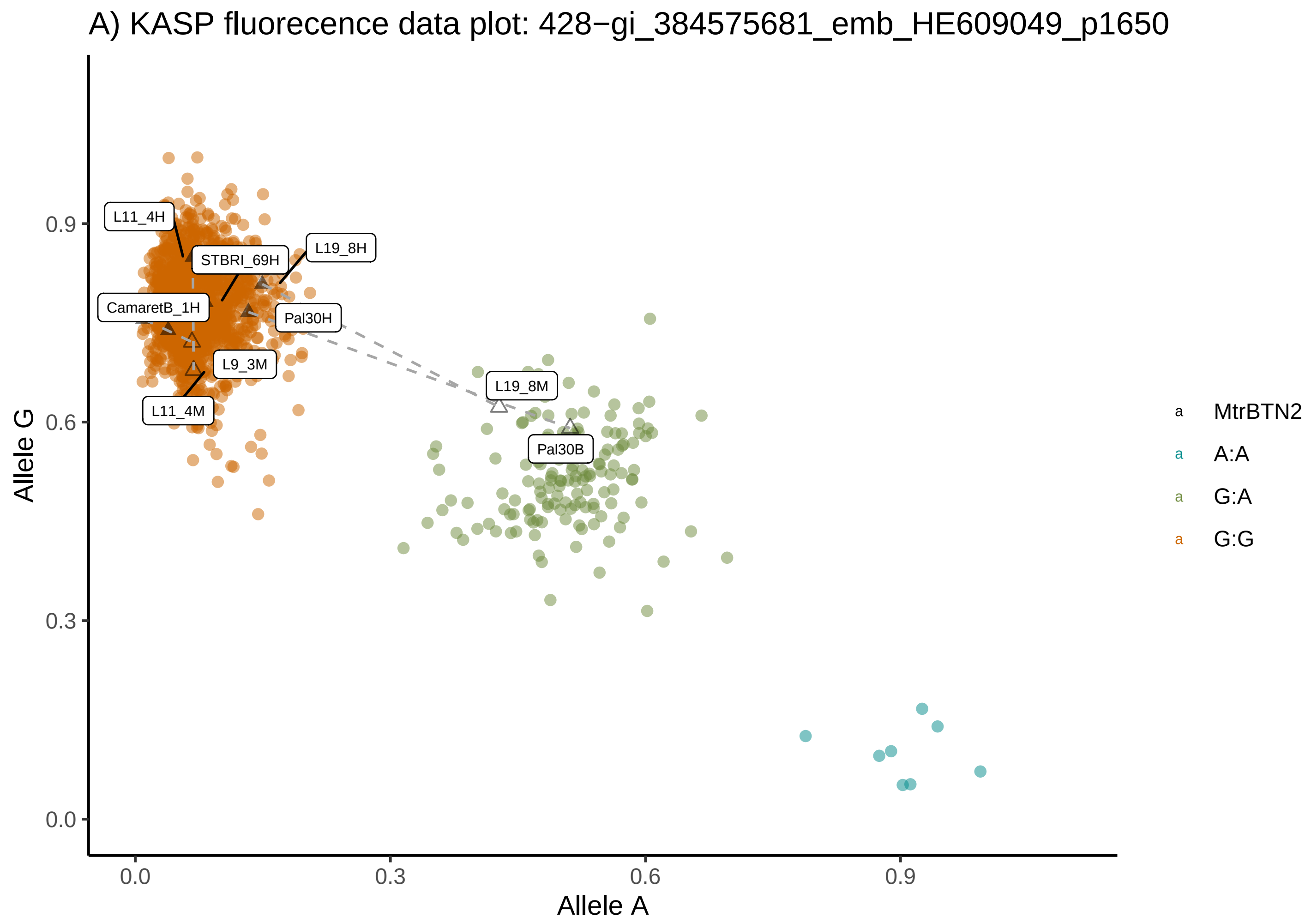
A) KASP fluorecence data plot: 428−gi_384575681_emb_HE609049_p1650
Allele G
Allele A
0.0
0.3
0.6
0.9
L11_4H
STBRI_69H
L19_8H
CamaretB_1H
Pal30H
L9_3M
L11_4M
L19_8M
Pal30B
MtrBTN2
A:A
G:A
G:G

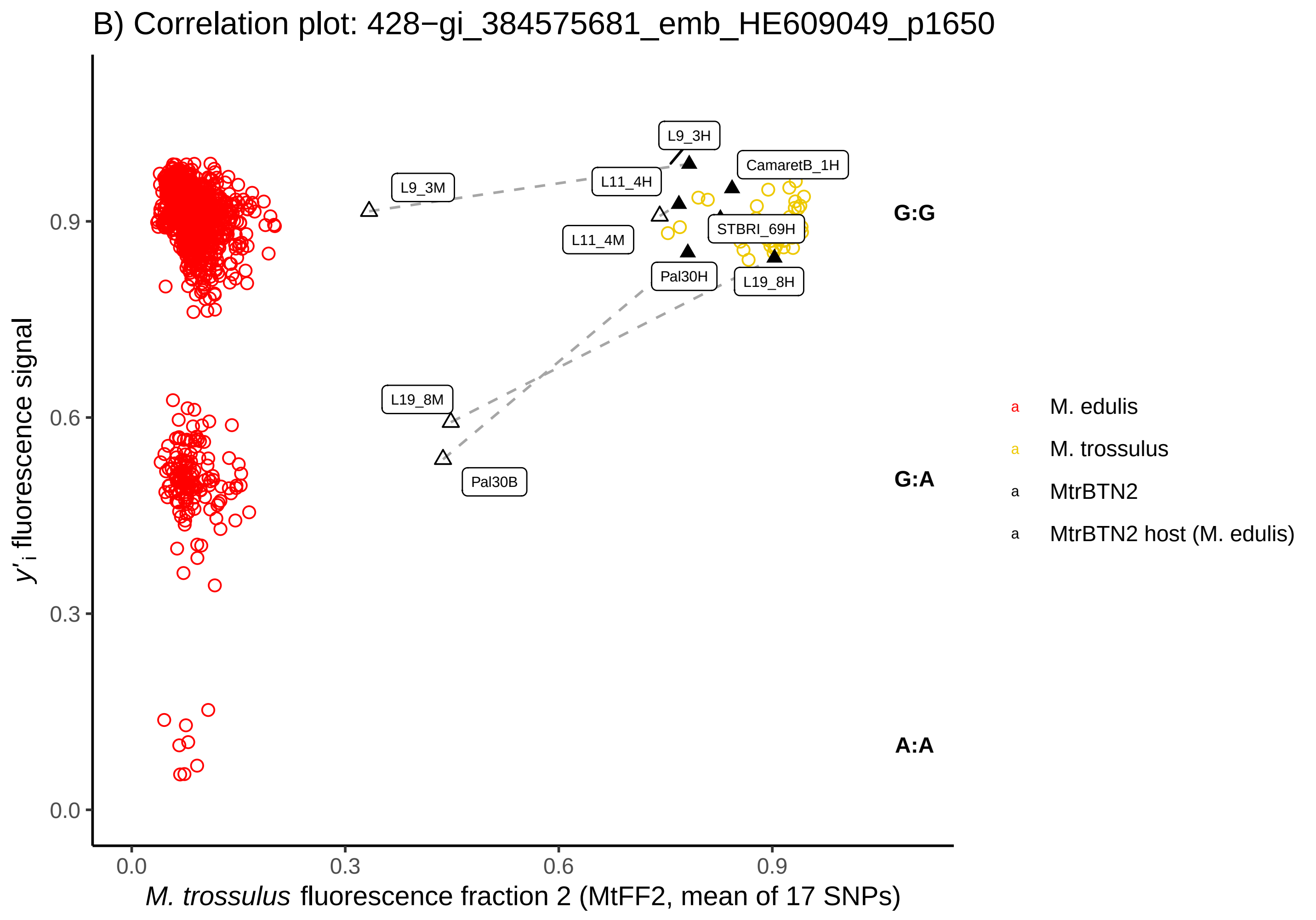
B) Correlation plot: 428−gi_384575681_emb_HE609049_p1650
y'i fluorescence signal
M. trossulus fluorescence fraction 2 (MtFF2, mean of 17 SNPs)
0.0
0.3
0.6
0.9
L9_3H
L11_4H
CamaretB_1H
L9_3M
L11_4M
STBRI_69H
Pal30H
L19_8H
L19_8M
Pal30B
G:G
G:A
A:A
M. edulis
M. trossulus
MtrBTN2
MtrBTN2 host (M. edulis)

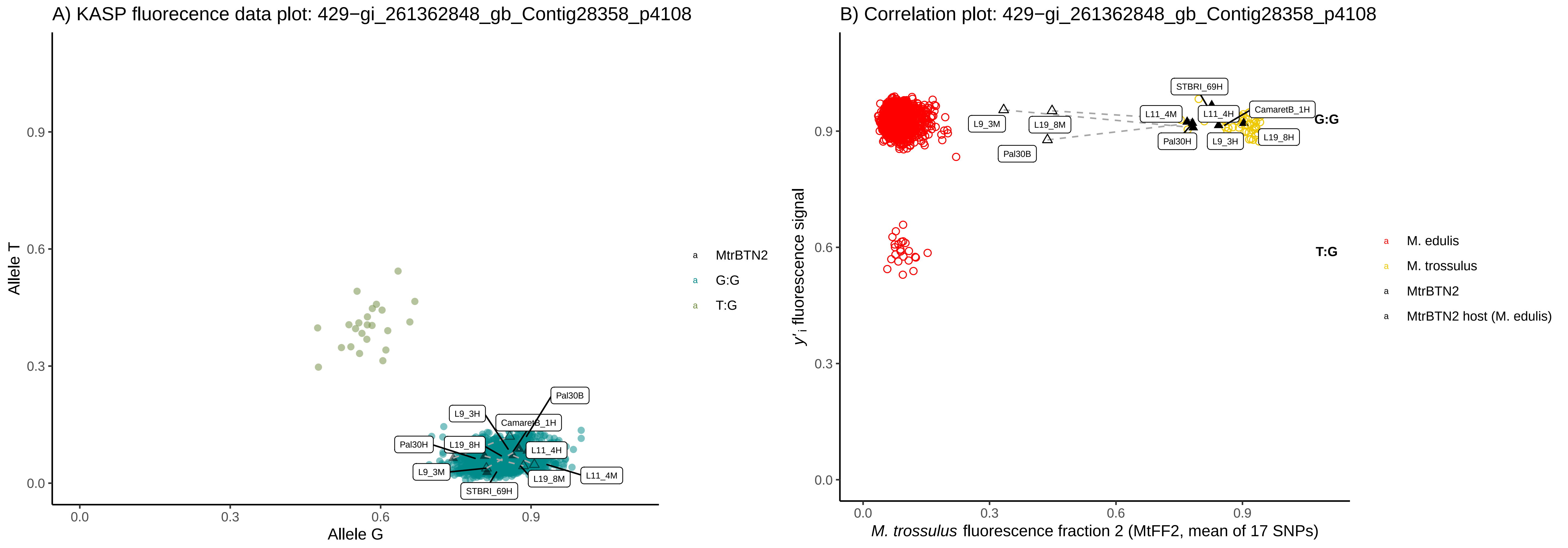
A) KASP fluorecence data plot: 429−gi_261362848_gb_Contig28358_p4108
Allele T
Allele G
0.0
0.3
0.6
0.9
Pal30B
L9_3H
CamaretB_1H
Pal30H
L19_8H
L11_4H
L9_3M
L19_8M
L11_4M
STBRI_69H
MtrBTN2
G:G
T:G
B) Correlation plot: 429−gi_261362848_gb_Contig28358_p4108
y'i fluorescence signal
M. trossulus fluorescence fraction 2 (MtFF2, mean of 17 SNPs)
STBRI_69H
L11_4M
L11_4H
CamaretB_1H
L9_3M
L19_8M
G:G
Pal30H
L9_3H
L19_8H
Pal30B
T:G
M. edulis
M. trossulus
MtrBTN2
MtrBTN2 host (M. edulis)

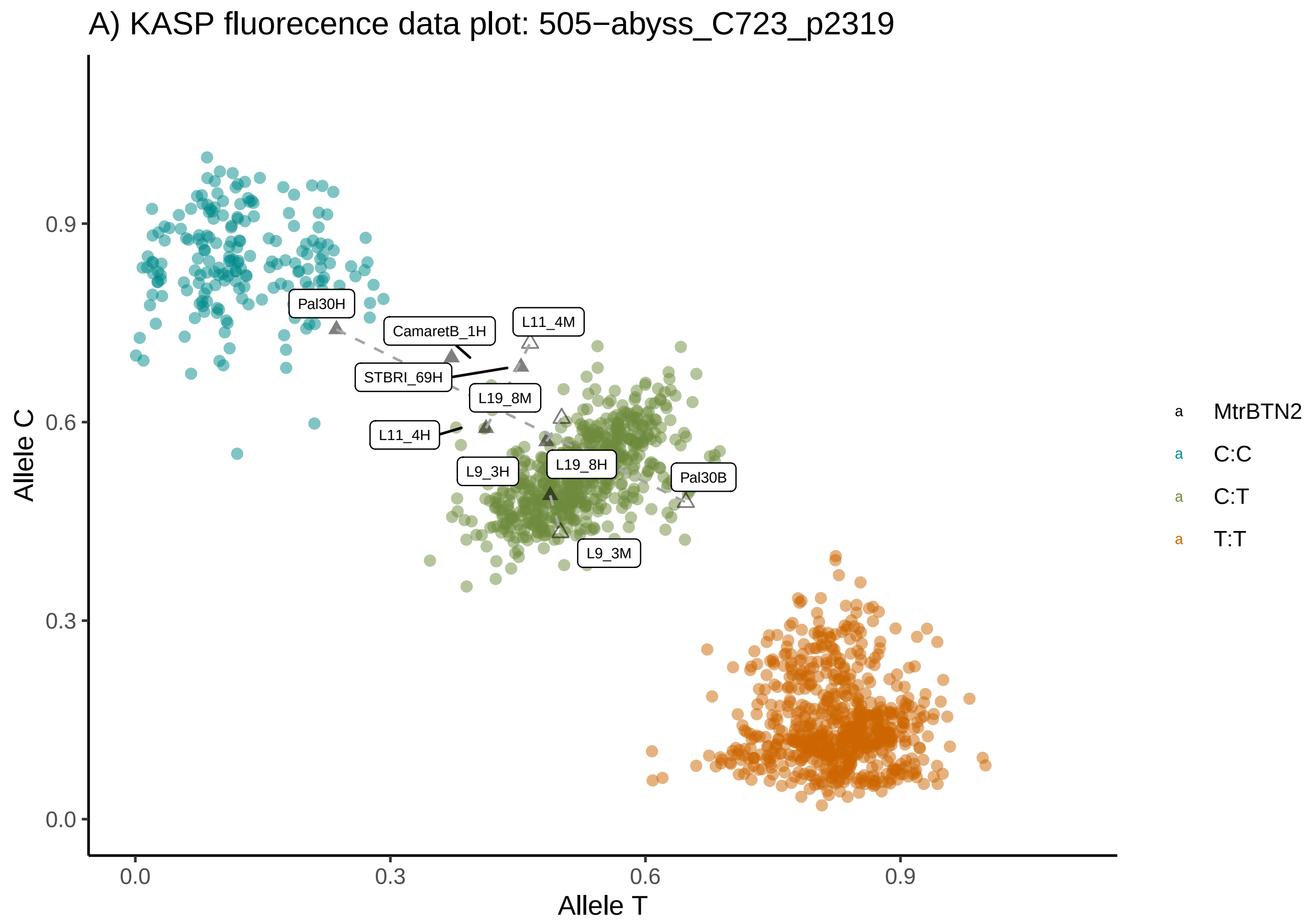
A) KASP fluorecence data plot: 505−abyss_C723_p2319
Allele C
Allele T
0.0
0.3
0.6
0.9
Pal30H
CamaretB_1H
L11_4M
STBRI_69H
L19_8M
L11_4H
L9_3H
L19_8H
Pal30B
L9_3M
MtrBTN2
C:C
C:T
T:T

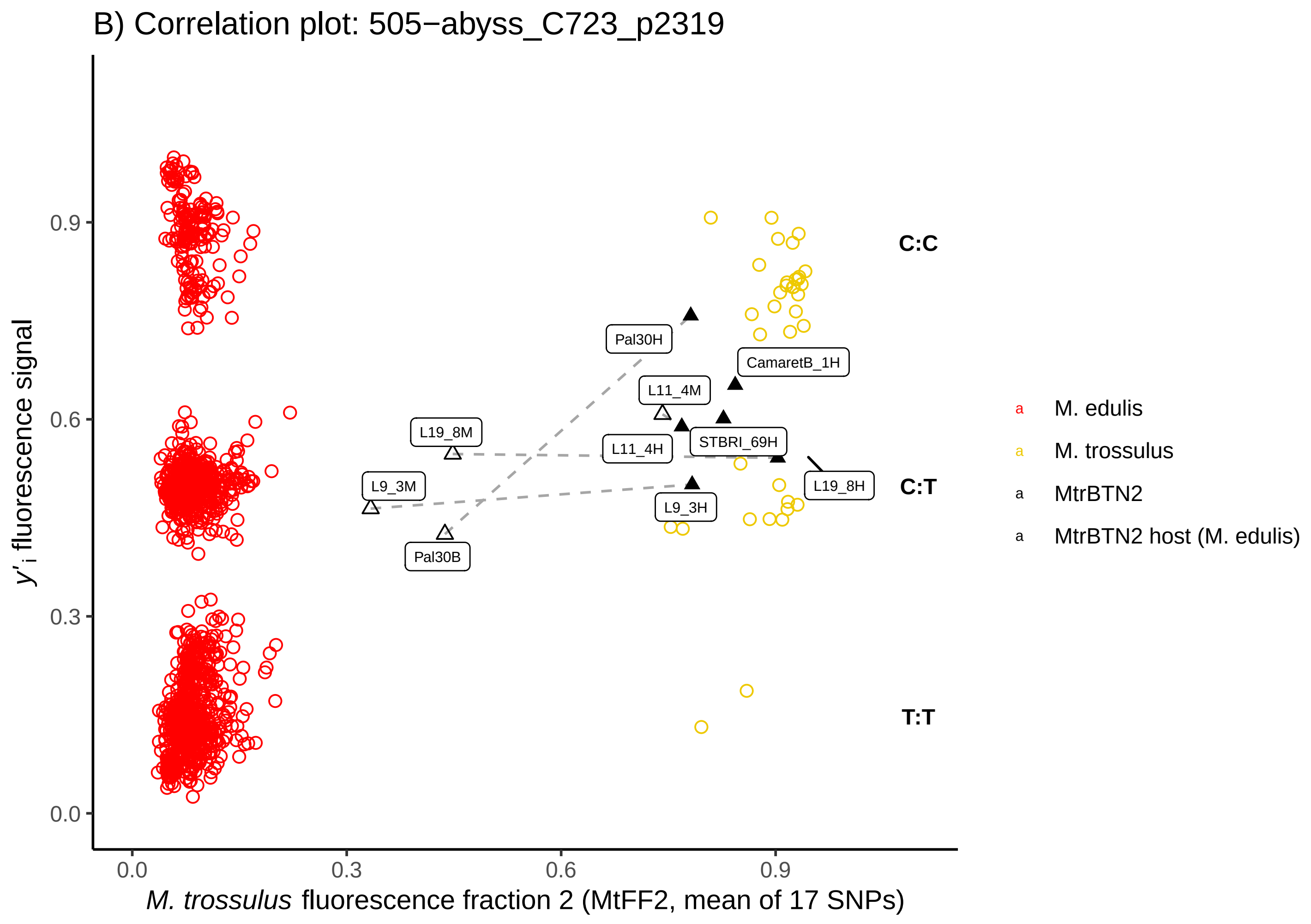
B) Correlation plot: 505−abyss_C723_p2319
y'i fluorescence signal
M. trossulus fluorescence fraction 2 (MtFF2, mean of 17 SNPs)
0.0
0.3
0.6
0.9
C:C
C:T
T:T
Pal30H
CamaretB_1H
L11_4M
L19_8M
L11_4H
STBRI_69H
L19_8H
L9_3M
L9_3H
Pal30B
M. edulis
M. trossulus
MtrBTN2
MtrBTN2 host (M. edulis)

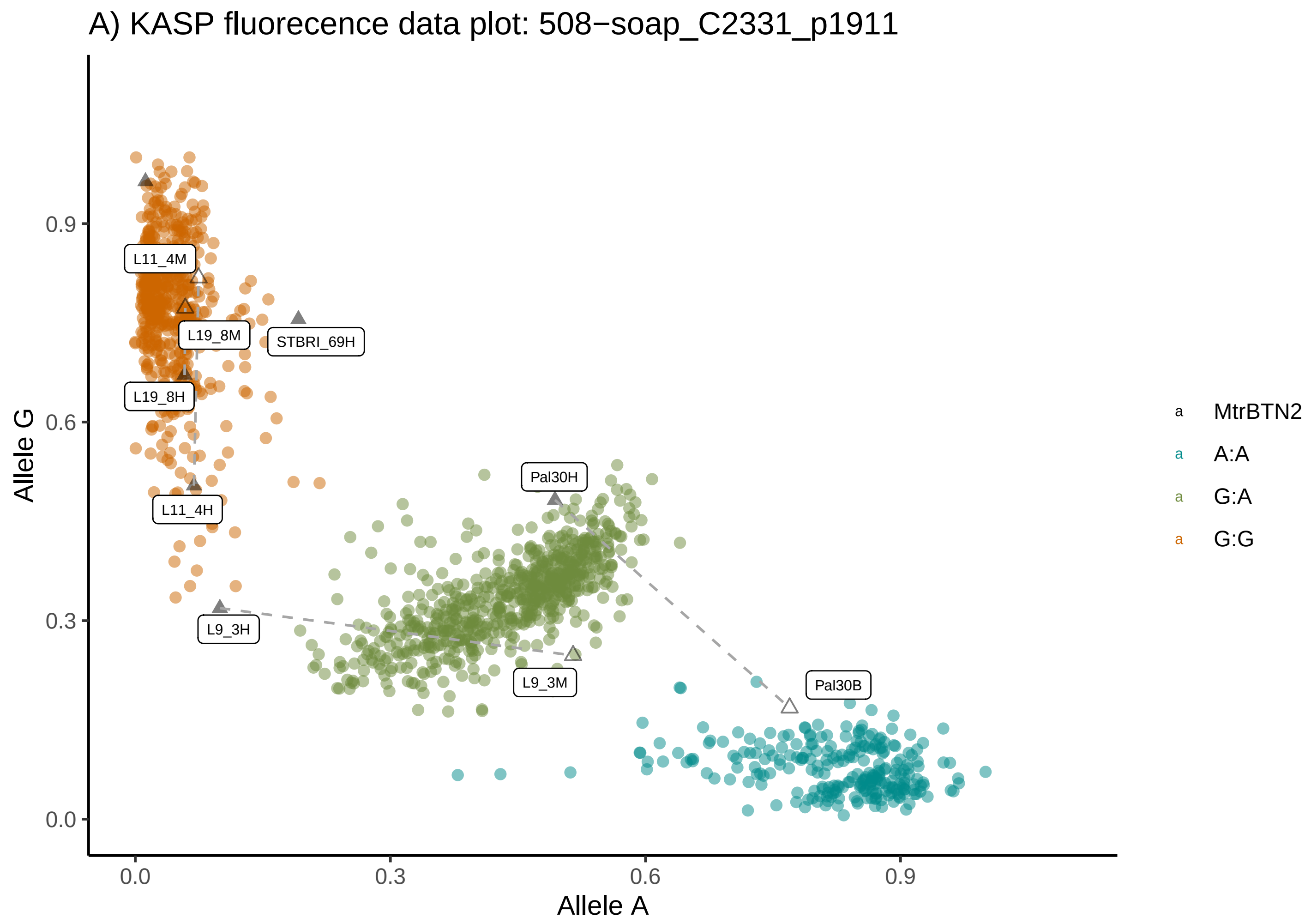
A) KASP fluorecence data plot: 508−soap_C2331_p1911
Allele G
Allele A
0.0
0.3
0.6
0.9
L11_4M
L19_8M
STBRI_69H
L19_8H
L11_4H
Pal30H
L9_3H
L9_3M
Pal30B
MtrBTN2
A:A
G:A
G:G

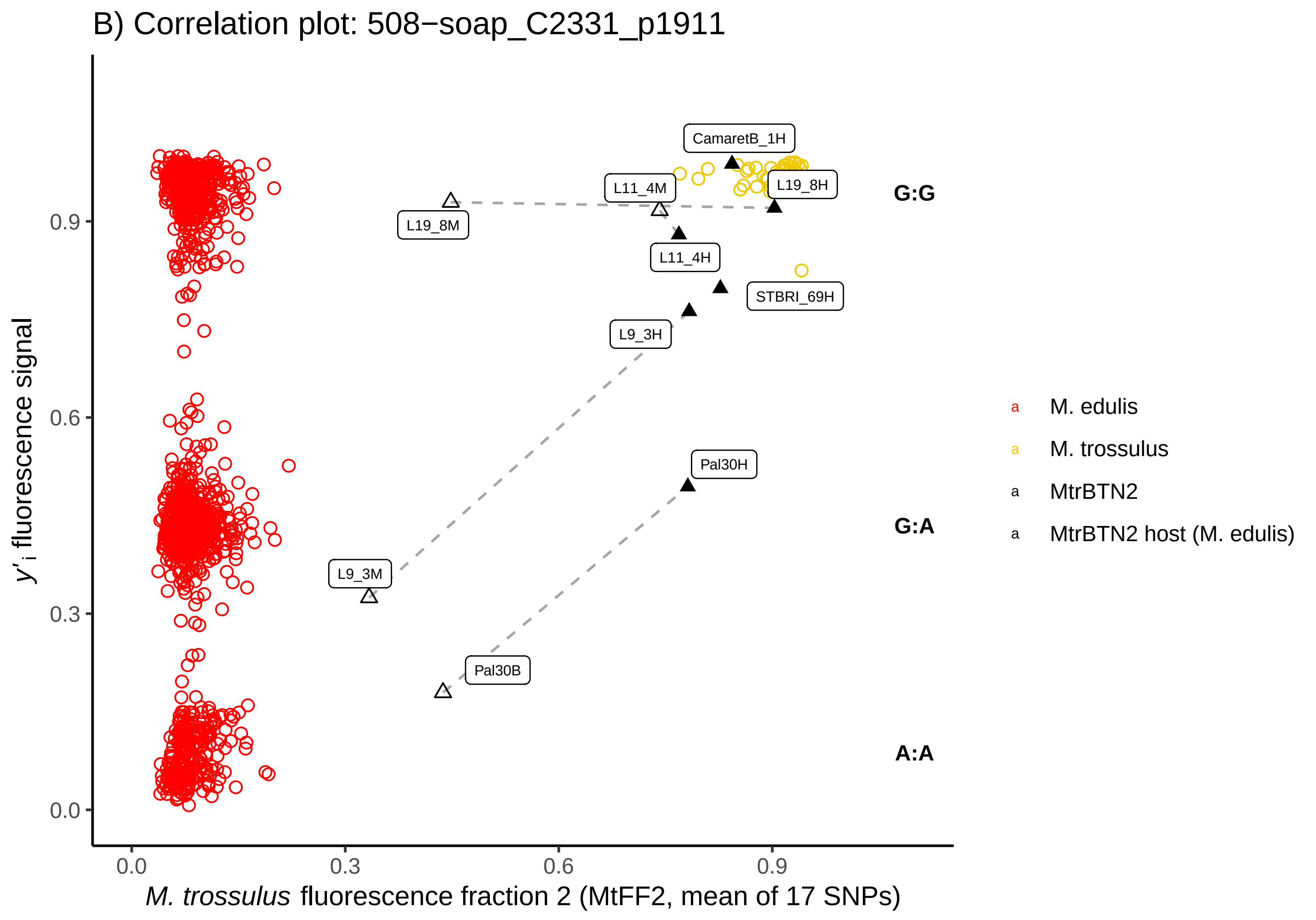
B) Correlation plot: 508−soap_C2331_p1911
y'i fluorescence signal
M. trossulus fluorescence fraction 2 (MtFF2, mean of 17 SNPs)
0.0
0.3
0.6
0.9
CamaretB_1H
L11_4M
L19_8H
L19_8M
L11_4H
STBRI_69H
L9_3H
Pal30H
L9_3M
Pal30B
G:G
G:A
A:A
M. edulis
M. trossulus
MtrBTN2
MtrBTN2 host (M. edulis)

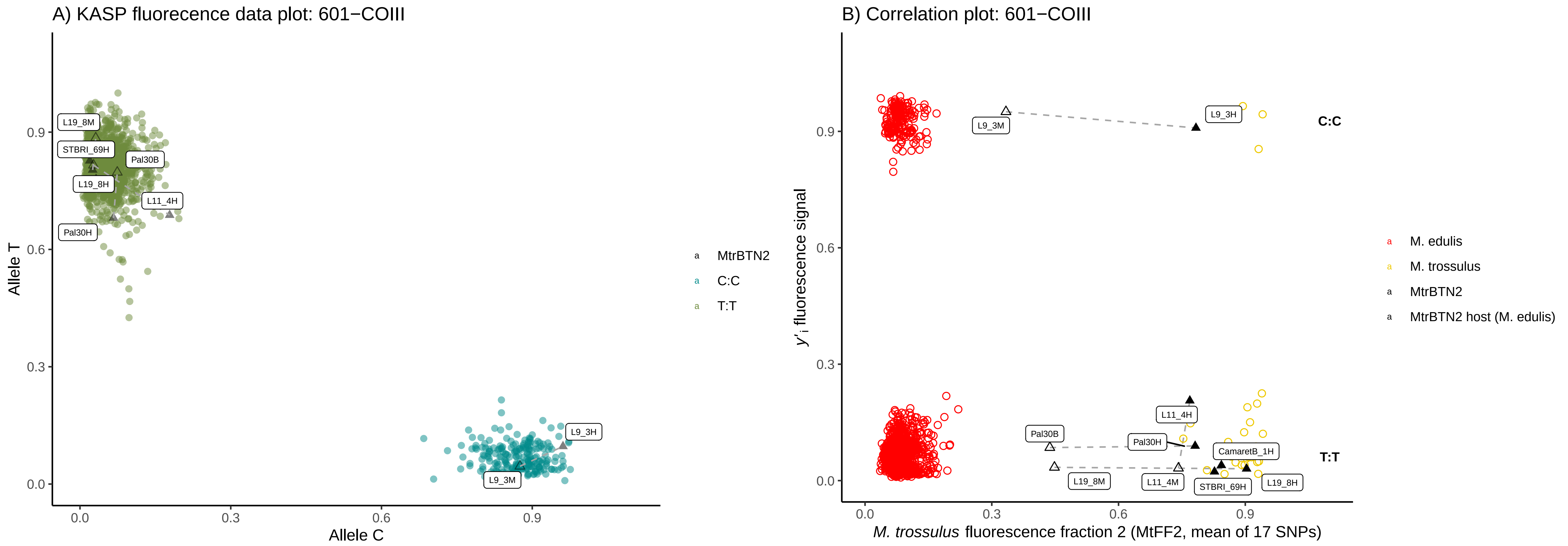
A) KASP fluorecence data plot: 601−COIII
Allele T
Allele C
0.0
0.3
0.6
0.9
L19_8M
STBRI_69H
Pal30B
L19_8H
L11_4H
Pal30H
L9_3H
L9_3M
MtrBTN2
C:C
T:T
B) Correlation plot: 601−COIII
y'i fluorescence signal
M. trossulus fluorescence fraction 2 (MtFF2, mean of 17 SNPs)
L9_3M
L9_3H
C:C
T:T
L11_4H
Pal30B
Pal30H
CamaretB_1H
L19_8M
L11_4M
STBRI_69H
L19_8H
M. edulis
M. trossulus
MtrBTN2
MtrBTN2 host (M. edulis)

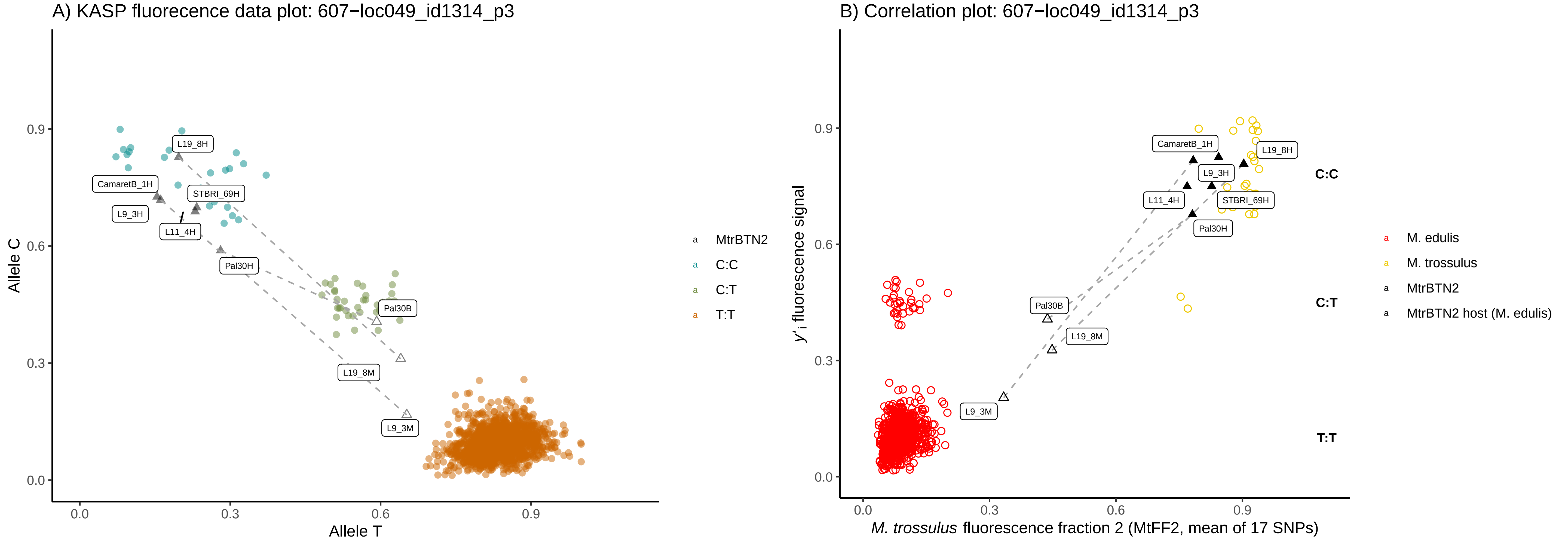
A) KASP fluorecence data plot: 607−loc049_id1314_p3
Allele C
Allele T
0.0
0.3
0.6
0.9
L19_8H
CamaretB_1H
STBRI_69H
L9_3H
L11_4H
Pal30H
Pal30B
L19_8M
L9_3M
MtrBTN2
C:C
C:T
T:T
B) Correlation plot: 607−loc049_id1314_p3
y'i fluorescence signal
M. trossulus fluorescence fraction 2 (MtFF2, mean of 17 SNPs)
CamaretB_1H
L19_8H
L9_3H
L11_4H
STBRI_69H
Pal30H
Pal30B
L19_8M
L9_3M
C:C
C:T
T:T
M. edulis
M. trossulus
MtrBTN2
MtrBTN2 host (M. edulis)

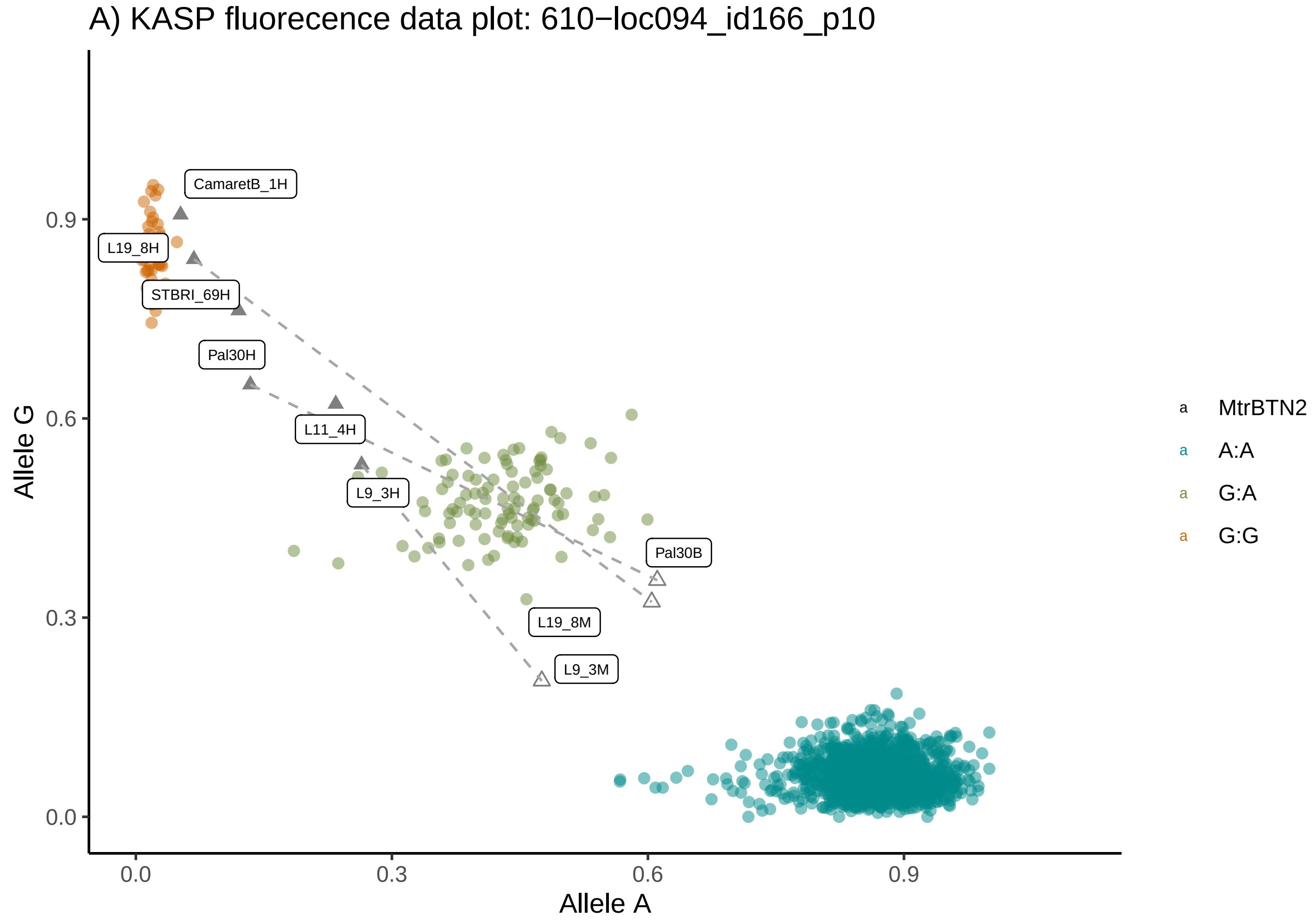
A) KASP fluorecence data plot: 610−loc094_id166_p10
Allele G
Allele A
0.0
0.3
0.6
0.9
CamaretB_1H
L19_8H
STBRI_69H
Pal30H
L11_4H
L9_3H
Pal30B
L19_8M
L9_3M
MtrBTN2
A:A
G:A
G:G

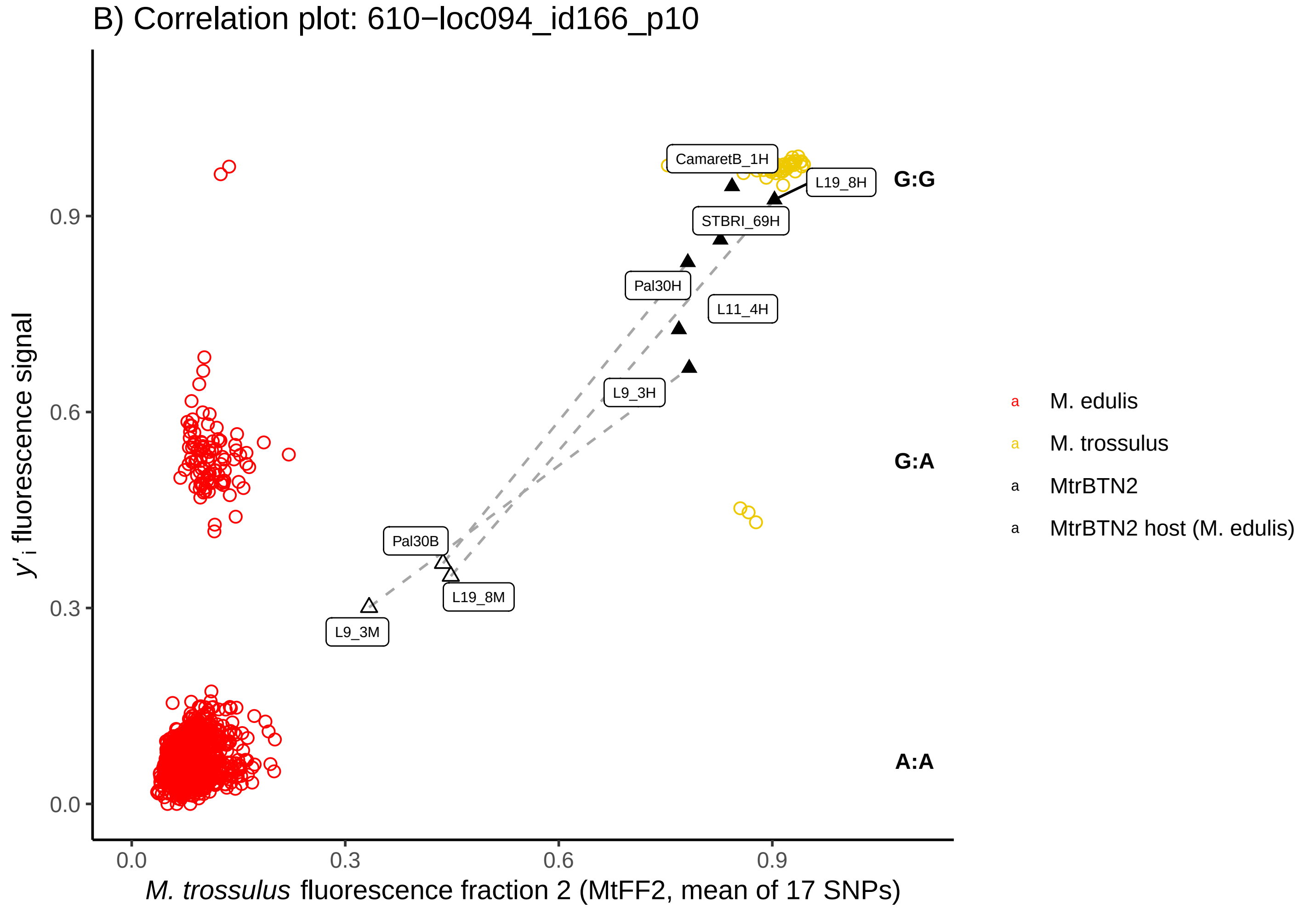
B) Correlation plot: 610−loc094_id166_p10
y'i fluorescence signal
M. trossulus fluorescence fraction 2 (MtFF2, mean of 17 SNPs)
0.0
0.3
0.6
0.9
CamaretB_1H
L19_8H
STBRI_69H
Pal30H
L11_4H
L9_3H
Pal30B
L19_8M
L9_3M
G:G
G:A
A:A
M. edulis
M. trossulus
MtrBTN2
MtrBTN2 host (M. edulis)

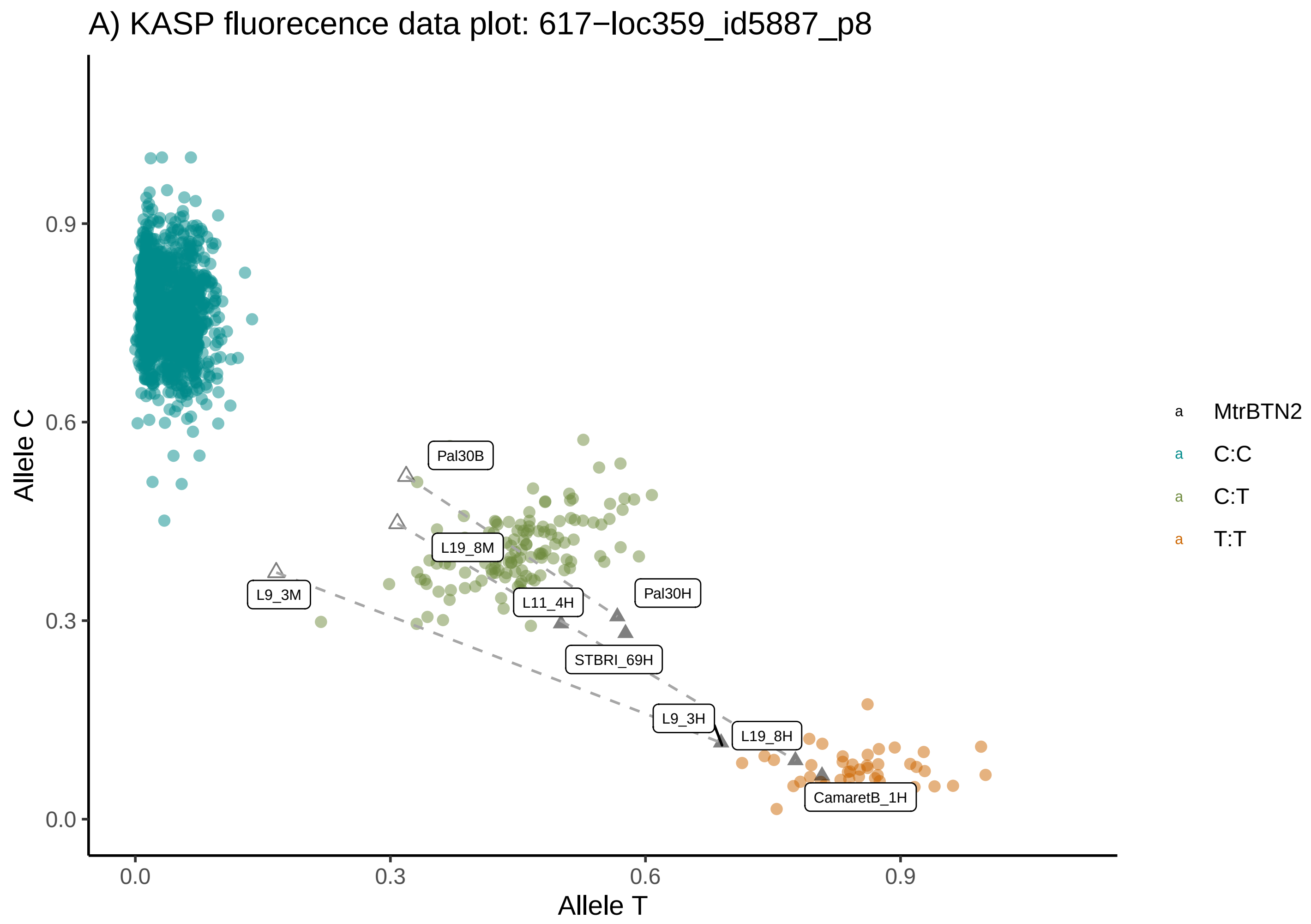
A) KASP fluorecence data plot: 617−loc359_id5887_p8
Allele C
Allele T
0.0
0.3
0.6
0.9
Pal30B
L19_8M
L9_3M
L11_4H
Pal30H
STBRI_69H
L9_3H
L19_8H
CamaretB_1H
MtrBTN2
C:C
C:T
T:T

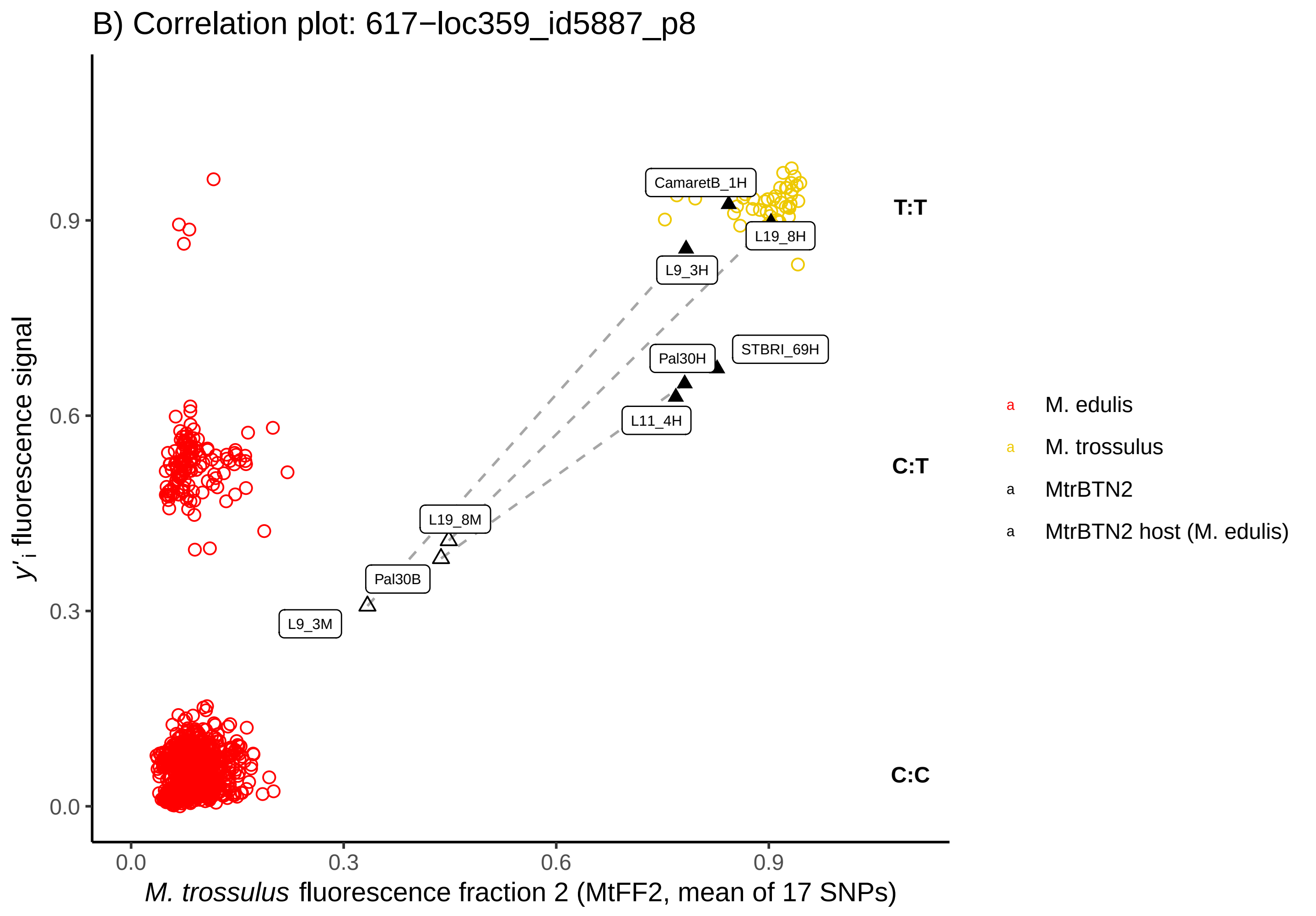
B) Correlation plot: 617−loc359_id5887_p8
y'i fluorescence signal
M. trossulus fluorescence fraction 2 (MtFF2, mean of 17 SNPs)
0.0
0.3
0.6
0.9
CamaretB_1H
L19_8H
L9_3H
Pal30H
STBRI_69H
L11_4H
L19_8M
Pal30B
L9_3M
T:T
C:T
C:C
M. edulis
M. trossulus
MtrBTN2
MtrBTN2 host (M. edulis)

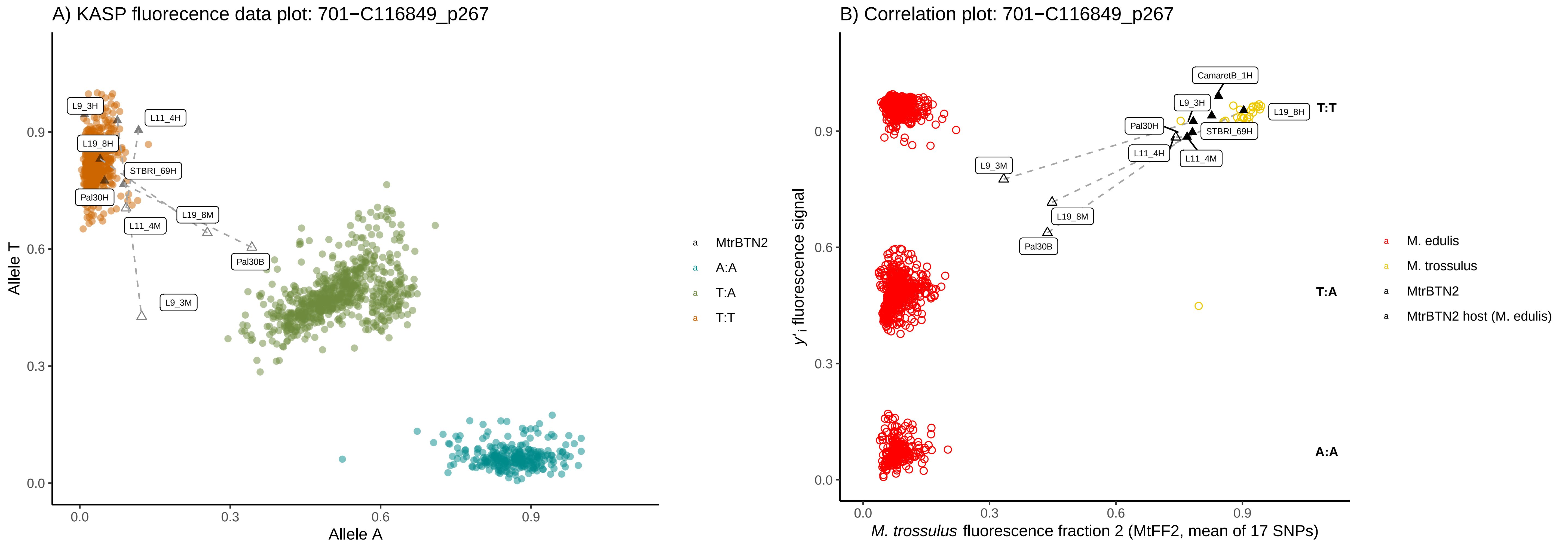
A) KASP fluorecence data plot: 701−C116849_p267
Allele T
Allele A
0.0
0.3
0.6
0.9
L9_3H
L11_4H
L19_8H
STBRI_69H
Pal30H
L19_8M
L11_4M
Pal30B
L9_3M
MtrBTN2
A:A
T:A
T:T
B) Correlation plot: 701−C116849_p267
y'i fluorescence signal
M. trossulus fluorescence fraction 2 (MtFF2, mean of 17 SNPs)
CamaretB_1H
L9_3H
L19_8H
Pal30H
STBRI_69H
L11_4H
L11_4M
L9_3M
L19_8M
Pal30B
T:T
T:A
A:A
M. edulis
M. trossulus
MtrBTN2
MtrBTN2 host (M. edulis)

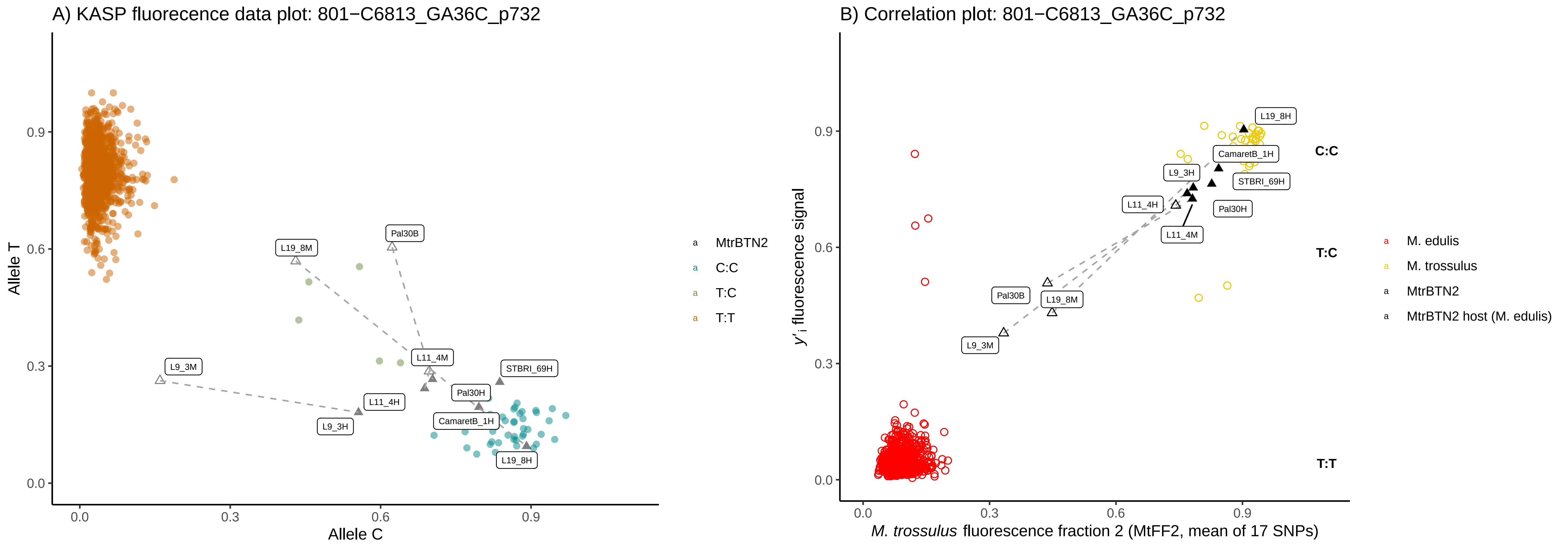

A) KASP fluorecence data plot: 801−C6813_GA36C_p732
Allele T
Allele C
0.0
0.3
0.6
0.9
L19_8M
Pal30B
L9_3M
L11_4M
STBRI_69H
L11_4H
Pal30H
L9_3H
CamaretB_1H
L19_8H
MtrBTN2
C:C
T:C
T:T
B) Correlation plot: 801−C6813_GA36C_p732
y'i fluorescence signal
M. trossulus fluorescence fraction 2 (MtFF2, mean of 17 SNPs)
C:C
T:C
T:T
M. edulis
M. trossulus
MtrBTN2
MtrBTN2 host (M. edulis)

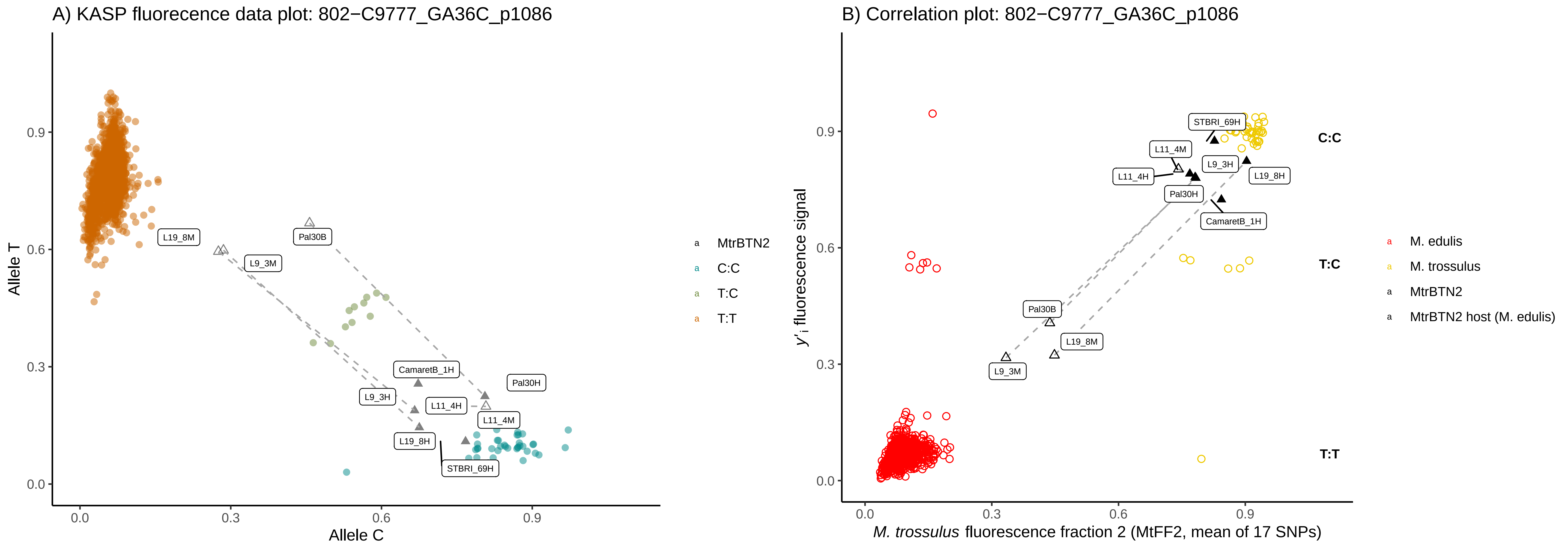
A) KASP fluorecence data plot: 802−C9777_GA36C_p1086
Allele T
Allele C
0.0
0.3
0.6
0.9
L19_8M
L9_3M
Pal30B
CamaretB_1H
Pal30H
L9_3H
L11_4H
L11_4M
L19_8H
STBRI_69H
MtrBTN2
C:C
T:C
T:T
B) Correlation plot: 802−C9777_GA36C_p1086
y'i fluorescence signal
M. trossulus fluorescence fraction 2 (MtFF2, mean of 17 SNPs)
STBRI_69H
L11_4M
L9_3H
L11_4H
L19_8H
Pal30H
CamaretB_1H
Pal30B
L19_8M
L9_3M
C:C
T:C
T:T
M. edulis
M. trossulus
MtrBTN2
MtrBTN2 host (M. edulis)

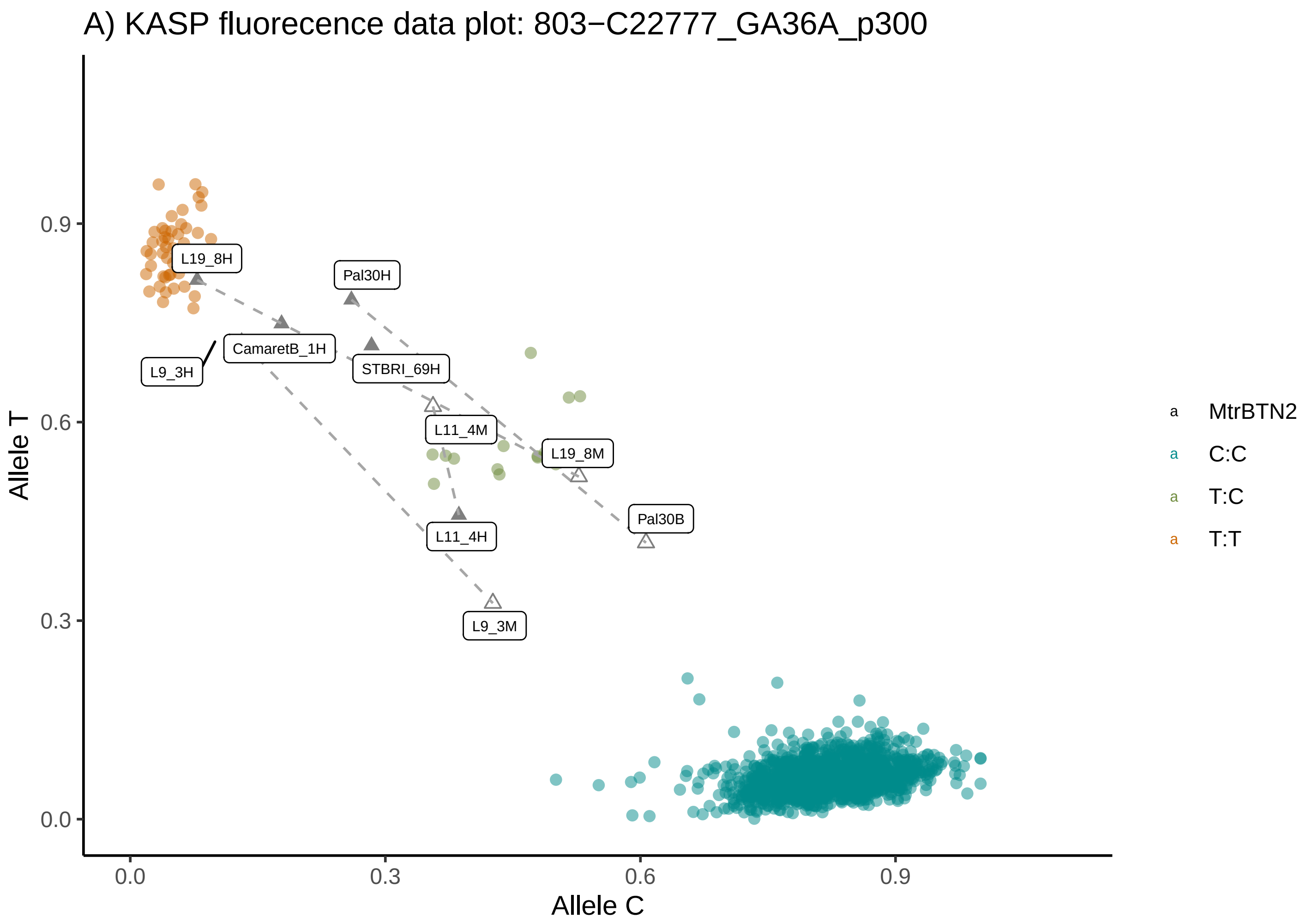
A) KASP fluorecence data plot: 803−C22777_GA36A_p300
Allele T
Allele C
0.0
0.3
0.6
0.9
L19_8H
Pal30H
CamaretB_1H
L9_3H
STBRI_69H
L11_4M
L19_8M
Pal30B
L11_4H
L9_3M
MtrBTN2
C:C
T:C
T:T

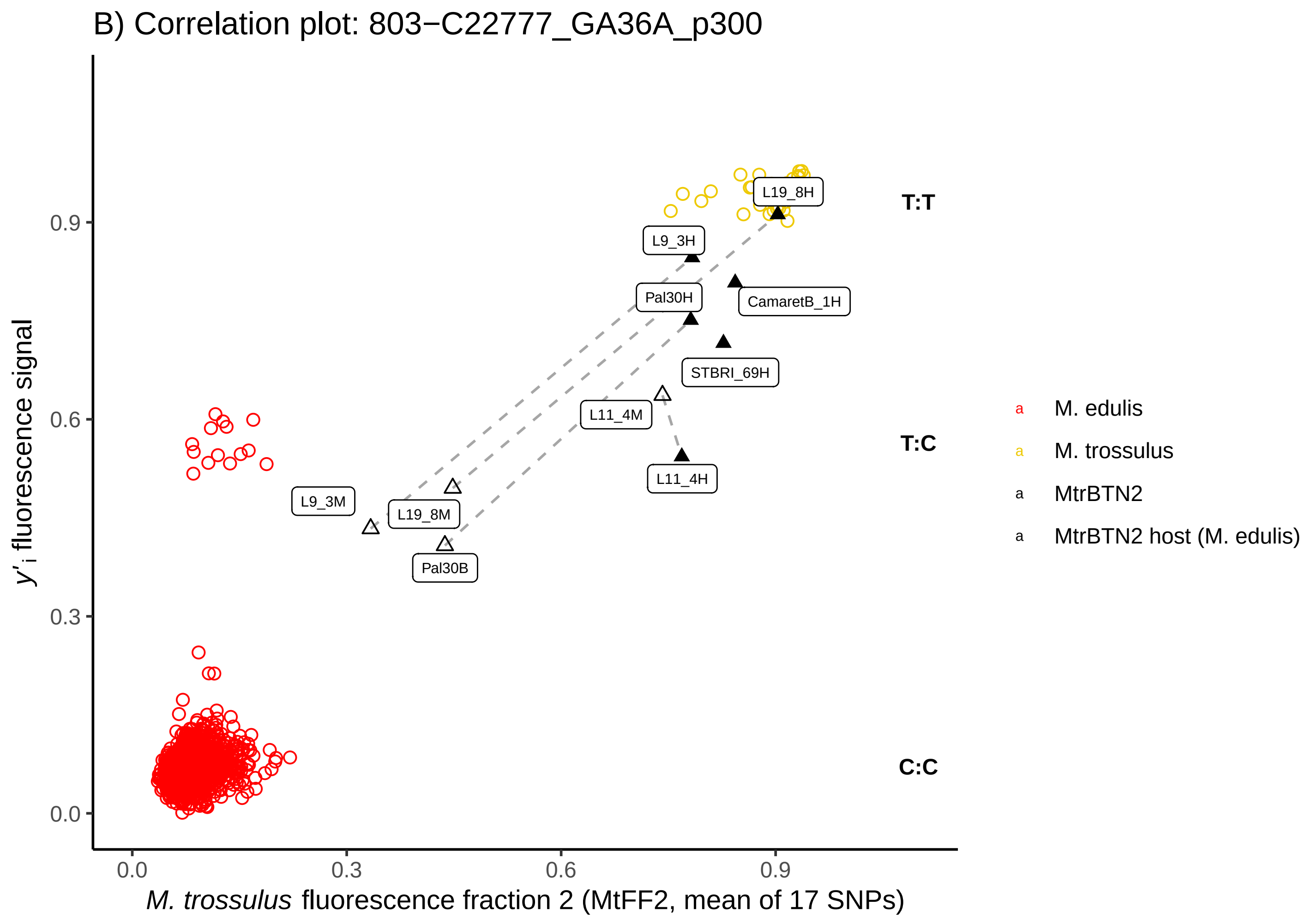
B) Correlation plot: 803−C22777_GA36A_p300
y'i fluorescence signal
M. trossulus fluorescence fraction 2 (MtFF2, mean of 17 SNPs)
0.0
0.3
0.6
0.9
L19_8H
L9_3H
Pal30H
CamaretB_1H
STBRI_69H
L11_4M
L11_4H
L9_3M
L19_8M
Pal30B
T:T
T:C
C:C
M. edulis
M. trossulus
MtrBTN2
MtrBTN2 host (M. edulis)

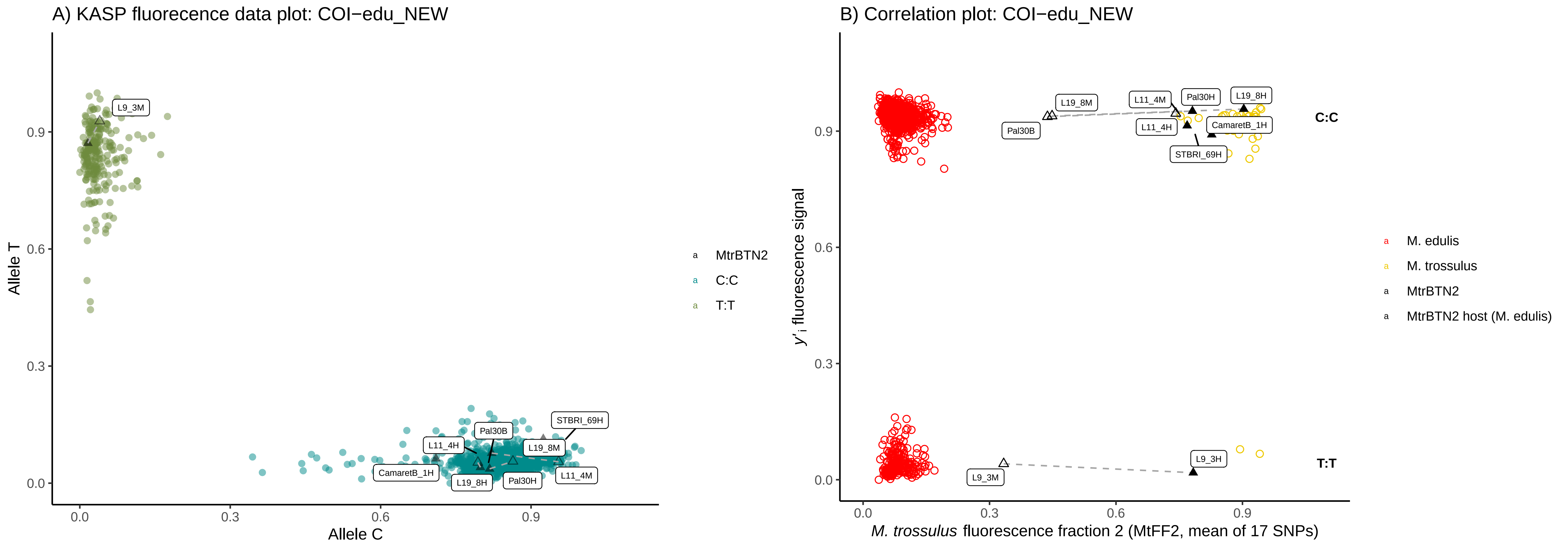
A) KASP fluorecence data plot: COI−edu_NEW
B) Correlation plot: COI−edu_NEW
Allele T
Allele C
0.0
0.3
0.6
0.9
L9_3M
L11_4H
Pal30B
STBRI_69H
L19_8M
CamaretB_1H
L19_8H
Pal30H
L11_4M
MtrBTN2
C:C
T:T
y'i fluorescence signal
M. trossulus fluorescence fraction 2 (MtFF2, mean of 17 SNPs)
L19_8M
L11_4M
Pal30H
L19_8H
Pal30B
L11_4H
CamaretB_1H
STBRI_69H
L9_3H
L9_3M
C:C
T:T
M. edulis
M. trossulus
MtrBTN2
MtrBTN2 host (M. edulis)
